# Identifying Site-specific Superoxide and Hydrogen Peroxide Production Rates from the Mitochondrial Electron Transport System Using a Computational Strategy

**DOI:** 10.1101/2021.07.01.450791

**Authors:** Quynh V. Duong, Yan Levitsky, Maria J. Dessinger, Jason N. Bazil

## Abstract

Mitochondrial reactive oxygen species (ROS) play important roles in cellular signaling; however, certain pathological conditions such as ischemia/reperfusion (I/R) injury disrupt ROS homeostasis and contribute to cell death. A major impediment to developing therapeutic measures against oxidative stress induced cellular damage is the lack of a quantitative framework to identify the specific sources and regulatory mechanisms of mitochondrial ROS production. We developed a thermodynamically consistent, mass-and-charge balanced, kinetic model of mitochondrial ROS homeostasis focused on redox sites of electron transport chain complexes I, II, and III. The model was calibrated and validated using comprehensive data sets relevant to ROS homeostasis. The model predicts that complex I ROS production dominates other sources under conditions favoring a high membrane potential with elevated NADH and QH_2_ levels. In general, complex I contributes to significant levels of ROS production under pathological conditions, while complexes II and III are responsible for basal levels of ROS production, especially when QH_2_ levels are elevated. The model also reveals that hydrogen peroxide production by complex I underlies the non-linear relationship between ROS emission and O_2_ at low O_2_ concentrations. Lastly, the model highlights the need to quantify scavenging system activity under different conditions to establish a complete picture of mitochondrial ROS homeostasis. In summary, we describe the individual contributions of the ETS complex redox sites to total ROS emission in mitochondria respiring under various combinations of NADH- and Q-linked respiratory fuels under varying work rates.

## I. Introduction

Reactive oxygen species (ROS) were once considered by-products of cellular respiration but are now recognized as important signaling molecules ^1–4^. Under pathological conditions such as ischemia/reperfusion (I/R) injury, elevated ROS levels contribute to cell death ^5^. This type of injury occurs after a period of partial or complete loss of tissue blood flow (ischemia) followed by the restoration of normal flow (reperfusion). Although reperfusion is necessary to salvage ischemic tissue, it leads to significant tissue damage in a ROS-dependent manner ^5^. This unfortunate and common event is also associated with clinical situations such as organ transplantation ^6^ and hypovolemic shock ^7^. Though I/R injury can affect any organ, metabolically active tissue such as brain and myocardium are the most sensitive to this injury. Despite enormous efforts to ameliorate the detrimental effects of reperfusion-dependent oxidative stress, pharmacological interventions have not produced effective clinical treatment options ^5,8,9^. A more complete understanding of redox site-specific ROS emission in I/R injury may lead to the design of efficient pharmacologic interventions.

In most mammalian cells, mitochondria are the primary source of ROS ^10–12^. As such, mitochondrial ROS homeostasis is essential to maintaining mitochondrial and cellular physiology. In essence, ROS production and elimination act as countering forces which determine the mitochondrial net ROS emission. The mechanisms underlying ROS production have been reviewed extensively elsewhere ^13,14^, and only key details are provided here. The electron transport system (ETS) complexes I and III are accepted as the major mitochondrial ROS producers; however, their respective contributions to total ROS emission vary according to the bioenergetic state of the organelle ^10^. Matrix and non-ETS inner membrane proteins can also produce ROS, but their contribution is limited ^15–17^. Under physiological conditions, electron flux through the ETS favors forward electron transport (FET). In this operating mode, pyruvate (P) from glucose metabolism is oxidized through the Krebs cycle to generate NADH, which enters the ETS at complex I. When succinate (S) is oxidized, electrons enter the ETS at complex II. Despite the different entry points, electrons converge at the quinone (Q) pool before they are passed onto cytochrome c and ultimately molecular oxygen (O_2_). This is the normal flow of electrons during energy transduction. During certain pathological states such as anoxia, the ETS circuitry is rewired. Here, the forward electron flow through complex I is supported by reversal of complex II while the activities of complexes III and IV dramatically fall near zero ^18,19^. Reorganization of electron flux causes succinate and QH_2_ to accumulate as fumarate reduction at complex II replaces O_2_ reduction at complex IV. Upon reperfusion, electron flow normalizes but the reduced Q pool leads to thermodynamic and kinetic processes favoring ETS operation in reverse electron transport (RET) mode. In this mode, complex I enters a near equilibrium state with nearly all its redox centers fully reduced, producing ROS at significantly elevated rates. Thus, ROS production is enhanced under RET compared to FET with the former playing a significant role in I/R-induced oxidative stress.

The majority of O_2_ is fully reduced to water at complex IV; however, one- or two-electron reduction of O_2_ by an ETS redox center upstream of complex IV produces superoxide or hydrogen peroxide, respectively. In complex I, the redox centers include a flavin mononucleotide (FMN) at the NADH oxidase site (site I_F_) ^20^, a semiquinone (SQ) at the Q reductase site (site I_Q_) ^21^ and a chain of iron-sulfur (Fe-S) clusters that rapidly relay electrons between the two ^22,23^. Likewise, complex II harbors a flavin adenine dinucleotide (FAD) at the succinate oxidase site (II_F_), a Q reductase site (II_Q_), and a chain of Fe-S clusters ^24,25^. The mammalian complex III redox centers include cytochrome c_1_, a high- and low-potential b type heme, the Reiske iron-sulfur protein (ISP) and two quinone binding sites (Q_N_ and Q_P_) ^26^. Superoxide arising from complex III is generally accepted to originate from the SQ at the quinone binding site proximal to the intermembrane space (Q_p_) ^27^.

The topic of site-specific ROS production has attracted experimental researchers for many years. Both pharmacological and genetic approaches have been utilized to dissect the contribution of redox centers to total mitochondrial ROS. While complexes I and III are the major ROS producers, the contribution of specific redox centers at physiologically relevant metabolic states remains elusive. For example, both site I_F_ ^28–30^ and I_Q_ ^31^ are proposed as major ROS-producing sites of complex I. Complex II was widely accepted as a negligible source of ETS ROS ^32^. However, several more recent studies report that complex II produces significant amounts of ROS under appropriate conditions ^33,34^. Such discrepancies in experimental data arise from the caveats inherent to experimental studies and have contributed to unsuccessful efforts to target oxidative stress in clinical settings. In particular, the use of chemical inhibitors and genetic models inevitably alter the native electron distribution in unpredictable patterns, leading to epiphenomena and conflicting experimental results. Differences in experimental conditions including species, tissues, developmental stage, etc. introduce additional confounding factors that are often neglected and make comparing results from different studies difficult. Lastly, the pursuit is further impeded by the lack of a robust method to distinguish species-specific ROS without interfering with other mitochondrial and extra-mitochondrial processes.

Computational modeling is a useful platform to address these challenges. A computational model serves as a quantitative framework to analyze mitochondrial bioenergetic data from a single, unified perspective. Several models of mitochondria exist at varying levels of complexity, focus, and approach ^35–42^. Some investigate the kinetics underlying ROS production by individual ETS complexes ^38–40^ while others integrate mitochondrial ROS production and elimination in a substrate-specific context ^41,42^. However, to our knowledge, none consistently reproduces a wide range of experimental data using a unified, coherent, and consistent framework. Thus, the origin of superoxide and hydrogen peroxide from the ETS remains elusive from a computational or experimental perspective.

We herein developed, analyzed, and corroborated a comprehensive model that simulates mitochondrial ROS homeostasis under a range of bioenergetic conditions. This model is focused on ROS from the ETS and is referred to as the ETS-ROS model. Modules relevant to ROS originating from the ETS were individually developed, parameterized, and corroborated against a variety of data sets ^38–40^. The model includes the primary ROS producers in mitochondria with their biochemical reactions constrained thermodynamically and balanced with respect to mass and charge. These modules were then incorporated into a single integrated framework, producing a model that is coherent and operates within biophysical constraints using a single, consistent set of parameters. The ETS-ROS model reproduces experimental data of not only net ROS production but also other mitochondrial bioenergetic variables. Using this approach, we have identified the regulation of ROS production by substrate metabolism *via* alterations to the mitochondrial membrane potential along with the NAD and Q pool redox states. The model also reveals that the scavenging system is saturable with a K_M_ for H_2_O_2_ in the nM range. Therefore, under pathological conditions such as I/R injury, cellular oxidative stress is a result of ROS production overwhelming ROS elimination.

## II. Materials and Methods

### 1. Computational model

#### General approach to modeling and processes included

Our modeling approach is modular and contains enzyme kinetic and ROS production models of complexes I ^38^, II ^39^ and III ^40^ previously published by our group. These models are biophysically detailed and thermodynamically consistent. They were individually calibrated and corroborated against a wide range of kinetic data sets. In this ETS-ROS model, we opted to explicitly model part of the Krebs cycle while lumping other parts. The explicit parts of the model consist of the enzymatic reactions that support succinate, fumarate, and malate transport and oxidation. Pyruvate transport is explicitly modeled; however, pyruvate dehydrogenase and the upper branch of the Krebs cycle (citrate synthase through succinyl-CoA synthetase) were modeled as a lumped reaction. This lumped reaction was sufficient to explain the experimental data. The detailed reactions for the lumped portion of the model are excluded to stay faithful to the parsimonious principle. In addition, our in-house experimental data reveal a moderate level of malic enzyme (ME) activity in mitochondria isolated from guinea pig ventricular cardiomyocytes (Fig. S1). Several other studies also support the expression of ME in cardiac tissue of guinea pigs ^43,44^, and we found this reaction necessary to fit the model to the calibration data sets (Table 2). A summary of processes that are explicitly modeled are shown in Figure 1. Notations of the redox centers included in the text are described in Table 1.

**Table 1.**
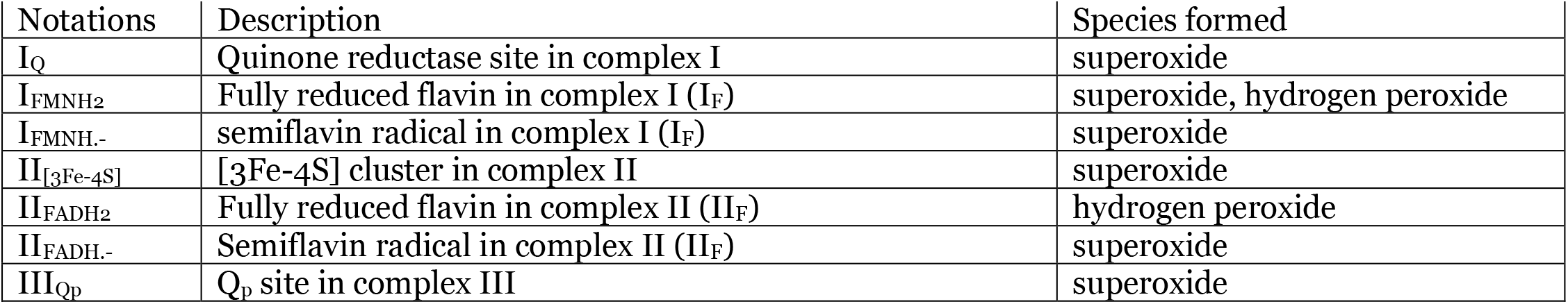
Description of the ETS redox centers included in the model

**Table 2.** Summary of data used in this study

| Modelling Stage | Data Sets | References |
| --- | --- | --- |
| Calibration | $J_{O_2} - P/M, S/R$ | This study |
| | $J_{O_2} - S$ | Duong *et al.* 45 |
| | $J_{H_2O_2} - P/M, S/R$ | This study |
| | $J_{H_2O_2} - S$ | Duong *et al.* 45 |
| | $J_{H_2O_2}$ v.s. $[O_2]$ | This study |
| | $J_{O_2} - L-M$ | Duong *et al.* 45 |
| Corroboration | $P/M/S - J_{O_2}, J_{H_2O_2}$ | This study |
| | $NADH - P/M, S/R, S, P/M/S$ | This study |

L-M: L-malate; P: pyruvate; S: succinate; R: rotenone

**Figure 1.**
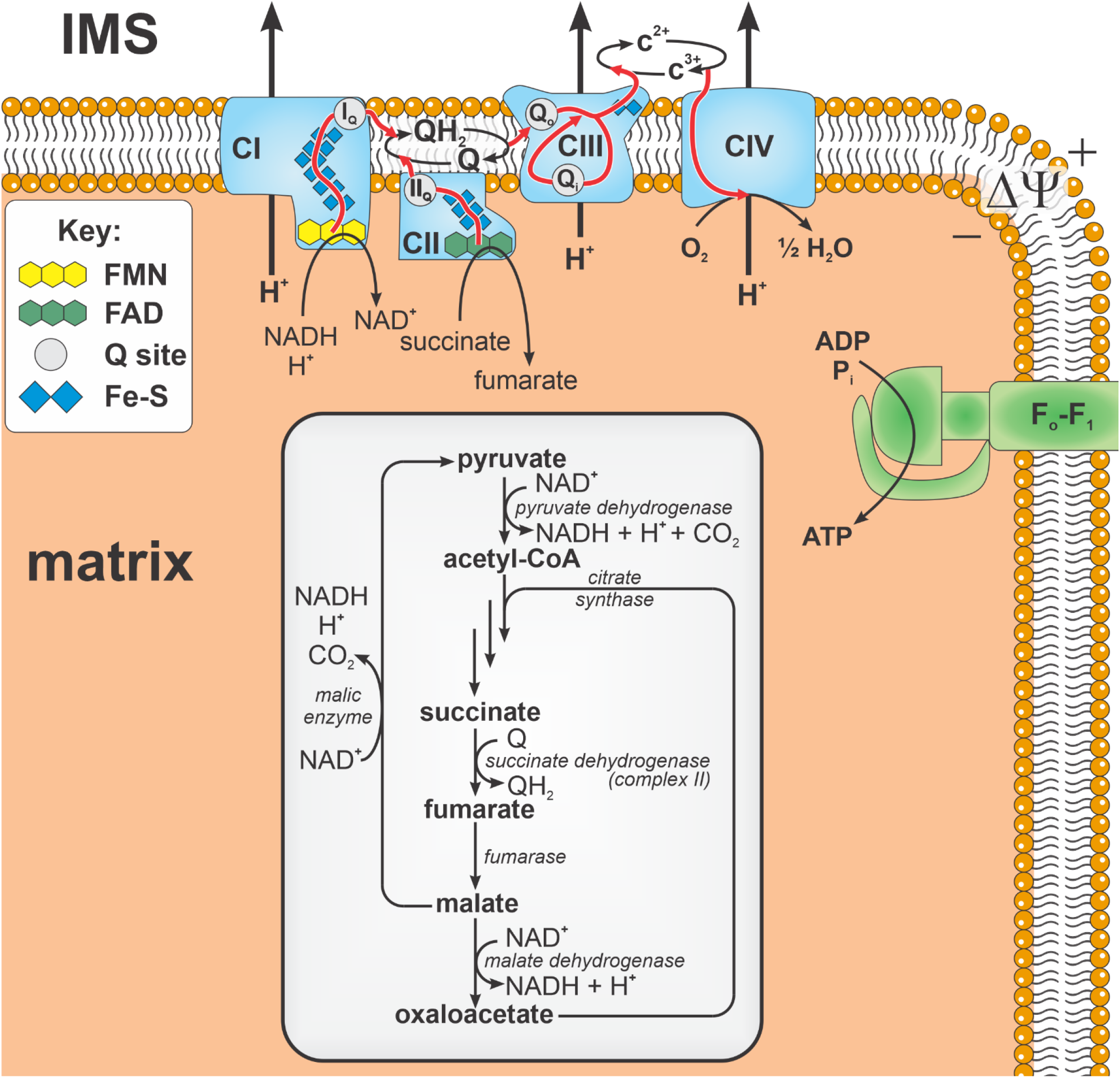
Schematic of modeled biochemical reactions and phenomena. The model consists of a partially lumped Kreb’s cycle with non-lumped enzymatic reactions shown in the light gray box. Black arrows denote chemical reactions. Red arrows denote electron flux through redox centers. FMN = flavin mononucleotide, FAD = flavin adenine dinucleotide, Q = ubiquinone, QH_2_ = ubiquinol, Fe-S = iron sulfur clusters, IMS = intermembrane space, CI – CIV = complexes I – IV, c = cytochrome *c*, Q site = quinol/quinone binding site.

#### Model simulations

The model was numerically simulated using MATLAB (R2020b). The parameter optimization was performed on a Dell desktop PC (64-bit operating system and x64-based processor Intel® core™ i7-7700 CPU @3.60GHz and 16 GB RAM) using the Parallel Computing Toolbox. A parallelized simulated annealing algorithm was first used to globally search for feasible parameters which were then refined using a local, gradient-based optimization algorithm. When fitting to the calibration data, the difference between model output and data were normalized with the respetive uncertainty in the data and minimzed using the standard least squares method.

#### Data sets

The intricate relationship between substrate utilization and ROS homeostasis necessitates that model calibration includes experimental data describing both NADH- and QH_2_-linked substrate supported respiration and ROS production. The calibration data set includes the oxygen consumption rates (J_O2_, nmol/mg/min), net hydrogen peroxide emission rates (J_H2O2_, pmol/mg/min) and NADH reduction (%) under NADH- and QH_2_-linked supported respiration in both FET and RET modes. Forward electron transport occurs when respiration is supported by pyruvate/L-malate (P/M) and S in the presence of rotenone (S/R). Reverse electron transport occurs under QH_2_-linked supported respiration when rotenone is absent (S). The calibrated model is subsequently challenged against novel data sets, those which were not used for model calibration. The first corroboration data set includes the J_O2_ and J_H2O2_ data obtained when respiration is supported by both NADH- and QH_2_-linked substrates (P/M/S). The second data set recapitulates the monotonic relationship of J_H2O2_ on oxygen concentrations in FET mode. The list of all the data sets utilized in the construction, calibration and corroboration of the model is summarized in Table 2.

The majority of the experimental data used for model calibration were from a previous publication of ours ^45^. In this study, we measured the oxygen consumption rate (J_O2_, nmol mg^−1^ min^−1^) and net hydrogen peroxide emission rate (J_H2O2_, pmol mg^−1^ min^−1^) of mitochondria respiring on P/M, S and S/R. Briefly, all experiments were performed using a standard KCl-based mitochondrial suspension buffer containing 130 mM KCl, 5 mM K_2_HPO_4_, 20 mM MOPS, 1 mM MgCl_2_, 1 mM EGTA and 0.1% w/v BSA (pH of 7.1 at 37 °C). For general methods and protocols regarding mitochondrial isolation and bioenergetic experiments, see our prior works ^45,46^. When used, the final concentrations of mitochondria, substrates, and inhibitors were 0.1 mg/mL mitochondria, 5 mM pyruvate (P), 1 mM L-malate (M), 10 mM succinate (S), 1 μM rotenone (R), and 500 μM ADP. The data were analyzed using the software MATLAB®.

#### Model parameters

The fixed model parameters are obtained from thermodynamic and kinetic data obtained from prior work ^18,38–40^. The adjustable parameter set consists of 14 parameters related to both ROS kinetics and substrate metabolism. We chose the smallest set of adjustable parameters necessary to simulate the calibration data sets. The ROS kinetic parameters are related to mitochondrial production and clearance of superoxide and hydrogen peroxide. The capacity of mitochondria to remove hydrogen peroxide is interchangeably referred to as the scavenging capacity in this paper. The scavenging system is modeled in a Michaelis-Menten like fashion - saturable and dependent on the hydrogen peroxide concentration. The ETS parameters include the enzyme contents for complex I, II, and III, the estimation of the semiquinone stability constant in complex III and the apparent rotenone binding constant to complex I. The remaining adjustable parameters are concerned with metabolite transport and substrate utilization (Table 3). Normalized local sensitivity coefficients are computed to inform the contribution of each adjustable parameter to the model output in a similar manner previously described ^38–40^. These values are calculated from model outputs coincident with the data and associated dynamics. The sensitivity analysis reveals that the complex I content is the most sensitive parameter followed by complex III content and the H_2_O_2_ affinity of the matrix scavenging system. These results are not unexpected considering the tight association between respiratory state and ROS emission.

**Table 3.** Model adjustable parameters

| Parameter | Definition | Value | Unit | Sensitivity | Rank |
| --- | --- | --- | --- | --- | --- |
| $Etot_{CI}$ | Complex I content | 0.167 | nmol mg<sup>-1</sup> | 0.118 | 1 |
| $Etot_{CII}$ | Complex II content | 0.725 | nmol mg<sup>-1</sup> | 0.038 | 8 |
| $Etot_{CIII}$ | Complex III content | 0.372 | nmol mg<sup>-1</sup> | 0.093 | 3 |
| $V_{max}^{DCC}$ | Dicarboxylate carrier max transport rate | 5450 | nmol mg<sup>-1</sup> min<sup>-1</sup> | 0.007 | 12 |
| $V_{max}^{FH}$ | Fumarate hydratase max rate | 6860 | nmol mg<sup>-1</sup> min<sup>-1</sup> | 0.028 | 11 |
| $X_{DH}$ | Dehydrogenase activity | 7210 | nmol mg<sup>-1</sup> min<sup>-1</sup> | 0.033 | 10 |
| $K_{DH}$ | Phosphoennergetics feedback constant for matrix dehydrogenase activity | 0.124 | mM | 0.002 | 13 |
| $r_{DH}$ | NADH/NAD<sup>+</sup> feedback constant for matrix dehydrogenase activity | 7.60x10<sup>-3</sup> | - | 0.001 | 14 |
| $k_{fME}$ | Malic enzyme rate constant | 3.02x10<sup>6</sup> | nmol mg<sup>-1</sup> min<sup>-1</sup> M<sup>-2</sup> | 0.043 | 7 |
| $V_{max}^{scavenge}$ | Maximum matrix scavenging rate | 31.6 | nmol mg<sup>-1</sup> min<sup>-1</sup> | 0.062 | 5 |
| $K_M^{scavenge}$ | $[H_2O_2]$ at half maximum scavenging rate | 106 | nM | 0.099 | 2 |
| $k_{Em,H_2O_2}$ | $H_2O_2$ permeability rate constant | 5.50x10<sup>5</sup> | min<sup>-1</sup> | 0.062 | 4 |
| $K_{CIII}^{SQ}$ | Complex III semiquinone stability constant | 1.40x10<sup>-12</sup> | - | 0.060 | 6 |
| $K_{CI}^{rot}$ | Complex I rotenone binding constant | 1.48 | pM | 0.037 | 9 |

### 2. Experimental Methods

#### Oxygen Consumption and Net Hydrogen Peroxide Emission Rates

The Oroboros Oxygraph (O2k) was used to simultaneously measure J_O2_ (nmol mg^−1^ min^−1^) and J_H2O2_ (pmol mg^−1^ min^−1^), described in detail in our previous publications ^45^. Briefly, J_H2O2_ values were measured using the Amplex UltraRed/Horseradish Peroxidase/Superoxide Dismutase assay. The reduction of hydrogen peroxide is coupled with the oxidation of Amplex UltraRed to resorufin. Resorufin fluorescence is converted to hydrogen peroxide concentration by using a hydrogen peroxide calibration curve obtained each experiment day from freshly prepared hydrogen peroxide standards. A post-hoc analysis of our previous J_H2O2_ data sets using S as the substrate reveals that S-dependent J_H2O2_ is highly sensitive to the respiratory control ratio (RCR). Higher J_H2O2_ values correspond to RCR (energized with P/M and in the presence of 4 μM Ca^2+^) greater than 18. Thus, we repeated J_H2O2_ measurements using mitochondria with an RCR above 18. Mitochondria were energized with L-M, S or the P/M/S combination (leak state). An ADP bolus was added to stimulate oxidative phosphorylation (oxphos state) after 5 minutes under L-M supported respiration, 2.5 minutes under S- and 2 minutes under P/M/S-supported respiration. These time periods were chosen to allow measurement of steady-state J_O2_ while maintaining adequate oxygen supply for oxidative phosphorylation turnover.

#### NADH Fluorescence Measurement

The fluorescence of NADH was monitored on an Olis DM-245 spectrofluorometer. Fluorescence was monitored with λ_excitation_ = 355 nm (8 nm bandpass filter) and λ _emission_ = 450 nm (13 nm bandpass filter). Baseline fluorescence was monitored for 1 minute followed by mitochondria addition. Substrates were added after 5 minutes of equilibration. A bolus of ADP stimulated oxidative phosphorylation. Fluorescence minima were obtained by uncoupling mitochondria with FCCP to produce maximally oxidized nicotinamide pools. Maximal fluorescence was obtained in the presence of rotenone to produce maximally reduced nicotinamide pools.

## III. Results

### 1. Model Calibration

The experimental fidelity of the ETS-ROS model is critically related to its ability to predict the *in vivo* response of the ETS under a wide variety of physiological and pathophysiological scenarios. Including both FET and RET modes of ROS production data into model calibration was essential to identify the parameters associated with site-specific ROS fluxes under these diverse experimental conditions. In the absence of exogenous ADP, the substrate combinations of P/M and S/R induce FET whereas RET was favored with S as the sole substrate. We used the P/M/S data, which consist of both FET and RET processes, to corroborate the model.

#### Oxygen consumption and net hydrogen peroxide emission rates

Outputs from the calibrated model were compared to experimentally determined J_O2_ (nmol/mg/min) and J_H2O2_ (pmol/mg/min), as shown in Fig. 2 and summarized in Table 4. In general, the model captures the dynamic profile of O_2_ consumption as well as H_2_O_2_ emission rates in a range of energy demand conditions. In the leak state, respiration rates are low and H_2_O_2_ emission rates are high. This is general for all substrate conditions and is due to membrane polarization and highly reduced mitochondrial redox couples (e.g., high NADH/NAD^+^). When succinate is present, the QH_2_/Q ratio is large and H_2_O_2_ emission rates reach even higher values. The highly reduced redox centers in the ETS proteins thermodynamically and kinetically favor elevated ROS emission rates.

**Figure 2.**
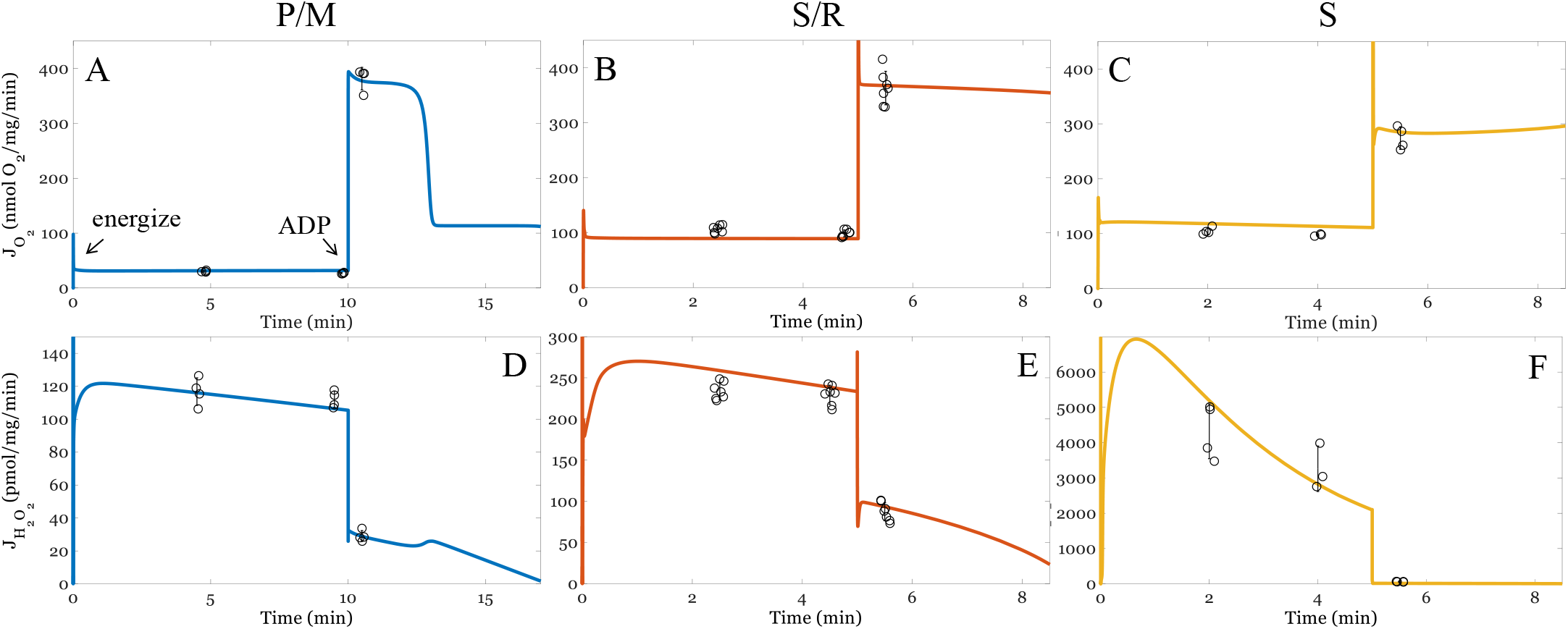
Model simulations of oxygen consumption rates (J_O2_, nmol/mg/min, *A-C*) and net hydrogen peroxide emission rates (J_H2O2_, pmol/mg/min, *D-F*) in forward electron transport (FET) and reverse electron transport (RET) modes. FET occurs when the substrates are P/M (A and D, blue) or S/R (B and E, orange). RET occurs when S is the substrate and rotenone is absent (C and F, yellow). The model outputs are represented by the solid lines. Individual data points are represented by the open black circles; error bars represent the standard deviations. Experimental data are obtained using mitochondria isolated from the ventricular cardiomyocytes of guinea pigs, as described in our previous work ^45^. As shown, the model outputs are within the experimental ranges for both J_O2_ and J_H2O2_ in both FET and RET modes.

**Table 4.** Experimental and model J_O2_ (nmol/mg/min) and J_H2O2_ (pmol/mg/min)

| Substrates | Respiratory state | Experimental $J_{O_2}$ | Model $J_{O_2}$ | Experimental $J_{H_2O_2}$ | Model $J_{H_2O_2}$ |
| --- | --- | --- | --- | --- | --- |
| Pyruvate/L-malate (P/M) | 5-minute leak | 30.3 ± 1.4 | 31 | 117 ± 8 | 116 |
| | 10-minute leak | 27 ± 1.1 | 31 | 112 ± 5 | 106 |
| | oxphos | 381.4 ± 20.6 | 377 | 29 ± 4 | 29 |
| Succinate/Rotenone (S/R) | 2.5-minute leak | 106.4 ± 6.7 | 89 | 234 ± 10 | 259 |
| | 5.0-minute leak | 98.5 ± 6.3 | 89 | 229 ± 12 | 236 |
| | oxphos | 363.3 ± 30.6 | 367 | 87 ± 11 | 91 |
| Succinate (S) | 2.5-minute leak | 106.9 ± 8.6 | 118 | 4,300 ± 775 | 4,995 |
| | 5.0-minute leak | 103.6 ± 4.5 | 113 | 3,700 ± 991 | 2,707 |
| | oxphos | 265.4 ± 36.3 | 284 | 61 ± 7 | 14 |

When ADP is added, mitochondria enter the oxphos state and dramatically increase their respiratory rate. In this state, redox couples become more oxidized and thus produce ROS at a lower rate compared to leak state. This relationship is also consistent across substrate conditions. Relative to the S/R condition, the absence of rotenone for the S condition did not affect leak-state J_O2_ but slightly depressed oxphos-state J_O2_. This results from NAD pool oxidation leading to oxaloacetate accumulation and complex II inhibition due to malate dehydrogenase turnover ^45,47^. Importantly, the H_2_O_2_ emission rate is dependent on O_2_ levels (will be discussed in detail below and briefly here). In these simulations, we modeled the buffer O_2_ dynamics as we will show later. Our initial buffer O_2_ concentration was 188 μM in a 2 mL volume of uniformly dispersed isolated mitochondria. In our conditions, a respiratory rate of 100 nmol O_2_/mg/min corresponds a rate of 10 μM/min which will deplete all the O_2_ in 18.8 min. These simulation results show the model not only reproduces the respiratory dynamics associated with various substrates and mitochondrial work rates, but it also simulates free radical emission rates under both FET and RET operating modes.

#### NADH reduction states

An essential bioenergetic variable associated with H_2_O_2_ emission is the NAD redox state. The changes to the NAD^+^ pool redox state vary with substrate delivery, ETS flux, and phosphorylation potential, reflecting the overall redox balance of the mitochondrial matrix. As shown in Figure 3, the model is capable of faithfully reproducing experimentally observed NADH dynamics of mitochondria respiring in the leak and oxphos states with P/M or S/R as substrates. Succinate-supported respiration with rotenone yielded a more reduced NADH pool in both leak and oxphos states compared to the P/M substrates. With P/M as the substrates, turnover of NADH at complex I favors a more oxidized NADH pool in leak (80%) and oxphos (30%) states relative to S/R (87.5% and 90%, respectively). The rapid NADH reduction dynamics seen with S/R are a result of residual malic enzyme and site I_Q_ activity. We used 1 μM rotenone in these experiments, and the reported apparent dissociation constant of rotenone binding at the I_Q_ site is 20 nM ^48^. In that study, the I_50_ value was determined using submitochondrial particles with a CoQ_10_ analogue. To properly identify the rotenone dissociation constant, kinetic and free radical studies using intact mitochondria with the native Q molecule used by complex I are necessary. Thus, we opted to fit this parameter with our calibration data sets. The resultant model simulations predict that complex I operates at 2% of its maximum capacity. That said, assuming simple inhibitor kinetics and the above stated I_50_ value for rotenone, residual complex I activity is expected to be ~2% which fits with our model simulations. With the J_O2_, J_H2O2_, and now the NADH calibrations, we next sought to challenge our model with an additional substrate combination without adjusting any model parameter other than setting the initial conditions (substrate concentrations).

**Figure 3.**
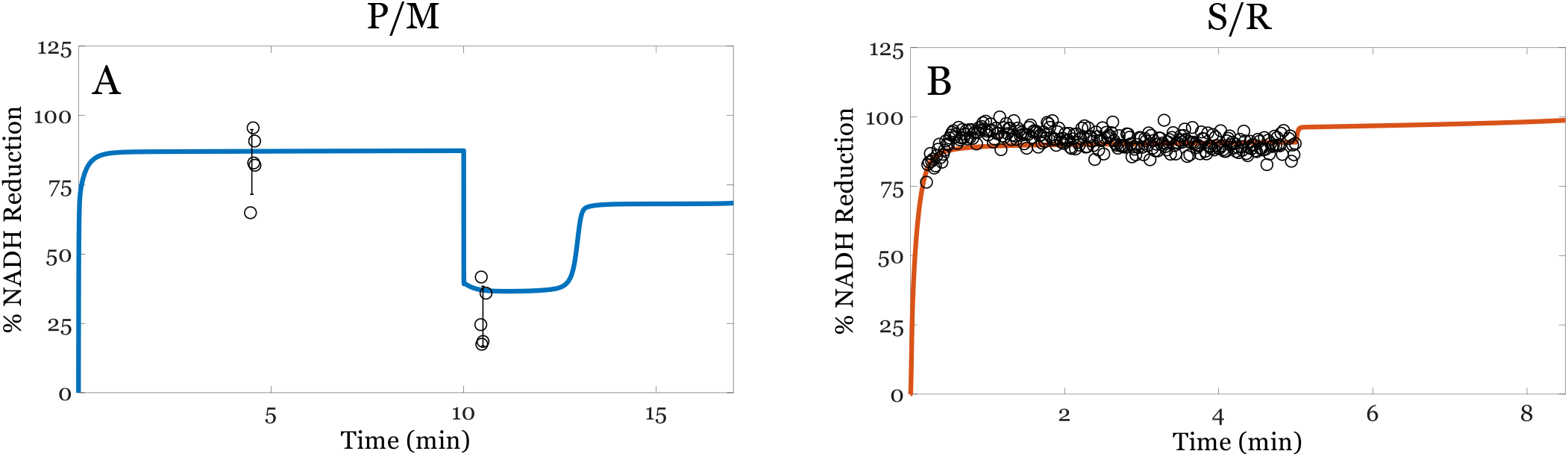
Model simulations of NADH reduction (%). A) Pyruvate/L-malate (P/M). B) Succinate with rotenone (S/R). The model outputs are represented by solid lines and within the experimental ranges. Individual data points are included; error bars represent the standard deviations. Experimental data are obtained using mitochondria isolated from the ventricular cardiomyocytes of guinea pigs. As shown, the model outputs are within the experimental ranges. Overall, succinate-supported respiration in the presence of rotenone maintains a more reduced NADH pool compared to P/M.

### 2. Model Corroboration

#### Oxygen consumption, net hydrogen peroxide production rates, and oxygen concentration dynamics under both NADH- and QH_2_-linked substrate metabolism

The final adjustable parameter set (Table 2) was used to simulate the model and predict the J_O2_, J_H2O2_ and [O_2_] dynamics when mitochondria were energized with a combination of NADH- and QH_2_-linked substrates (P/M/S). Under this condition, electrons entering the ETS at complexes I and II converge at the Q pool ^49^. Such conditions favor the reduction of the NAD^+^ pool, the Q pool, and a high membrane potential, resulting in ROS generation at complex I through RET. As shown in Figure 4, the ETS-ROS model outputs are consistent with experimental data collected from the P/M/S condition. Experimentally determined leak-state J_O2_ (nmol/mg/min) are 129 ± 5 at 2 minutes and 121 ± 12 at 4 minutes. The oxphos-state J_O2_ is 729 ± 91 nmol/mg/min. The corresponding simulated J_O2_ (nmol/mg/min) are 116 at 2-minute leak state, 114 at 4-minute leak state and 654 during oxphos (Fig. 4A). Experimentally measured J_H2O2_ (pmol/mg/min) are 4,304 ± 499 at 2-minute leak state, 3,843 ± 713 at 4-minute leak state and 82 ± 7 during oxphos. The corresponding model predictions are 4,449, 3,100, and 42 pmol/mg/min (Fig. 4B). In general, J_O2_ and J_H2O2_ were significantly elevated compared to the calibration substrate mixtures which favor electron entry at either complex I or II. With the P/M/S substrate combination, both complex I and complex II are fully engaged to deliver electrons into the Q pool during oxphos. This will increase the respiration rate with the high substrate delivery favoring reduction of mitochondrial redox couples. The model predicts the resting and active respiratory states with very good accuracy. The small underestimation of ROS emission in the oxphos state is due to minor sources we did not include in this model.

**Figure 4.**
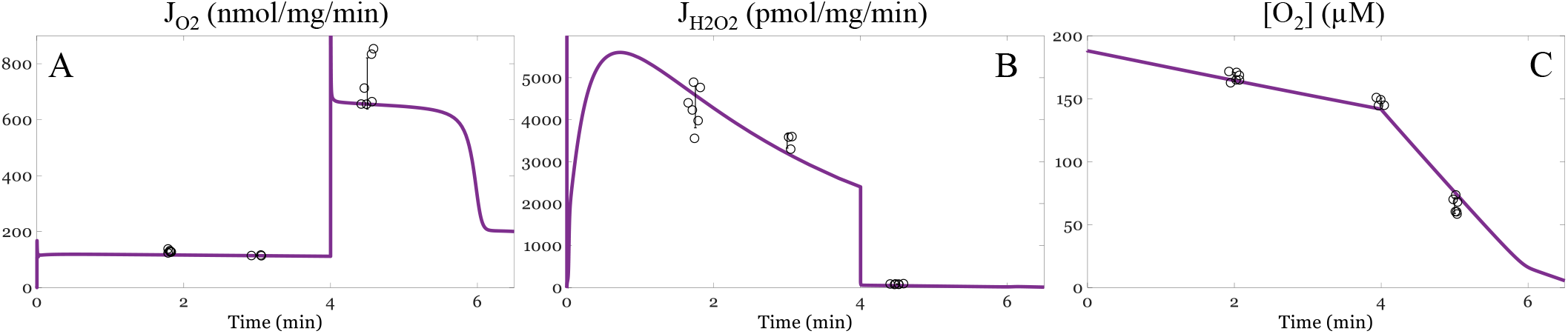
Model simulations of oxygen consumption rates (J_O2_, nmol/mg/min), hydrogen peroxide emission rates (J_H2O2_, pmol/mg/min) and oxygen concentration dynamics when the substrates include both pyruvate/L-malate (P/M) and succinate (S). The model outputs are represented by the solid lines. Individual data points are shown by the open black circles; error bars represent the standard deviations. Experimental data are obtained using mitochondria isolated from the ventricular cardiomyocytes of guinea pigs, as described in our previous work ^45^. Isolated mitochondria (0.1 mg/mL) were energized with P/M/S. The respiratory and net hydrogen peroxide emission rates were quantified simultaneously using the O2k oxygraph.

#### Monotonic relationship between J_H2O2_ on [O_2_]

Although J_O2_ is essentially independent of [O_2_] until it drops below 1 μM, the monotonic relationship between J_H2O2_ and [O_2_] has been described by several groups ^45,50,51^. To test whether the ETS-ROS model can adequately capture this dependence, we compared the simulated J_H2O2_ to experimentally determined values at varying [O_2_]. As shown, the model simulations are consistent with experimental data when mitochondria are energized with NADH-linked substrates and QH_2_-linked substrates with RET inhibition. Further, the monotonic relationship between J_H2O2_ and [O_2_] is dependent on substrate utilization. In the S/R condition, the net J_H2O2_ is more sensitive to changes in [O_2_], and a non-linearity is observed at low [O_2_]. Model analysis reveals this non-linearity is a manifestation of how mitochondrial produce hydrogen peroxide at the lower [O_2_] (Fig. 5B). The rapid drop in hydrogen peroxide production as [O_2_] approaches zero is due to a decreased availability of the reduced I_FMN_. In our model, flavin sites must be unoccupied before they are able to react with O_2_. This occurs because of the following: First, the reduced I_FMN_ preferentially binds to NAD^+^ over NADH ^38,52^. Second, the kinetic pressures between NADH production (via residual malic enzyme activity) and NADH consumption (via ROS production reactions) results in a gradual but complete reduction of the NAD pool as the [O_2_] required for ROS production approaches zero. Consequently, the net J_H2O2_ vs [O_2_] is non-linear with the S/R combination which we will now explicitly show in the next section.

**Figure 5.**
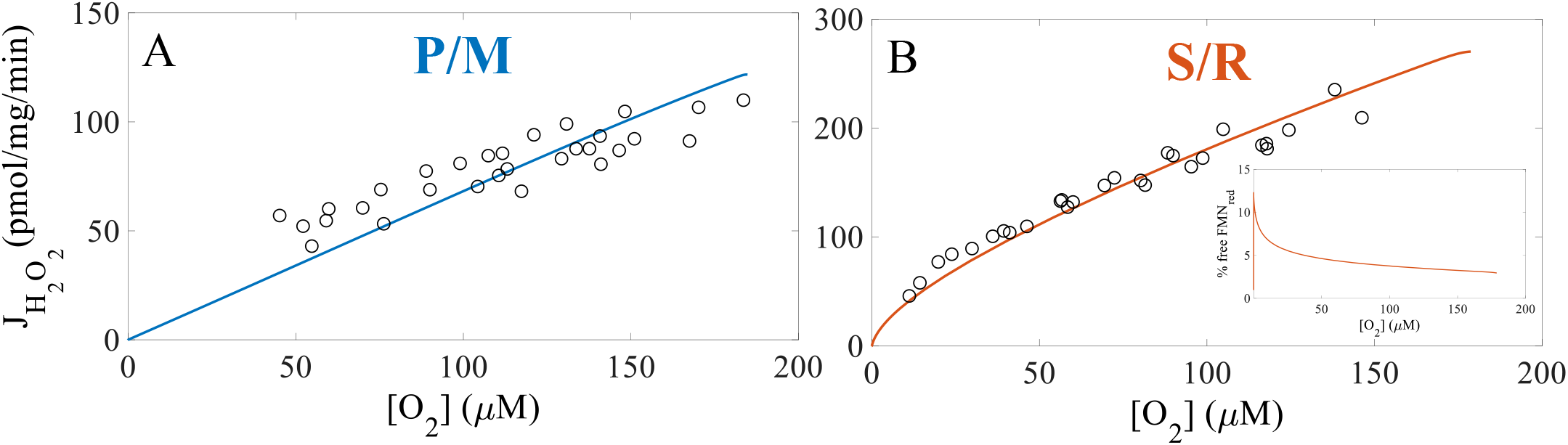
Model simulation of the monotonic relationship between net hydrogen peroxide emission rates (J_H2O2_, pmol/mg/min) and oxygen concentrations (μM). Open, black circles represent experimental data; error bars represent the standard deviations. Model simulations are shown by the solid lines. Experimental data and simulated values were leak-state J_H2O2_ values at varying oxygen concentrations. A) Mitochondria were energized with P/M. B) Mitochondria were energized with S and RET is inhibited (S/R). As shown, the model is able to capture the monotonic dependence between J_H2O2_ and [O_2_] for both substrates. Importantly, the monotonicity appears linear with P/M and non-linear with S/R as the substrates.

### 3. Model Predictions

The high level of consistency between model simulations and experimental data demonstrate that our model is structurally sound and appropriately calibrated. We next used the ETS-ROS model to 1) investigate the linear and non-linear relationship of [O_2_] on J_H2O2_, 2) predict the effects of substrate utilization on site-specific and species-specific contributions to mitochondrial ROS production during different respiratory states and ETS flux directions, 3) assess the degree of mitochondrial scavenging activity and percent utilization during the experimental conditions in the calibration and corroboration data sets, and 4) elucidate the impact on the rate of ROS emission of the redox states of the NAD and Q pools, along with the mitochondrial membrane potential. These model predictions provide valuable mechanistic information about mitochondrial redox homeostasis that are experimentally inaccessible.

#### Hydrogen peroxide production by site I_F_ underlies the kinetics of net ROS emission rates at low [O_2_]

Preliminary model analysis of the monotonic dependence between net J_H2O2_ and [O_2_] reveals that, when mitochondria are energized by succinate and RET is inhibited with rotenone, H_2_O_2_ production is non-linearly sensitive to changes in [O_2_] (Fig. 5B). We previously explained this phenomenon with an empirical model which was unable to identify redox center contributions to this behavior ^45^. Using our computational ETS-ROS model, we dissected the individual processes underlying the net J_H2O2_ measurements as shown in Fig. 6. Here, the total ROS (J_ROS_) is defined with respect to 2-electron equivalents and equals the sum of hydrogen peroxide (J_H2O2_) plus half of the superoxide flux (J_SO_/2). The scavenging capacity shown in right y-axis of panels shown in Fig. 6A and 6D reflect the fraction scavenging capacity utilization. In the P/M condition, the scavenging system activity is only a few percent of the maximum rate of H_2_O_2_ clearance. In contrast, the scavenging system activity increases to 15% when mitochondria are energized with S/R. This increase in scavenging activity is expected as mitochondria respiring in the S/R condition produce more ROS compared to the P/M condition (Fig. 6A and 6D). This goes up even further when S or P/M/S are used as respiratory fuels, as we will show later. In the presence of P/M, a linear dependence between total J_ROS_ and [O_2_] was observed, and H_2_O_2_ production is low (Fig. 6A). However, J_ROS_ exhibit a non-linear dependence on [O_2_] under S/R-supported respiration (Fig. 6D). The primary reason for this difference is the H_2_O_2_ producing activity of the I_F_ site.

**Figure 6.**
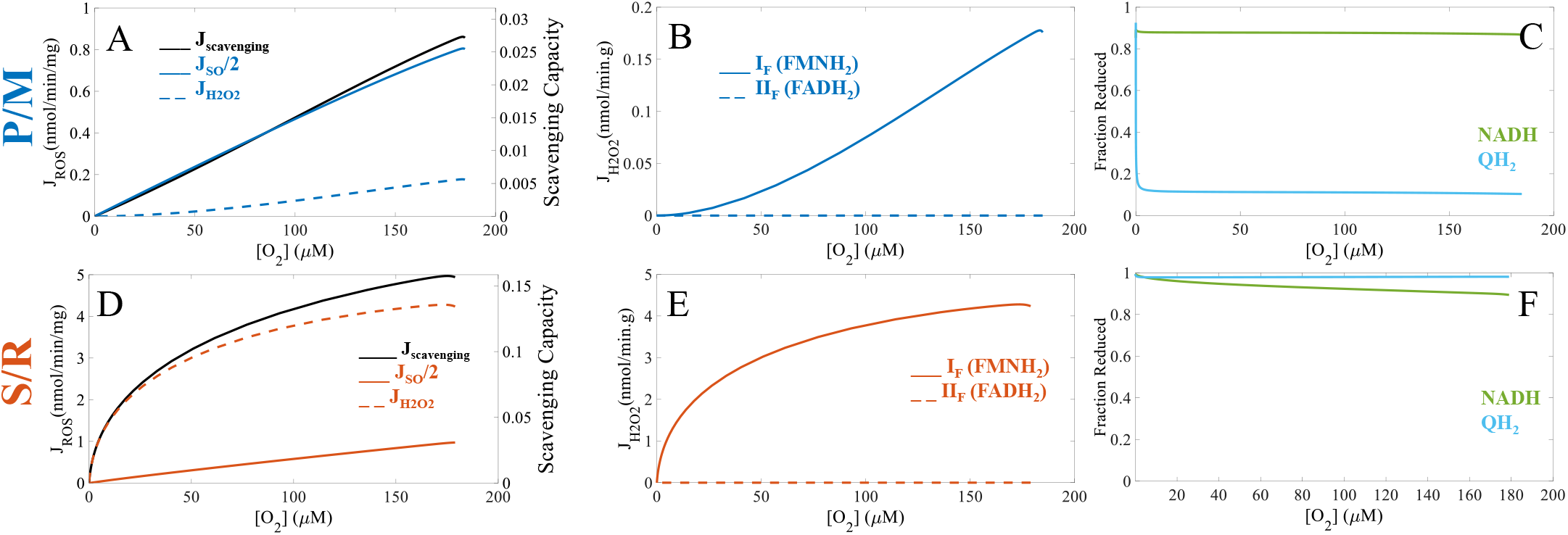
Model analysis of site-specific H_2_O_2_ production and bioenergetics variables critical to mitochondrial ROS production over a range of [O_2_] (μM). To further understand the substrate-specific monotonicity associated with the net J_H2O2_ on [O_2_] (Fig. 5), model analysis was performed to extract additional mechanistic insights. The total ROS, superoxide and hydrogen peroxide fluxes are shown with active scavenging under P/M (A) and S/R (D) simulations. Total ROS flux (J_ROS_, solid lines) is defined as the sum of hydrogen peroxide flux (J_H2O2_, dotted lines) and half of the superoxide flux (J_SO_, dashed lines). The fully reduced flavin of complex I contributes to most J_H2O2_ under both P/M-(B) and S/R-(E) supported respiration. It also underlies the non-linearity associated with JROS at low [O_2_] in the S/R simulation. The model further demonstrates that the redox states of the NADH and the Q pools are highly sensitive to experimental conditions (C and F). Rotenone inhibits QH_2_ oxidation at site I_Q_, which indirectly prevents effective NADH oxidation at site I_F_. The NADH pool is, thus, significantly reduced in the S/R simulation, creating a condition conducive for H_2_O_2_ production.

Regardless of the substrates, most H_2_O_2_ originates from the fully reduced flavin of complex I which produces the non-linear relationship between J_ROS_ and [O_2_] for S/R supported respiration (Fig. 6B and 6E). The combination of highly reduced NAD^+^ and Q pools redox states explains the J_H2O2_ profile associated with each substrate combinations (Fig. 6C and 6F). In the P/M simulation, the NADH pool is less reduced, and the Q pool is significantly oxidized. In contrast, rotenone in the S/R simulation prevents quinol oxidation at site I_Q_ and NADH oxidation at site I_F_, maintaining the NADH and Q pools in mostly reduced states. As seen in Fig. 6F, the NADH level at high [O_2_] is 90% reduced and rises to 100% as [O_2_] approaches zero. At high [O_2_], most of the reduced flavin is bound up with NAD^+^ despite it being only 10% of the NAD^+^ pool. This is because NAD^+^ binds with about 10-fold higher affinity than NADH to the reduced flavin on complex I. However, as O_2_ is depleted, the NAD^+^ pool becomes progressively reduced freeing the binding site near the reduced flavin to maintain H_2_O_2_ production at elevated rates until anoxia (Fig. 5B, inset).

#### Site-specific superoxide and hydrogen peroxide production rates

As shown in Figures 7 and 8, the site-specific contributions to total superoxide and hydrogen peroxide production are determined by substrate combinations, respiratory states, and ETS flux directions. The site-specific superoxide production rates are also shown in Table 5 whereas site-specific hydrogen peroxide production rates are included in the text below.

**Figure 7.**
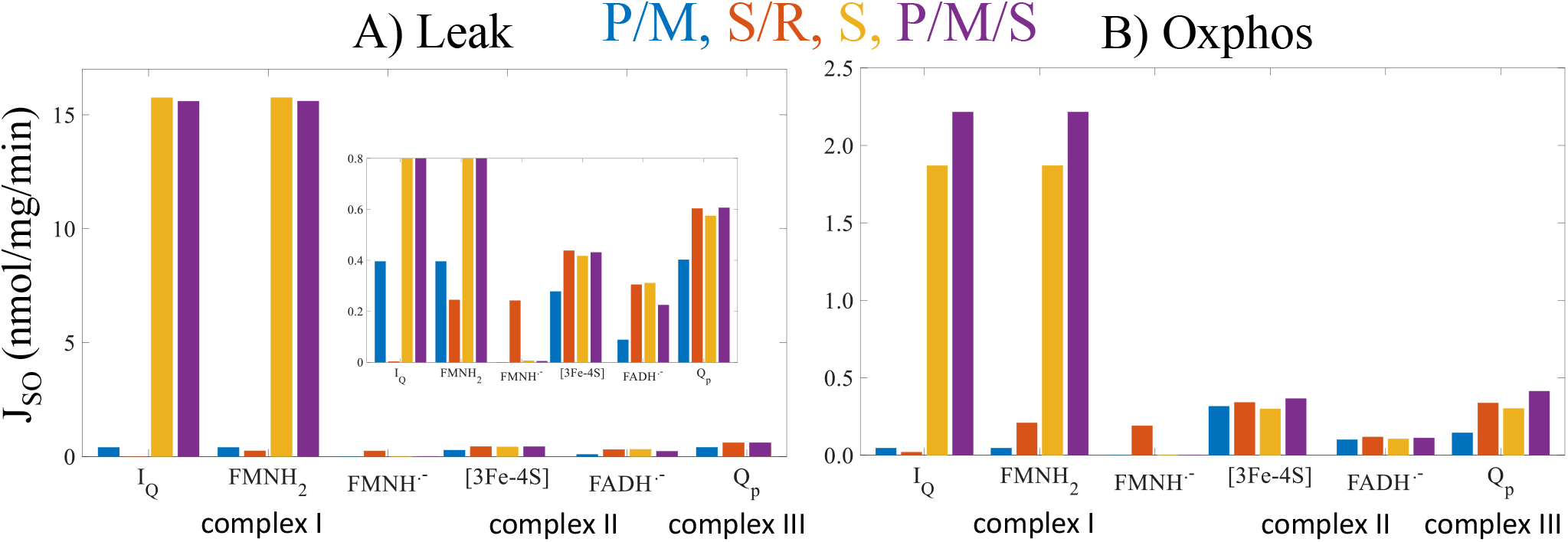
Model predictions of superoxide production at ETS redox centers in leak (A) and oxphos (B) states. In non-phosphorylating mitochondria (leak) energized with P/M or S/R, forward electron transport (FET) is favored whereas S and P/M/S favor reverse electron transport (RET). Complex I redox centers include the quinone reductase site (I_Q_) and the flavin mononucleotide (I_FMNH2_ = reduced flavin mononucleotide, I_FMNH.-_ = flavin mononucleotide radical). Complex II redox centers are the [3Fe-4S] cluster (II_[3Fe-4S]_) and the flavin adenine dinucleotide (II_FADH.-_ = flavin adenine dinucleotide radical). The Q_p_ site resides in complex III (III_Qp_). Overall, superoxide production is lowest under P/M-supported respiration and minimized by phosphorylation. The highly reduced Q pool and a high membrane potential under S- and P/M/S-supported respiration maintain sites I_Q_ and I_F_ in the highly reduced states regardless of respiratory states, favoring superoxide production by these sites I_Q_ and I_F_ and making complex I the greatest combined source of superoxide.

**Figure 8.**
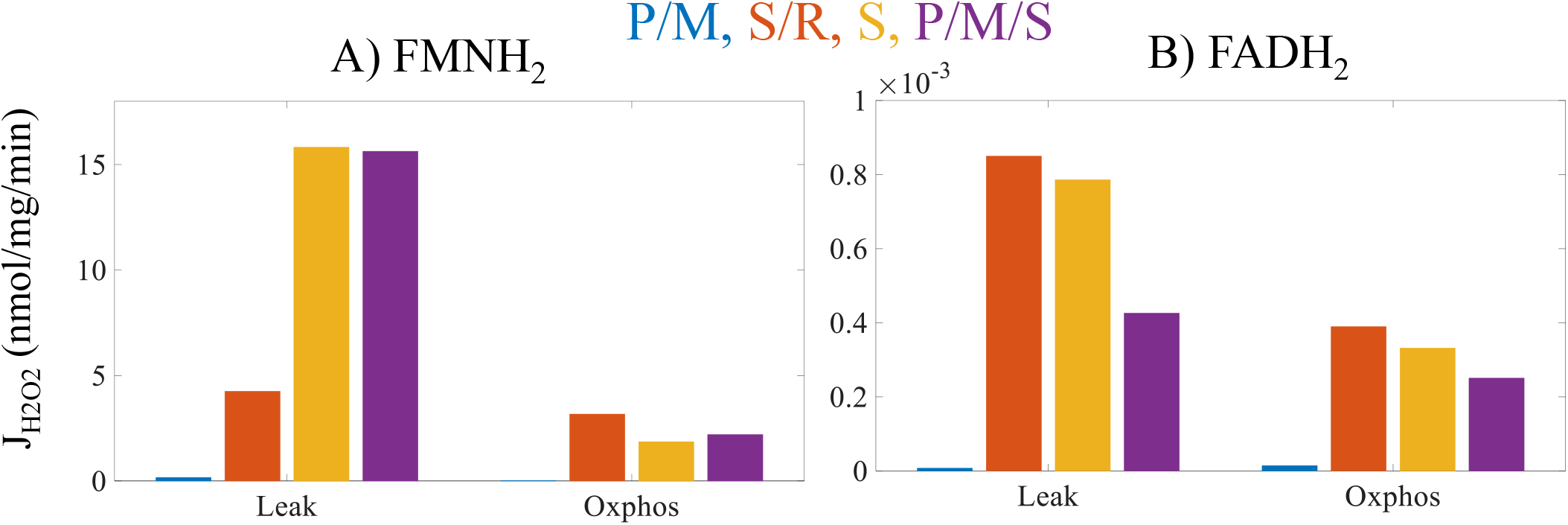
Model predictions of hydrogen peroxide production at ETS redox centers at leak (A) and oxphos (B) states. Only the I_FMNH2_ and II_FADH2_ are considered as hydrogen peroxide production requires 2 electrons. Left axis values are for I_FMNH2_, and right axis values are for II_FADH2_. In non-phosphorylating mitochondria energized with P/M or S/R, forward electron transport (FET) is favored whereas S and P/M/S favor reverse electron transport (RET). Overall, hydrogen peroxide production is lowest under P/M-supported respiration and minimized by phosphorylation, similar to superoxide production. Regardless of the respiratory states and electron flux directions, most hydrogen peroxide originates from the I_FMNH2_.

**Table 5.** Model predictions of site-specific superoxide production (J_SO_, nmol/mg/min) during leak and oxphos states supported by different substrates.

| State | Leak |  |  |  | Oxphos |  |  |  |
| --- | --- | --- | --- | --- | --- | --- | --- | --- |
| Site / Substrates | P/M | S/R | S | P/M/S | P/M | S/R | S | P/M/S |
| $I_Q$ | 0.395 | 0.003 | 15.7 | 15.6 | 0.046 | 0.019 | 1.87 | 2.22 |
| $I_{FMNH_2}$ | 0.395 | 0.244 | 15.7 | 15.6 | 0.046 | 0.209 | 1.87 | 2.22 |
| $I_{FMNH_-}$ | ~0 | 0.241 | 0.006 | 0.004 | ~0 | 0.190 | 0.001 | 0.001 |
| $II_{[3Fe-4S]}$ | 0.277 | 0.436 | 0.417 | 0.431 | 0.315 | 0.340 | 0.299 | 0.365 |
| $II_{FADH_-}$ | 0.087 | 0.303 | 0.311 | 0.224 | 0.10 | 0.117 | 0.106 | 0.111 |
| $III_{Qp}$ | 0.401 | 0.601 | 0.574 | 0.606 | 0.145 | 0.337 | 0.301 | 0.413 |

Values less than $10^{-4}$ are shown as ~0.

During leak state with substrate combinations favoring RET (S and P/M/S), ROS is produced chiefly at sites I_Q_ and I_FMNH2_ (Fig. 7A). The rates from these sites exceed 25 times that of the next highest site. When operating in FET mode, these two sites are joined by III_Qp_ as the top three ROS producers. In the S/R condition, sites I_Q_ and I_FMNH2_ are replaced by both CII sites as top ROS producers when their individual contributions are combined. Both complex II sites increase their individual ROS production rates due to the presence of a highly reduced Q pool. During oxphos, bioenergetic conditions are quite different and the mitochondrial redox couples become more oxidized, in general.

Figure 7B shows that the ROS production rates in the oxphos state from all sites are dramatically reduced with a similar profile relative to the leak state. Notably, sites I_Q_ and I_FMNH2_ remain the sites with the highest capacity to produce ROS. Despite high ETS turnover, complex I redox centers are maintained in a reduced state, producing high ROS emission rates. In the oxphos state, the NAD^+^ pool becomes more oxidized, but the Q pool gets more reduced. This is due to the activation of parts of the TCA cycle via metabolite and allosteric modulatory changes ^18^. As a result, complex II and III sites do not significantly drop below leak-state rates despite a large drop in membrane potential and NADH levels. The high rates of sites I_Q_ and I_FMNH2_ in S and P/M/S conditions during the oxphos state is somewhat surprising. During leak state, the FMN of complex I is operating in a near-equilibrium state under highly reducing conditions. The I_Q_ site feeds electrons into the complex to support ROS production at both sites. In contrast, the sites at complex I are displaced from equilibrium during oxphos and contribute to electron flow down to O_2_. Yet, in this state, these sites still produce ROS at a rate of around five times or more when compared to the other sites. The sites that produce H_2_O_2_ show a similar pattern.

As direct H_2_O_2_ production by the respiratory chain is a two-electron redox reaction, only the I_FMNH2_ and II_FADH2_ are considered as ETS sources of H_2_O_2_ in the model. Regardless of respiratory states and substrate sources, almost all H_2_O_2_ produced by mitochondria originate from the I_FMNH2_ site (Fig. 8). Site II_FADH2_ produces very little H_2_O_2_ in the presence of fumarate ^39^. Transitioning to oxphos from leak state leads to a more dramatic reduction J_H2O2_ relative to J_SO_. This is because J_H2O2_ relies more heavily on NADH rather than QH_2_ or the membrane potential ^38,39^. In the S/R condition, the mitochondrial NAD pool is more reduced relative to the P/M condition due to the i) presence of malic enzyme activity and ii) inhibition of complex I. Under S and P/M/S conditions, the model predicts that the hydrogen peroxide production rate from site I_FMNH2_ is similar. Consequently, most of the decrease in total hydrogen peroxide after the transition to oxphos results from decreased production by this site. However, phosphorylating mitochondria energized by P/M/S produce more hydrogen peroxide from the I_FMNH2_ site compared to S alone. In the P/M/S condition, pyruvate metabolism produces NADH through the TCA cycle which resutls in a more reduced NAD^+^ pool. Consequently, the I_FMNH2_ site is more reduced under P/M/S-supported respiration, enhancing hydrogen peroxide arising from this site.

#### Effects of substrate utilization and electron transport mode on scavenging activity

The activity of the scavenging systems’ response to changes in ROS production remains an enigma for the same reasons that the origins of mitochondrial ROS under native conditions are inconclusive. Therefore, due to the lack of quantitative data, we modeled the scavenging system assuming that it is saturable and responsive to changes in matrix H_2_O_2_ production. Because regenerating the scavenging system relies on NAD(P)H, ROS scavenging has a limited capacity ^10^. Thus, the responsive nature of the scavenging system to matrix H_2_O_2_ is necessary to keep net ROS production in the physiological range. In this model, these assumptions were adequate as the model outputs in the calibration and corroboration stages are consistent with experimental data. Thus, we used the model to predict the rate of ROS scavenging in the experimental conditions simulated in Figure 1. The model reveals that scavenging activity is lower in FET mode, which is unsurprising since it is consistent with lower ROS production. In RET mode, the scavenging activity is operating near maximum and ROS emission rates increase nearly 40-fold. Thus, changes in scavenging capacity or H_2_O_2_ sensitivity can result in elevated ROS emission rates that will lead to cellular oxidative stress.

#### Primary determinants of ROS production

The corroborated model is further used to explore the effects of the mitochondrial membrane potential along with the NADH and Q pool redox state on J_SO_ and J_H2O2_ from individual complexes (Fig. 10). In this simulation, we simulated the model until reaching the steady state with respect to mitochondrial state variables under the P/M condition. We clamped substrates at saturating levels and oxygen at a physiological level of 20 μM ^10^ in this simulation. We then perturbed the membrane potential, NAD^+^ redox state, and Q pool redox state while keeping the other state variables fixed and computed the resultant free radical fluxes. This simulation exemplifies the utility of model in testing hypotheses that are not experimentally feasible.

The model predicts a maximum rate of ROS production occurs when the membrane potential is high and the NAD^+^ and Q pools are extremely reduced (Fig. 10). In general, at high membrane potentials, mitochondrial ROS production is more sensitive to NADH levels than QH_2_ levels. When NADH levels begin to exceed 90%, ROS production exponentially increases. That said, QH_2_ levels still significantly affect total ROS produciton showing saturation at more oxidized levels. Of the H_2_O_2_ producing sites, complex I produces significantly more ROS compared to complex II. Its maxima lie along the 100% NADH and 220 mV membrane potential plane where QH_2_ is maximally reduced. In contrast to the H_2_O_2_ producings sites, the rates of the SO producing sites are distributed differently. Complex I dominates in RET mode and contributes little to SO production under more oxidized conditions. Complex II produces the least amount of SO under these conditions and is independent of NADH. As expected from our prior study, complex II SO production is very sensitive to QH_2_ levels. In contrast, complex III produces a moderate amount of SO and contributes to the basal level of SO production by the mitochondrial respiratory chain. As with the other ETS pumps, this complex produces most SO when the membrane potential, NADH levels, and QH_2_ levels are high.

## IV. Discussion

### 1. Model predictions of site-specific ROS from the ETS complexes

Site-specific ROS production has inspired many experimental studies and led to a wealth of experimental data. However, the data are limited to experimental conditions wherein inhibitors are present. The use of different systems, experimental conditions and techniques has also led to discrepancies among these studies. Simpler systems, such as purified enzymes, have the advantage that fewer variables need to be accounted for in interpreting experimental results. But the conclusions may not be applicable when these results are integrated into biochemical reaction networks. On the contrary, a more intact system affords an environment more similar to *in vivo* conditions but introduces more hidden variables. Perhaps the biggest hidden variable in all mitochondrial studies is the dynamic redox state of the Q pool. There is currently no experimental method that can be used to quantify the real-time dynamic changes of the Q pool without sample destruction. Additionally, factors that affect thermodynamics such as temperature, buffer ionic strength and buffer pH can all regulate enzyme activities but are often neglected. These changes can lead to large changes in pathway fluxes and study outcomes. A computational approach can be applied to fully realize unified analyses at high temporal and spatial resolutions. Thus, different experimental conditions are analyzed using a unified framework to reconcile contradictory reports. This quantitative framework not only checks for the internal consistency of data but also enables extrapolation to experimentally untestable space for hypothesis testing or generation.

While several mitochondrial ROS models exist ^41,42,53–56^, none can reproduce the wide variety of available mitochondrial bioenergetic data quantitatively and consistently. These models either lack the biophysical details to simulate the enzymatic reactions associated with both high and low electron flux regimes ^42,56^ or exclude complex II as a ROS-producing component ^53^. Our ETS-ROS model was constructed using validated models of complexes I, II, and III. Each of these models contains the biophysical details necessary to simulate a wide variety of bioenergetic data ^38–40^. The models were calibrated and validated before they were integrated into the ETS-ROS model. The integrated model was again calibrated (Figs. 2, 3) and validated (Figs. 4, 5) before the model was used to generate mechanistic predictions (Figs. 7–10). Only processes that are supported by experimental data are explicitly modeled, and sensitivity analysis was performed on the integrated model to refine the adjustable parameters (Table 3).

Using the validated model, we identify the ROS species and their origin (Figs. 8, 9) under the different metabolic conditions of our experimental data (Fig. 2). We found that the III_Qp_ site, site I_FMNH2_ and site I_Q_ as a semiquinone are the primary sources of superoxide in non-phosphorylating mitochondria energized with P/M (Fig. 7, Table 5). In non-phosphorylating mitochondria, site I_FMNH.-_ gives rise to a considerable amount of superoxide under QH_2_-supported respiration when RET is inhibited (S/R). These model predictions reconcile experimental results regarding complex I from different studies. For example, Galkin and Brandt ^30^ and Grivennikova & Vinogradov ^29^ independently concluded that both the fully reduced flavin (FMNH_2_) and the flavin radical (FMNH^.-^) can give rise to superoxide. Kussmaul and Hirst also found that site I_F_ is a primary source of superoxide but only when fully reduced ^28^. In contrast, Lambert and Brand reported that site I_Q_ is the predominant source of superoxide originating from Complex I ^21^. The Ohnishi group found that both sites I_Q_ and I_F_ are ROS sources when they exist in the radical forms ^31^. Thus, computational modeling shows that the differences among experimental data are likely due to different experimental conditions and that even seemingly contradictory results may be true under the right conditions.

**Figure 9.**
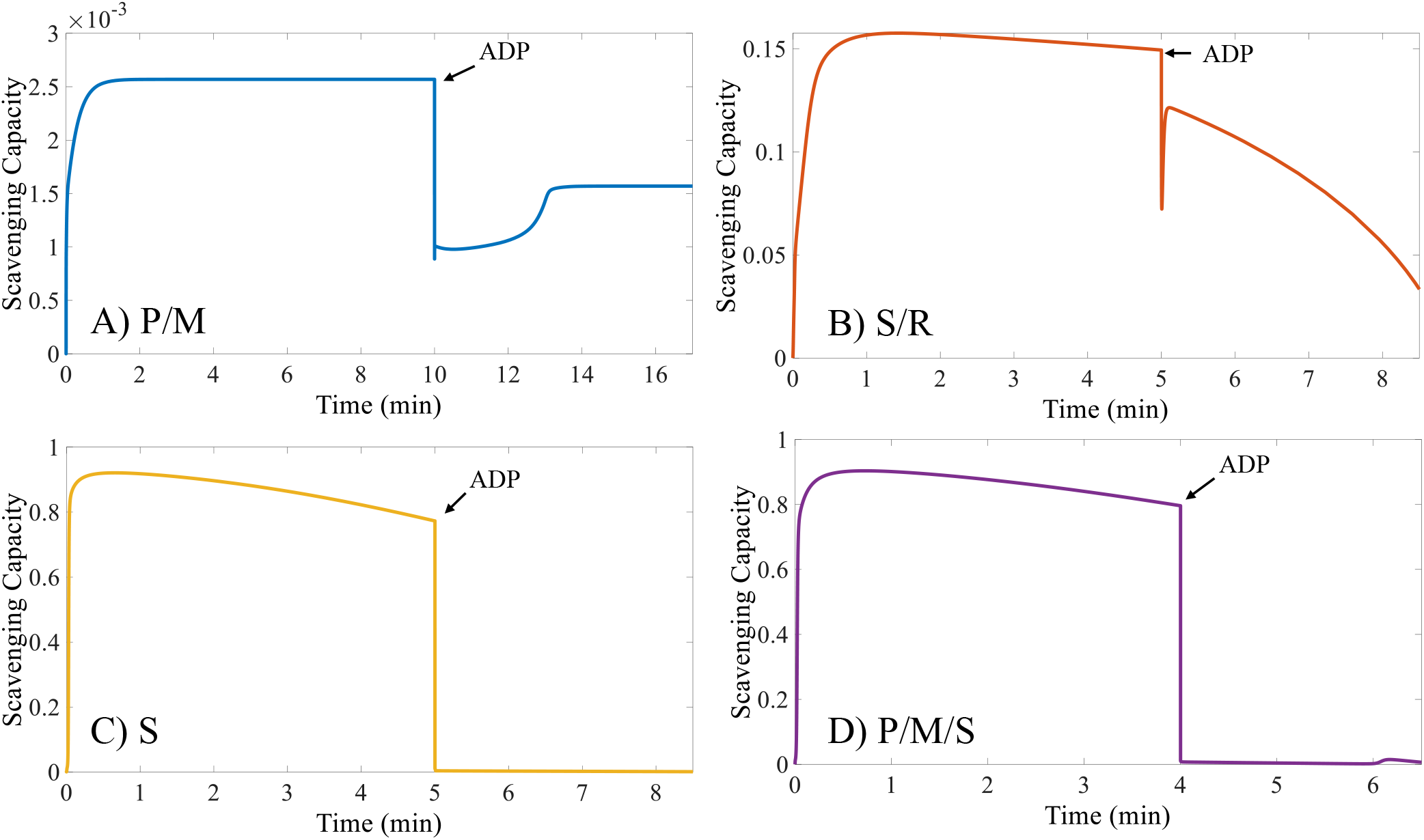
Model predictions of active scavenging under different metabolic conditions. Forward electron transport (FET) occurs under P/M- and S/R-supported respiration (*A, B*). Reverse electron transport occurs under S- and P/M/S-supported respiration (*C, D*). ADP addition results in maximal oxidative phosphorylation (oxphos state). During oxphos, the scavenging activity decreases regardless of the electron source and transport mode. Under RET mode, the scavenging activity is significantly increased but unable to keep the net J_H2O2_ at comparable levels to under FET. The model predictions of active scavenging, thus, together with the changes in J_H2O2_ and membrane potential support the assumptions that the scavenging activity is saturable and responds to changes in ROS production.

**Figure 10.**
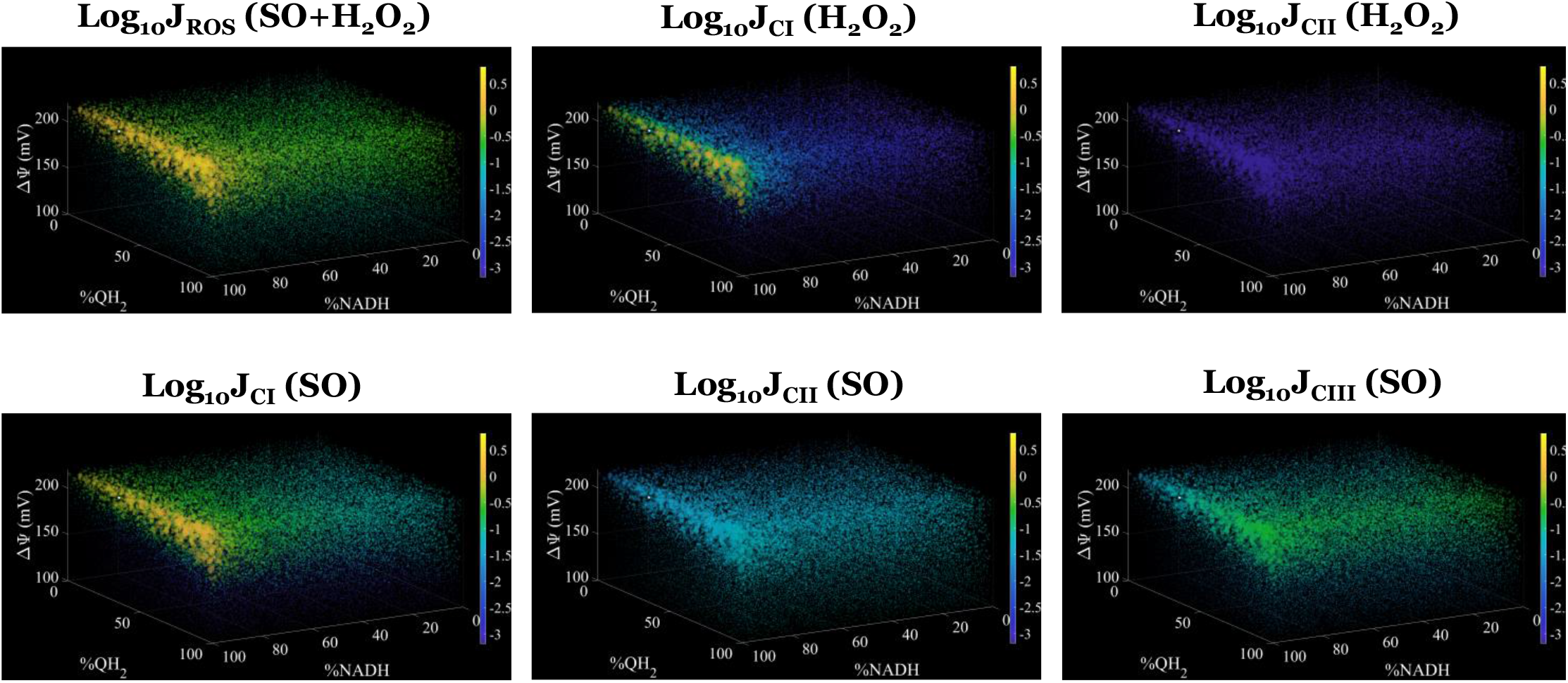
Model predictions of superoxide and hydrogen peroxide fluxes (J_SO_ and J_H2O2_, respectively) from all and individual ETS complexes under P/M supported respiration. The state variables at 5-minute steady state were fixed and the membrane potential (ΔΨ), %NADH, and %QH_2_ were adjusted in the ranges shown. Model outputs are the log_10_ values of each flux with units of nmol/mg/min. Also, model outputs are represented as colored dots where the size of the dot is proportional to the model output’s value relative to the colorbar.

Like complex I, the identity and origin of ROS produced by complex II remain unresolved although until recently, it was thought to be a negligible source of mitochondrial ROS. Quinlan *et al*. found that site II_F_ produces comparable amounts of ROS to site I_Q_ in the presence of fatty acids ^20^. The notion that site II_F_ produces most ROS from Complex II is shared by Siebels and Drose ^34^. However, others contend that significant amounts of ROS originate from the site II_Q_ ^57^ and the ISC near the Q site ^58^. We used the model to identify conditions that favor ROS production by complex II and the contributions of its redox centers. In particular, the model identifies site II_[3Fe-4S]_ as the primary source of superoxide arising from complex II. In FET, superoxide arising from site II_FADH.-_ becomes more significant only under QH_2_-supported respiration (S/R). Thus, our model supports the experimental results that the ISC near the Q site and site II_F_ give rise to most of complex II’s ROS. Moreover, the model predicts that FET in non-phosphorylating mitochondria favors superoxide production by complex III. The bc1 complex also contributes significantly to total superoxide flux in phosphorylating mitochondria although its contribution is more pronounced in FET mode. Unfortunately, all these predictions remain difficult to experimentally test due to the lack of a robust, quantitative, and straight-forward method to directly measure them without interfering with other processes.

### 2. Future directions

In developing this model, we focused on the known primary sources of mitochondrial ROS, the ETS complexes. Electron flavoprotein ^59–61^, dihydrolipoamide dehydrogenase ^15,16,62^, and other mitochondrial dehydrogenases ^63,64^ are additional sources of ROS that may become significant in different physiological conditions and tissues. In the next evolution of the ETS-ROS model, we will expand the model to include two major modifications. The first is to expand the model to explicitly include the full TCA cycle with additional enzymatic processes uncovered during the development of this model (e.g., malic enzyme). This requires additional experiments to collect data on metabolite profiles under the various conditions. Second, we will develop models of beta-oxidation that are able to simulate both kinetic turnover and ROS production from electron flavoprotein and other associated ROS processes. This will, again, require additional experiments focused on fatty acid metabolism under various conditions spanning across the anticipated physiological domain. Lastly, we will revisit the scavenging system to include glutathione and thioredoxin scavenging enzymes. Despite the high level of consistency between model simulations and experimental data (Figs. 2–5), this approach is incapable of partitioning the roles of the glutathione and peroxiredoxin systems involved in the ROS detoxifying pathway. Addressing these aspects require additional experimental data under the prevailing conditions to properly calibrate the model. Individual kinetic models of each of the four primary enzymes responsible for most of the H_2_O_2_ scavenging are already developed, but they remain untested in an integrated model ^65–67^. In all cases, each future generation of the model will be more refined and enable testing of hypotheses that are experimentally infeasible.

### 3. Conclusions

In conclusion, we have developed, analyzed, and corroborated a model of mitochondrial ROS that identifies the species-specific ROS originating from the ETS redox centers under varying respiratory states and electron flux directions. Each ROS-producing module is constructed in a thermodynamically faithful manner and tested prior to being unified in a single platform ^38–40^. The biophysical details included in the model are supported by experimental data and further refined by the process of sensitivity analysis. Being modular, parsimonious, and consistent, this modeling approach enables the individual modules within the model to work harmoniously with each other in a thermodynamically faithful manner. This approach also facilitates the inclusion of other mitochondrial physiology processes that are related to ROS homeostasis, as mentioned above, when data become available. These qualities distinguish this model from existing models which also attempt to recapitulate mitochondrial ROS homeostasis ^41,42^ as ours is capable of consistently reproducing experimental data on various aspects of mitochondrial physiology and making predictions that are physiologically relevant.

## Supporting information

Supplemental Figs and Equations

Model Codes

## Data availability

All data, model equations, and model codes described in this manuscript are given in the main body or supplement.

## Author Contributions

Concept and design: JNB, QVD

Acquisition of data: JNB, QVD, MJD, YL

Analysis and interpretation of data: JNB, QVD, YL

Development of computer code: JNB, QVD

Drafting or revising the article: JNB, QVD, YL

## Abbreviations

SDH: succinate dehydrogenase
ROS: reactive oxygen species
ISC: iron sulfur center
SQ: semiquinone
O_2_^•-^: superoxide
H_2_O_2_: hydrogen peroxide
I/R: ischemia/reperfusion
PMS: phenazine methosulfate
TMPD: N,N,N′,N′-tetramethyl-p-phenylenediamine

