## Supplemental Figs and Equations for "Identifying Site-specific Superoxide and Hydrogen Peroxide Production Rates from the Mitochondrial Electron Transport System Using a Computational Strategy"

**Running title**: Cardiac electron transport system sources of free radical production

**Keywords**: Electron transport system (ETS), mitochondria, reactive oxygen species, enzyme kinetics, oxidative stress, computational biology, ischemia/reperfusion injury, forward electron transport, reverse electron transport

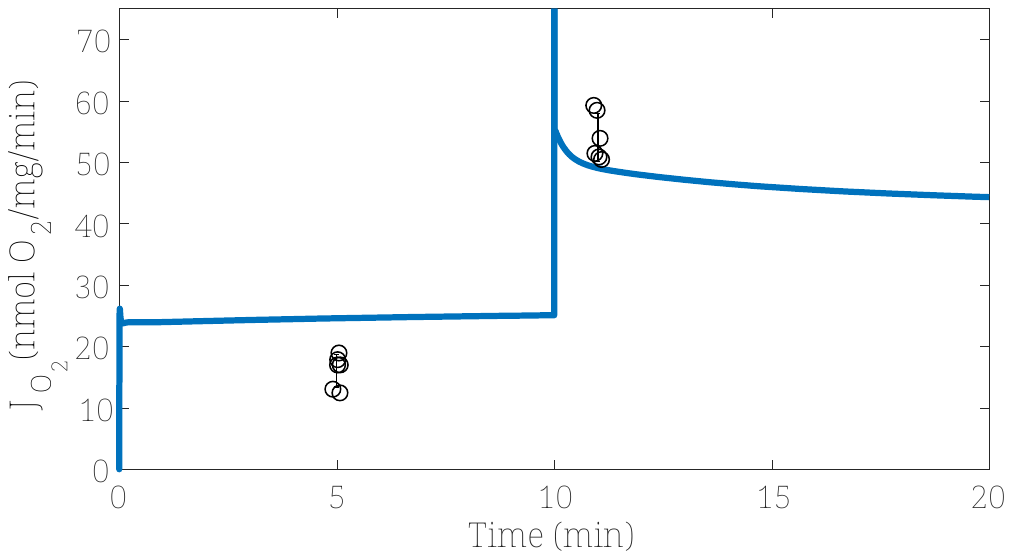

Figure S1. Malate-dependent respiratory dynamics in the absence and presence of saturating ADP. The model slightly overpredicts leak state J_O2_ and slightly underpredicts oxphos J_O2_. This is due to the simplistic nature the malic enzyme reaction was modeled. These fits are expected to improve with a more rigorous malic enzyme kinetic model including allosteric control mechanisms and a more complete TCA cycle.

Malic enzyme reaction is modeled using the following simple equation:

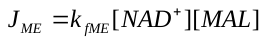

This reaction produces pyruvate, NADH, and a CO_2_ as products.

**Model Codes Description**

Generate_Figures.m – mfile used to reproduce the figures in the article

ETS_ROS_model.m – mfile of the system of differential algebraic equations governing model behavior including individual flux models for individual biochemical reactions

data.mat – mat file containing experimental data structure

parameters.mat – mat file containing model parameter structure

RESULTS.mat – mat file containing pre-ran model simulation results

rng_ii.mat – mat file containing random number selection for 4D plots shown in article
