## Supplementary material for "Identifying Site-specific Superoxide and Hydrogen Peroxide Production Rates from the Mitochondrial Electron Transport System Using a Computational Strategy": Model Codes: ETS_ROS_model.m

```
%% Computing time derivatives of state variables

function varargout = ETS_ROS_model(t,x,p)
% initialize LHS
f = zeros(size(x,1),size(x,2));

%% Compute binding polynomial array
p.BP = ones(size(x,1),size(x,2));
p.BP(p.i.Lmtx,:) = BP(p.if.ligands,'mtx',x,p);
p.BP(p.i.Lcyt,:) = BP(p.if.ligands,'cyt',x,p);

%% Compute Algebriac Constraint Residuals

% Compute other Matrix Conserved Species
f(p.i.ATP_x,:) = x(p.i.ATP_x,:) - (p.Atot - x(p.i.ADP_x,:));
f(p.i.NADH_x,:) = x(p.i.NADH_x,:) - (p.Ntot - x(p.i.NAD_x,:));
f(p.i.UQH2_x,:) = x(p.i.UQH2_x,:) - (p.Qtot - x(p.i.UQ_x,:));
f(p.i.c2_x,:) = x(p.i.c2_x,:) - (p.Ctot - x(p.i.c3_x,:));

%% Compute ETS Fluxes

% Complex I: NADH + UQ + 5H_m = NAD + UQH2 + 4H_c
[J.CI.JNADH,J.CI.JQ,J.CI.JSO,J.CI.JH2O2,J.CI.d] = CI(x,p); % nmol/mg/min

% Complex II: SUC + Q = FUM + QH2
[J.CII.JSUC, J.CII.JSO, J.CII.JH2O2, J.CII.JQH2, ~, J.CII.Xinact, J.CII.d] = CII(x,p); % nmol/mg/min

% Complex III: UQH2 + 2Cytc(ox) = UQ + 2Cytc(red) + 2H_c
[J.CIII.Jc3, J.CIII.JQp, J.CIII.JQn, J.CIII.JSO,J.CIII.d] = CIII(x,p); % nmol/mg/min

% Complex IV: 2Cytc(red) + 1/2O2 + 4H_x = 2Cytc(ox) + H20 + 2H_c
J.CIV = CIV(x,p); % nmol/mg/min

% Proton leak: H_c = H_x
J.mHleak = mHleak(x,p); % nmol/mg/min

%% Compute TCA Cycle and Related Reaction Rates

% partially lumped TCA cycle
% OAA + PYR + 3NAD + ADP + Pi ==> SUC + ATP + 3NADH + 3H+
J.MitoDH = MitoDH(x,p); % nmol/mg/min

% malic enzyme (decarboxylating)
% MAL + NAD = PYR + NADH + CO2
J.ME = ME(x,p); % nmol/mg/min

% DCC
% SUC|MAL_x + Pi_c = SUC|MAL)_c + Pi_x
J.DCC = DCC(x,p); % nmol/mg/min

% FH
% FUM + H2O = MAL
J.FH = FH(x,p); % nmol/mg/min

% MDH
% MAL + NAD = OAA + NADH + H+
J.MDH = MDH(x,p); % nmol/mg/min

% MCT
% H_c + PYR_c = PYR_x + H_x
J.MCT = MCT(x,p); % nmol/mg/min

%% Compute ATP/ADP cycle Fluxes

% ADP + Pi + nH = ATP + H2O
J.F1FO = F1FO(x,p); % nmol/mg/min

% ADP_c + ATP_x = ADP_x + ATP_c
J.ANT = ANT(x,p); % nmol/mg/min 

% HPi_c + H_c = HPi_x + H_x
J.PiC = PiC(x,p); % nmol/mg/min

%% Compute K+ Cycle Fluxes

% K_x + H_c = K_c + H_x
J.mKHE = KHE(x,p); % nmol/mg/min

%% Na+/Ca+ Cycle Fluxes

% Ca_c = Ca_x
J.MCU = MCU(x,p); % nmol/mg/min

% Ca_x + 3Na_c = Ca_c + 3Na_x
J.NCLX = NCLX(x,p); % nmol/mg/min

% Na_x + H_c = Na_c + H_x
J.mNHE = NHE(x,p); % nmol/mg/min

%% Calcium Phosphate Buffering

% Ca + Pi = CaPi (stiochiometry not shown)
J.CaPi = CaPi(x,p,p.CaPi.kf,p.CaPi.K,p.CaPi.n,p.CaPi.m); % nmol/mg/min, (non-saturable model)

%% ROS Scavenging

% 2HO2 = O2 + H2O2
J.SODx = p.kf_SODx*x(p.i.SO_x,:); % nmol/mg/min 

% 2HO2 = O2 + H2O2
J.SODc = p.kf_SODc*x(p.i.SO_c,:); % M/min (arb high rate)

% 2H + H2O2 = 2H2O
J.H2O2scavenge = p.Vmax_scavening*x(p.i.H2O2_x,:)./(p.Km_H2O2 + x(p.i.H2O2_x,:)); % nmol/mg/min

% H2O2_x = H2O2_c
J.H2O2perm = p.kEm_H2O2*(x(p.i.H2O2_x,:) - x(p.i.H2O2_c,:)); % nmol/mg/min

% H2O2_c = Reso (flourescent reporter molecule)
J.HrP = p.kf_HrP*x(p.i.H2O2_c,:); % M/min (arb high rate)

% Net O2 consumption from respiratory chain including ROS metabolism
J.dO2 = - J.CIV/2/p.Vol.H2O_cyt ...
    - (J.CI.JSO + J.CIII.JSO + J.CII.JSO)/p.Vol.H2O_cyt ...
    - (J.CI.JH2O2 + J.CII.JH2O2)/p.Vol.H2O_cyt ...
    + J.SODx/p.Vol.H2O_cyt + J.SODc; % M/min

%% Compute Time Derivatives for Matrix non-cation State Variables

% dPsi
f(p.i.dPsi,:) = (4*J.CI.JNADH + 1*J.CIII.Jc3 + 4*J.CIV - p.F1FO.nH*J.F1FO - J.ANT - J.mHleak)/p.Cmito; % mV/min

% Matrix species
f(p.i.ADP_x,:) = (J.ANT - J.F1FO - J.MitoDH)/p.Vol.H2O_mtx; % M/min
f(p.i.Pi_x,:) = (J.PiC + J.DCC.Pi - J.F1FO - J.MitoDH)/p.Vol.H2O_mtx; % M/min
f(p.i.NAD_x,:) = (-3*J.MitoDH - J.MDH + J.CI.JNADH - J.ME)/p.Vol.H2O_mtx; % M/min
f(p.i.UQ_x,:) = (- J.CI.JNADH + 1/2*J.CIII.Jc3 - J.CII.JQH2)/p.Vol.lipid_imm; % M/min
f(p.i.SO_x,:) = (J.CI.JSO + 1/2*J.CIII.JSO + J.CII.JSO - 2*J.SODx)/p.Vol.H2O_mtx; % M/min
f(p.i.H2O2_x,:) = (J.CI.JH2O2 + J.CII.JH2O2 + J.SODx - J.H2O2scavenge - J.H2O2perm)/p.Vol.H2O_mtx; % M/min
f(p.i.PYR_x,:) = (J.MCT - J.MitoDH  + J.ME)/p.Vol.H2O_mtx; % M/min
f(p.i.SUC_x,:) = (J.DCC.SUC - J.CII.JSUC + J.MitoDH)/p.Vol.H2O_mtx; % M/min
f(p.i.FUM_x,:) = (J.CII.JSUC - J.FH)/p.Vol.H2O_mtx; % M/min
f(p.i.MAL_x,:) = (J.FH + J.DCC.MAL - J.MDH  - J.ME)/p.Vol.H2O_mtx; % M/min
f(p.i.OAA_x,:) = (J.MDH - J.MitoDH)/p.Vol.H2O_mtx; % M/min
f(p.i.O2_x,:) = J.dO2 + 1e-9; % allow small amount of back diffusion to prevent negative O2 during anoxia simulations

%% Compute Matrix Cation Derivatives Based on Rapid Equilibrium Dynamics

f = mCation_dXdt(x,f,p,J); % M/min

%% Compute Time Derivatives for IMS non-cation State Variables

f(p.i.c3_x,:) = (- 1*J.CIII.Jc3 + 2*J.CIV)/p.Vol.H2O_ims; % M/min

%% Reactant Injection (adding bolus amounts to buffer)
J = InjectReactants(t,x,p,J); % M/min

%% External ATPase

J.ATPase = ATPase(x,p); % M/min

%% Compute Derivative for cytosolic/buffer space

% Buffer/Cytosolic species
f(p.i.Pi_c,:) = (J.Inject(p.if.Pi,:) - J.PiC - J.DCC.Pi)/p.Vol.H2O_cyt + J.ATPase; % M/min
f(p.i.ATP_c,:) = (J.Inject(p.if.ATP,:) + J.ANT)/p.Vol.H2O_cyt - J.ATPase; % M/min
f(p.i.ADP_c,:) = (J.Inject(p.if.ADP,:) - J.ANT)/p.Vol.H2O_cyt + J.ATPase; % M/min
f(p.i.SO_c,:) = (+ 1/2*J.CIII.JSO)/p.Vol.H2O_cyt  - 2*J.SODc; % M/min
f(p.i.H2O2_c,:) = J.SODc + J.H2O2perm/p.Vol.H2O_cyt - J.HrP; % M/min
f(p.i.Reso,:) = J.HrP; % M/min
f(p.i.PYR_c,:) = (-J.MCT)/p.Vol.H2O_cyt; % M/min
f(p.i.SUC_c,:) = -(J.DCC.SUC)/p.Vol.H2O_cyt; % M/min
f(p.i.MAL_c,:) = -(J.DCC.MAL)/p.Vol.H2O_cyt; % M/min

%% Compute Cytosolic Cation Derivatives Based on Rapid Equilibrium Dynamics

f = cCation_dXdt(x,f,p,J); % M/min

%% Clamp buffer/cytosolic concentrations to generate steady state
switch p.exp
case 0 
f(p.i.cyt,:) = 0; % clamp all buffer variables
case 1
f([p.i.H_c,p.i.K_c,p.i.Na_c,p.i.Mg_c,p.i.Ca_c],:)=0; % clamp all buffer free cation variables
case 2
f(p.i.H_c,:) = 0; % only fix buffer pH
case 3 % fix cytosolic substrates, O2, and pH
f([p.i.H_c,p.i.PYR_c,p.i.SUC_c,p.i.MAL_c,p.i.O2_x],:)=0; % clamp all buffer free cation variables
end

%% extra output arguments
varargout={f,J};
end

%% Compute Ion Derivatives Using Rapid Equilibrium Approximation Method

function f = mCation_dXdt(x,f,p,J)
%% Compute Time Derivatives for Cations
H_x = x(p.i.H_x,:);     % M
K_x = x(p.i.K_x,:);     % M
Na_x = x(p.i.Na_x,:);   % M
Mg_x = x(p.i.Mg_x,:);   % M
Ca_x = x(p.i.Ca_x,:);   % M

%% Define Ligand Vector
Li = x(p.i.Lmtx,:); % M

%% Define Ligand Derivatives
dLidt = f(p.i.Lmtx,:); % M/min
% set dLidt for algebraic constraints equal to negative of counterpart
dLidt(1,:) = -f(p.i.ADP_x,:); % indices hard coded
dLidt(5,:) = -f(p.i.NAD_x,:); % indices hard coded

%% Partial Derivatives of binding polynomials:
Pm_L = p.BP(p.i.Lmtx,:);
Pm_L2 = Pm_L.^2;
Pm_HBH = (1 + 1./p.KD(p.if.ligands,p.if.K)*K_x + 1./p.KD(p.if.ligands,p.if.Na)*Na_x ...
    + 1./p.KD(p.if.ligands,p.if.Mg)*Mg_x + 1./p.KD(p.if.ligands,p.if.Ca)*Ca_x);
Pm_KBK = (1 + 1./p.KD(p.if.ligands,p.if.H)*H_x + 1./p.KD(p.if.ligands,p.if.Na)*Na_x ...
    + 1./p.KD(p.if.ligands,p.if.Mg)*Mg_x + 1./p.KD(p.if.ligands,p.if.Ca)*Ca_x);
Pm_NBN = (1 + 1./p.KD(p.if.ligands,p.if.H)*H_x + 1./p.KD(p.if.ligands,p.if.K)*K_x ...
    + 1./p.KD(p.if.ligands,p.if.Mg)*Mg_x + 1./p.KD(p.if.ligands,p.if.Ca)*Ca_x);
Pm_MBM = (1 + 1./p.KD(p.if.ligands,p.if.H)*H_x + 1./p.KD(p.if.ligands,p.if.K)*K_x ...
    + 1./p.KD(p.if.ligands,p.if.Na)*Na_x + 1./p.KD(p.if.ligands,p.if.Ca)*Ca_x);
Pm_CBC = (1 + 1./p.KD(p.if.ligands,p.if.H)*H_x + 1./p.KD(p.if.ligands,p.if.K)*K_x ...
    + 1./p.KD(p.if.ligands,p.if.Na)*Na_x + 1./p.KD(p.if.ligands,p.if.Mg)*Mg_x);

pHBpH = + sum(Pm_HBH.*Li./Pm_L2./(p.KD(p.if.ligands,p.if.H)*ones(1,size(Li,2))),1);
pHBpK = - H_x.*sum(Li./Pm_L2 ...
    ./(p.KD(p.if.ligands,p.if.H).*p.KD(p.if.ligands,p.if.K)*ones(1,size(Li,2))),1);
pHBpNa = - H_x.*sum(Li./Pm_L2 ...
    ./(p.KD(p.if.ligands,p.if.H).*p.KD(p.if.ligands,p.if.Na)*ones(1,size(Li,2))),1);
pHBpMg = - H_x.*sum(Li./Pm_L2 ...
    ./(p.KD(p.if.ligands,p.if.H).*p.KD(p.if.ligands,p.if.Mg)*ones(1,size(Li,2))),1);
pHBpCa = - H_x.*sum(Li./Pm_L2 ...
    ./(p.KD(p.if.ligands,p.if.H).*p.KD(p.if.ligands,p.if.Ca)*ones(1,size(Li,2))),1);

pKBpH = - K_x.*sum(Li./Pm_L2 ...
    ./(p.KD(p.if.ligands,p.if.K).*p.KD(p.if.ligands,p.if.H)*ones(1,size(Li,2))),1);
pKBpK = + sum(Pm_KBK.*Li./Pm_L2./(p.KD(p.if.ligands,p.if.K)*ones(1,size(Li,2))),1);
pKBpNa = - K_x.*sum(Li./Pm_L2 ...
    ./(p.KD(p.if.ligands,p.if.K).*p.KD(p.if.ligands,p.if.Na)*ones(1,size(Li,2))),1);
pKBpMg = - K_x.*sum(Li./Pm_L2 ...
    ./(p.KD(p.if.ligands,p.if.K).*p.KD(p.if.ligands,p.if.Mg)*ones(1,size(Li,2))),1);
pKBpCa = - K_x.*sum(Li./Pm_L2 ...
    ./(p.KD(p.if.ligands,p.if.K).*p.KD(p.if.ligands,p.if.Ca)*ones(1,size(Li,2))),1);

pNaBpH = - Na_x.*sum(Li./Pm_L2 ...
    ./(p.KD(p.if.ligands,p.if.Na).*p.KD(p.if.ligands,p.if.H)*ones(1,size(Li,2))),1);
pNaBpK = - Na_x.*sum(Li./Pm_L2 ...
    ./(p.KD(p.if.ligands,p.if.Na).*p.KD(p.if.ligands,p.if.K)*ones(1,size(Li,2))),1);
pNaBpNa = + sum(Pm_NBN.*Li./Pm_L2./(p.KD(p.if.ligands,p.if.Na)*ones(1,size(Li,2))),1);
pNaBpMg = - Na_x.*sum(Li./Pm_L2 ...
    ./(p.KD(p.if.ligands,p.if.Na).*p.KD(p.if.ligands,p.if.Mg)*ones(1,size(Li,2))),1);
pNaBpCa = - Na_x.*sum(Li./Pm_L2 ...
    ./(p.KD(p.if.ligands,p.if.Na).*p.KD(p.if.ligands,p.if.Ca)*ones(1,size(Li,2))),1);

pMgBpH = - Mg_x.*sum(Li./Pm_L2 ...
    ./(p.KD(p.if.ligands,p.if.Mg).*p.KD(p.if.ligands,p.if.H)*ones(1,size(Li,2))),1);
pMgBpK = - Mg_x.*sum(Li./Pm_L2 ...
    ./(p.KD(p.if.ligands,p.if.Mg).*p.KD(p.if.ligands,p.if.K)*ones(1,size(Li,2))),1);
pMgBpNa = - Mg_x.*sum(Li./Pm_L2 ...
    ./(p.KD(p.if.ligands,p.if.Mg).*p.KD(p.if.ligands,p.if.Na)*ones(1,size(Li,2))),1);
pMgBpMg = + sum(Pm_MBM.*Li./Pm_L2./(p.KD(p.if.ligands,p.if.Mg)*ones(1,size(Li,2))),1);
pMgBpCa = - Mg_x.*sum(Li./Pm_L2 ...
    ./(p.KD(p.if.ligands,p.if.Mg).*p.KD(p.if.ligands,p.if.Ca)*ones(1,size(Li,2))),1);

pCaBpH = - Ca_x.*sum(Li./Pm_L2 ...
    ./(p.KD(p.if.ligands,p.if.Ca).*p.KD(p.if.ligands,p.if.H)*ones(1,size(Li,2))),1);
pCaBpK = - Ca_x.*sum(Li./Pm_L2 ...
    ./(p.KD(p.if.ligands,p.if.Ca).*p.KD(p.if.ligands,p.if.K)*ones(1,size(Li,2))),1);
pCaBpNa = - Ca_x.*sum(Li./Pm_L2 ...
    ./(p.KD(p.if.ligands,p.if.Ca).*p.KD(p.if.ligands,p.if.Na)*ones(1,size(Li,2))),1);
pCaBpMg = - Ca_x.*sum(Li./Pm_L2 ...
    ./(p.KD(p.if.ligands,p.if.Ca).*p.KD(p.if.ligands,p.if.Mg)*ones(1,size(Li,2))),1);
pCaBpCa = + sum(Pm_CBC.*Li./Pm_L2 ...
    ./(p.KD(p.if.ligands,p.if.Ca)*ones(1,size(Li,2))),1);

%% PHI's:
PhiH = - H_x.*sum(dLidt./Pm_L ...
    ./(p.KD(p.if.ligands,p.if.H)*ones(1,size(Li,2)))) ...
    + ((1).*(J.MitoDH) + 1*J.MCT + 1*J.MDH - 5*J.CI.JNADH ...
    - 1*J.CIII.Jc3 - 4*J.CIV + (p.F1FO.nH-1)*J.F1FO + 2*J.PiC ...
    + J.mHleak + J.mKHE + J.mNHE)/p.Vol.H2O_mtx ...
    + 2*(p.CaPi.n - p.CaPi.m)*J.CaPi;
PhiK = - K_x.*sum(dLidt./Pm_L ...
    ./(p.KD(p.if.ligands,p.if.K)*ones(1,size(Li,2)))) ...
    + (-J.mKHE)/p.Vol.H2O_mtx;
PhiNa = - Na_x.*sum(dLidt./Pm_L ...
    ./(p.KD(p.if.ligands,p.if.Na)*ones(1,size(Li,2))))...
    + (3*J.NCLX - J.mNHE)/p.Vol.H2O_mtx;
PhiMg = - Mg_x.*sum(dLidt./Pm_L ...
    ./(p.KD(p.if.ligands,p.if.Mg)*ones(1,size(Li,2))));
PhiCa = - Ca_x.*sum(dLidt./Pm_L ...
    ./(p.KD(p.if.ligands,p.if.Ca)*ones(1,size(Li,2)))) ...
    + (J.MCU - J.NCLX)/p.Vol.H2O_mtx - p.CaPi.n*J.CaPi;


%% Endogenous buffering for [H+], [K+], [Na+], [Mg2+], [Ca2+], [Pi]
BH = p.Buff.Htot;       % M
K_BH = p.Buff.KD_H;     % M

BK = p.Buff.Ktot;       % M
K_BK = p.Buff.KD_K;     % M

BNa = p.Buff.Natot;     % M
K_BNa = p.Buff.KD_Na;   % M

BMg = p.Buff.Mgtot;     % M
K_BMg = p.Buff.KD_Mg;   % M

BCa = p.Buff.BCa;      % M
K_BCa = p.Buff.K_BCa;  % M

%% Buffering Terms:
DH = 1 + pHBpH + BH./K_BH./(1+H_x./K_BH).^2;
DK = 1 + pKBpK + BK./K_BK./(1+K_x./K_BK).^2;
DNa = 1 + pNaBpNa + BNa./K_BNa./(1+Na_x./K_BNa).^2;
DMg = 1 + pMgBpMg + BMg./K_BMg./(1+Mg_x./K_BMg).^2;
DCa = 1 + pCaBpCa + BCa./K_BCa./(1+Ca_x./K_BCa).^2;

%% Solution to linear cation eqns.
D=(DH.*DK.*DMg.*pCaBpNa.*pNaBpCa + DH.*DK.*DNa.*pCaBpMg.*pMgBpCa + DCa.*DH.*DK.*pMgBpNa.*pNaBpMg + DK.*DMg.*DNa.*pCaBpH.*pHBpCa + DH.*DMg.*DNa.*pCaBpK.*pKBpCa + DCa.*DK.*DMg.*pHBpNa.*pNaBpH + DCa.*DH.*DMg.*pKBpNa.*pNaBpK + DCa.*DK.*DNa.*pHBpMg.*pMgBpH + DCa.*DH.*DNa.*pKBpMg.*pMgBpK + DCa.*DMg.*DNa.*pHBpK.*pKBpH - DH.*DK.*pCaBpNa.*pMgBpCa.*pNaBpMg - DH.*DK.*pCaBpMg.*pMgBpNa.*pNaBpCa - DK.*DMg.*pCaBpH.*pHBpNa.*pNaBpCa - DK.*DMg.*pCaBpNa.*pHBpCa.*pNaBpH - DH.*DMg.*pCaBpK.*pKBpNa.*pNaBpCa - DH.*DMg.*pCaBpNa.*pKBpCa.*pNaBpK - DK.*DNa.*pCaBpH.*pHBpMg.*pMgBpCa - DK.*DNa.*pCaBpMg.*pHBpCa.*pMgBpH - DH.*DNa.*pCaBpK.*pKBpMg.*pMgBpCa - DH.*DNa.*pCaBpMg.*pKBpCa.*pMgBpK - DCa.*DK.*pHBpNa.*pMgBpH.*pNaBpMg - DCa.*DK.*pHBpMg.*pMgBpNa.*pNaBpH - DCa.*DH.*pKBpNa.*pMgBpK.*pNaBpMg - DCa.*DH.*pKBpMg.*pMgBpNa.*pNaBpK - DMg.*DNa.*pCaBpH.*pHBpK.*pKBpCa - DMg.*DNa.*pCaBpK.*pHBpCa.*pKBpH - DCa.*DMg.*pHBpK.*pKBpNa.*pNaBpH - DCa.*DMg.*pHBpNa.*pKBpH.*pNaBpK - DCa.*DNa.*pHBpK.*pKBpMg.*pMgBpH - DCa.*DNa.*pHBpMg.*pKBpH.*pMgBpK - DK.*pCaBpH.*pHBpCa.*pMgBpNa.*pNaBpMg + DK.*pCaBpH.*pHBpNa.*pMgBpCa.*pNaBpMg + DK.*pCaBpH.*pHBpMg.*pMgBpNa.*pNaBpCa + DK.*pCaBpNa.*pHBpCa.*pMgBpH.*pNaBpMg - DK.*pCaBpNa.*pHBpMg.*pMgBpH.*pNaBpCa + DK.*pCaBpNa.*pHBpMg.*pMgBpCa.*pNaBpH + DK.*pCaBpMg.*pHBpCa.*pMgBpNa.*pNaBpH + DK.*pCaBpMg.*pHBpNa.*pMgBpH.*pNaBpCa - DK.*pCaBpMg.*pHBpNa.*pMgBpCa.*pNaBpH - DH.*pCaBpK.*pKBpCa.*pMgBpNa.*pNaBpMg + DH.*pCaBpK.*pKBpNa.*pMgBpCa.*pNaBpMg + DH.*pCaBpK.*pKBpMg.*pMgBpNa.*pNaBpCa + DH.*pCaBpNa.*pKBpCa.*pMgBpK.*pNaBpMg - DH.*pCaBpNa.*pKBpMg.*pMgBpK.*pNaBpCa + DH.*pCaBpNa.*pKBpMg.*pMgBpCa.*pNaBpK + DH.*pCaBpMg.*pKBpCa.*pMgBpNa.*pNaBpK + DH.*pCaBpMg.*pKBpNa.*pMgBpK.*pNaBpCa - DH.*pCaBpMg.*pKBpNa.*pMgBpCa.*pNaBpK + DMg.*pCaBpH.*pHBpK.*pKBpNa.*pNaBpCa - DMg.*pCaBpH.*pHBpCa.*pKBpNa.*pNaBpK + DMg.*pCaBpH.*pHBpNa.*pKBpCa.*pNaBpK + DMg.*pCaBpK.*pHBpCa.*pKBpNa.*pNaBpH + DMg.*pCaBpK.*pHBpNa.*pKBpH.*pNaBpCa - DMg.*pCaBpK.*pHBpNa.*pKBpCa.*pNaBpH - DMg.*pCaBpNa.*pHBpK.*pKBpH.*pNaBpCa + DMg.*pCaBpNa.*pHBpK.*pKBpCa.*pNaBpH + DMg.*pCaBpNa.*pHBpCa.*pKBpH.*pNaBpK + DNa.*pCaBpH.*pHBpK.*pKBpMg.*pMgBpCa - DNa.*pCaBpH.*pHBpCa.*pKBpMg.*pMgBpK + DNa.*pCaBpH.*pHBpMg.*pKBpCa.*pMgBpK + DNa.*pCaBpK.*pHBpCa.*pKBpMg.*pMgBpH + DNa.*pCaBpK.*pHBpMg.*pKBpH.*pMgBpCa - DNa.*pCaBpK.*pHBpMg.*pKBpCa.*pMgBpH - DNa.*pCaBpMg.*pHBpK.*pKBpH.*pMgBpCa + DNa.*pCaBpMg.*pHBpK.*pKBpCa.*pMgBpH + DNa.*pCaBpMg.*pHBpCa.*pKBpH.*pMgBpK - DCa.*pHBpK.*pKBpH.*pMgBpNa.*pNaBpMg + DCa.*pHBpK.*pKBpNa.*pMgBpH.*pNaBpMg + DCa.*pHBpK.*pKBpMg.*pMgBpNa.*pNaBpH + DCa.*pHBpNa.*pKBpH.*pMgBpK.*pNaBpMg + DCa.*pHBpNa.*pKBpMg.*pMgBpH.*pNaBpK - DCa.*pHBpNa.*pKBpMg.*pMgBpK.*pNaBpH + DCa.*pHBpMg.*pKBpH.*pMgBpNa.*pNaBpK - DCa.*pHBpMg.*pKBpNa.*pMgBpH.*pNaBpK + DCa.*pHBpMg.*pKBpNa.*pMgBpK.*pNaBpH + pCaBpH.*pHBpK.*pKBpCa.*pMgBpNa.*pNaBpMg - pCaBpH.*pHBpK.*pKBpNa.*pMgBpCa.*pNaBpMg - pCaBpH.*pHBpK.*pKBpMg.*pMgBpNa.*pNaBpCa + pCaBpH.*pHBpCa.*pKBpNa.*pMgBpK.*pNaBpMg + pCaBpH.*pHBpCa.*pKBpMg.*pMgBpNa.*pNaBpK - pCaBpH.*pHBpNa.*pKBpCa.*pMgBpK.*pNaBpMg + pCaBpH.*pHBpNa.*pKBpMg.*pMgBpK.*pNaBpCa - pCaBpH.*pHBpNa.*pKBpMg.*pMgBpCa.*pNaBpK - pCaBpH.*pHBpMg.*pKBpCa.*pMgBpNa.*pNaBpK - pCaBpH.*pHBpMg.*pKBpNa.*pMgBpK.*pNaBpCa + pCaBpH.*pHBpMg.*pKBpNa.*pMgBpCa.*pNaBpK + pCaBpK.*pHBpCa.*pKBpH.*pMgBpNa.*pNaBpMg - pCaBpK.*pHBpCa.*pKBpNa.*pMgBpH.*pNaBpMg - pCaBpK.*pHBpCa.*pKBpMg.*pMgBpNa.*pNaBpH - pCaBpK.*pHBpNa.*pKBpH.*pMgBpCa.*pNaBpMg + pCaBpK.*pHBpNa.*pKBpCa.*pMgBpH.*pNaBpMg - pCaBpK.*pHBpNa.*pKBpMg.*pMgBpH.*pNaBpCa + pCaBpK.*pHBpNa.*pKBpMg.*pMgBpCa.*pNaBpH - pCaBpK.*pHBpMg.*pKBpH.*pMgBpNa.*pNaBpCa + pCaBpK.*pHBpMg.*pKBpCa.*pMgBpNa.*pNaBpH + pCaBpK.*pHBpMg.*pKBpNa.*pMgBpH.*pNaBpCa - pCaBpK.*pHBpMg.*pKBpNa.*pMgBpCa.*pNaBpH + pCaBpNa.*pHBpK.*pKBpH.*pMgBpCa.*pNaBpMg - pCaBpNa.*pHBpK.*pKBpCa.*pMgBpH.*pNaBpMg + pCaBpNa.*pHBpK.*pKBpMg.*pMgBpH.*pNaBpCa - pCaBpNa.*pHBpK.*pKBpMg.*pMgBpCa.*pNaBpH - pCaBpNa.*pHBpCa.*pKBpH.*pMgBpK.*pNaBpMg - pCaBpNa.*pHBpCa.*pKBpMg.*pMgBpH.*pNaBpK + pCaBpNa.*pHBpCa.*pKBpMg.*pMgBpK.*pNaBpH + pCaBpNa.*pHBpMg.*pKBpH.*pMgBpK.*pNaBpCa - pCaBpNa.*pHBpMg.*pKBpH.*pMgBpCa.*pNaBpK + pCaBpNa.*pHBpMg.*pKBpCa.*pMgBpH.*pNaBpK - pCaBpNa.*pHBpMg.*pKBpCa.*pMgBpK.*pNaBpH + pCaBpMg.*pHBpK.*pKBpH.*pMgBpNa.*pNaBpCa - pCaBpMg.*pHBpK.*pKBpCa.*pMgBpNa.*pNaBpH - pCaBpMg.*pHBpK.*pKBpNa.*pMgBpH.*pNaBpCa + pCaBpMg.*pHBpK.*pKBpNa.*pMgBpCa.*pNaBpH - pCaBpMg.*pHBpCa.*pKBpH.*pMgBpNa.*pNaBpK + pCaBpMg.*pHBpCa.*pKBpNa.*pMgBpH.*pNaBpK - pCaBpMg.*pHBpCa.*pKBpNa.*pMgBpK.*pNaBpH - pCaBpMg.*pHBpNa.*pKBpH.*pMgBpK.*pNaBpCa + pCaBpMg.*pHBpNa.*pKBpH.*pMgBpCa.*pNaBpK - pCaBpMg.*pHBpNa.*pKBpCa.*pMgBpH.*pNaBpK + pCaBpMg.*pHBpNa.*pKBpCa.*pMgBpK.*pNaBpH - DCa.*DH.*DK.*DMg.*DNa);
dHdt=-(DK.*DMg.*PhiNa.*pCaBpNa.*pHBpCa + DCa.*DK.*PhiNa.*pHBpMg.*pMgBpNa + DCa.*DMg.*PhiNa.*pHBpK.*pKBpNa + DK.*DNa.*PhiMg.*pCaBpMg.*pHBpCa + DCa.*DK.*PhiMg.*pHBpNa.*pNaBpMg + DCa.*DNa.*PhiMg.*pHBpK.*pKBpMg + DMg.*DNa.*PhiK.*pCaBpK.*pHBpCa + DCa.*DMg.*PhiK.*pHBpNa.*pNaBpK + DCa.*DNa.*PhiK.*pHBpMg.*pMgBpK - DK.*DMg.*PhiH.*pCaBpNa.*pNaBpCa - DK.*DNa.*PhiH.*pCaBpMg.*pMgBpCa - DCa.*DK.*PhiH.*pMgBpNa.*pNaBpMg - DMg.*DNa.*PhiH.*pCaBpK.*pKBpCa - DCa.*DMg.*PhiH.*pKBpNa.*pNaBpK - DCa.*DNa.*PhiH.*pKBpMg.*pMgBpK + DK.*DMg.*PhiCa.*pHBpNa.*pNaBpCa + DK.*DNa.*PhiCa.*pHBpMg.*pMgBpCa + DMg.*DNa.*PhiCa.*pHBpK.*pKBpCa - DK.*PhiNa.*pCaBpNa.*pHBpMg.*pMgBpCa - DK.*PhiNa.*pCaBpMg.*pHBpCa.*pMgBpNa + DK.*PhiNa.*pCaBpMg.*pHBpNa.*pMgBpCa - DMg.*PhiNa.*pCaBpK.*pHBpCa.*pKBpNa + DMg.*PhiNa.*pCaBpK.*pHBpNa.*pKBpCa - DMg.*PhiNa.*pCaBpNa.*pHBpK.*pKBpCa - DCa.*PhiNa.*pHBpK.*pKBpMg.*pMgBpNa + DCa.*PhiNa.*pHBpNa.*pKBpMg.*pMgBpK - DCa.*PhiNa.*pHBpMg.*pKBpNa.*pMgBpK - DK.*PhiMg.*pCaBpNa.*pHBpCa.*pNaBpMg + DK.*PhiMg.*pCaBpNa.*pHBpMg.*pNaBpCa - DK.*PhiMg.*pCaBpMg.*pHBpNa.*pNaBpCa - DNa.*PhiMg.*pCaBpK.*pHBpCa.*pKBpMg + DNa.*PhiMg.*pCaBpK.*pHBpMg.*pKBpCa - DNa.*PhiMg.*pCaBpMg.*pHBpK.*pKBpCa - DCa.*PhiMg.*pHBpK.*pKBpNa.*pNaBpMg - DCa.*PhiMg.*pHBpNa.*pKBpMg.*pNaBpK + DCa.*PhiMg.*pHBpMg.*pKBpNa.*pNaBpK - DMg.*PhiK.*pCaBpK.*pHBpNa.*pNaBpCa + DMg.*PhiK.*pCaBpNa.*pHBpK.*pNaBpCa - DMg.*PhiK.*pCaBpNa.*pHBpCa.*pNaBpK - DNa.*PhiK.*pCaBpK.*pHBpMg.*pMgBpCa + DNa.*PhiK.*pCaBpMg.*pHBpK.*pMgBpCa - DNa.*PhiK.*pCaBpMg.*pHBpCa.*pMgBpK + DCa.*PhiK.*pHBpK.*pMgBpNa.*pNaBpMg - DCa.*PhiK.*pHBpNa.*pMgBpK.*pNaBpMg - DCa.*PhiK.*pHBpMg.*pMgBpNa.*pNaBpK + DK.*PhiH.*pCaBpNa.*pMgBpCa.*pNaBpMg + DK.*PhiH.*pCaBpMg.*pMgBpNa.*pNaBpCa + DMg.*PhiH.*pCaBpK.*pKBpNa.*pNaBpCa + DMg.*PhiH.*pCaBpNa.*pKBpCa.*pNaBpK + DNa.*PhiH.*pCaBpK.*pKBpMg.*pMgBpCa + DNa.*PhiH.*pCaBpMg.*pKBpCa.*pMgBpK + DCa.*PhiH.*pKBpNa.*pMgBpK.*pNaBpMg + DCa.*PhiH.*pKBpMg.*pMgBpNa.*pNaBpK + DK.*PhiCa.*pHBpCa.*pMgBpNa.*pNaBpMg - DK.*PhiCa.*pHBpNa.*pMgBpCa.*pNaBpMg - DK.*PhiCa.*pHBpMg.*pMgBpNa.*pNaBpCa - DMg.*PhiCa.*pHBpK.*pKBpNa.*pNaBpCa + DMg.*PhiCa.*pHBpCa.*pKBpNa.*pNaBpK - DMg.*PhiCa.*pHBpNa.*pKBpCa.*pNaBpK - DNa.*PhiCa.*pHBpK.*pKBpMg.*pMgBpCa + DNa.*PhiCa.*pHBpCa.*pKBpMg.*pMgBpK - DNa.*PhiCa.*pHBpMg.*pKBpCa.*pMgBpK + PhiNa.*pCaBpK.*pHBpCa.*pKBpMg.*pMgBpNa - PhiNa.*pCaBpK.*pHBpNa.*pKBpMg.*pMgBpCa - PhiNa.*pCaBpK.*pHBpMg.*pKBpCa.*pMgBpNa + PhiNa.*pCaBpK.*pHBpMg.*pKBpNa.*pMgBpCa + PhiNa.*pCaBpNa.*pHBpK.*pKBpMg.*pMgBpCa - PhiNa.*pCaBpNa.*pHBpCa.*pKBpMg.*pMgBpK + PhiNa.*pCaBpNa.*pHBpMg.*pKBpCa.*pMgBpK + PhiNa.*pCaBpMg.*pHBpK.*pKBpCa.*pMgBpNa - PhiNa.*pCaBpMg.*pHBpK.*pKBpNa.*pMgBpCa + PhiNa.*pCaBpMg.*pHBpCa.*pKBpNa.*pMgBpK - PhiNa.*pCaBpMg.*pHBpNa.*pKBpCa.*pMgBpK + PhiMg.*pCaBpK.*pHBpCa.*pKBpNa.*pNaBpMg - PhiMg.*pCaBpK.*pHBpNa.*pKBpCa.*pNaBpMg + PhiMg.*pCaBpK.*pHBpNa.*pKBpMg.*pNaBpCa - PhiMg.*pCaBpK.*pHBpMg.*pKBpNa.*pNaBpCa + PhiMg.*pCaBpNa.*pHBpK.*pKBpCa.*pNaBpMg - PhiMg.*pCaBpNa.*pHBpK.*pKBpMg.*pNaBpCa + PhiMg.*pCaBpNa.*pHBpCa.*pKBpMg.*pNaBpK - PhiMg.*pCaBpNa.*pHBpMg.*pKBpCa.*pNaBpK + PhiMg.*pCaBpMg.*pHBpK.*pKBpNa.*pNaBpCa - PhiMg.*pCaBpMg.*pHBpCa.*pKBpNa.*pNaBpK + PhiMg.*pCaBpMg.*pHBpNa.*pKBpCa.*pNaBpK - PhiK.*pCaBpK.*pHBpCa.*pMgBpNa.*pNaBpMg + PhiK.*pCaBpK.*pHBpNa.*pMgBpCa.*pNaBpMg + PhiK.*pCaBpK.*pHBpMg.*pMgBpNa.*pNaBpCa - PhiK.*pCaBpNa.*pHBpK.*pMgBpCa.*pNaBpMg + PhiK.*pCaBpNa.*pHBpCa.*pMgBpK.*pNaBpMg - PhiK.*pCaBpNa.*pHBpMg.*pMgBpK.*pNaBpCa + PhiK.*pCaBpNa.*pHBpMg.*pMgBpCa.*pNaBpK - PhiK.*pCaBpMg.*pHBpK.*pMgBpNa.*pNaBpCa + PhiK.*pCaBpMg.*pHBpCa.*pMgBpNa.*pNaBpK + PhiK.*pCaBpMg.*pHBpNa.*pMgBpK.*pNaBpCa - PhiK.*pCaBpMg.*pHBpNa.*pMgBpCa.*pNaBpK + PhiH.*pCaBpK.*pKBpCa.*pMgBpNa.*pNaBpMg - PhiH.*pCaBpK.*pKBpNa.*pMgBpCa.*pNaBpMg - PhiH.*pCaBpK.*pKBpMg.*pMgBpNa.*pNaBpCa - PhiH.*pCaBpNa.*pKBpCa.*pMgBpK.*pNaBpMg + PhiH.*pCaBpNa.*pKBpMg.*pMgBpK.*pNaBpCa - PhiH.*pCaBpNa.*pKBpMg.*pMgBpCa.*pNaBpK - PhiH.*pCaBpMg.*pKBpCa.*pMgBpNa.*pNaBpK - PhiH.*pCaBpMg.*pKBpNa.*pMgBpK.*pNaBpCa + PhiH.*pCaBpMg.*pKBpNa.*pMgBpCa.*pNaBpK - PhiCa.*pHBpK.*pKBpCa.*pMgBpNa.*pNaBpMg + PhiCa.*pHBpK.*pKBpNa.*pMgBpCa.*pNaBpMg + PhiCa.*pHBpK.*pKBpMg.*pMgBpNa.*pNaBpCa - PhiCa.*pHBpCa.*pKBpNa.*pMgBpK.*pNaBpMg - PhiCa.*pHBpCa.*pKBpMg.*pMgBpNa.*pNaBpK + PhiCa.*pHBpNa.*pKBpCa.*pMgBpK.*pNaBpMg - PhiCa.*pHBpNa.*pKBpMg.*pMgBpK.*pNaBpCa + PhiCa.*pHBpNa.*pKBpMg.*pMgBpCa.*pNaBpK + PhiCa.*pHBpMg.*pKBpCa.*pMgBpNa.*pNaBpK + PhiCa.*pHBpMg.*pKBpNa.*pMgBpK.*pNaBpCa - PhiCa.*pHBpMg.*pKBpNa.*pMgBpCa.*pNaBpK + DCa.*DK.*DMg.*DNa.*PhiH - DCa.*DK.*DMg.*PhiNa.*pHBpNa - DCa.*DK.*DNa.*PhiMg.*pHBpMg - DCa.*DMg.*DNa.*PhiK.*pHBpK - DK.*DMg.*DNa.*PhiCa.*pHBpCa)./D;
dNadt=(DH.*DK.*PhiNa.*pCaBpMg.*pMgBpCa + DK.*DMg.*PhiNa.*pCaBpH.*pHBpCa + DH.*DMg.*PhiNa.*pCaBpK.*pKBpCa + DCa.*DK.*PhiNa.*pHBpMg.*pMgBpH + DCa.*DH.*PhiNa.*pKBpMg.*pMgBpK + DCa.*DMg.*PhiNa.*pHBpK.*pKBpH - DH.*DK.*PhiMg.*pCaBpMg.*pNaBpCa - DCa.*DK.*PhiMg.*pHBpMg.*pNaBpH - DCa.*DH.*PhiMg.*pKBpMg.*pNaBpK - DH.*DMg.*PhiK.*pCaBpK.*pNaBpCa - DCa.*DH.*PhiK.*pMgBpK.*pNaBpMg - DCa.*DMg.*PhiK.*pHBpK.*pNaBpH - DK.*DMg.*PhiH.*pCaBpH.*pNaBpCa - DCa.*DK.*PhiH.*pMgBpH.*pNaBpMg - DCa.*DMg.*PhiH.*pKBpH.*pNaBpK - DH.*DK.*PhiCa.*pMgBpCa.*pNaBpMg - DK.*DMg.*PhiCa.*pHBpCa.*pNaBpH - DH.*DMg.*PhiCa.*pKBpCa.*pNaBpK - DK.*PhiNa.*pCaBpH.*pHBpMg.*pMgBpCa - DK.*PhiNa.*pCaBpMg.*pHBpCa.*pMgBpH - DH.*PhiNa.*pCaBpK.*pKBpMg.*pMgBpCa - DH.*PhiNa.*pCaBpMg.*pKBpCa.*pMgBpK - DMg.*PhiNa.*pCaBpH.*pHBpK.*pKBpCa - DMg.*PhiNa.*pCaBpK.*pHBpCa.*pKBpH - DCa.*PhiNa.*pHBpK.*pKBpMg.*pMgBpH - DCa.*PhiNa.*pHBpMg.*pKBpH.*pMgBpK - DK.*PhiMg.*pCaBpH.*pHBpCa.*pNaBpMg + DK.*PhiMg.*pCaBpH.*pHBpMg.*pNaBpCa + DK.*PhiMg.*pCaBpMg.*pHBpCa.*pNaBpH - DH.*PhiMg.*pCaBpK.*pKBpCa.*pNaBpMg + DH.*PhiMg.*pCaBpK.*pKBpMg.*pNaBpCa + DH.*PhiMg.*pCaBpMg.*pKBpCa.*pNaBpK - DCa.*PhiMg.*pHBpK.*pKBpH.*pNaBpMg + DCa.*PhiMg.*pHBpK.*pKBpMg.*pNaBpH + DCa.*PhiMg.*pHBpMg.*pKBpH.*pNaBpK + DH.*PhiK.*pCaBpK.*pMgBpCa.*pNaBpMg + DH.*PhiK.*pCaBpMg.*pMgBpK.*pNaBpCa - DH.*PhiK.*pCaBpMg.*pMgBpCa.*pNaBpK + DMg.*PhiK.*pCaBpH.*pHBpK.*pNaBpCa - DMg.*PhiK.*pCaBpH.*pHBpCa.*pNaBpK + DMg.*PhiK.*pCaBpK.*pHBpCa.*pNaBpH + DCa.*PhiK.*pHBpK.*pMgBpH.*pNaBpMg - DCa.*PhiK.*pHBpMg.*pMgBpH.*pNaBpK + DCa.*PhiK.*pHBpMg.*pMgBpK.*pNaBpH + DK.*PhiH.*pCaBpH.*pMgBpCa.*pNaBpMg + DK.*PhiH.*pCaBpMg.*pMgBpH.*pNaBpCa - DK.*PhiH.*pCaBpMg.*pMgBpCa.*pNaBpH + DMg.*PhiH.*pCaBpH.*pKBpCa.*pNaBpK + DMg.*PhiH.*pCaBpK.*pKBpH.*pNaBpCa - DMg.*PhiH.*pCaBpK.*pKBpCa.*pNaBpH + DCa.*PhiH.*pKBpH.*pMgBpK.*pNaBpMg + DCa.*PhiH.*pKBpMg.*pMgBpH.*pNaBpK - DCa.*PhiH.*pKBpMg.*pMgBpK.*pNaBpH + DK.*PhiCa.*pHBpCa.*pMgBpH.*pNaBpMg - DK.*PhiCa.*pHBpMg.*pMgBpH.*pNaBpCa + DK.*PhiCa.*pHBpMg.*pMgBpCa.*pNaBpH + DH.*PhiCa.*pKBpCa.*pMgBpK.*pNaBpMg - DH.*PhiCa.*pKBpMg.*pMgBpK.*pNaBpCa + DH.*PhiCa.*pKBpMg.*pMgBpCa.*pNaBpK - DMg.*PhiCa.*pHBpK.*pKBpH.*pNaBpCa + DMg.*PhiCa.*pHBpK.*pKBpCa.*pNaBpH + DMg.*PhiCa.*pHBpCa.*pKBpH.*pNaBpK + PhiNa.*pCaBpH.*pHBpK.*pKBpMg.*pMgBpCa - PhiNa.*pCaBpH.*pHBpCa.*pKBpMg.*pMgBpK + PhiNa.*pCaBpH.*pHBpMg.*pKBpCa.*pMgBpK + PhiNa.*pCaBpK.*pHBpCa.*pKBpMg.*pMgBpH + PhiNa.*pCaBpK.*pHBpMg.*pKBpH.*pMgBpCa - PhiNa.*pCaBpK.*pHBpMg.*pKBpCa.*pMgBpH - PhiNa.*pCaBpMg.*pHBpK.*pKBpH.*pMgBpCa + PhiNa.*pCaBpMg.*pHBpK.*pKBpCa.*pMgBpH + PhiNa.*pCaBpMg.*pHBpCa.*pKBpH.*pMgBpK + PhiMg.*pCaBpH.*pHBpK.*pKBpCa.*pNaBpMg - PhiMg.*pCaBpH.*pHBpK.*pKBpMg.*pNaBpCa + PhiMg.*pCaBpH.*pHBpCa.*pKBpMg.*pNaBpK - PhiMg.*pCaBpH.*pHBpMg.*pKBpCa.*pNaBpK + PhiMg.*pCaBpK.*pHBpCa.*pKBpH.*pNaBpMg - PhiMg.*pCaBpK.*pHBpCa.*pKBpMg.*pNaBpH - PhiMg.*pCaBpK.*pHBpMg.*pKBpH.*pNaBpCa + PhiMg.*pCaBpK.*pHBpMg.*pKBpCa.*pNaBpH + PhiMg.*pCaBpMg.*pHBpK.*pKBpH.*pNaBpCa - PhiMg.*pCaBpMg.*pHBpK.*pKBpCa.*pNaBpH - PhiMg.*pCaBpMg.*pHBpCa.*pKBpH.*pNaBpK - PhiK.*pCaBpH.*pHBpK.*pMgBpCa.*pNaBpMg + PhiK.*pCaBpH.*pHBpCa.*pMgBpK.*pNaBpMg - PhiK.*pCaBpH.*pHBpMg.*pMgBpK.*pNaBpCa + PhiK.*pCaBpH.*pHBpMg.*pMgBpCa.*pNaBpK - PhiK.*pCaBpK.*pHBpCa.*pMgBpH.*pNaBpMg + PhiK.*pCaBpK.*pHBpMg.*pMgBpH.*pNaBpCa - PhiK.*pCaBpK.*pHBpMg.*pMgBpCa.*pNaBpH - PhiK.*pCaBpMg.*pHBpK.*pMgBpH.*pNaBpCa + PhiK.*pCaBpMg.*pHBpK.*pMgBpCa.*pNaBpH + PhiK.*pCaBpMg.*pHBpCa.*pMgBpH.*pNaBpK - PhiK.*pCaBpMg.*pHBpCa.*pMgBpK.*pNaBpH - PhiH.*pCaBpH.*pKBpCa.*pMgBpK.*pNaBpMg + PhiH.*pCaBpH.*pKBpMg.*pMgBpK.*pNaBpCa - PhiH.*pCaBpH.*pKBpMg.*pMgBpCa.*pNaBpK - PhiH.*pCaBpK.*pKBpH.*pMgBpCa.*pNaBpMg + PhiH.*pCaBpK.*pKBpCa.*pMgBpH.*pNaBpMg - PhiH.*pCaBpK.*pKBpMg.*pMgBpH.*pNaBpCa + PhiH.*pCaBpK.*pKBpMg.*pMgBpCa.*pNaBpH - PhiH.*pCaBpMg.*pKBpH.*pMgBpK.*pNaBpCa + PhiH.*pCaBpMg.*pKBpH.*pMgBpCa.*pNaBpK - PhiH.*pCaBpMg.*pKBpCa.*pMgBpH.*pNaBpK + PhiH.*pCaBpMg.*pKBpCa.*pMgBpK.*pNaBpH + PhiCa.*pHBpK.*pKBpH.*pMgBpCa.*pNaBpMg - PhiCa.*pHBpK.*pKBpCa.*pMgBpH.*pNaBpMg + PhiCa.*pHBpK.*pKBpMg.*pMgBpH.*pNaBpCa - PhiCa.*pHBpK.*pKBpMg.*pMgBpCa.*pNaBpH - PhiCa.*pHBpCa.*pKBpH.*pMgBpK.*pNaBpMg - PhiCa.*pHBpCa.*pKBpMg.*pMgBpH.*pNaBpK + PhiCa.*pHBpCa.*pKBpMg.*pMgBpK.*pNaBpH + PhiCa.*pHBpMg.*pKBpH.*pMgBpK.*pNaBpCa - PhiCa.*pHBpMg.*pKBpH.*pMgBpCa.*pNaBpK + PhiCa.*pHBpMg.*pKBpCa.*pMgBpH.*pNaBpK - PhiCa.*pHBpMg.*pKBpCa.*pMgBpK.*pNaBpH - DCa.*DH.*DK.*DMg.*PhiNa + DCa.*DH.*DK.*PhiMg.*pNaBpMg + DCa.*DH.*DMg.*PhiK.*pNaBpK + DCa.*DK.*DMg.*PhiH.*pNaBpH + DH.*DK.*DMg.*PhiCa.*pNaBpCa)./D;
dKdt=-(DH.*DMg.*PhiNa.*pCaBpNa.*pKBpCa + DCa.*DH.*PhiNa.*pKBpMg.*pMgBpNa + DCa.*DMg.*PhiNa.*pHBpNa.*pKBpH + DH.*DNa.*PhiMg.*pCaBpMg.*pKBpCa + DCa.*DH.*PhiMg.*pKBpNa.*pNaBpMg + DCa.*DNa.*PhiMg.*pHBpMg.*pKBpH - DH.*DMg.*PhiK.*pCaBpNa.*pNaBpCa - DH.*DNa.*PhiK.*pCaBpMg.*pMgBpCa - DCa.*DH.*PhiK.*pMgBpNa.*pNaBpMg - DMg.*DNa.*PhiK.*pCaBpH.*pHBpCa - DCa.*DMg.*PhiK.*pHBpNa.*pNaBpH - DCa.*DNa.*PhiK.*pHBpMg.*pMgBpH + DMg.*DNa.*PhiH.*pCaBpH.*pKBpCa + DCa.*DMg.*PhiH.*pKBpNa.*pNaBpH + DCa.*DNa.*PhiH.*pKBpMg.*pMgBpH + DH.*DMg.*PhiCa.*pKBpNa.*pNaBpCa + DH.*DNa.*PhiCa.*pKBpMg.*pMgBpCa + DMg.*DNa.*PhiCa.*pHBpCa.*pKBpH - DH.*PhiNa.*pCaBpNa.*pKBpMg.*pMgBpCa - DH.*PhiNa.*pCaBpMg.*pKBpCa.*pMgBpNa + DH.*PhiNa.*pCaBpMg.*pKBpNa.*pMgBpCa + DMg.*PhiNa.*pCaBpH.*pHBpCa.*pKBpNa - DMg.*PhiNa.*pCaBpH.*pHBpNa.*pKBpCa - DMg.*PhiNa.*pCaBpNa.*pHBpCa.*pKBpH - DCa.*PhiNa.*pHBpNa.*pKBpMg.*pMgBpH - DCa.*PhiNa.*pHBpMg.*pKBpH.*pMgBpNa + DCa.*PhiNa.*pHBpMg.*pKBpNa.*pMgBpH - DH.*PhiMg.*pCaBpNa.*pKBpCa.*pNaBpMg + DH.*PhiMg.*pCaBpNa.*pKBpMg.*pNaBpCa - DH.*PhiMg.*pCaBpMg.*pKBpNa.*pNaBpCa + DNa.*PhiMg.*pCaBpH.*pHBpCa.*pKBpMg - DNa.*PhiMg.*pCaBpH.*pHBpMg.*pKBpCa - DNa.*PhiMg.*pCaBpMg.*pHBpCa.*pKBpH - DCa.*PhiMg.*pHBpNa.*pKBpH.*pNaBpMg + DCa.*PhiMg.*pHBpNa.*pKBpMg.*pNaBpH - DCa.*PhiMg.*pHBpMg.*pKBpNa.*pNaBpH + DH.*PhiK.*pCaBpNa.*pMgBpCa.*pNaBpMg + DH.*PhiK.*pCaBpMg.*pMgBpNa.*pNaBpCa + DMg.*PhiK.*pCaBpH.*pHBpNa.*pNaBpCa + DMg.*PhiK.*pCaBpNa.*pHBpCa.*pNaBpH + DNa.*PhiK.*pCaBpH.*pHBpMg.*pMgBpCa + DNa.*PhiK.*pCaBpMg.*pHBpCa.*pMgBpH + DCa.*PhiK.*pHBpNa.*pMgBpH.*pNaBpMg + DCa.*PhiK.*pHBpMg.*pMgBpNa.*pNaBpH - DMg.*PhiH.*pCaBpH.*pKBpNa.*pNaBpCa + DMg.*PhiH.*pCaBpNa.*pKBpH.*pNaBpCa - DMg.*PhiH.*pCaBpNa.*pKBpCa.*pNaBpH - DNa.*PhiH.*pCaBpH.*pKBpMg.*pMgBpCa + DNa.*PhiH.*pCaBpMg.*pKBpH.*pMgBpCa - DNa.*PhiH.*pCaBpMg.*pKBpCa.*pMgBpH + DCa.*PhiH.*pKBpH.*pMgBpNa.*pNaBpMg - DCa.*PhiH.*pKBpNa.*pMgBpH.*pNaBpMg - DCa.*PhiH.*pKBpMg.*pMgBpNa.*pNaBpH + DH.*PhiCa.*pKBpCa.*pMgBpNa.*pNaBpMg - DH.*PhiCa.*pKBpNa.*pMgBpCa.*pNaBpMg - DH.*PhiCa.*pKBpMg.*pMgBpNa.*pNaBpCa - DMg.*PhiCa.*pHBpCa.*pKBpNa.*pNaBpH - DMg.*PhiCa.*pHBpNa.*pKBpH.*pNaBpCa + DMg.*PhiCa.*pHBpNa.*pKBpCa.*pNaBpH - DNa.*PhiCa.*pHBpCa.*pKBpMg.*pMgBpH - DNa.*PhiCa.*pHBpMg.*pKBpH.*pMgBpCa + DNa.*PhiCa.*pHBpMg.*pKBpCa.*pMgBpH - PhiNa.*pCaBpH.*pHBpCa.*pKBpMg.*pMgBpNa + PhiNa.*pCaBpH.*pHBpNa.*pKBpMg.*pMgBpCa + PhiNa.*pCaBpH.*pHBpMg.*pKBpCa.*pMgBpNa - PhiNa.*pCaBpH.*pHBpMg.*pKBpNa.*pMgBpCa + PhiNa.*pCaBpNa.*pHBpCa.*pKBpMg.*pMgBpH + PhiNa.*pCaBpNa.*pHBpMg.*pKBpH.*pMgBpCa - PhiNa.*pCaBpNa.*pHBpMg.*pKBpCa.*pMgBpH + PhiNa.*pCaBpMg.*pHBpCa.*pKBpH.*pMgBpNa - PhiNa.*pCaBpMg.*pHBpCa.*pKBpNa.*pMgBpH - PhiNa.*pCaBpMg.*pHBpNa.*pKBpH.*pMgBpCa + PhiNa.*pCaBpMg.*pHBpNa.*pKBpCa.*pMgBpH - PhiMg.*pCaBpH.*pHBpCa.*pKBpNa.*pNaBpMg + PhiMg.*pCaBpH.*pHBpNa.*pKBpCa.*pNaBpMg - PhiMg.*pCaBpH.*pHBpNa.*pKBpMg.*pNaBpCa + PhiMg.*pCaBpH.*pHBpMg.*pKBpNa.*pNaBpCa + PhiMg.*pCaBpNa.*pHBpCa.*pKBpH.*pNaBpMg - PhiMg.*pCaBpNa.*pHBpCa.*pKBpMg.*pNaBpH - PhiMg.*pCaBpNa.*pHBpMg.*pKBpH.*pNaBpCa + PhiMg.*pCaBpNa.*pHBpMg.*pKBpCa.*pNaBpH + PhiMg.*pCaBpMg.*pHBpCa.*pKBpNa.*pNaBpH + PhiMg.*pCaBpMg.*pHBpNa.*pKBpH.*pNaBpCa - PhiMg.*pCaBpMg.*pHBpNa.*pKBpCa.*pNaBpH + PhiK.*pCaBpH.*pHBpCa.*pMgBpNa.*pNaBpMg - PhiK.*pCaBpH.*pHBpNa.*pMgBpCa.*pNaBpMg - PhiK.*pCaBpH.*pHBpMg.*pMgBpNa.*pNaBpCa - PhiK.*pCaBpNa.*pHBpCa.*pMgBpH.*pNaBpMg + PhiK.*pCaBpNa.*pHBpMg.*pMgBpH.*pNaBpCa - PhiK.*pCaBpNa.*pHBpMg.*pMgBpCa.*pNaBpH - PhiK.*pCaBpMg.*pHBpCa.*pMgBpNa.*pNaBpH - PhiK.*pCaBpMg.*pHBpNa.*pMgBpH.*pNaBpCa + PhiK.*pCaBpMg.*pHBpNa.*pMgBpCa.*pNaBpH - PhiH.*pCaBpH.*pKBpCa.*pMgBpNa.*pNaBpMg + PhiH.*pCaBpH.*pKBpNa.*pMgBpCa.*pNaBpMg + PhiH.*pCaBpH.*pKBpMg.*pMgBpNa.*pNaBpCa - PhiH.*pCaBpNa.*pKBpH.*pMgBpCa.*pNaBpMg + PhiH.*pCaBpNa.*pKBpCa.*pMgBpH.*pNaBpMg - PhiH.*pCaBpNa.*pKBpMg.*pMgBpH.*pNaBpCa + PhiH.*pCaBpNa.*pKBpMg.*pMgBpCa.*pNaBpH - PhiH.*pCaBpMg.*pKBpH.*pMgBpNa.*pNaBpCa + PhiH.*pCaBpMg.*pKBpCa.*pMgBpNa.*pNaBpH + PhiH.*pCaBpMg.*pKBpNa.*pMgBpH.*pNaBpCa - PhiH.*pCaBpMg.*pKBpNa.*pMgBpCa.*pNaBpH - PhiCa.*pHBpCa.*pKBpH.*pMgBpNa.*pNaBpMg + PhiCa.*pHBpCa.*pKBpNa.*pMgBpH.*pNaBpMg + PhiCa.*pHBpCa.*pKBpMg.*pMgBpNa.*pNaBpH + PhiCa.*pHBpNa.*pKBpH.*pMgBpCa.*pNaBpMg - PhiCa.*pHBpNa.*pKBpCa.*pMgBpH.*pNaBpMg + PhiCa.*pHBpNa.*pKBpMg.*pMgBpH.*pNaBpCa - PhiCa.*pHBpNa.*pKBpMg.*pMgBpCa.*pNaBpH + PhiCa.*pHBpMg.*pKBpH.*pMgBpNa.*pNaBpCa - PhiCa.*pHBpMg.*pKBpCa.*pMgBpNa.*pNaBpH - PhiCa.*pHBpMg.*pKBpNa.*pMgBpH.*pNaBpCa + PhiCa.*pHBpMg.*pKBpNa.*pMgBpCa.*pNaBpH + DCa.*DH.*DMg.*DNa.*PhiK - DCa.*DH.*DMg.*PhiNa.*pKBpNa - DCa.*DH.*DNa.*PhiMg.*pKBpMg - DCa.*DMg.*DNa.*PhiH.*pKBpH - DH.*DMg.*DNa.*PhiCa.*pKBpCa)./D;
dMgdt=-(DH.*DK.*PhiNa.*pCaBpNa.*pMgBpCa + DCa.*DK.*PhiNa.*pHBpNa.*pMgBpH + DCa.*DH.*PhiNa.*pKBpNa.*pMgBpK - DH.*DK.*PhiMg.*pCaBpNa.*pNaBpCa - DK.*DNa.*PhiMg.*pCaBpH.*pHBpCa - DH.*DNa.*PhiMg.*pCaBpK.*pKBpCa - DCa.*DK.*PhiMg.*pHBpNa.*pNaBpH - DCa.*DH.*PhiMg.*pKBpNa.*pNaBpK - DCa.*DNa.*PhiMg.*pHBpK.*pKBpH + DH.*DNa.*PhiK.*pCaBpK.*pMgBpCa + DCa.*DH.*PhiK.*pMgBpNa.*pNaBpK + DCa.*DNa.*PhiK.*pHBpK.*pMgBpH + DK.*DNa.*PhiH.*pCaBpH.*pMgBpCa + DCa.*DK.*PhiH.*pMgBpNa.*pNaBpH + DCa.*DNa.*PhiH.*pKBpH.*pMgBpK + DH.*DK.*PhiCa.*pMgBpNa.*pNaBpCa + DK.*DNa.*PhiCa.*pHBpCa.*pMgBpH + DH.*DNa.*PhiCa.*pKBpCa.*pMgBpK + DK.*PhiNa.*pCaBpH.*pHBpCa.*pMgBpNa - DK.*PhiNa.*pCaBpH.*pHBpNa.*pMgBpCa - DK.*PhiNa.*pCaBpNa.*pHBpCa.*pMgBpH + DH.*PhiNa.*pCaBpK.*pKBpCa.*pMgBpNa - DH.*PhiNa.*pCaBpK.*pKBpNa.*pMgBpCa - DH.*PhiNa.*pCaBpNa.*pKBpCa.*pMgBpK + DCa.*PhiNa.*pHBpK.*pKBpH.*pMgBpNa - DCa.*PhiNa.*pHBpK.*pKBpNa.*pMgBpH - DCa.*PhiNa.*pHBpNa.*pKBpH.*pMgBpK + DK.*PhiMg.*pCaBpH.*pHBpNa.*pNaBpCa + DK.*PhiMg.*pCaBpNa.*pHBpCa.*pNaBpH + DH.*PhiMg.*pCaBpK.*pKBpNa.*pNaBpCa + DH.*PhiMg.*pCaBpNa.*pKBpCa.*pNaBpK + DNa.*PhiMg.*pCaBpH.*pHBpK.*pKBpCa + DNa.*PhiMg.*pCaBpK.*pHBpCa.*pKBpH + DCa.*PhiMg.*pHBpK.*pKBpNa.*pNaBpH + DCa.*PhiMg.*pHBpNa.*pKBpH.*pNaBpK - DH.*PhiK.*pCaBpK.*pMgBpNa.*pNaBpCa + DH.*PhiK.*pCaBpNa.*pMgBpK.*pNaBpCa - DH.*PhiK.*pCaBpNa.*pMgBpCa.*pNaBpK - DNa.*PhiK.*pCaBpH.*pHBpK.*pMgBpCa + DNa.*PhiK.*pCaBpH.*pHBpCa.*pMgBpK - DNa.*PhiK.*pCaBpK.*pHBpCa.*pMgBpH - DCa.*PhiK.*pHBpK.*pMgBpNa.*pNaBpH - DCa.*PhiK.*pHBpNa.*pMgBpH.*pNaBpK + DCa.*PhiK.*pHBpNa.*pMgBpK.*pNaBpH - DK.*PhiH.*pCaBpH.*pMgBpNa.*pNaBpCa + DK.*PhiH.*pCaBpNa.*pMgBpH.*pNaBpCa - DK.*PhiH.*pCaBpNa.*pMgBpCa.*pNaBpH - DNa.*PhiH.*pCaBpH.*pKBpCa.*pMgBpK - DNa.*PhiH.*pCaBpK.*pKBpH.*pMgBpCa + DNa.*PhiH.*pCaBpK.*pKBpCa.*pMgBpH - DCa.*PhiH.*pKBpH.*pMgBpNa.*pNaBpK + DCa.*PhiH.*pKBpNa.*pMgBpH.*pNaBpK - DCa.*PhiH.*pKBpNa.*pMgBpK.*pNaBpH - DK.*PhiCa.*pHBpCa.*pMgBpNa.*pNaBpH - DK.*PhiCa.*pHBpNa.*pMgBpH.*pNaBpCa + DK.*PhiCa.*pHBpNa.*pMgBpCa.*pNaBpH - DH.*PhiCa.*pKBpCa.*pMgBpNa.*pNaBpK - DH.*PhiCa.*pKBpNa.*pMgBpK.*pNaBpCa + DH.*PhiCa.*pKBpNa.*pMgBpCa.*pNaBpK + DNa.*PhiCa.*pHBpK.*pKBpH.*pMgBpCa - DNa.*PhiCa.*pHBpK.*pKBpCa.*pMgBpH - DNa.*PhiCa.*pHBpCa.*pKBpH.*pMgBpK - PhiNa.*pCaBpH.*pHBpK.*pKBpCa.*pMgBpNa + PhiNa.*pCaBpH.*pHBpK.*pKBpNa.*pMgBpCa - PhiNa.*pCaBpH.*pHBpCa.*pKBpNa.*pMgBpK + PhiNa.*pCaBpH.*pHBpNa.*pKBpCa.*pMgBpK - PhiNa.*pCaBpK.*pHBpCa.*pKBpH.*pMgBpNa + PhiNa.*pCaBpK.*pHBpCa.*pKBpNa.*pMgBpH + PhiNa.*pCaBpK.*pHBpNa.*pKBpH.*pMgBpCa - PhiNa.*pCaBpK.*pHBpNa.*pKBpCa.*pMgBpH - PhiNa.*pCaBpNa.*pHBpK.*pKBpH.*pMgBpCa + PhiNa.*pCaBpNa.*pHBpK.*pKBpCa.*pMgBpH + PhiNa.*pCaBpNa.*pHBpCa.*pKBpH.*pMgBpK - PhiMg.*pCaBpH.*pHBpK.*pKBpNa.*pNaBpCa + PhiMg.*pCaBpH.*pHBpCa.*pKBpNa.*pNaBpK - PhiMg.*pCaBpH.*pHBpNa.*pKBpCa.*pNaBpK - PhiMg.*pCaBpK.*pHBpCa.*pKBpNa.*pNaBpH - PhiMg.*pCaBpK.*pHBpNa.*pKBpH.*pNaBpCa + PhiMg.*pCaBpK.*pHBpNa.*pKBpCa.*pNaBpH + PhiMg.*pCaBpNa.*pHBpK.*pKBpH.*pNaBpCa - PhiMg.*pCaBpNa.*pHBpK.*pKBpCa.*pNaBpH - PhiMg.*pCaBpNa.*pHBpCa.*pKBpH.*pNaBpK + PhiK.*pCaBpH.*pHBpK.*pMgBpNa.*pNaBpCa - PhiK.*pCaBpH.*pHBpCa.*pMgBpNa.*pNaBpK - PhiK.*pCaBpH.*pHBpNa.*pMgBpK.*pNaBpCa + PhiK.*pCaBpH.*pHBpNa.*pMgBpCa.*pNaBpK + PhiK.*pCaBpK.*pHBpCa.*pMgBpNa.*pNaBpH + PhiK.*pCaBpK.*pHBpNa.*pMgBpH.*pNaBpCa - PhiK.*pCaBpK.*pHBpNa.*pMgBpCa.*pNaBpH - PhiK.*pCaBpNa.*pHBpK.*pMgBpH.*pNaBpCa + PhiK.*pCaBpNa.*pHBpK.*pMgBpCa.*pNaBpH + PhiK.*pCaBpNa.*pHBpCa.*pMgBpH.*pNaBpK - PhiK.*pCaBpNa.*pHBpCa.*pMgBpK.*pNaBpH + PhiH.*pCaBpH.*pKBpCa.*pMgBpNa.*pNaBpK + PhiH.*pCaBpH.*pKBpNa.*pMgBpK.*pNaBpCa - PhiH.*pCaBpH.*pKBpNa.*pMgBpCa.*pNaBpK + PhiH.*pCaBpK.*pKBpH.*pMgBpNa.*pNaBpCa - PhiH.*pCaBpK.*pKBpCa.*pMgBpNa.*pNaBpH - PhiH.*pCaBpK.*pKBpNa.*pMgBpH.*pNaBpCa + PhiH.*pCaBpK.*pKBpNa.*pMgBpCa.*pNaBpH - PhiH.*pCaBpNa.*pKBpH.*pMgBpK.*pNaBpCa + PhiH.*pCaBpNa.*pKBpH.*pMgBpCa.*pNaBpK - PhiH.*pCaBpNa.*pKBpCa.*pMgBpH.*pNaBpK + PhiH.*pCaBpNa.*pKBpCa.*pMgBpK.*pNaBpH - PhiCa.*pHBpK.*pKBpH.*pMgBpNa.*pNaBpCa + PhiCa.*pHBpK.*pKBpCa.*pMgBpNa.*pNaBpH + PhiCa.*pHBpK.*pKBpNa.*pMgBpH.*pNaBpCa - PhiCa.*pHBpK.*pKBpNa.*pMgBpCa.*pNaBpH + PhiCa.*pHBpCa.*pKBpH.*pMgBpNa.*pNaBpK - PhiCa.*pHBpCa.*pKBpNa.*pMgBpH.*pNaBpK + PhiCa.*pHBpCa.*pKBpNa.*pMgBpK.*pNaBpH + PhiCa.*pHBpNa.*pKBpH.*pMgBpK.*pNaBpCa - PhiCa.*pHBpNa.*pKBpH.*pMgBpCa.*pNaBpK + PhiCa.*pHBpNa.*pKBpCa.*pMgBpH.*pNaBpK - PhiCa.*pHBpNa.*pKBpCa.*pMgBpK.*pNaBpH + DCa.*DH.*DK.*DNa.*PhiMg - DCa.*DH.*DK.*PhiNa.*pMgBpNa - DCa.*DH.*DNa.*PhiK.*pMgBpK - DCa.*DK.*DNa.*PhiH.*pMgBpH - DH.*DK.*DNa.*PhiCa.*pMgBpCa)./D;
dCadt=-(DH.*DK.*PhiNa.*pCaBpMg.*pMgBpNa + DK.*DMg.*PhiNa.*pCaBpH.*pHBpNa + DH.*DMg.*PhiNa.*pCaBpK.*pKBpNa + DH.*DK.*PhiMg.*pCaBpNa.*pNaBpMg + DK.*DNa.*PhiMg.*pCaBpH.*pHBpMg + DH.*DNa.*PhiMg.*pCaBpK.*pKBpMg + DH.*DMg.*PhiK.*pCaBpNa.*pNaBpK + DH.*DNa.*PhiK.*pCaBpMg.*pMgBpK + DMg.*DNa.*PhiK.*pCaBpH.*pHBpK + DK.*DMg.*PhiH.*pCaBpNa.*pNaBpH + DK.*DNa.*PhiH.*pCaBpMg.*pMgBpH + DMg.*DNa.*PhiH.*pCaBpK.*pKBpH - DH.*DK.*PhiCa.*pMgBpNa.*pNaBpMg - DK.*DMg.*PhiCa.*pHBpNa.*pNaBpH - DH.*DMg.*PhiCa.*pKBpNa.*pNaBpK - DK.*DNa.*PhiCa.*pHBpMg.*pMgBpH - DH.*DNa.*PhiCa.*pKBpMg.*pMgBpK - DMg.*DNa.*PhiCa.*pHBpK.*pKBpH - DK.*PhiNa.*pCaBpH.*pHBpMg.*pMgBpNa + DK.*PhiNa.*pCaBpNa.*pHBpMg.*pMgBpH - DK.*PhiNa.*pCaBpMg.*pHBpNa.*pMgBpH - DH.*PhiNa.*pCaBpK.*pKBpMg.*pMgBpNa + DH.*PhiNa.*pCaBpNa.*pKBpMg.*pMgBpK - DH.*PhiNa.*pCaBpMg.*pKBpNa.*pMgBpK - DMg.*PhiNa.*pCaBpH.*pHBpK.*pKBpNa - DMg.*PhiNa.*pCaBpK.*pHBpNa.*pKBpH + DMg.*PhiNa.*pCaBpNa.*pHBpK.*pKBpH - DK.*PhiMg.*pCaBpH.*pHBpNa.*pNaBpMg - DK.*PhiMg.*pCaBpNa.*pHBpMg.*pNaBpH + DK.*PhiMg.*pCaBpMg.*pHBpNa.*pNaBpH - DH.*PhiMg.*pCaBpK.*pKBpNa.*pNaBpMg - DH.*PhiMg.*pCaBpNa.*pKBpMg.*pNaBpK + DH.*PhiMg.*pCaBpMg.*pKBpNa.*pNaBpK - DNa.*PhiMg.*pCaBpH.*pHBpK.*pKBpMg - DNa.*PhiMg.*pCaBpK.*pHBpMg.*pKBpH + DNa.*PhiMg.*pCaBpMg.*pHBpK.*pKBpH + DH.*PhiK.*pCaBpK.*pMgBpNa.*pNaBpMg - DH.*PhiK.*pCaBpNa.*pMgBpK.*pNaBpMg - DH.*PhiK.*pCaBpMg.*pMgBpNa.*pNaBpK - DMg.*PhiK.*pCaBpH.*pHBpNa.*pNaBpK + DMg.*PhiK.*pCaBpK.*pHBpNa.*pNaBpH - DMg.*PhiK.*pCaBpNa.*pHBpK.*pNaBpH - DNa.*PhiK.*pCaBpH.*pHBpMg.*pMgBpK + DNa.*PhiK.*pCaBpK.*pHBpMg.*pMgBpH - DNa.*PhiK.*pCaBpMg.*pHBpK.*pMgBpH + DK.*PhiH.*pCaBpH.*pMgBpNa.*pNaBpMg - DK.*PhiH.*pCaBpNa.*pMgBpH.*pNaBpMg - DK.*PhiH.*pCaBpMg.*pMgBpNa.*pNaBpH + DMg.*PhiH.*pCaBpH.*pKBpNa.*pNaBpK - DMg.*PhiH.*pCaBpK.*pKBpNa.*pNaBpH - DMg.*PhiH.*pCaBpNa.*pKBpH.*pNaBpK + DNa.*PhiH.*pCaBpH.*pKBpMg.*pMgBpK - DNa.*PhiH.*pCaBpK.*pKBpMg.*pMgBpH - DNa.*PhiH.*pCaBpMg.*pKBpH.*pMgBpK + DK.*PhiCa.*pHBpNa.*pMgBpH.*pNaBpMg + DK.*PhiCa.*pHBpMg.*pMgBpNa.*pNaBpH + DH.*PhiCa.*pKBpNa.*pMgBpK.*pNaBpMg + DH.*PhiCa.*pKBpMg.*pMgBpNa.*pNaBpK + DMg.*PhiCa.*pHBpK.*pKBpNa.*pNaBpH + DMg.*PhiCa.*pHBpNa.*pKBpH.*pNaBpK + DNa.*PhiCa.*pHBpK.*pKBpMg.*pMgBpH + DNa.*PhiCa.*pHBpMg.*pKBpH.*pMgBpK + PhiNa.*pCaBpH.*pHBpK.*pKBpMg.*pMgBpNa - PhiNa.*pCaBpH.*pHBpNa.*pKBpMg.*pMgBpK + PhiNa.*pCaBpH.*pHBpMg.*pKBpNa.*pMgBpK + PhiNa.*pCaBpK.*pHBpNa.*pKBpMg.*pMgBpH + PhiNa.*pCaBpK.*pHBpMg.*pKBpH.*pMgBpNa - PhiNa.*pCaBpK.*pHBpMg.*pKBpNa.*pMgBpH - PhiNa.*pCaBpNa.*pHBpK.*pKBpMg.*pMgBpH - PhiNa.*pCaBpNa.*pHBpMg.*pKBpH.*pMgBpK - PhiNa.*pCaBpMg.*pHBpK.*pKBpH.*pMgBpNa + PhiNa.*pCaBpMg.*pHBpK.*pKBpNa.*pMgBpH + PhiNa.*pCaBpMg.*pHBpNa.*pKBpH.*pMgBpK + PhiMg.*pCaBpH.*pHBpK.*pKBpNa.*pNaBpMg + PhiMg.*pCaBpH.*pHBpNa.*pKBpMg.*pNaBpK - PhiMg.*pCaBpH.*pHBpMg.*pKBpNa.*pNaBpK + PhiMg.*pCaBpK.*pHBpNa.*pKBpH.*pNaBpMg - PhiMg.*pCaBpK.*pHBpNa.*pKBpMg.*pNaBpH + PhiMg.*pCaBpK.*pHBpMg.*pKBpNa.*pNaBpH - PhiMg.*pCaBpNa.*pHBpK.*pKBpH.*pNaBpMg + PhiMg.*pCaBpNa.*pHBpK.*pKBpMg.*pNaBpH + PhiMg.*pCaBpNa.*pHBpMg.*pKBpH.*pNaBpK - PhiMg.*pCaBpMg.*pHBpK.*pKBpNa.*pNaBpH - PhiMg.*pCaBpMg.*pHBpNa.*pKBpH.*pNaBpK - PhiK.*pCaBpH.*pHBpK.*pMgBpNa.*pNaBpMg + PhiK.*pCaBpH.*pHBpNa.*pMgBpK.*pNaBpMg + PhiK.*pCaBpH.*pHBpMg.*pMgBpNa.*pNaBpK - PhiK.*pCaBpK.*pHBpNa.*pMgBpH.*pNaBpMg - PhiK.*pCaBpK.*pHBpMg.*pMgBpNa.*pNaBpH + PhiK.*pCaBpNa.*pHBpK.*pMgBpH.*pNaBpMg - PhiK.*pCaBpNa.*pHBpMg.*pMgBpH.*pNaBpK + PhiK.*pCaBpNa.*pHBpMg.*pMgBpK.*pNaBpH + PhiK.*pCaBpMg.*pHBpK.*pMgBpNa.*pNaBpH + PhiK.*pCaBpMg.*pHBpNa.*pMgBpH.*pNaBpK - PhiK.*pCaBpMg.*pHBpNa.*pMgBpK.*pNaBpH - PhiH.*pCaBpH.*pKBpNa.*pMgBpK.*pNaBpMg - PhiH.*pCaBpH.*pKBpMg.*pMgBpNa.*pNaBpK - PhiH.*pCaBpK.*pKBpH.*pMgBpNa.*pNaBpMg + PhiH.*pCaBpK.*pKBpNa.*pMgBpH.*pNaBpMg + PhiH.*pCaBpK.*pKBpMg.*pMgBpNa.*pNaBpH + PhiH.*pCaBpNa.*pKBpH.*pMgBpK.*pNaBpMg + PhiH.*pCaBpNa.*pKBpMg.*pMgBpH.*pNaBpK - PhiH.*pCaBpNa.*pKBpMg.*pMgBpK.*pNaBpH + PhiH.*pCaBpMg.*pKBpH.*pMgBpNa.*pNaBpK - PhiH.*pCaBpMg.*pKBpNa.*pMgBpH.*pNaBpK + PhiH.*pCaBpMg.*pKBpNa.*pMgBpK.*pNaBpH + PhiCa.*pHBpK.*pKBpH.*pMgBpNa.*pNaBpMg - PhiCa.*pHBpK.*pKBpNa.*pMgBpH.*pNaBpMg - PhiCa.*pHBpK.*pKBpMg.*pMgBpNa.*pNaBpH - PhiCa.*pHBpNa.*pKBpH.*pMgBpK.*pNaBpMg - PhiCa.*pHBpNa.*pKBpMg.*pMgBpH.*pNaBpK + PhiCa.*pHBpNa.*pKBpMg.*pMgBpK.*pNaBpH - PhiCa.*pHBpMg.*pKBpH.*pMgBpNa.*pNaBpK + PhiCa.*pHBpMg.*pKBpNa.*pMgBpH.*pNaBpK - PhiCa.*pHBpMg.*pKBpNa.*pMgBpK.*pNaBpH + DH.*DK.*DMg.*DNa.*PhiCa - DH.*DK.*DMg.*PhiNa.*pCaBpNa - DH.*DK.*DNa.*PhiMg.*pCaBpMg - DH.*DMg.*DNa.*PhiK.*pCaBpK - DK.*DMg.*DNa.*PhiH.*pCaBpH)./D;

% rate units defined by Phi units
f(p.i.H_x,:) = dHdt;
f(p.i.Na_x,:) = dNadt;
f(p.i.K_x,:) = dKdt;
f(p.i.Mg_x,:) = dMgdt;
f(p.i.Ca_x,:) = dCadt;

end

%% Compute Extra-Matrix Ion Derivatives Using Rapid Equilibrium Approximation

function f = cCation_dXdt(x,f,p,J)
%% Compute Time Derivatives for Cations
H_c = x(p.i.H_c,:);     % M
K_c = x(p.i.K_c,:);     % M
Na_c = x(p.i.Na_c,:);   % M
Mg_c = x(p.i.Mg_c,:);   % M
Ca_c = x(p.i.Ca_c,:);   % M

%% Define Ligand Vector
Li = x(p.i.Lcyt,:); % M

%% Define Ligand Derivatives
dLidt = f(p.i.Lcyt,:); % M/min

%% Partial Derivatives of binding polynomials:
Pm_L = p.BP(p.i.Lcyt,:);
Pm_L2 = Pm_L.^2;
Pm_HBH = (1 + 1./p.KD(p.if.ligands,p.if.K)*K_c + 1./p.KD(p.if.ligands,p.if.Na)*Na_c ...
    + 1./p.KD(p.if.ligands,p.if.Mg)*Mg_c + 1./p.KD(p.if.ligands,p.if.Ca)*Ca_c);
Pm_KBK = (1 + 1./p.KD(p.if.ligands,p.if.H)*H_c + 1./p.KD(p.if.ligands,p.if.Na)*Na_c ...
    + 1./p.KD(p.if.ligands,p.if.Mg)*Mg_c + 1./p.KD(p.if.ligands,p.if.Ca)*Ca_c);
Pm_NBN = (1 + 1./p.KD(p.if.ligands,p.if.H)*H_c + 1./p.KD(p.if.ligands,p.if.K)*K_c ...
    + 1./p.KD(p.if.ligands,p.if.Mg)*Mg_c + 1./p.KD(p.if.ligands,p.if.Ca)*Ca_c);
Pm_MBM = (1 + 1./p.KD(p.if.ligands,p.if.H)*H_c + 1./p.KD(p.if.ligands,p.if.K)*K_c ...
    + 1./p.KD(p.if.ligands,p.if.Na)*Na_c + 1./p.KD(p.if.ligands,p.if.Ca)*Ca_c);
Pm_CBC = (1 + 1./p.KD(p.if.ligands,p.if.H)*H_c + 1./p.KD(p.if.ligands,p.if.K)*K_c ...
    + 1./p.KD(p.if.ligands,p.if.Na)*Na_c + 1./p.KD(p.if.ligands,p.if.Mg)*Mg_c);

pHBpH = + sum(Pm_HBH.*Li./Pm_L2./(p.KD(p.if.ligands,p.if.H)*ones(1,size(Li,2))),1);
pHBpK = - H_c.*sum(Li./Pm_L2 ...
    ./(p.KD(p.if.ligands,p.if.H).*p.KD(p.if.ligands,p.if.K)*ones(1,size(Li,2))),1);
pHBpNa = - H_c.*sum(Li./Pm_L2 ...
    ./(p.KD(p.if.ligands,p.if.H).*p.KD(p.if.ligands,p.if.Na)*ones(1,size(Li,2))),1);
pHBpMg = - H_c.*sum(Li./Pm_L2 ...
    ./(p.KD(p.if.ligands,p.if.H).*p.KD(p.if.ligands,p.if.Mg)*ones(1,size(Li,2))),1);
pHBpCa = - H_c.*sum(Li./Pm_L2 ...
    ./(p.KD(p.if.ligands,p.if.H).*p.KD(p.if.ligands,p.if.Ca)*ones(1,size(Li,2))),1);

pKBpH = - K_c.*sum(Li./Pm_L2 ...
    ./(p.KD(p.if.ligands,p.if.K).*p.KD(p.if.ligands,p.if.H)*ones(1,size(Li,2))),1);
pKBpK = + sum(Pm_KBK.*Li./Pm_L2./(p.KD(p.if.ligands,p.if.K)*ones(1,size(Li,2))),1);
pKBpNa = - K_c.*sum(Li./Pm_L2 ...
    ./(p.KD(p.if.ligands,p.if.K).*p.KD(p.if.ligands,p.if.Na)*ones(1,size(Li,2))),1);
pKBpMg = - K_c.*sum(Li./Pm_L2 ...
    ./(p.KD(p.if.ligands,p.if.K).*p.KD(p.if.ligands,p.if.Mg)*ones(1,size(Li,2))),1);
pKBpCa = - K_c.*sum(Li./Pm_L2 ...
    ./(p.KD(p.if.ligands,p.if.K).*p.KD(p.if.ligands,p.if.Ca)*ones(1,size(Li,2))),1);

pNaBpH = - Na_c.*sum(Li./Pm_L2 ...
    ./(p.KD(p.if.ligands,p.if.Na).*p.KD(p.if.ligands,p.if.H)*ones(1,size(Li,2))),1);
pNaBpK = - Na_c.*sum(Li./Pm_L2 ...
    ./(p.KD(p.if.ligands,p.if.Na).*p.KD(p.if.ligands,p.if.K)*ones(1,size(Li,2))),1);
pNaBpNa = + sum(Pm_NBN.*Li./Pm_L2./(p.KD(p.if.ligands,p.if.Na)*ones(1,size(Li,2))),1);
pNaBpMg = - Na_c.*sum(Li./Pm_L2 ...
    ./(p.KD(p.if.ligands,p.if.Na).*p.KD(p.if.ligands,p.if.Mg)*ones(1,size(Li,2))),1);
pNaBpCa = - Na_c.*sum(Li./Pm_L2 ...
    ./(p.KD(p.if.ligands,p.if.Na).*p.KD(p.if.ligands,p.if.Ca)*ones(1,size(Li,2))),1);

pMgBpH = - Mg_c.*sum(Li./Pm_L2 ...
    ./(p.KD(p.if.ligands,p.if.Mg).*p.KD(p.if.ligands,p.if.H)*ones(1,size(Li,2))),1);
pMgBpK = - Mg_c.*sum(Li./Pm_L2 ...
    ./(p.KD(p.if.ligands,p.if.Mg).*p.KD(p.if.ligands,p.if.K)*ones(1,size(Li,2))),1);
pMgBpNa = - Mg_c.*sum(Li./Pm_L2 ...
    ./(p.KD(p.if.ligands,p.if.Mg).*p.KD(p.if.ligands,p.if.Na)*ones(1,size(Li,2))),1);
pMgBpMg = + sum(Pm_MBM.*Li./Pm_L2./(p.KD(p.if.ligands,p.if.Mg)*ones(1,size(Li,2))),1);
pMgBpCa = - Mg_c.*sum(Li./Pm_L2 ...
    ./(p.KD(p.if.ligands,p.if.Mg).*p.KD(p.if.ligands,p.if.Ca)*ones(1,size(Li,2))),1);

pCaBpH = - Ca_c.*sum(Li./Pm_L2 ...
    ./(p.KD(p.if.ligands,p.if.Ca).*p.KD(p.if.ligands,p.if.H)*ones(1,size(Li,2))),1);
pCaBpK = - Ca_c.*sum(Li./Pm_L2 ...
    ./(p.KD(p.if.ligands,p.if.Ca).*p.KD(p.if.ligands,p.if.K)*ones(1,size(Li,2))),1);
pCaBpNa = - Ca_c.*sum(Li./Pm_L2 ...
    ./(p.KD(p.if.ligands,p.if.Ca).*p.KD(p.if.ligands,p.if.Na)*ones(1,size(Li,2))),1);
pCaBpMg = - Ca_c.*sum(Li./Pm_L2 ...
    ./(p.KD(p.if.ligands,p.if.Ca).*p.KD(p.if.ligands,p.if.Mg)*ones(1,size(Li,2))),1);
pCaBpCa = + sum(Pm_CBC.*Li./Pm_L2 ...
    ./(p.KD(p.if.ligands,p.if.Ca)*ones(1,size(Li,2))),1);

%% PHI's:
PhiH = - H_c.*sum(dLidt./Pm_L ...
    ./(p.KD(p.if.ligands,p.if.H)*ones(1,size(Li,2)))) ...
    - ( - 5*J.CI.JNADH - 1*J.CIII.Jc3 - 4*J.CIV ...
    + (p.F1FO.nH-1)*J.F1FO + 2*J.PiC + J.mHleak ...
    + 1*J.mKHE + J.mNHE ...
    - J.Inject(p.if.H,:))/p.Vol.H2O_cyt; % add CHE
PhiK = - K_c.*sum(dLidt./Pm_L ...
    ./(p.KD(p.if.ligands,p.if.K)*ones(1,size(Li,2)))) ...
    - ( - J.mKHE - J.Inject(p.if.K,:))/p.Vol.H2O_cyt;
PhiNa = - Na_c.*sum(dLidt./Pm_L ...
    ./(p.KD(p.if.ligands,p.if.Na)*ones(1,size(Li,2))))...
    - (3*J.NCLX - J.mNHE - J.Inject(p.if.Na,:))/p.Vol.H2O_cyt;
PhiMg = - Mg_c.*sum(dLidt./Pm_L ...
    ./(p.KD(p.if.ligands,p.if.Mg)*ones(1,size(Li,2)))) ...
    + J.Inject(p.if.Mg,:)/p.Vol.H2O_cyt;
PhiCa = - Ca_c.*sum(dLidt./Pm_L ...
    ./(p.KD(p.if.ligands,p.if.Ca)*ones(1,size(Li,2)))) ...
    - (J.MCU - J.NCLX - J.Inject(p.if.Ca,:))/p.Vol.H2O_cyt; % add CHE

%% additional buffers for [H+], [K+], [Na+], [Mg2+] and [Ca2+]
BH = p.cBuff.Htot;      % M
K_BH = p.cBuff.KD_H; % M

% extra buffers added to cytosol
EDTA = p.cBuff.EDTA;   % M, total
EGTA = p.cBuff.EGTA;   % M, total

% EDTA binding poly (must include 2nd protonation state)
P_EDTA = 1 + H_c./p.EB.KD(p.if2.EDTA,p.if2.H) ...
    + H_c.^2./p.EB.KD(p.if2.EDTA,p.if2.H)./p.EB.KD(p.if2.EDTA,p.if2.HH)...
    + Na_c./p.EB.KD(p.if2.EDTA,p.if2.Na) ...
    + K_c./p.EB.KD(p.if2.EDTA,p.if2.K) ...
    + Mg_c./p.EB.KD(p.if2.EDTA,p.if2.Mg) ...
    + Ca_c./p.EB.KD(p.if2.EDTA,p.if2.Ca);

% EGTA binding poly (must include 2nd protonation state)
P_EGTA = 1 + H_c./p.EB.KD(p.if2.EGTA,p.if2.H) ...
    + H_c.^2./p.EB.KD(p.if2.EGTA,p.if2.H)./p.EB.KD(p.if2.EGTA,p.if2.HH)...
    + Na_c./p.EB.KD(p.if2.EGTA,p.if2.Na) ...
    + K_c./p.EB.KD(p.if2.EGTA,p.if2.K) ...
    + Mg_c./p.EB.KD(p.if2.EGTA,p.if2.Mg) ...
    + Ca_c./p.EB.KD(p.if2.EGTA,p.if2.Ca);

% K and Na binding to OM proteins/lipids negligible
BK=0;K_BK=inf;BNa=0;K_BNa=inf;

% Currently, do not allow Mg/Ca to bind to OM proteins/lipids
BMg=0;K_BMg=inf;

% extra-matrix buffering from respiration buffer constituents
BCa = p.cBuff.Ca.Btot;
K_BCa = p.cBuff.Ca.K;
n_BCa = p.cBuff.Ca.n;

% Compute partial derivatives of EGTA and EDTA species
% here HEGTA is taken to be HEGTA+H2EGTA
% and HEDTA = HEDTA + H2EDTA
pHEGTApH = EGTA/p.EB.KD(p.if2.EGTA,p.if2.H)./P_EGTA.^2.*((P_EGTA ...
    - H_c/p.EB.KD(p.if2.EGTA,p.if2.H) ...
    - 2*H_c.^2/p.EB.KD(p.if2.EGTA,p.if2.H)/p.EB.KD(p.if2.EGTA,p.if2.HH))...
    + H_c/p.EB.KD(p.if2.EGTA,p.if2.HH).*(2*P_EGTA ...
    - H_c/p.EB.KD(p.if2.EGTA,p.if2.H) ...
    - H_c.^2/p.EB.KD(p.if2.EGTA,p.if2.H)/p.EB.KD(p.if2.EGTA,p.if2.HH)));
pHEGTApNa = -H_c.*EGTA/p.EB.KD(p.if2.EGTA,p.if2.H) ...
    /p.EB.KD(p.if2.EGTA,p.if2.Na)./P_EGTA.^2.*(1 ...
    + H_c/p.EB.KD(p.if2.EGTA,p.if2.HH));
pHEGTApK = -H_c.*EGTA/p.EB.KD(p.if2.EGTA,p.if2.H) ...
    /p.EB.KD(p.if2.EGTA,p.if2.K)./P_EGTA.^2.*(1 ...
    + H_c/p.EB.KD(p.if2.EGTA,p.if2.HH));
pHEGTApMg = -H_c.*EGTA/p.EB.KD(p.if2.EGTA,p.if2.H) ...
    /p.EB.KD(p.if2.EGTA,p.if2.Mg)./P_EGTA.^2.*(1 ...
    + H_c/p.EB.KD(p.if2.EGTA,p.if2.HH));
pHEGTApCa = -H_c.*EGTA/p.EB.KD(p.if2.EGTA,p.if2.H) ...
    /p.EB.KD(p.if2.EGTA,p.if2.Ca)./P_EGTA.^2.*(1 ...
    + H_c/p.EB.KD(p.if2.EGTA,p.if2.HH));

pNaEGTApH = -Na_c.*EGTA/p.EB.KD(p.if2.EGTA,p.if2.Na)./P_EGTA.^2 ...
    .*(1/p.EB.KD(p.if2.EGTA,p.if2.H) ...
    + 2*H_c/p.EB.KD(p.if2.EGTA,p.if2.H)/p.EB.KD(p.if2.EGTA,p.if2.HH));
pNaEGTApNa = EGTA/p.EB.KD(p.if2.EGTA,p.if2.Na)./P_EGTA.^2.*(P_EGTA ...
    - Na_c/p.EB.KD(p.if2.EGTA,p.if2.Na));
pNaEGTApK = -Na_c.*EGTA/p.EB.KD(p.if2.EGTA,p.if2.Na)./P_EGTA.^2 ...
    /p.EB.KD(p.if2.EGTA,p.if2.K);
pNaEGTApMg = -Na_c.*EGTA/p.EB.KD(p.if2.EGTA,p.if2.Na)./P_EGTA.^2 ...
    /p.EB.KD(p.if2.EGTA,p.if2.Mg);
pNaEGTApCa = -Na_c.*EGTA/p.EB.KD(p.if2.EGTA,p.if2.Na)./P_EGTA.^2 ...
    /p.EB.KD(p.if2.EGTA,p.if2.Ca);

pKEGTApH = -K_c.*EGTA/p.EB.KD(p.if2.EGTA,p.if2.K)./P_EGTA.^2 ...
    .*(1/p.EB.KD(p.if2.EGTA,p.if2.H) ...
    + 2*H_c/p.EB.KD(p.if2.EGTA,p.if2.H)/p.EB.KD(p.if2.EGTA,p.if2.HH));
pKEGTApNa = -K_c.*EGTA/p.EB.KD(p.if2.EGTA,p.if2.K)./P_EGTA.^2 ...
    /p.EB.KD(p.if2.EGTA,p.if2.Na);
pKEGTApK = EGTA/p.EB.KD(p.if2.EGTA,p.if2.K)./P_EGTA.^2.*(P_EGTA ...
    - K_c/p.EB.KD(p.if2.EGTA,p.if2.K));
pKEGTApMg = -K_c.*EGTA/p.EB.KD(p.if2.EGTA,p.if2.K)./P_EGTA.^2 ...
    /p.EB.KD(p.if2.EGTA,p.if2.Mg);
pKEGTApCa = -K_c.*EGTA/p.EB.KD(p.if2.EGTA,p.if2.K)./P_EGTA.^2 ...
    /p.EB.KD(p.if2.EGTA,p.if2.Ca);

pMgEGTApH = -Mg_c.*EGTA/p.EB.KD(p.if2.EGTA,p.if2.Mg)./P_EGTA.^2 ...
    .*(1/p.EB.KD(p.if2.EGTA,p.if2.H) ...
    + 2*H_c/p.EB.KD(p.if2.EGTA,p.if2.H)/p.EB.KD(p.if2.EGTA,p.if2.HH));
pMgEGTApNa = -Mg_c.*EGTA/p.EB.KD(p.if2.EGTA,p.if2.Mg)./P_EGTA.^2 ...
    /p.EB.KD(p.if2.EGTA,p.if2.Na);
pMgEGTApK = -Mg_c.*EGTA/p.EB.KD(p.if2.EGTA,p.if2.Mg)./P_EGTA.^2 ...
    /p.EB.KD(p.if2.EGTA,p.if2.K);
pMgEGTApMg = EGTA/p.EB.KD(p.if2.EGTA,p.if2.Mg)./P_EGTA.^2.*(P_EGTA ...
    - Mg_c/p.EB.KD(p.if2.EGTA,p.if2.Mg));
pMgEGTApCa = -Mg_c.*EGTA/p.EB.KD(p.if2.EGTA,p.if2.Mg)./P_EGTA.^2 ...
    /p.EB.KD(p.if2.EGTA,p.if2.Ca);

pCaEGTApH = -Ca_c.*EGTA/p.EB.KD(p.if2.EGTA,p.if2.Ca)./P_EGTA.^2 ...
    .*(1/p.EB.KD(p.if2.EGTA,p.if2.H) ...
    + 2*H_c/p.EB.KD(p.if2.EGTA,p.if2.H)/p.EB.KD(p.if2.EGTA,p.if2.HH));
pCaEGTApNa = -Ca_c.*EGTA/p.EB.KD(p.if2.EGTA,p.if2.Ca)./P_EGTA.^2 ...
    /p.EB.KD(p.if2.EGTA,p.if2.Na);
pCaEGTApK = -Ca_c.*EGTA/p.EB.KD(p.if2.EGTA,p.if2.Ca)./P_EGTA.^2 ...
    /p.EB.KD(p.if2.EGTA,p.if2.K);
pCaEGTApMg = -Ca_c.*EGTA/p.EB.KD(p.if2.EGTA,p.if2.Ca)./P_EGTA.^2 ...
    /p.EB.KD(p.if2.EGTA,p.if2.Mg);
pCaEGTApCa = EGTA/p.EB.KD(p.if2.EGTA,p.if2.Ca)./P_EGTA.^2.*(P_EGTA ...
    - Ca_c/p.EB.KD(p.if2.EGTA,p.if2.Ca));

pHEDTApH = EDTA/p.EB.KD(p.if2.EDTA,p.if2.H)./P_EDTA.^2.*((P_EDTA ...
    - H_c/p.EB.KD(p.if2.EDTA,p.if2.H) ...
    - 2*H_c.^2/p.EB.KD(p.if2.EDTA,p.if2.H)/p.EB.KD(p.if2.EDTA,p.if2.HH))...
    + H_c/p.EB.KD(p.if2.EDTA,p.if2.HH).*(2*P_EDTA ...
    - H_c/p.EB.KD(p.if2.EDTA,p.if2.H) ...
    - H_c.^2/p.EB.KD(p.if2.EDTA,p.if2.H)/p.EB.KD(p.if2.EDTA,p.if2.HH)));
pHEDTApNa = -H_c.*EDTA/p.EB.KD(p.if2.EDTA,p.if2.H) ...
    /p.EB.KD(p.if2.EDTA,p.if2.Na)./P_EDTA.^2.*(1 ...
    + H_c/p.EB.KD(p.if2.EDTA,p.if2.HH));
pHEDTApK = -H_c.*EDTA/p.EB.KD(p.if2.EDTA,p.if2.H) ...
    /p.EB.KD(p.if2.EDTA,p.if2.K)./P_EDTA.^2.*(1 ...
    + H_c/p.EB.KD(p.if2.EDTA,p.if2.HH));
pHEDTApMg = -H_c.*EDTA/p.EB.KD(p.if2.EDTA,p.if2.H) ...
    /p.EB.KD(p.if2.EDTA,p.if2.Mg)./P_EDTA.^2.*(1 ...
    + H_c/p.EB.KD(p.if2.EDTA,p.if2.HH));
pHEDTApCa = -H_c.*EDTA/p.EB.KD(p.if2.EDTA,p.if2.H) ...
    /p.EB.KD(p.if2.EDTA,p.if2.Ca)./P_EDTA.^2.*(1 ...
    + H_c/p.EB.KD(p.if2.EDTA,p.if2.HH));

pNaEDTApH = -Na_c.*EDTA/p.EB.KD(p.if2.EDTA,p.if2.Na)./P_EDTA.^2 ...
    .*(1/p.EB.KD(p.if2.EDTA,p.if2.H) ...
    + 2*H_c/p.EB.KD(p.if2.EDTA,p.if2.H)/p.EB.KD(p.if2.EDTA,p.if2.HH));
pNaEDTApNa = EDTA/p.EB.KD(p.if2.EDTA,p.if2.Na)./P_EDTA.^2.*(P_EDTA ...
    - Na_c/p.EB.KD(p.if2.EDTA,p.if2.Na));
pNaEDTApK = -Na_c.*EDTA/p.EB.KD(p.if2.EDTA,p.if2.Na)./P_EDTA.^2 ...
    /p.EB.KD(p.if2.EDTA,p.if2.K);
pNaEDTApMg = -Na_c.*EDTA/p.EB.KD(p.if2.EDTA,p.if2.Na)./P_EDTA.^2 ...
    /p.EB.KD(p.if2.EDTA,p.if2.Mg);
pNaEDTApCa = -Na_c.*EDTA/p.EB.KD(p.if2.EDTA,p.if2.Na)./P_EDTA.^2 ...
    /p.EB.KD(p.if2.EDTA,p.if2.Ca);

pKEDTApH = -K_c.*EDTA/p.EB.KD(p.if2.EDTA,p.if2.K)./P_EDTA.^2 ...
    .*(1/p.EB.KD(p.if2.EDTA,p.if2.H) ...
    + 2*H_c/p.EB.KD(p.if2.EDTA,p.if2.H)/p.EB.KD(p.if2.EDTA,p.if2.HH));
pKEDTApNa = -K_c.*EDTA/p.EB.KD(p.if2.EDTA,p.if2.K)./P_EDTA.^2 ...
    /p.EB.KD(p.if2.EDTA,p.if2.Na);
pKEDTApK = EDTA/p.EB.KD(p.if2.EDTA,p.if2.K)./P_EDTA.^2.*(P_EDTA ...
    - K_c/p.EB.KD(p.if2.EDTA,p.if2.K));
pKEDTApMg = -K_c.*EDTA/p.EB.KD(p.if2.EDTA,p.if2.K)./P_EDTA.^2 ...
    /p.EB.KD(p.if2.EDTA,p.if2.Mg);
pKEDTApCa = -K_c.*EDTA/p.EB.KD(p.if2.EDTA,p.if2.K)./P_EDTA.^2 ...
    /p.EB.KD(p.if2.EDTA,p.if2.Ca);

pMgEDTApH = -Mg_c.*EDTA/p.EB.KD(p.if2.EDTA,p.if2.Mg)./P_EDTA.^2 ...
    .*(1/p.EB.KD(p.if2.EDTA,p.if2.H) ...
    + 2*H_c/p.EB.KD(p.if2.EDTA,p.if2.H)/p.EB.KD(p.if2.EDTA,p.if2.HH));
pMgEDTApNa = -Mg_c.*EDTA/p.EB.KD(p.if2.EDTA,p.if2.Mg)./P_EDTA.^2 ...
    /p.EB.KD(p.if2.EDTA,p.if2.Na);
pMgEDTApK = -Mg_c.*EDTA/p.EB.KD(p.if2.EDTA,p.if2.Mg)./P_EDTA.^2 ...
    /p.EB.KD(p.if2.EDTA,p.if2.K);
pMgEDTApMg = EDTA/p.EB.KD(p.if2.EDTA,p.if2.Mg)./P_EDTA.^2.*(P_EDTA ...
    - Mg_c/p.EB.KD(p.if2.EDTA,p.if2.Mg));
pMgEDTApCa = -Mg_c.*EDTA/p.EB.KD(p.if2.EDTA,p.if2.Mg)./P_EDTA.^2 ...
    /p.EB.KD(p.if2.EDTA,p.if2.Ca);

pCaEDTApH = -Ca_c.*EDTA/p.EB.KD(p.if2.EDTA,p.if2.Ca)./P_EDTA.^2 ...
    .*(1/p.EB.KD(p.if2.EDTA,p.if2.H) ...
    + 2*H_c/p.EB.KD(p.if2.EDTA,p.if2.H)/p.EB.KD(p.if2.EDTA,p.if2.HH));
pCaEDTApNa = -Ca_c.*EDTA/p.EB.KD(p.if2.EDTA,p.if2.Ca)./P_EDTA.^2 ...
    /p.EB.KD(p.if2.EDTA,p.if2.Na);
pCaEDTApK = -Ca_c.*EDTA/p.EB.KD(p.if2.EDTA,p.if2.Ca)./P_EDTA.^2 ...
    /p.EB.KD(p.if2.EDTA,p.if2.K);
pCaEDTApMg = -Ca_c.*EDTA/p.EB.KD(p.if2.EDTA,p.if2.Ca)./P_EDTA.^2 ...
    /p.EB.KD(p.if2.EDTA,p.if2.Mg);
pCaEDTApCa = EDTA/p.EB.KD(p.if2.EDTA,p.if2.Ca)./P_EDTA.^2.*(P_EDTA ...
    - Ca_c/p.EB.KD(p.if2.EDTA,p.if2.Ca));

% add to partial derivatives already computed
pHBpH = pHBpH + pHEGTApH + pHEDTApH;
pHBpNa = pHBpNa + pHEGTApNa + pHEDTApNa;
pHBpK = pHBpK + pHEGTApK + pHEDTApK;
pHBpMg = pHBpMg + pHEGTApMg + pHEDTApMg;
pHBpCa = pHBpCa + pHEGTApCa + pHEDTApCa;

pKBpH = pKBpH + pKEGTApH + pKEDTApH;
pKBpK = pKBpK + pKEGTApK + pKEDTApK;
pKBpNa = pKBpNa + pKEGTApNa + pKEDTApNa;
pKBpMg = pKBpMg + pKEGTApMg + pKEDTApMg;
pKBpCa = pKBpCa + pKEGTApCa + pKEDTApCa;

pNaBpH = pNaBpH + pNaEGTApH + pNaEDTApH;
pNaBpK = pNaBpK  + pNaEGTApK + pNaEDTApK;
pNaBpNa = pNaBpNa + pNaEGTApNa + pNaEDTApNa;
pNaBpMg = pNaBpMg + pNaEGTApMg + pNaEDTApMg;
pNaBpCa = pNaBpCa + pNaEGTApCa + pNaEDTApCa;

pMgBpH = pMgBpH + pMgEGTApH + pMgEDTApH;
pMgBpK = pMgBpK + pMgEGTApK + pMgEDTApK;
pMgBpNa = pMgBpNa + pMgEGTApNa + pMgEDTApNa;
pMgBpMg = pMgBpMg + pMgEGTApMg + pMgEDTApMg;
pMgBpCa = pMgBpCa + pMgEGTApCa + pMgEDTApCa;

pCaBpH = pCaBpH + pCaEGTApH + pCaEDTApH;
pCaBpK = pCaBpK + pCaEGTApK + pCaEDTApK;
pCaBpNa = pCaBpNa + pCaEGTApNa + pCaEDTApNa;
pCaBpMg = pCaBpMg + pCaEGTApMg + pCaEDTApMg;
pCaBpCa = pCaBpCa + pCaEGTApCa + pCaEDTApCa;

%% Buffering Terms:
DH = 1 + pHBpH + BH./K_BH./(1+H_c./K_BH).^2;
DK = 1 + pKBpK + BK./K_BK./(1+K_c./K_BK).^2;
DNa = 1 + pNaBpNa + BNa./K_BNa./(1+Na_c./K_BNa).^2;
DMg = 1 + pMgBpMg + BMg./K_BMg./(1+Mg_c./K_BMg).^2;
DCa = 1 + pCaBpCa + n_BCa^2*Ca_c.^(n_BCa-1).*BCa/K_BCa^n_BCa./(1+(Ca_c/K_BCa).^n_BCa).^2;

%% analytical solution to cation eqns. based on Vinnakota et al., 2008
dHdt = -(DK.*DMg.*PhiNa.*pCaBpNa.*pHBpCa + DCa.*DK.*PhiNa.*pHBpMg.*pMgBpNa + DCa.*DMg.*PhiNa.*pHBpK.*pKBpNa + DK.*DNa.*PhiMg.*pCaBpMg.*pHBpCa + DCa.*DK.*PhiMg.*pHBpNa.*pNaBpMg + DCa.*DNa.*PhiMg.*pHBpK.*pKBpMg + DMg.*DNa.*PhiK.*pCaBpK.*pHBpCa + DCa.*DMg.*PhiK.*pHBpNa.*pNaBpK + DCa.*DNa.*PhiK.*pHBpMg.*pMgBpK - DK.*DMg.*PhiH.*pCaBpNa.*pNaBpCa - DK.*DNa.*PhiH.*pCaBpMg.*pMgBpCa - DCa.*DK.*PhiH.*pMgBpNa.*pNaBpMg - DMg.*DNa.*PhiH.*pCaBpK.*pKBpCa - DCa.*DMg.*PhiH.*pKBpNa.*pNaBpK - DCa.*DNa.*PhiH.*pKBpMg.*pMgBpK + DK.*DMg.*PhiCa.*pHBpNa.*pNaBpCa + DK.*DNa.*PhiCa.*pHBpMg.*pMgBpCa + DMg.*DNa.*PhiCa.*pHBpK.*pKBpCa - DK.*PhiNa.*pCaBpNa.*pHBpMg.*pMgBpCa - DK.*PhiNa.*pCaBpMg.*pHBpCa.*pMgBpNa + DK.*PhiNa.*pCaBpMg.*pHBpNa.*pMgBpCa - DMg.*PhiNa.*pCaBpK.*pHBpCa.*pKBpNa + DMg.*PhiNa.*pCaBpK.*pHBpNa.*pKBpCa - DMg.*PhiNa.*pCaBpNa.*pHBpK.*pKBpCa - DCa.*PhiNa.*pHBpK.*pKBpMg.*pMgBpNa + DCa.*PhiNa.*pHBpNa.*pKBpMg.*pMgBpK - DCa.*PhiNa.*pHBpMg.*pKBpNa.*pMgBpK - DK.*PhiMg.*pCaBpNa.*pHBpCa.*pNaBpMg + DK.*PhiMg.*pCaBpNa.*pHBpMg.*pNaBpCa - DK.*PhiMg.*pCaBpMg.*pHBpNa.*pNaBpCa - DNa.*PhiMg.*pCaBpK.*pHBpCa.*pKBpMg + DNa.*PhiMg.*pCaBpK.*pHBpMg.*pKBpCa - DNa.*PhiMg.*pCaBpMg.*pHBpK.*pKBpCa - DCa.*PhiMg.*pHBpK.*pKBpNa.*pNaBpMg - DCa.*PhiMg.*pHBpNa.*pKBpMg.*pNaBpK + DCa.*PhiMg.*pHBpMg.*pKBpNa.*pNaBpK - DMg.*PhiK.*pCaBpK.*pHBpNa.*pNaBpCa + DMg.*PhiK.*pCaBpNa.*pHBpK.*pNaBpCa - DMg.*PhiK.*pCaBpNa.*pHBpCa.*pNaBpK - DNa.*PhiK.*pCaBpK.*pHBpMg.*pMgBpCa + DNa.*PhiK.*pCaBpMg.*pHBpK.*pMgBpCa - DNa.*PhiK.*pCaBpMg.*pHBpCa.*pMgBpK + DCa.*PhiK.*pHBpK.*pMgBpNa.*pNaBpMg - DCa.*PhiK.*pHBpNa.*pMgBpK.*pNaBpMg - DCa.*PhiK.*pHBpMg.*pMgBpNa.*pNaBpK + DK.*PhiH.*pCaBpNa.*pMgBpCa.*pNaBpMg + DK.*PhiH.*pCaBpMg.*pMgBpNa.*pNaBpCa + DMg.*PhiH.*pCaBpK.*pKBpNa.*pNaBpCa + DMg.*PhiH.*pCaBpNa.*pKBpCa.*pNaBpK + DNa.*PhiH.*pCaBpK.*pKBpMg.*pMgBpCa + DNa.*PhiH.*pCaBpMg.*pKBpCa.*pMgBpK + DCa.*PhiH.*pKBpNa.*pMgBpK.*pNaBpMg + DCa.*PhiH.*pKBpMg.*pMgBpNa.*pNaBpK + DK.*PhiCa.*pHBpCa.*pMgBpNa.*pNaBpMg - DK.*PhiCa.*pHBpNa.*pMgBpCa.*pNaBpMg - DK.*PhiCa.*pHBpMg.*pMgBpNa.*pNaBpCa - DMg.*PhiCa.*pHBpK.*pKBpNa.*pNaBpCa + DMg.*PhiCa.*pHBpCa.*pKBpNa.*pNaBpK - DMg.*PhiCa.*pHBpNa.*pKBpCa.*pNaBpK - DNa.*PhiCa.*pHBpK.*pKBpMg.*pMgBpCa + DNa.*PhiCa.*pHBpCa.*pKBpMg.*pMgBpK - DNa.*PhiCa.*pHBpMg.*pKBpCa.*pMgBpK + PhiNa.*pCaBpK.*pHBpCa.*pKBpMg.*pMgBpNa - PhiNa.*pCaBpK.*pHBpNa.*pKBpMg.*pMgBpCa - PhiNa.*pCaBpK.*pHBpMg.*pKBpCa.*pMgBpNa + PhiNa.*pCaBpK.*pHBpMg.*pKBpNa.*pMgBpCa + PhiNa.*pCaBpNa.*pHBpK.*pKBpMg.*pMgBpCa - PhiNa.*pCaBpNa.*pHBpCa.*pKBpMg.*pMgBpK + PhiNa.*pCaBpNa.*pHBpMg.*pKBpCa.*pMgBpK + PhiNa.*pCaBpMg.*pHBpK.*pKBpCa.*pMgBpNa - PhiNa.*pCaBpMg.*pHBpK.*pKBpNa.*pMgBpCa + PhiNa.*pCaBpMg.*pHBpCa.*pKBpNa.*pMgBpK - PhiNa.*pCaBpMg.*pHBpNa.*pKBpCa.*pMgBpK + PhiMg.*pCaBpK.*pHBpCa.*pKBpNa.*pNaBpMg - PhiMg.*pCaBpK.*pHBpNa.*pKBpCa.*pNaBpMg + PhiMg.*pCaBpK.*pHBpNa.*pKBpMg.*pNaBpCa - PhiMg.*pCaBpK.*pHBpMg.*pKBpNa.*pNaBpCa + PhiMg.*pCaBpNa.*pHBpK.*pKBpCa.*pNaBpMg - PhiMg.*pCaBpNa.*pHBpK.*pKBpMg.*pNaBpCa + PhiMg.*pCaBpNa.*pHBpCa.*pKBpMg.*pNaBpK - PhiMg.*pCaBpNa.*pHBpMg.*pKBpCa.*pNaBpK + PhiMg.*pCaBpMg.*pHBpK.*pKBpNa.*pNaBpCa - PhiMg.*pCaBpMg.*pHBpCa.*pKBpNa.*pNaBpK + PhiMg.*pCaBpMg.*pHBpNa.*pKBpCa.*pNaBpK - PhiK.*pCaBpK.*pHBpCa.*pMgBpNa.*pNaBpMg + PhiK.*pCaBpK.*pHBpNa.*pMgBpCa.*pNaBpMg + PhiK.*pCaBpK.*pHBpMg.*pMgBpNa.*pNaBpCa - PhiK.*pCaBpNa.*pHBpK.*pMgBpCa.*pNaBpMg + PhiK.*pCaBpNa.*pHBpCa.*pMgBpK.*pNaBpMg - PhiK.*pCaBpNa.*pHBpMg.*pMgBpK.*pNaBpCa + PhiK.*pCaBpNa.*pHBpMg.*pMgBpCa.*pNaBpK - PhiK.*pCaBpMg.*pHBpK.*pMgBpNa.*pNaBpCa + PhiK.*pCaBpMg.*pHBpCa.*pMgBpNa.*pNaBpK + PhiK.*pCaBpMg.*pHBpNa.*pMgBpK.*pNaBpCa - PhiK.*pCaBpMg.*pHBpNa.*pMgBpCa.*pNaBpK + PhiH.*pCaBpK.*pKBpCa.*pMgBpNa.*pNaBpMg - PhiH.*pCaBpK.*pKBpNa.*pMgBpCa.*pNaBpMg - PhiH.*pCaBpK.*pKBpMg.*pMgBpNa.*pNaBpCa - PhiH.*pCaBpNa.*pKBpCa.*pMgBpK.*pNaBpMg + PhiH.*pCaBpNa.*pKBpMg.*pMgBpK.*pNaBpCa - PhiH.*pCaBpNa.*pKBpMg.*pMgBpCa.*pNaBpK - PhiH.*pCaBpMg.*pKBpCa.*pMgBpNa.*pNaBpK - PhiH.*pCaBpMg.*pKBpNa.*pMgBpK.*pNaBpCa + PhiH.*pCaBpMg.*pKBpNa.*pMgBpCa.*pNaBpK - PhiCa.*pHBpK.*pKBpCa.*pMgBpNa.*pNaBpMg + PhiCa.*pHBpK.*pKBpNa.*pMgBpCa.*pNaBpMg + PhiCa.*pHBpK.*pKBpMg.*pMgBpNa.*pNaBpCa - PhiCa.*pHBpCa.*pKBpNa.*pMgBpK.*pNaBpMg - PhiCa.*pHBpCa.*pKBpMg.*pMgBpNa.*pNaBpK + PhiCa.*pHBpNa.*pKBpCa.*pMgBpK.*pNaBpMg - PhiCa.*pHBpNa.*pKBpMg.*pMgBpK.*pNaBpCa + PhiCa.*pHBpNa.*pKBpMg.*pMgBpCa.*pNaBpK + PhiCa.*pHBpMg.*pKBpCa.*pMgBpNa.*pNaBpK + PhiCa.*pHBpMg.*pKBpNa.*pMgBpK.*pNaBpCa - PhiCa.*pHBpMg.*pKBpNa.*pMgBpCa.*pNaBpK + DCa.*DK.*DMg.*DNa.*PhiH - DCa.*DK.*DMg.*PhiNa.*pHBpNa - DCa.*DK.*DNa.*PhiMg.*pHBpMg - DCa.*DMg.*DNa.*PhiK.*pHBpK - DK.*DMg.*DNa.*PhiCa.*pHBpCa)./(DH.*DK.*DMg.*pCaBpNa.*pNaBpCa + DH.*DK.*DNa.*pCaBpMg.*pMgBpCa + DCa.*DH.*DK.*pMgBpNa.*pNaBpMg + DK.*DMg.*DNa.*pCaBpH.*pHBpCa + DH.*DMg.*DNa.*pCaBpK.*pKBpCa + DCa.*DK.*DMg.*pHBpNa.*pNaBpH + DCa.*DH.*DMg.*pKBpNa.*pNaBpK + DCa.*DK.*DNa.*pHBpMg.*pMgBpH + DCa.*DH.*DNa.*pKBpMg.*pMgBpK + DCa.*DMg.*DNa.*pHBpK.*pKBpH - DH.*DK.*pCaBpNa.*pMgBpCa.*pNaBpMg - DH.*DK.*pCaBpMg.*pMgBpNa.*pNaBpCa - DK.*DMg.*pCaBpH.*pHBpNa.*pNaBpCa - DK.*DMg.*pCaBpNa.*pHBpCa.*pNaBpH - DH.*DMg.*pCaBpK.*pKBpNa.*pNaBpCa - DH.*DMg.*pCaBpNa.*pKBpCa.*pNaBpK - DK.*DNa.*pCaBpH.*pHBpMg.*pMgBpCa - DK.*DNa.*pCaBpMg.*pHBpCa.*pMgBpH - DH.*DNa.*pCaBpK.*pKBpMg.*pMgBpCa - DH.*DNa.*pCaBpMg.*pKBpCa.*pMgBpK - DCa.*DK.*pHBpNa.*pMgBpH.*pNaBpMg - DCa.*DK.*pHBpMg.*pMgBpNa.*pNaBpH - DCa.*DH.*pKBpNa.*pMgBpK.*pNaBpMg - DCa.*DH.*pKBpMg.*pMgBpNa.*pNaBpK - DMg.*DNa.*pCaBpH.*pHBpK.*pKBpCa - DMg.*DNa.*pCaBpK.*pHBpCa.*pKBpH - DCa.*DMg.*pHBpK.*pKBpNa.*pNaBpH - DCa.*DMg.*pHBpNa.*pKBpH.*pNaBpK - DCa.*DNa.*pHBpK.*pKBpMg.*pMgBpH - DCa.*DNa.*pHBpMg.*pKBpH.*pMgBpK - DK.*pCaBpH.*pHBpCa.*pMgBpNa.*pNaBpMg + DK.*pCaBpH.*pHBpNa.*pMgBpCa.*pNaBpMg + DK.*pCaBpH.*pHBpMg.*pMgBpNa.*pNaBpCa + DK.*pCaBpNa.*pHBpCa.*pMgBpH.*pNaBpMg - DK.*pCaBpNa.*pHBpMg.*pMgBpH.*pNaBpCa + DK.*pCaBpNa.*pHBpMg.*pMgBpCa.*pNaBpH + DK.*pCaBpMg.*pHBpCa.*pMgBpNa.*pNaBpH + DK.*pCaBpMg.*pHBpNa.*pMgBpH.*pNaBpCa - DK.*pCaBpMg.*pHBpNa.*pMgBpCa.*pNaBpH - DH.*pCaBpK.*pKBpCa.*pMgBpNa.*pNaBpMg + DH.*pCaBpK.*pKBpNa.*pMgBpCa.*pNaBpMg + DH.*pCaBpK.*pKBpMg.*pMgBpNa.*pNaBpCa + DH.*pCaBpNa.*pKBpCa.*pMgBpK.*pNaBpMg - DH.*pCaBpNa.*pKBpMg.*pMgBpK.*pNaBpCa + DH.*pCaBpNa.*pKBpMg.*pMgBpCa.*pNaBpK + DH.*pCaBpMg.*pKBpCa.*pMgBpNa.*pNaBpK + DH.*pCaBpMg.*pKBpNa.*pMgBpK.*pNaBpCa - DH.*pCaBpMg.*pKBpNa.*pMgBpCa.*pNaBpK + DMg.*pCaBpH.*pHBpK.*pKBpNa.*pNaBpCa - DMg.*pCaBpH.*pHBpCa.*pKBpNa.*pNaBpK + DMg.*pCaBpH.*pHBpNa.*pKBpCa.*pNaBpK + DMg.*pCaBpK.*pHBpCa.*pKBpNa.*pNaBpH + DMg.*pCaBpK.*pHBpNa.*pKBpH.*pNaBpCa - DMg.*pCaBpK.*pHBpNa.*pKBpCa.*pNaBpH - DMg.*pCaBpNa.*pHBpK.*pKBpH.*pNaBpCa + DMg.*pCaBpNa.*pHBpK.*pKBpCa.*pNaBpH + DMg.*pCaBpNa.*pHBpCa.*pKBpH.*pNaBpK + DNa.*pCaBpH.*pHBpK.*pKBpMg.*pMgBpCa - DNa.*pCaBpH.*pHBpCa.*pKBpMg.*pMgBpK + DNa.*pCaBpH.*pHBpMg.*pKBpCa.*pMgBpK + DNa.*pCaBpK.*pHBpCa.*pKBpMg.*pMgBpH + DNa.*pCaBpK.*pHBpMg.*pKBpH.*pMgBpCa - DNa.*pCaBpK.*pHBpMg.*pKBpCa.*pMgBpH - DNa.*pCaBpMg.*pHBpK.*pKBpH.*pMgBpCa + DNa.*pCaBpMg.*pHBpK.*pKBpCa.*pMgBpH + DNa.*pCaBpMg.*pHBpCa.*pKBpH.*pMgBpK - DCa.*pHBpK.*pKBpH.*pMgBpNa.*pNaBpMg + DCa.*pHBpK.*pKBpNa.*pMgBpH.*pNaBpMg + DCa.*pHBpK.*pKBpMg.*pMgBpNa.*pNaBpH + DCa.*pHBpNa.*pKBpH.*pMgBpK.*pNaBpMg + DCa.*pHBpNa.*pKBpMg.*pMgBpH.*pNaBpK - DCa.*pHBpNa.*pKBpMg.*pMgBpK.*pNaBpH + DCa.*pHBpMg.*pKBpH.*pMgBpNa.*pNaBpK - DCa.*pHBpMg.*pKBpNa.*pMgBpH.*pNaBpK + DCa.*pHBpMg.*pKBpNa.*pMgBpK.*pNaBpH + pCaBpH.*pHBpK.*pKBpCa.*pMgBpNa.*pNaBpMg - pCaBpH.*pHBpK.*pKBpNa.*pMgBpCa.*pNaBpMg - pCaBpH.*pHBpK.*pKBpMg.*pMgBpNa.*pNaBpCa + pCaBpH.*pHBpCa.*pKBpNa.*pMgBpK.*pNaBpMg + pCaBpH.*pHBpCa.*pKBpMg.*pMgBpNa.*pNaBpK - pCaBpH.*pHBpNa.*pKBpCa.*pMgBpK.*pNaBpMg + pCaBpH.*pHBpNa.*pKBpMg.*pMgBpK.*pNaBpCa - pCaBpH.*pHBpNa.*pKBpMg.*pMgBpCa.*pNaBpK - pCaBpH.*pHBpMg.*pKBpCa.*pMgBpNa.*pNaBpK - pCaBpH.*pHBpMg.*pKBpNa.*pMgBpK.*pNaBpCa + pCaBpH.*pHBpMg.*pKBpNa.*pMgBpCa.*pNaBpK + pCaBpK.*pHBpCa.*pKBpH.*pMgBpNa.*pNaBpMg - pCaBpK.*pHBpCa.*pKBpNa.*pMgBpH.*pNaBpMg - pCaBpK.*pHBpCa.*pKBpMg.*pMgBpNa.*pNaBpH - pCaBpK.*pHBpNa.*pKBpH.*pMgBpCa.*pNaBpMg + pCaBpK.*pHBpNa.*pKBpCa.*pMgBpH.*pNaBpMg - pCaBpK.*pHBpNa.*pKBpMg.*pMgBpH.*pNaBpCa + pCaBpK.*pHBpNa.*pKBpMg.*pMgBpCa.*pNaBpH - pCaBpK.*pHBpMg.*pKBpH.*pMgBpNa.*pNaBpCa + pCaBpK.*pHBpMg.*pKBpCa.*pMgBpNa.*pNaBpH + pCaBpK.*pHBpMg.*pKBpNa.*pMgBpH.*pNaBpCa - pCaBpK.*pHBpMg.*pKBpNa.*pMgBpCa.*pNaBpH + pCaBpNa.*pHBpK.*pKBpH.*pMgBpCa.*pNaBpMg - pCaBpNa.*pHBpK.*pKBpCa.*pMgBpH.*pNaBpMg + pCaBpNa.*pHBpK.*pKBpMg.*pMgBpH.*pNaBpCa - pCaBpNa.*pHBpK.*pKBpMg.*pMgBpCa.*pNaBpH - pCaBpNa.*pHBpCa.*pKBpH.*pMgBpK.*pNaBpMg - pCaBpNa.*pHBpCa.*pKBpMg.*pMgBpH.*pNaBpK + pCaBpNa.*pHBpCa.*pKBpMg.*pMgBpK.*pNaBpH + pCaBpNa.*pHBpMg.*pKBpH.*pMgBpK.*pNaBpCa - pCaBpNa.*pHBpMg.*pKBpH.*pMgBpCa.*pNaBpK + pCaBpNa.*pHBpMg.*pKBpCa.*pMgBpH.*pNaBpK - pCaBpNa.*pHBpMg.*pKBpCa.*pMgBpK.*pNaBpH + pCaBpMg.*pHBpK.*pKBpH.*pMgBpNa.*pNaBpCa - pCaBpMg.*pHBpK.*pKBpCa.*pMgBpNa.*pNaBpH - pCaBpMg.*pHBpK.*pKBpNa.*pMgBpH.*pNaBpCa + pCaBpMg.*pHBpK.*pKBpNa.*pMgBpCa.*pNaBpH - pCaBpMg.*pHBpCa.*pKBpH.*pMgBpNa.*pNaBpK + pCaBpMg.*pHBpCa.*pKBpNa.*pMgBpH.*pNaBpK - pCaBpMg.*pHBpCa.*pKBpNa.*pMgBpK.*pNaBpH - pCaBpMg.*pHBpNa.*pKBpH.*pMgBpK.*pNaBpCa + pCaBpMg.*pHBpNa.*pKBpH.*pMgBpCa.*pNaBpK - pCaBpMg.*pHBpNa.*pKBpCa.*pMgBpH.*pNaBpK + pCaBpMg.*pHBpNa.*pKBpCa.*pMgBpK.*pNaBpH - DCa.*DH.*DK.*DMg.*DNa);
dKdt = -(DH.*DMg.*PhiNa.*pCaBpNa.*pKBpCa + DCa.*DH.*PhiNa.*pKBpMg.*pMgBpNa + DCa.*DMg.*PhiNa.*pHBpNa.*pKBpH + DH.*DNa.*PhiMg.*pCaBpMg.*pKBpCa + DCa.*DH.*PhiMg.*pKBpNa.*pNaBpMg + DCa.*DNa.*PhiMg.*pHBpMg.*pKBpH - DH.*DMg.*PhiK.*pCaBpNa.*pNaBpCa - DH.*DNa.*PhiK.*pCaBpMg.*pMgBpCa - DCa.*DH.*PhiK.*pMgBpNa.*pNaBpMg - DMg.*DNa.*PhiK.*pCaBpH.*pHBpCa - DCa.*DMg.*PhiK.*pHBpNa.*pNaBpH - DCa.*DNa.*PhiK.*pHBpMg.*pMgBpH + DMg.*DNa.*PhiH.*pCaBpH.*pKBpCa + DCa.*DMg.*PhiH.*pKBpNa.*pNaBpH + DCa.*DNa.*PhiH.*pKBpMg.*pMgBpH + DH.*DMg.*PhiCa.*pKBpNa.*pNaBpCa + DH.*DNa.*PhiCa.*pKBpMg.*pMgBpCa + DMg.*DNa.*PhiCa.*pHBpCa.*pKBpH - DH.*PhiNa.*pCaBpNa.*pKBpMg.*pMgBpCa - DH.*PhiNa.*pCaBpMg.*pKBpCa.*pMgBpNa + DH.*PhiNa.*pCaBpMg.*pKBpNa.*pMgBpCa + DMg.*PhiNa.*pCaBpH.*pHBpCa.*pKBpNa - DMg.*PhiNa.*pCaBpH.*pHBpNa.*pKBpCa - DMg.*PhiNa.*pCaBpNa.*pHBpCa.*pKBpH - DCa.*PhiNa.*pHBpNa.*pKBpMg.*pMgBpH - DCa.*PhiNa.*pHBpMg.*pKBpH.*pMgBpNa + DCa.*PhiNa.*pHBpMg.*pKBpNa.*pMgBpH - DH.*PhiMg.*pCaBpNa.*pKBpCa.*pNaBpMg + DH.*PhiMg.*pCaBpNa.*pKBpMg.*pNaBpCa - DH.*PhiMg.*pCaBpMg.*pKBpNa.*pNaBpCa + DNa.*PhiMg.*pCaBpH.*pHBpCa.*pKBpMg - DNa.*PhiMg.*pCaBpH.*pHBpMg.*pKBpCa - DNa.*PhiMg.*pCaBpMg.*pHBpCa.*pKBpH - DCa.*PhiMg.*pHBpNa.*pKBpH.*pNaBpMg + DCa.*PhiMg.*pHBpNa.*pKBpMg.*pNaBpH - DCa.*PhiMg.*pHBpMg.*pKBpNa.*pNaBpH + DH.*PhiK.*pCaBpNa.*pMgBpCa.*pNaBpMg + DH.*PhiK.*pCaBpMg.*pMgBpNa.*pNaBpCa + DMg.*PhiK.*pCaBpH.*pHBpNa.*pNaBpCa + DMg.*PhiK.*pCaBpNa.*pHBpCa.*pNaBpH + DNa.*PhiK.*pCaBpH.*pHBpMg.*pMgBpCa + DNa.*PhiK.*pCaBpMg.*pHBpCa.*pMgBpH + DCa.*PhiK.*pHBpNa.*pMgBpH.*pNaBpMg + DCa.*PhiK.*pHBpMg.*pMgBpNa.*pNaBpH - DMg.*PhiH.*pCaBpH.*pKBpNa.*pNaBpCa + DMg.*PhiH.*pCaBpNa.*pKBpH.*pNaBpCa - DMg.*PhiH.*pCaBpNa.*pKBpCa.*pNaBpH - DNa.*PhiH.*pCaBpH.*pKBpMg.*pMgBpCa + DNa.*PhiH.*pCaBpMg.*pKBpH.*pMgBpCa - DNa.*PhiH.*pCaBpMg.*pKBpCa.*pMgBpH + DCa.*PhiH.*pKBpH.*pMgBpNa.*pNaBpMg - DCa.*PhiH.*pKBpNa.*pMgBpH.*pNaBpMg - DCa.*PhiH.*pKBpMg.*pMgBpNa.*pNaBpH + DH.*PhiCa.*pKBpCa.*pMgBpNa.*pNaBpMg - DH.*PhiCa.*pKBpNa.*pMgBpCa.*pNaBpMg - DH.*PhiCa.*pKBpMg.*pMgBpNa.*pNaBpCa - DMg.*PhiCa.*pHBpCa.*pKBpNa.*pNaBpH - DMg.*PhiCa.*pHBpNa.*pKBpH.*pNaBpCa + DMg.*PhiCa.*pHBpNa.*pKBpCa.*pNaBpH - DNa.*PhiCa.*pHBpCa.*pKBpMg.*pMgBpH - DNa.*PhiCa.*pHBpMg.*pKBpH.*pMgBpCa + DNa.*PhiCa.*pHBpMg.*pKBpCa.*pMgBpH - PhiNa.*pCaBpH.*pHBpCa.*pKBpMg.*pMgBpNa + PhiNa.*pCaBpH.*pHBpNa.*pKBpMg.*pMgBpCa + PhiNa.*pCaBpH.*pHBpMg.*pKBpCa.*pMgBpNa - PhiNa.*pCaBpH.*pHBpMg.*pKBpNa.*pMgBpCa + PhiNa.*pCaBpNa.*pHBpCa.*pKBpMg.*pMgBpH + PhiNa.*pCaBpNa.*pHBpMg.*pKBpH.*pMgBpCa - PhiNa.*pCaBpNa.*pHBpMg.*pKBpCa.*pMgBpH + PhiNa.*pCaBpMg.*pHBpCa.*pKBpH.*pMgBpNa - PhiNa.*pCaBpMg.*pHBpCa.*pKBpNa.*pMgBpH - PhiNa.*pCaBpMg.*pHBpNa.*pKBpH.*pMgBpCa + PhiNa.*pCaBpMg.*pHBpNa.*pKBpCa.*pMgBpH - PhiMg.*pCaBpH.*pHBpCa.*pKBpNa.*pNaBpMg + PhiMg.*pCaBpH.*pHBpNa.*pKBpCa.*pNaBpMg - PhiMg.*pCaBpH.*pHBpNa.*pKBpMg.*pNaBpCa + PhiMg.*pCaBpH.*pHBpMg.*pKBpNa.*pNaBpCa + PhiMg.*pCaBpNa.*pHBpCa.*pKBpH.*pNaBpMg - PhiMg.*pCaBpNa.*pHBpCa.*pKBpMg.*pNaBpH - PhiMg.*pCaBpNa.*pHBpMg.*pKBpH.*pNaBpCa + PhiMg.*pCaBpNa.*pHBpMg.*pKBpCa.*pNaBpH + PhiMg.*pCaBpMg.*pHBpCa.*pKBpNa.*pNaBpH + PhiMg.*pCaBpMg.*pHBpNa.*pKBpH.*pNaBpCa - PhiMg.*pCaBpMg.*pHBpNa.*pKBpCa.*pNaBpH + PhiK.*pCaBpH.*pHBpCa.*pMgBpNa.*pNaBpMg - PhiK.*pCaBpH.*pHBpNa.*pMgBpCa.*pNaBpMg - PhiK.*pCaBpH.*pHBpMg.*pMgBpNa.*pNaBpCa - PhiK.*pCaBpNa.*pHBpCa.*pMgBpH.*pNaBpMg + PhiK.*pCaBpNa.*pHBpMg.*pMgBpH.*pNaBpCa - PhiK.*pCaBpNa.*pHBpMg.*pMgBpCa.*pNaBpH - PhiK.*pCaBpMg.*pHBpCa.*pMgBpNa.*pNaBpH - PhiK.*pCaBpMg.*pHBpNa.*pMgBpH.*pNaBpCa + PhiK.*pCaBpMg.*pHBpNa.*pMgBpCa.*pNaBpH - PhiH.*pCaBpH.*pKBpCa.*pMgBpNa.*pNaBpMg + PhiH.*pCaBpH.*pKBpNa.*pMgBpCa.*pNaBpMg + PhiH.*pCaBpH.*pKBpMg.*pMgBpNa.*pNaBpCa - PhiH.*pCaBpNa.*pKBpH.*pMgBpCa.*pNaBpMg + PhiH.*pCaBpNa.*pKBpCa.*pMgBpH.*pNaBpMg - PhiH.*pCaBpNa.*pKBpMg.*pMgBpH.*pNaBpCa + PhiH.*pCaBpNa.*pKBpMg.*pMgBpCa.*pNaBpH - PhiH.*pCaBpMg.*pKBpH.*pMgBpNa.*pNaBpCa + PhiH.*pCaBpMg.*pKBpCa.*pMgBpNa.*pNaBpH + PhiH.*pCaBpMg.*pKBpNa.*pMgBpH.*pNaBpCa - PhiH.*pCaBpMg.*pKBpNa.*pMgBpCa.*pNaBpH - PhiCa.*pHBpCa.*pKBpH.*pMgBpNa.*pNaBpMg + PhiCa.*pHBpCa.*pKBpNa.*pMgBpH.*pNaBpMg + PhiCa.*pHBpCa.*pKBpMg.*pMgBpNa.*pNaBpH + PhiCa.*pHBpNa.*pKBpH.*pMgBpCa.*pNaBpMg - PhiCa.*pHBpNa.*pKBpCa.*pMgBpH.*pNaBpMg + PhiCa.*pHBpNa.*pKBpMg.*pMgBpH.*pNaBpCa - PhiCa.*pHBpNa.*pKBpMg.*pMgBpCa.*pNaBpH + PhiCa.*pHBpMg.*pKBpH.*pMgBpNa.*pNaBpCa - PhiCa.*pHBpMg.*pKBpCa.*pMgBpNa.*pNaBpH - PhiCa.*pHBpMg.*pKBpNa.*pMgBpH.*pNaBpCa + PhiCa.*pHBpMg.*pKBpNa.*pMgBpCa.*pNaBpH + DCa.*DH.*DMg.*DNa.*PhiK - DCa.*DH.*DMg.*PhiNa.*pKBpNa - DCa.*DH.*DNa.*PhiMg.*pKBpMg - DCa.*DMg.*DNa.*PhiH.*pKBpH - DH.*DMg.*DNa.*PhiCa.*pKBpCa)./(DH.*DK.*DMg.*pCaBpNa.*pNaBpCa + DH.*DK.*DNa.*pCaBpMg.*pMgBpCa + DCa.*DH.*DK.*pMgBpNa.*pNaBpMg + DK.*DMg.*DNa.*pCaBpH.*pHBpCa + DH.*DMg.*DNa.*pCaBpK.*pKBpCa + DCa.*DK.*DMg.*pHBpNa.*pNaBpH + DCa.*DH.*DMg.*pKBpNa.*pNaBpK + DCa.*DK.*DNa.*pHBpMg.*pMgBpH + DCa.*DH.*DNa.*pKBpMg.*pMgBpK + DCa.*DMg.*DNa.*pHBpK.*pKBpH - DH.*DK.*pCaBpNa.*pMgBpCa.*pNaBpMg - DH.*DK.*pCaBpMg.*pMgBpNa.*pNaBpCa - DK.*DMg.*pCaBpH.*pHBpNa.*pNaBpCa - DK.*DMg.*pCaBpNa.*pHBpCa.*pNaBpH - DH.*DMg.*pCaBpK.*pKBpNa.*pNaBpCa - DH.*DMg.*pCaBpNa.*pKBpCa.*pNaBpK - DK.*DNa.*pCaBpH.*pHBpMg.*pMgBpCa - DK.*DNa.*pCaBpMg.*pHBpCa.*pMgBpH - DH.*DNa.*pCaBpK.*pKBpMg.*pMgBpCa - DH.*DNa.*pCaBpMg.*pKBpCa.*pMgBpK - DCa.*DK.*pHBpNa.*pMgBpH.*pNaBpMg - DCa.*DK.*pHBpMg.*pMgBpNa.*pNaBpH - DCa.*DH.*pKBpNa.*pMgBpK.*pNaBpMg - DCa.*DH.*pKBpMg.*pMgBpNa.*pNaBpK - DMg.*DNa.*pCaBpH.*pHBpK.*pKBpCa - DMg.*DNa.*pCaBpK.*pHBpCa.*pKBpH - DCa.*DMg.*pHBpK.*pKBpNa.*pNaBpH - DCa.*DMg.*pHBpNa.*pKBpH.*pNaBpK - DCa.*DNa.*pHBpK.*pKBpMg.*pMgBpH - DCa.*DNa.*pHBpMg.*pKBpH.*pMgBpK - DK.*pCaBpH.*pHBpCa.*pMgBpNa.*pNaBpMg + DK.*pCaBpH.*pHBpNa.*pMgBpCa.*pNaBpMg + DK.*pCaBpH.*pHBpMg.*pMgBpNa.*pNaBpCa + DK.*pCaBpNa.*pHBpCa.*pMgBpH.*pNaBpMg - DK.*pCaBpNa.*pHBpMg.*pMgBpH.*pNaBpCa + DK.*pCaBpNa.*pHBpMg.*pMgBpCa.*pNaBpH + DK.*pCaBpMg.*pHBpCa.*pMgBpNa.*pNaBpH + DK.*pCaBpMg.*pHBpNa.*pMgBpH.*pNaBpCa - DK.*pCaBpMg.*pHBpNa.*pMgBpCa.*pNaBpH - DH.*pCaBpK.*pKBpCa.*pMgBpNa.*pNaBpMg + DH.*pCaBpK.*pKBpNa.*pMgBpCa.*pNaBpMg + DH.*pCaBpK.*pKBpMg.*pMgBpNa.*pNaBpCa + DH.*pCaBpNa.*pKBpCa.*pMgBpK.*pNaBpMg - DH.*pCaBpNa.*pKBpMg.*pMgBpK.*pNaBpCa + DH.*pCaBpNa.*pKBpMg.*pMgBpCa.*pNaBpK + DH.*pCaBpMg.*pKBpCa.*pMgBpNa.*pNaBpK + DH.*pCaBpMg.*pKBpNa.*pMgBpK.*pNaBpCa - DH.*pCaBpMg.*pKBpNa.*pMgBpCa.*pNaBpK + DMg.*pCaBpH.*pHBpK.*pKBpNa.*pNaBpCa - DMg.*pCaBpH.*pHBpCa.*pKBpNa.*pNaBpK + DMg.*pCaBpH.*pHBpNa.*pKBpCa.*pNaBpK + DMg.*pCaBpK.*pHBpCa.*pKBpNa.*pNaBpH + DMg.*pCaBpK.*pHBpNa.*pKBpH.*pNaBpCa - DMg.*pCaBpK.*pHBpNa.*pKBpCa.*pNaBpH - DMg.*pCaBpNa.*pHBpK.*pKBpH.*pNaBpCa + DMg.*pCaBpNa.*pHBpK.*pKBpCa.*pNaBpH + DMg.*pCaBpNa.*pHBpCa.*pKBpH.*pNaBpK + DNa.*pCaBpH.*pHBpK.*pKBpMg.*pMgBpCa - DNa.*pCaBpH.*pHBpCa.*pKBpMg.*pMgBpK + DNa.*pCaBpH.*pHBpMg.*pKBpCa.*pMgBpK + DNa.*pCaBpK.*pHBpCa.*pKBpMg.*pMgBpH + DNa.*pCaBpK.*pHBpMg.*pKBpH.*pMgBpCa - DNa.*pCaBpK.*pHBpMg.*pKBpCa.*pMgBpH - DNa.*pCaBpMg.*pHBpK.*pKBpH.*pMgBpCa + DNa.*pCaBpMg.*pHBpK.*pKBpCa.*pMgBpH + DNa.*pCaBpMg.*pHBpCa.*pKBpH.*pMgBpK - DCa.*pHBpK.*pKBpH.*pMgBpNa.*pNaBpMg + DCa.*pHBpK.*pKBpNa.*pMgBpH.*pNaBpMg + DCa.*pHBpK.*pKBpMg.*pMgBpNa.*pNaBpH + DCa.*pHBpNa.*pKBpH.*pMgBpK.*pNaBpMg + DCa.*pHBpNa.*pKBpMg.*pMgBpH.*pNaBpK - DCa.*pHBpNa.*pKBpMg.*pMgBpK.*pNaBpH + DCa.*pHBpMg.*pKBpH.*pMgBpNa.*pNaBpK - DCa.*pHBpMg.*pKBpNa.*pMgBpH.*pNaBpK + DCa.*pHBpMg.*pKBpNa.*pMgBpK.*pNaBpH + pCaBpH.*pHBpK.*pKBpCa.*pMgBpNa.*pNaBpMg - pCaBpH.*pHBpK.*pKBpNa.*pMgBpCa.*pNaBpMg - pCaBpH.*pHBpK.*pKBpMg.*pMgBpNa.*pNaBpCa + pCaBpH.*pHBpCa.*pKBpNa.*pMgBpK.*pNaBpMg + pCaBpH.*pHBpCa.*pKBpMg.*pMgBpNa.*pNaBpK - pCaBpH.*pHBpNa.*pKBpCa.*pMgBpK.*pNaBpMg + pCaBpH.*pHBpNa.*pKBpMg.*pMgBpK.*pNaBpCa - pCaBpH.*pHBpNa.*pKBpMg.*pMgBpCa.*pNaBpK - pCaBpH.*pHBpMg.*pKBpCa.*pMgBpNa.*pNaBpK - pCaBpH.*pHBpMg.*pKBpNa.*pMgBpK.*pNaBpCa + pCaBpH.*pHBpMg.*pKBpNa.*pMgBpCa.*pNaBpK + pCaBpK.*pHBpCa.*pKBpH.*pMgBpNa.*pNaBpMg - pCaBpK.*pHBpCa.*pKBpNa.*pMgBpH.*pNaBpMg - pCaBpK.*pHBpCa.*pKBpMg.*pMgBpNa.*pNaBpH - pCaBpK.*pHBpNa.*pKBpH.*pMgBpCa.*pNaBpMg + pCaBpK.*pHBpNa.*pKBpCa.*pMgBpH.*pNaBpMg - pCaBpK.*pHBpNa.*pKBpMg.*pMgBpH.*pNaBpCa + pCaBpK.*pHBpNa.*pKBpMg.*pMgBpCa.*pNaBpH - pCaBpK.*pHBpMg.*pKBpH.*pMgBpNa.*pNaBpCa + pCaBpK.*pHBpMg.*pKBpCa.*pMgBpNa.*pNaBpH + pCaBpK.*pHBpMg.*pKBpNa.*pMgBpH.*pNaBpCa - pCaBpK.*pHBpMg.*pKBpNa.*pMgBpCa.*pNaBpH + pCaBpNa.*pHBpK.*pKBpH.*pMgBpCa.*pNaBpMg - pCaBpNa.*pHBpK.*pKBpCa.*pMgBpH.*pNaBpMg + pCaBpNa.*pHBpK.*pKBpMg.*pMgBpH.*pNaBpCa - pCaBpNa.*pHBpK.*pKBpMg.*pMgBpCa.*pNaBpH - pCaBpNa.*pHBpCa.*pKBpH.*pMgBpK.*pNaBpMg - pCaBpNa.*pHBpCa.*pKBpMg.*pMgBpH.*pNaBpK + pCaBpNa.*pHBpCa.*pKBpMg.*pMgBpK.*pNaBpH + pCaBpNa.*pHBpMg.*pKBpH.*pMgBpK.*pNaBpCa - pCaBpNa.*pHBpMg.*pKBpH.*pMgBpCa.*pNaBpK + pCaBpNa.*pHBpMg.*pKBpCa.*pMgBpH.*pNaBpK - pCaBpNa.*pHBpMg.*pKBpCa.*pMgBpK.*pNaBpH + pCaBpMg.*pHBpK.*pKBpH.*pMgBpNa.*pNaBpCa - pCaBpMg.*pHBpK.*pKBpCa.*pMgBpNa.*pNaBpH - pCaBpMg.*pHBpK.*pKBpNa.*pMgBpH.*pNaBpCa + pCaBpMg.*pHBpK.*pKBpNa.*pMgBpCa.*pNaBpH - pCaBpMg.*pHBpCa.*pKBpH.*pMgBpNa.*pNaBpK + pCaBpMg.*pHBpCa.*pKBpNa.*pMgBpH.*pNaBpK - pCaBpMg.*pHBpCa.*pKBpNa.*pMgBpK.*pNaBpH - pCaBpMg.*pHBpNa.*pKBpH.*pMgBpK.*pNaBpCa + pCaBpMg.*pHBpNa.*pKBpH.*pMgBpCa.*pNaBpK - pCaBpMg.*pHBpNa.*pKBpCa.*pMgBpH.*pNaBpK + pCaBpMg.*pHBpNa.*pKBpCa.*pMgBpK.*pNaBpH - DCa.*DH.*DK.*DMg.*DNa);
dNadt = (DH.*DK.*PhiNa.*pCaBpMg.*pMgBpCa + DK.*DMg.*PhiNa.*pCaBpH.*pHBpCa + DH.*DMg.*PhiNa.*pCaBpK.*pKBpCa + DCa.*DK.*PhiNa.*pHBpMg.*pMgBpH + DCa.*DH.*PhiNa.*pKBpMg.*pMgBpK + DCa.*DMg.*PhiNa.*pHBpK.*pKBpH - DH.*DK.*PhiMg.*pCaBpMg.*pNaBpCa - DCa.*DK.*PhiMg.*pHBpMg.*pNaBpH - DCa.*DH.*PhiMg.*pKBpMg.*pNaBpK - DH.*DMg.*PhiK.*pCaBpK.*pNaBpCa - DCa.*DH.*PhiK.*pMgBpK.*pNaBpMg - DCa.*DMg.*PhiK.*pHBpK.*pNaBpH - DK.*DMg.*PhiH.*pCaBpH.*pNaBpCa - DCa.*DK.*PhiH.*pMgBpH.*pNaBpMg - DCa.*DMg.*PhiH.*pKBpH.*pNaBpK - DH.*DK.*PhiCa.*pMgBpCa.*pNaBpMg - DK.*DMg.*PhiCa.*pHBpCa.*pNaBpH - DH.*DMg.*PhiCa.*pKBpCa.*pNaBpK - DK.*PhiNa.*pCaBpH.*pHBpMg.*pMgBpCa - DK.*PhiNa.*pCaBpMg.*pHBpCa.*pMgBpH - DH.*PhiNa.*pCaBpK.*pKBpMg.*pMgBpCa - DH.*PhiNa.*pCaBpMg.*pKBpCa.*pMgBpK - DMg.*PhiNa.*pCaBpH.*pHBpK.*pKBpCa - DMg.*PhiNa.*pCaBpK.*pHBpCa.*pKBpH - DCa.*PhiNa.*pHBpK.*pKBpMg.*pMgBpH - DCa.*PhiNa.*pHBpMg.*pKBpH.*pMgBpK - DK.*PhiMg.*pCaBpH.*pHBpCa.*pNaBpMg + DK.*PhiMg.*pCaBpH.*pHBpMg.*pNaBpCa + DK.*PhiMg.*pCaBpMg.*pHBpCa.*pNaBpH - DH.*PhiMg.*pCaBpK.*pKBpCa.*pNaBpMg + DH.*PhiMg.*pCaBpK.*pKBpMg.*pNaBpCa + DH.*PhiMg.*pCaBpMg.*pKBpCa.*pNaBpK - DCa.*PhiMg.*pHBpK.*pKBpH.*pNaBpMg + DCa.*PhiMg.*pHBpK.*pKBpMg.*pNaBpH + DCa.*PhiMg.*pHBpMg.*pKBpH.*pNaBpK + DH.*PhiK.*pCaBpK.*pMgBpCa.*pNaBpMg + DH.*PhiK.*pCaBpMg.*pMgBpK.*pNaBpCa - DH.*PhiK.*pCaBpMg.*pMgBpCa.*pNaBpK + DMg.*PhiK.*pCaBpH.*pHBpK.*pNaBpCa - DMg.*PhiK.*pCaBpH.*pHBpCa.*pNaBpK + DMg.*PhiK.*pCaBpK.*pHBpCa.*pNaBpH + DCa.*PhiK.*pHBpK.*pMgBpH.*pNaBpMg - DCa.*PhiK.*pHBpMg.*pMgBpH.*pNaBpK + DCa.*PhiK.*pHBpMg.*pMgBpK.*pNaBpH + DK.*PhiH.*pCaBpH.*pMgBpCa.*pNaBpMg + DK.*PhiH.*pCaBpMg.*pMgBpH.*pNaBpCa - DK.*PhiH.*pCaBpMg.*pMgBpCa.*pNaBpH + DMg.*PhiH.*pCaBpH.*pKBpCa.*pNaBpK + DMg.*PhiH.*pCaBpK.*pKBpH.*pNaBpCa - DMg.*PhiH.*pCaBpK.*pKBpCa.*pNaBpH + DCa.*PhiH.*pKBpH.*pMgBpK.*pNaBpMg + DCa.*PhiH.*pKBpMg.*pMgBpH.*pNaBpK - DCa.*PhiH.*pKBpMg.*pMgBpK.*pNaBpH + DK.*PhiCa.*pHBpCa.*pMgBpH.*pNaBpMg - DK.*PhiCa.*pHBpMg.*pMgBpH.*pNaBpCa + DK.*PhiCa.*pHBpMg.*pMgBpCa.*pNaBpH + DH.*PhiCa.*pKBpCa.*pMgBpK.*pNaBpMg - DH.*PhiCa.*pKBpMg.*pMgBpK.*pNaBpCa + DH.*PhiCa.*pKBpMg.*pMgBpCa.*pNaBpK - DMg.*PhiCa.*pHBpK.*pKBpH.*pNaBpCa + DMg.*PhiCa.*pHBpK.*pKBpCa.*pNaBpH + DMg.*PhiCa.*pHBpCa.*pKBpH.*pNaBpK + PhiNa.*pCaBpH.*pHBpK.*pKBpMg.*pMgBpCa - PhiNa.*pCaBpH.*pHBpCa.*pKBpMg.*pMgBpK + PhiNa.*pCaBpH.*pHBpMg.*pKBpCa.*pMgBpK + PhiNa.*pCaBpK.*pHBpCa.*pKBpMg.*pMgBpH + PhiNa.*pCaBpK.*pHBpMg.*pKBpH.*pMgBpCa - PhiNa.*pCaBpK.*pHBpMg.*pKBpCa.*pMgBpH - PhiNa.*pCaBpMg.*pHBpK.*pKBpH.*pMgBpCa + PhiNa.*pCaBpMg.*pHBpK.*pKBpCa.*pMgBpH + PhiNa.*pCaBpMg.*pHBpCa.*pKBpH.*pMgBpK + PhiMg.*pCaBpH.*pHBpK.*pKBpCa.*pNaBpMg - PhiMg.*pCaBpH.*pHBpK.*pKBpMg.*pNaBpCa + PhiMg.*pCaBpH.*pHBpCa.*pKBpMg.*pNaBpK - PhiMg.*pCaBpH.*pHBpMg.*pKBpCa.*pNaBpK + PhiMg.*pCaBpK.*pHBpCa.*pKBpH.*pNaBpMg - PhiMg.*pCaBpK.*pHBpCa.*pKBpMg.*pNaBpH - PhiMg.*pCaBpK.*pHBpMg.*pKBpH.*pNaBpCa + PhiMg.*pCaBpK.*pHBpMg.*pKBpCa.*pNaBpH + PhiMg.*pCaBpMg.*pHBpK.*pKBpH.*pNaBpCa - PhiMg.*pCaBpMg.*pHBpK.*pKBpCa.*pNaBpH - PhiMg.*pCaBpMg.*pHBpCa.*pKBpH.*pNaBpK - PhiK.*pCaBpH.*pHBpK.*pMgBpCa.*pNaBpMg + PhiK.*pCaBpH.*pHBpCa.*pMgBpK.*pNaBpMg - PhiK.*pCaBpH.*pHBpMg.*pMgBpK.*pNaBpCa + PhiK.*pCaBpH.*pHBpMg.*pMgBpCa.*pNaBpK - PhiK.*pCaBpK.*pHBpCa.*pMgBpH.*pNaBpMg + PhiK.*pCaBpK.*pHBpMg.*pMgBpH.*pNaBpCa - PhiK.*pCaBpK.*pHBpMg.*pMgBpCa.*pNaBpH - PhiK.*pCaBpMg.*pHBpK.*pMgBpH.*pNaBpCa + PhiK.*pCaBpMg.*pHBpK.*pMgBpCa.*pNaBpH + PhiK.*pCaBpMg.*pHBpCa.*pMgBpH.*pNaBpK - PhiK.*pCaBpMg.*pHBpCa.*pMgBpK.*pNaBpH - PhiH.*pCaBpH.*pKBpCa.*pMgBpK.*pNaBpMg + PhiH.*pCaBpH.*pKBpMg.*pMgBpK.*pNaBpCa - PhiH.*pCaBpH.*pKBpMg.*pMgBpCa.*pNaBpK - PhiH.*pCaBpK.*pKBpH.*pMgBpCa.*pNaBpMg + PhiH.*pCaBpK.*pKBpCa.*pMgBpH.*pNaBpMg - PhiH.*pCaBpK.*pKBpMg.*pMgBpH.*pNaBpCa + PhiH.*pCaBpK.*pKBpMg.*pMgBpCa.*pNaBpH - PhiH.*pCaBpMg.*pKBpH.*pMgBpK.*pNaBpCa + PhiH.*pCaBpMg.*pKBpH.*pMgBpCa.*pNaBpK - PhiH.*pCaBpMg.*pKBpCa.*pMgBpH.*pNaBpK + PhiH.*pCaBpMg.*pKBpCa.*pMgBpK.*pNaBpH + PhiCa.*pHBpK.*pKBpH.*pMgBpCa.*pNaBpMg - PhiCa.*pHBpK.*pKBpCa.*pMgBpH.*pNaBpMg + PhiCa.*pHBpK.*pKBpMg.*pMgBpH.*pNaBpCa - PhiCa.*pHBpK.*pKBpMg.*pMgBpCa.*pNaBpH - PhiCa.*pHBpCa.*pKBpH.*pMgBpK.*pNaBpMg - PhiCa.*pHBpCa.*pKBpMg.*pMgBpH.*pNaBpK + PhiCa.*pHBpCa.*pKBpMg.*pMgBpK.*pNaBpH + PhiCa.*pHBpMg.*pKBpH.*pMgBpK.*pNaBpCa - PhiCa.*pHBpMg.*pKBpH.*pMgBpCa.*pNaBpK + PhiCa.*pHBpMg.*pKBpCa.*pMgBpH.*pNaBpK - PhiCa.*pHBpMg.*pKBpCa.*pMgBpK.*pNaBpH - DCa.*DH.*DK.*DMg.*PhiNa + DCa.*DH.*DK.*PhiMg.*pNaBpMg + DCa.*DH.*DMg.*PhiK.*pNaBpK + DCa.*DK.*DMg.*PhiH.*pNaBpH + DH.*DK.*DMg.*PhiCa.*pNaBpCa)./(DH.*DK.*DMg.*pCaBpNa.*pNaBpCa + DH.*DK.*DNa.*pCaBpMg.*pMgBpCa + DCa.*DH.*DK.*pMgBpNa.*pNaBpMg + DK.*DMg.*DNa.*pCaBpH.*pHBpCa + DH.*DMg.*DNa.*pCaBpK.*pKBpCa + DCa.*DK.*DMg.*pHBpNa.*pNaBpH + DCa.*DH.*DMg.*pKBpNa.*pNaBpK + DCa.*DK.*DNa.*pHBpMg.*pMgBpH + DCa.*DH.*DNa.*pKBpMg.*pMgBpK + DCa.*DMg.*DNa.*pHBpK.*pKBpH - DH.*DK.*pCaBpNa.*pMgBpCa.*pNaBpMg - DH.*DK.*pCaBpMg.*pMgBpNa.*pNaBpCa - DK.*DMg.*pCaBpH.*pHBpNa.*pNaBpCa - DK.*DMg.*pCaBpNa.*pHBpCa.*pNaBpH - DH.*DMg.*pCaBpK.*pKBpNa.*pNaBpCa - DH.*DMg.*pCaBpNa.*pKBpCa.*pNaBpK - DK.*DNa.*pCaBpH.*pHBpMg.*pMgBpCa - DK.*DNa.*pCaBpMg.*pHBpCa.*pMgBpH - DH.*DNa.*pCaBpK.*pKBpMg.*pMgBpCa - DH.*DNa.*pCaBpMg.*pKBpCa.*pMgBpK - DCa.*DK.*pHBpNa.*pMgBpH.*pNaBpMg - DCa.*DK.*pHBpMg.*pMgBpNa.*pNaBpH - DCa.*DH.*pKBpNa.*pMgBpK.*pNaBpMg - DCa.*DH.*pKBpMg.*pMgBpNa.*pNaBpK - DMg.*DNa.*pCaBpH.*pHBpK.*pKBpCa - DMg.*DNa.*pCaBpK.*pHBpCa.*pKBpH - DCa.*DMg.*pHBpK.*pKBpNa.*pNaBpH - DCa.*DMg.*pHBpNa.*pKBpH.*pNaBpK - DCa.*DNa.*pHBpK.*pKBpMg.*pMgBpH - DCa.*DNa.*pHBpMg.*pKBpH.*pMgBpK - DK.*pCaBpH.*pHBpCa.*pMgBpNa.*pNaBpMg + DK.*pCaBpH.*pHBpNa.*pMgBpCa.*pNaBpMg + DK.*pCaBpH.*pHBpMg.*pMgBpNa.*pNaBpCa + DK.*pCaBpNa.*pHBpCa.*pMgBpH.*pNaBpMg - DK.*pCaBpNa.*pHBpMg.*pMgBpH.*pNaBpCa + DK.*pCaBpNa.*pHBpMg.*pMgBpCa.*pNaBpH + DK.*pCaBpMg.*pHBpCa.*pMgBpNa.*pNaBpH + DK.*pCaBpMg.*pHBpNa.*pMgBpH.*pNaBpCa - DK.*pCaBpMg.*pHBpNa.*pMgBpCa.*pNaBpH - DH.*pCaBpK.*pKBpCa.*pMgBpNa.*pNaBpMg + DH.*pCaBpK.*pKBpNa.*pMgBpCa.*pNaBpMg + DH.*pCaBpK.*pKBpMg.*pMgBpNa.*pNaBpCa + DH.*pCaBpNa.*pKBpCa.*pMgBpK.*pNaBpMg - DH.*pCaBpNa.*pKBpMg.*pMgBpK.*pNaBpCa + DH.*pCaBpNa.*pKBpMg.*pMgBpCa.*pNaBpK + DH.*pCaBpMg.*pKBpCa.*pMgBpNa.*pNaBpK + DH.*pCaBpMg.*pKBpNa.*pMgBpK.*pNaBpCa - DH.*pCaBpMg.*pKBpNa.*pMgBpCa.*pNaBpK + DMg.*pCaBpH.*pHBpK.*pKBpNa.*pNaBpCa - DMg.*pCaBpH.*pHBpCa.*pKBpNa.*pNaBpK + DMg.*pCaBpH.*pHBpNa.*pKBpCa.*pNaBpK + DMg.*pCaBpK.*pHBpCa.*pKBpNa.*pNaBpH + DMg.*pCaBpK.*pHBpNa.*pKBpH.*pNaBpCa - DMg.*pCaBpK.*pHBpNa.*pKBpCa.*pNaBpH - DMg.*pCaBpNa.*pHBpK.*pKBpH.*pNaBpCa + DMg.*pCaBpNa.*pHBpK.*pKBpCa.*pNaBpH + DMg.*pCaBpNa.*pHBpCa.*pKBpH.*pNaBpK + DNa.*pCaBpH.*pHBpK.*pKBpMg.*pMgBpCa - DNa.*pCaBpH.*pHBpCa.*pKBpMg.*pMgBpK + DNa.*pCaBpH.*pHBpMg.*pKBpCa.*pMgBpK + DNa.*pCaBpK.*pHBpCa.*pKBpMg.*pMgBpH + DNa.*pCaBpK.*pHBpMg.*pKBpH.*pMgBpCa - DNa.*pCaBpK.*pHBpMg.*pKBpCa.*pMgBpH - DNa.*pCaBpMg.*pHBpK.*pKBpH.*pMgBpCa + DNa.*pCaBpMg.*pHBpK.*pKBpCa.*pMgBpH + DNa.*pCaBpMg.*pHBpCa.*pKBpH.*pMgBpK - DCa.*pHBpK.*pKBpH.*pMgBpNa.*pNaBpMg + DCa.*pHBpK.*pKBpNa.*pMgBpH.*pNaBpMg + DCa.*pHBpK.*pKBpMg.*pMgBpNa.*pNaBpH + DCa.*pHBpNa.*pKBpH.*pMgBpK.*pNaBpMg + DCa.*pHBpNa.*pKBpMg.*pMgBpH.*pNaBpK - DCa.*pHBpNa.*pKBpMg.*pMgBpK.*pNaBpH + DCa.*pHBpMg.*pKBpH.*pMgBpNa.*pNaBpK - DCa.*pHBpMg.*pKBpNa.*pMgBpH.*pNaBpK + DCa.*pHBpMg.*pKBpNa.*pMgBpK.*pNaBpH + pCaBpH.*pHBpK.*pKBpCa.*pMgBpNa.*pNaBpMg - pCaBpH.*pHBpK.*pKBpNa.*pMgBpCa.*pNaBpMg - pCaBpH.*pHBpK.*pKBpMg.*pMgBpNa.*pNaBpCa + pCaBpH.*pHBpCa.*pKBpNa.*pMgBpK.*pNaBpMg + pCaBpH.*pHBpCa.*pKBpMg.*pMgBpNa.*pNaBpK - pCaBpH.*pHBpNa.*pKBpCa.*pMgBpK.*pNaBpMg + pCaBpH.*pHBpNa.*pKBpMg.*pMgBpK.*pNaBpCa - pCaBpH.*pHBpNa.*pKBpMg.*pMgBpCa.*pNaBpK - pCaBpH.*pHBpMg.*pKBpCa.*pMgBpNa.*pNaBpK - pCaBpH.*pHBpMg.*pKBpNa.*pMgBpK.*pNaBpCa + pCaBpH.*pHBpMg.*pKBpNa.*pMgBpCa.*pNaBpK + pCaBpK.*pHBpCa.*pKBpH.*pMgBpNa.*pNaBpMg - pCaBpK.*pHBpCa.*pKBpNa.*pMgBpH.*pNaBpMg - pCaBpK.*pHBpCa.*pKBpMg.*pMgBpNa.*pNaBpH - pCaBpK.*pHBpNa.*pKBpH.*pMgBpCa.*pNaBpMg + pCaBpK.*pHBpNa.*pKBpCa.*pMgBpH.*pNaBpMg - pCaBpK.*pHBpNa.*pKBpMg.*pMgBpH.*pNaBpCa + pCaBpK.*pHBpNa.*pKBpMg.*pMgBpCa.*pNaBpH - pCaBpK.*pHBpMg.*pKBpH.*pMgBpNa.*pNaBpCa + pCaBpK.*pHBpMg.*pKBpCa.*pMgBpNa.*pNaBpH + pCaBpK.*pHBpMg.*pKBpNa.*pMgBpH.*pNaBpCa - pCaBpK.*pHBpMg.*pKBpNa.*pMgBpCa.*pNaBpH + pCaBpNa.*pHBpK.*pKBpH.*pMgBpCa.*pNaBpMg - pCaBpNa.*pHBpK.*pKBpCa.*pMgBpH.*pNaBpMg + pCaBpNa.*pHBpK.*pKBpMg.*pMgBpH.*pNaBpCa - pCaBpNa.*pHBpK.*pKBpMg.*pMgBpCa.*pNaBpH - pCaBpNa.*pHBpCa.*pKBpH.*pMgBpK.*pNaBpMg - pCaBpNa.*pHBpCa.*pKBpMg.*pMgBpH.*pNaBpK + pCaBpNa.*pHBpCa.*pKBpMg.*pMgBpK.*pNaBpH + pCaBpNa.*pHBpMg.*pKBpH.*pMgBpK.*pNaBpCa - pCaBpNa.*pHBpMg.*pKBpH.*pMgBpCa.*pNaBpK + pCaBpNa.*pHBpMg.*pKBpCa.*pMgBpH.*pNaBpK - pCaBpNa.*pHBpMg.*pKBpCa.*pMgBpK.*pNaBpH + pCaBpMg.*pHBpK.*pKBpH.*pMgBpNa.*pNaBpCa - pCaBpMg.*pHBpK.*pKBpCa.*pMgBpNa.*pNaBpH - pCaBpMg.*pHBpK.*pKBpNa.*pMgBpH.*pNaBpCa + pCaBpMg.*pHBpK.*pKBpNa.*pMgBpCa.*pNaBpH - pCaBpMg.*pHBpCa.*pKBpH.*pMgBpNa.*pNaBpK + pCaBpMg.*pHBpCa.*pKBpNa.*pMgBpH.*pNaBpK - pCaBpMg.*pHBpCa.*pKBpNa.*pMgBpK.*pNaBpH - pCaBpMg.*pHBpNa.*pKBpH.*pMgBpK.*pNaBpCa + pCaBpMg.*pHBpNa.*pKBpH.*pMgBpCa.*pNaBpK - pCaBpMg.*pHBpNa.*pKBpCa.*pMgBpH.*pNaBpK + pCaBpMg.*pHBpNa.*pKBpCa.*pMgBpK.*pNaBpH - DCa.*DH.*DK.*DMg.*DNa);
dMgdt = -(DH.*DK.*PhiNa.*pCaBpNa.*pMgBpCa + DCa.*DK.*PhiNa.*pHBpNa.*pMgBpH + DCa.*DH.*PhiNa.*pKBpNa.*pMgBpK - DH.*DK.*PhiMg.*pCaBpNa.*pNaBpCa - DK.*DNa.*PhiMg.*pCaBpH.*pHBpCa - DH.*DNa.*PhiMg.*pCaBpK.*pKBpCa - DCa.*DK.*PhiMg.*pHBpNa.*pNaBpH - DCa.*DH.*PhiMg.*pKBpNa.*pNaBpK - DCa.*DNa.*PhiMg.*pHBpK.*pKBpH + DH.*DNa.*PhiK.*pCaBpK.*pMgBpCa + DCa.*DH.*PhiK.*pMgBpNa.*pNaBpK + DCa.*DNa.*PhiK.*pHBpK.*pMgBpH + DK.*DNa.*PhiH.*pCaBpH.*pMgBpCa + DCa.*DK.*PhiH.*pMgBpNa.*pNaBpH + DCa.*DNa.*PhiH.*pKBpH.*pMgBpK + DH.*DK.*PhiCa.*pMgBpNa.*pNaBpCa + DK.*DNa.*PhiCa.*pHBpCa.*pMgBpH + DH.*DNa.*PhiCa.*pKBpCa.*pMgBpK + DK.*PhiNa.*pCaBpH.*pHBpCa.*pMgBpNa - DK.*PhiNa.*pCaBpH.*pHBpNa.*pMgBpCa - DK.*PhiNa.*pCaBpNa.*pHBpCa.*pMgBpH + DH.*PhiNa.*pCaBpK.*pKBpCa.*pMgBpNa - DH.*PhiNa.*pCaBpK.*pKBpNa.*pMgBpCa - DH.*PhiNa.*pCaBpNa.*pKBpCa.*pMgBpK + DCa.*PhiNa.*pHBpK.*pKBpH.*pMgBpNa - DCa.*PhiNa.*pHBpK.*pKBpNa.*pMgBpH - DCa.*PhiNa.*pHBpNa.*pKBpH.*pMgBpK + DK.*PhiMg.*pCaBpH.*pHBpNa.*pNaBpCa + DK.*PhiMg.*pCaBpNa.*pHBpCa.*pNaBpH + DH.*PhiMg.*pCaBpK.*pKBpNa.*pNaBpCa + DH.*PhiMg.*pCaBpNa.*pKBpCa.*pNaBpK + DNa.*PhiMg.*pCaBpH.*pHBpK.*pKBpCa + DNa.*PhiMg.*pCaBpK.*pHBpCa.*pKBpH + DCa.*PhiMg.*pHBpK.*pKBpNa.*pNaBpH + DCa.*PhiMg.*pHBpNa.*pKBpH.*pNaBpK - DH.*PhiK.*pCaBpK.*pMgBpNa.*pNaBpCa + DH.*PhiK.*pCaBpNa.*pMgBpK.*pNaBpCa - DH.*PhiK.*pCaBpNa.*pMgBpCa.*pNaBpK - DNa.*PhiK.*pCaBpH.*pHBpK.*pMgBpCa + DNa.*PhiK.*pCaBpH.*pHBpCa.*pMgBpK - DNa.*PhiK.*pCaBpK.*pHBpCa.*pMgBpH - DCa.*PhiK.*pHBpK.*pMgBpNa.*pNaBpH - DCa.*PhiK.*pHBpNa.*pMgBpH.*pNaBpK + DCa.*PhiK.*pHBpNa.*pMgBpK.*pNaBpH - DK.*PhiH.*pCaBpH.*pMgBpNa.*pNaBpCa + DK.*PhiH.*pCaBpNa.*pMgBpH.*pNaBpCa - DK.*PhiH.*pCaBpNa.*pMgBpCa.*pNaBpH - DNa.*PhiH.*pCaBpH.*pKBpCa.*pMgBpK - DNa.*PhiH.*pCaBpK.*pKBpH.*pMgBpCa + DNa.*PhiH.*pCaBpK.*pKBpCa.*pMgBpH - DCa.*PhiH.*pKBpH.*pMgBpNa.*pNaBpK + DCa.*PhiH.*pKBpNa.*pMgBpH.*pNaBpK - DCa.*PhiH.*pKBpNa.*pMgBpK.*pNaBpH - DK.*PhiCa.*pHBpCa.*pMgBpNa.*pNaBpH - DK.*PhiCa.*pHBpNa.*pMgBpH.*pNaBpCa + DK.*PhiCa.*pHBpNa.*pMgBpCa.*pNaBpH - DH.*PhiCa.*pKBpCa.*pMgBpNa.*pNaBpK - DH.*PhiCa.*pKBpNa.*pMgBpK.*pNaBpCa + DH.*PhiCa.*pKBpNa.*pMgBpCa.*pNaBpK + DNa.*PhiCa.*pHBpK.*pKBpH.*pMgBpCa - DNa.*PhiCa.*pHBpK.*pKBpCa.*pMgBpH - DNa.*PhiCa.*pHBpCa.*pKBpH.*pMgBpK - PhiNa.*pCaBpH.*pHBpK.*pKBpCa.*pMgBpNa + PhiNa.*pCaBpH.*pHBpK.*pKBpNa.*pMgBpCa - PhiNa.*pCaBpH.*pHBpCa.*pKBpNa.*pMgBpK + PhiNa.*pCaBpH.*pHBpNa.*pKBpCa.*pMgBpK - PhiNa.*pCaBpK.*pHBpCa.*pKBpH.*pMgBpNa + PhiNa.*pCaBpK.*pHBpCa.*pKBpNa.*pMgBpH + PhiNa.*pCaBpK.*pHBpNa.*pKBpH.*pMgBpCa - PhiNa.*pCaBpK.*pHBpNa.*pKBpCa.*pMgBpH - PhiNa.*pCaBpNa.*pHBpK.*pKBpH.*pMgBpCa + PhiNa.*pCaBpNa.*pHBpK.*pKBpCa.*pMgBpH + PhiNa.*pCaBpNa.*pHBpCa.*pKBpH.*pMgBpK - PhiMg.*pCaBpH.*pHBpK.*pKBpNa.*pNaBpCa + PhiMg.*pCaBpH.*pHBpCa.*pKBpNa.*pNaBpK - PhiMg.*pCaBpH.*pHBpNa.*pKBpCa.*pNaBpK - PhiMg.*pCaBpK.*pHBpCa.*pKBpNa.*pNaBpH - PhiMg.*pCaBpK.*pHBpNa.*pKBpH.*pNaBpCa + PhiMg.*pCaBpK.*pHBpNa.*pKBpCa.*pNaBpH + PhiMg.*pCaBpNa.*pHBpK.*pKBpH.*pNaBpCa - PhiMg.*pCaBpNa.*pHBpK.*pKBpCa.*pNaBpH - PhiMg.*pCaBpNa.*pHBpCa.*pKBpH.*pNaBpK + PhiK.*pCaBpH.*pHBpK.*pMgBpNa.*pNaBpCa - PhiK.*pCaBpH.*pHBpCa.*pMgBpNa.*pNaBpK - PhiK.*pCaBpH.*pHBpNa.*pMgBpK.*pNaBpCa + PhiK.*pCaBpH.*pHBpNa.*pMgBpCa.*pNaBpK + PhiK.*pCaBpK.*pHBpCa.*pMgBpNa.*pNaBpH + PhiK.*pCaBpK.*pHBpNa.*pMgBpH.*pNaBpCa - PhiK.*pCaBpK.*pHBpNa.*pMgBpCa.*pNaBpH - PhiK.*pCaBpNa.*pHBpK.*pMgBpH.*pNaBpCa + PhiK.*pCaBpNa.*pHBpK.*pMgBpCa.*pNaBpH + PhiK.*pCaBpNa.*pHBpCa.*pMgBpH.*pNaBpK - PhiK.*pCaBpNa.*pHBpCa.*pMgBpK.*pNaBpH + PhiH.*pCaBpH.*pKBpCa.*pMgBpNa.*pNaBpK + PhiH.*pCaBpH.*pKBpNa.*pMgBpK.*pNaBpCa - PhiH.*pCaBpH.*pKBpNa.*pMgBpCa.*pNaBpK + PhiH.*pCaBpK.*pKBpH.*pMgBpNa.*pNaBpCa - PhiH.*pCaBpK.*pKBpCa.*pMgBpNa.*pNaBpH - PhiH.*pCaBpK.*pKBpNa.*pMgBpH.*pNaBpCa + PhiH.*pCaBpK.*pKBpNa.*pMgBpCa.*pNaBpH - PhiH.*pCaBpNa.*pKBpH.*pMgBpK.*pNaBpCa + PhiH.*pCaBpNa.*pKBpH.*pMgBpCa.*pNaBpK - PhiH.*pCaBpNa.*pKBpCa.*pMgBpH.*pNaBpK + PhiH.*pCaBpNa.*pKBpCa.*pMgBpK.*pNaBpH - PhiCa.*pHBpK.*pKBpH.*pMgBpNa.*pNaBpCa + PhiCa.*pHBpK.*pKBpCa.*pMgBpNa.*pNaBpH + PhiCa.*pHBpK.*pKBpNa.*pMgBpH.*pNaBpCa - PhiCa.*pHBpK.*pKBpNa.*pMgBpCa.*pNaBpH + PhiCa.*pHBpCa.*pKBpH.*pMgBpNa.*pNaBpK - PhiCa.*pHBpCa.*pKBpNa.*pMgBpH.*pNaBpK + PhiCa.*pHBpCa.*pKBpNa.*pMgBpK.*pNaBpH + PhiCa.*pHBpNa.*pKBpH.*pMgBpK.*pNaBpCa - PhiCa.*pHBpNa.*pKBpH.*pMgBpCa.*pNaBpK + PhiCa.*pHBpNa.*pKBpCa.*pMgBpH.*pNaBpK - PhiCa.*pHBpNa.*pKBpCa.*pMgBpK.*pNaBpH + DCa.*DH.*DK.*DNa.*PhiMg - DCa.*DH.*DK.*PhiNa.*pMgBpNa - DCa.*DH.*DNa.*PhiK.*pMgBpK - DCa.*DK.*DNa.*PhiH.*pMgBpH - DH.*DK.*DNa.*PhiCa.*pMgBpCa)./(DH.*DK.*DMg.*pCaBpNa.*pNaBpCa + DH.*DK.*DNa.*pCaBpMg.*pMgBpCa + DCa.*DH.*DK.*pMgBpNa.*pNaBpMg + DK.*DMg.*DNa.*pCaBpH.*pHBpCa + DH.*DMg.*DNa.*pCaBpK.*pKBpCa + DCa.*DK.*DMg.*pHBpNa.*pNaBpH + DCa.*DH.*DMg.*pKBpNa.*pNaBpK + DCa.*DK.*DNa.*pHBpMg.*pMgBpH + DCa.*DH.*DNa.*pKBpMg.*pMgBpK + DCa.*DMg.*DNa.*pHBpK.*pKBpH - DH.*DK.*pCaBpNa.*pMgBpCa.*pNaBpMg - DH.*DK.*pCaBpMg.*pMgBpNa.*pNaBpCa - DK.*DMg.*pCaBpH.*pHBpNa.*pNaBpCa - DK.*DMg.*pCaBpNa.*pHBpCa.*pNaBpH - DH.*DMg.*pCaBpK.*pKBpNa.*pNaBpCa - DH.*DMg.*pCaBpNa.*pKBpCa.*pNaBpK - DK.*DNa.*pCaBpH.*pHBpMg.*pMgBpCa - DK.*DNa.*pCaBpMg.*pHBpCa.*pMgBpH - DH.*DNa.*pCaBpK.*pKBpMg.*pMgBpCa - DH.*DNa.*pCaBpMg.*pKBpCa.*pMgBpK - DCa.*DK.*pHBpNa.*pMgBpH.*pNaBpMg - DCa.*DK.*pHBpMg.*pMgBpNa.*pNaBpH - DCa.*DH.*pKBpNa.*pMgBpK.*pNaBpMg - DCa.*DH.*pKBpMg.*pMgBpNa.*pNaBpK - DMg.*DNa.*pCaBpH.*pHBpK.*pKBpCa - DMg.*DNa.*pCaBpK.*pHBpCa.*pKBpH - DCa.*DMg.*pHBpK.*pKBpNa.*pNaBpH - DCa.*DMg.*pHBpNa.*pKBpH.*pNaBpK - DCa.*DNa.*pHBpK.*pKBpMg.*pMgBpH - DCa.*DNa.*pHBpMg.*pKBpH.*pMgBpK - DK.*pCaBpH.*pHBpCa.*pMgBpNa.*pNaBpMg + DK.*pCaBpH.*pHBpNa.*pMgBpCa.*pNaBpMg + DK.*pCaBpH.*pHBpMg.*pMgBpNa.*pNaBpCa + DK.*pCaBpNa.*pHBpCa.*pMgBpH.*pNaBpMg - DK.*pCaBpNa.*pHBpMg.*pMgBpH.*pNaBpCa + DK.*pCaBpNa.*pHBpMg.*pMgBpCa.*pNaBpH + DK.*pCaBpMg.*pHBpCa.*pMgBpNa.*pNaBpH + DK.*pCaBpMg.*pHBpNa.*pMgBpH.*pNaBpCa - DK.*pCaBpMg.*pHBpNa.*pMgBpCa.*pNaBpH - DH.*pCaBpK.*pKBpCa.*pMgBpNa.*pNaBpMg + DH.*pCaBpK.*pKBpNa.*pMgBpCa.*pNaBpMg + DH.*pCaBpK.*pKBpMg.*pMgBpNa.*pNaBpCa + DH.*pCaBpNa.*pKBpCa.*pMgBpK.*pNaBpMg - DH.*pCaBpNa.*pKBpMg.*pMgBpK.*pNaBpCa + DH.*pCaBpNa.*pKBpMg.*pMgBpCa.*pNaBpK + DH.*pCaBpMg.*pKBpCa.*pMgBpNa.*pNaBpK + DH.*pCaBpMg.*pKBpNa.*pMgBpK.*pNaBpCa - DH.*pCaBpMg.*pKBpNa.*pMgBpCa.*pNaBpK + DMg.*pCaBpH.*pHBpK.*pKBpNa.*pNaBpCa - DMg.*pCaBpH.*pHBpCa.*pKBpNa.*pNaBpK + DMg.*pCaBpH.*pHBpNa.*pKBpCa.*pNaBpK + DMg.*pCaBpK.*pHBpCa.*pKBpNa.*pNaBpH + DMg.*pCaBpK.*pHBpNa.*pKBpH.*pNaBpCa - DMg.*pCaBpK.*pHBpNa.*pKBpCa.*pNaBpH - DMg.*pCaBpNa.*pHBpK.*pKBpH.*pNaBpCa + DMg.*pCaBpNa.*pHBpK.*pKBpCa.*pNaBpH + DMg.*pCaBpNa.*pHBpCa.*pKBpH.*pNaBpK + DNa.*pCaBpH.*pHBpK.*pKBpMg.*pMgBpCa - DNa.*pCaBpH.*pHBpCa.*pKBpMg.*pMgBpK + DNa.*pCaBpH.*pHBpMg.*pKBpCa.*pMgBpK + DNa.*pCaBpK.*pHBpCa.*pKBpMg.*pMgBpH + DNa.*pCaBpK.*pHBpMg.*pKBpH.*pMgBpCa - DNa.*pCaBpK.*pHBpMg.*pKBpCa.*pMgBpH - DNa.*pCaBpMg.*pHBpK.*pKBpH.*pMgBpCa + DNa.*pCaBpMg.*pHBpK.*pKBpCa.*pMgBpH + DNa.*pCaBpMg.*pHBpCa.*pKBpH.*pMgBpK - DCa.*pHBpK.*pKBpH.*pMgBpNa.*pNaBpMg + DCa.*pHBpK.*pKBpNa.*pMgBpH.*pNaBpMg + DCa.*pHBpK.*pKBpMg.*pMgBpNa.*pNaBpH + DCa.*pHBpNa.*pKBpH.*pMgBpK.*pNaBpMg + DCa.*pHBpNa.*pKBpMg.*pMgBpH.*pNaBpK - DCa.*pHBpNa.*pKBpMg.*pMgBpK.*pNaBpH + DCa.*pHBpMg.*pKBpH.*pMgBpNa.*pNaBpK - DCa.*pHBpMg.*pKBpNa.*pMgBpH.*pNaBpK + DCa.*pHBpMg.*pKBpNa.*pMgBpK.*pNaBpH + pCaBpH.*pHBpK.*pKBpCa.*pMgBpNa.*pNaBpMg - pCaBpH.*pHBpK.*pKBpNa.*pMgBpCa.*pNaBpMg - pCaBpH.*pHBpK.*pKBpMg.*pMgBpNa.*pNaBpCa + pCaBpH.*pHBpCa.*pKBpNa.*pMgBpK.*pNaBpMg + pCaBpH.*pHBpCa.*pKBpMg.*pMgBpNa.*pNaBpK - pCaBpH.*pHBpNa.*pKBpCa.*pMgBpK.*pNaBpMg + pCaBpH.*pHBpNa.*pKBpMg.*pMgBpK.*pNaBpCa - pCaBpH.*pHBpNa.*pKBpMg.*pMgBpCa.*pNaBpK - pCaBpH.*pHBpMg.*pKBpCa.*pMgBpNa.*pNaBpK - pCaBpH.*pHBpMg.*pKBpNa.*pMgBpK.*pNaBpCa + pCaBpH.*pHBpMg.*pKBpNa.*pMgBpCa.*pNaBpK + pCaBpK.*pHBpCa.*pKBpH.*pMgBpNa.*pNaBpMg - pCaBpK.*pHBpCa.*pKBpNa.*pMgBpH.*pNaBpMg - pCaBpK.*pHBpCa.*pKBpMg.*pMgBpNa.*pNaBpH - pCaBpK.*pHBpNa.*pKBpH.*pMgBpCa.*pNaBpMg + pCaBpK.*pHBpNa.*pKBpCa.*pMgBpH.*pNaBpMg - pCaBpK.*pHBpNa.*pKBpMg.*pMgBpH.*pNaBpCa + pCaBpK.*pHBpNa.*pKBpMg.*pMgBpCa.*pNaBpH - pCaBpK.*pHBpMg.*pKBpH.*pMgBpNa.*pNaBpCa + pCaBpK.*pHBpMg.*pKBpCa.*pMgBpNa.*pNaBpH + pCaBpK.*pHBpMg.*pKBpNa.*pMgBpH.*pNaBpCa - pCaBpK.*pHBpMg.*pKBpNa.*pMgBpCa.*pNaBpH + pCaBpNa.*pHBpK.*pKBpH.*pMgBpCa.*pNaBpMg - pCaBpNa.*pHBpK.*pKBpCa.*pMgBpH.*pNaBpMg + pCaBpNa.*pHBpK.*pKBpMg.*pMgBpH.*pNaBpCa - pCaBpNa.*pHBpK.*pKBpMg.*pMgBpCa.*pNaBpH - pCaBpNa.*pHBpCa.*pKBpH.*pMgBpK.*pNaBpMg - pCaBpNa.*pHBpCa.*pKBpMg.*pMgBpH.*pNaBpK + pCaBpNa.*pHBpCa.*pKBpMg.*pMgBpK.*pNaBpH + pCaBpNa.*pHBpMg.*pKBpH.*pMgBpK.*pNaBpCa - pCaBpNa.*pHBpMg.*pKBpH.*pMgBpCa.*pNaBpK + pCaBpNa.*pHBpMg.*pKBpCa.*pMgBpH.*pNaBpK - pCaBpNa.*pHBpMg.*pKBpCa.*pMgBpK.*pNaBpH + pCaBpMg.*pHBpK.*pKBpH.*pMgBpNa.*pNaBpCa - pCaBpMg.*pHBpK.*pKBpCa.*pMgBpNa.*pNaBpH - pCaBpMg.*pHBpK.*pKBpNa.*pMgBpH.*pNaBpCa + pCaBpMg.*pHBpK.*pKBpNa.*pMgBpCa.*pNaBpH - pCaBpMg.*pHBpCa.*pKBpH.*pMgBpNa.*pNaBpK + pCaBpMg.*pHBpCa.*pKBpNa.*pMgBpH.*pNaBpK - pCaBpMg.*pHBpCa.*pKBpNa.*pMgBpK.*pNaBpH - pCaBpMg.*pHBpNa.*pKBpH.*pMgBpK.*pNaBpCa + pCaBpMg.*pHBpNa.*pKBpH.*pMgBpCa.*pNaBpK - pCaBpMg.*pHBpNa.*pKBpCa.*pMgBpH.*pNaBpK + pCaBpMg.*pHBpNa.*pKBpCa.*pMgBpK.*pNaBpH - DCa.*DH.*DK.*DMg.*DNa);
dCadt = -(DH.*DK.*PhiNa.*pCaBpMg.*pMgBpNa + DK.*DMg.*PhiNa.*pCaBpH.*pHBpNa + DH.*DMg.*PhiNa.*pCaBpK.*pKBpNa + DH.*DK.*PhiMg.*pCaBpNa.*pNaBpMg + DK.*DNa.*PhiMg.*pCaBpH.*pHBpMg + DH.*DNa.*PhiMg.*pCaBpK.*pKBpMg + DH.*DMg.*PhiK.*pCaBpNa.*pNaBpK + DH.*DNa.*PhiK.*pCaBpMg.*pMgBpK + DMg.*DNa.*PhiK.*pCaBpH.*pHBpK + DK.*DMg.*PhiH.*pCaBpNa.*pNaBpH + DK.*DNa.*PhiH.*pCaBpMg.*pMgBpH + DMg.*DNa.*PhiH.*pCaBpK.*pKBpH - DH.*DK.*PhiCa.*pMgBpNa.*pNaBpMg - DK.*DMg.*PhiCa.*pHBpNa.*pNaBpH - DH.*DMg.*PhiCa.*pKBpNa.*pNaBpK - DK.*DNa.*PhiCa.*pHBpMg.*pMgBpH - DH.*DNa.*PhiCa.*pKBpMg.*pMgBpK - DMg.*DNa.*PhiCa.*pHBpK.*pKBpH - DK.*PhiNa.*pCaBpH.*pHBpMg.*pMgBpNa + DK.*PhiNa.*pCaBpNa.*pHBpMg.*pMgBpH - DK.*PhiNa.*pCaBpMg.*pHBpNa.*pMgBpH - DH.*PhiNa.*pCaBpK.*pKBpMg.*pMgBpNa + DH.*PhiNa.*pCaBpNa.*pKBpMg.*pMgBpK - DH.*PhiNa.*pCaBpMg.*pKBpNa.*pMgBpK - DMg.*PhiNa.*pCaBpH.*pHBpK.*pKBpNa - DMg.*PhiNa.*pCaBpK.*pHBpNa.*pKBpH + DMg.*PhiNa.*pCaBpNa.*pHBpK.*pKBpH - DK.*PhiMg.*pCaBpH.*pHBpNa.*pNaBpMg - DK.*PhiMg.*pCaBpNa.*pHBpMg.*pNaBpH + DK.*PhiMg.*pCaBpMg.*pHBpNa.*pNaBpH - DH.*PhiMg.*pCaBpK.*pKBpNa.*pNaBpMg - DH.*PhiMg.*pCaBpNa.*pKBpMg.*pNaBpK + DH.*PhiMg.*pCaBpMg.*pKBpNa.*pNaBpK - DNa.*PhiMg.*pCaBpH.*pHBpK.*pKBpMg - DNa.*PhiMg.*pCaBpK.*pHBpMg.*pKBpH + DNa.*PhiMg.*pCaBpMg.*pHBpK.*pKBpH + DH.*PhiK.*pCaBpK.*pMgBpNa.*pNaBpMg - DH.*PhiK.*pCaBpNa.*pMgBpK.*pNaBpMg - DH.*PhiK.*pCaBpMg.*pMgBpNa.*pNaBpK - DMg.*PhiK.*pCaBpH.*pHBpNa.*pNaBpK + DMg.*PhiK.*pCaBpK.*pHBpNa.*pNaBpH - DMg.*PhiK.*pCaBpNa.*pHBpK.*pNaBpH - DNa.*PhiK.*pCaBpH.*pHBpMg.*pMgBpK + DNa.*PhiK.*pCaBpK.*pHBpMg.*pMgBpH - DNa.*PhiK.*pCaBpMg.*pHBpK.*pMgBpH + DK.*PhiH.*pCaBpH.*pMgBpNa.*pNaBpMg - DK.*PhiH.*pCaBpNa.*pMgBpH.*pNaBpMg - DK.*PhiH.*pCaBpMg.*pMgBpNa.*pNaBpH + DMg.*PhiH.*pCaBpH.*pKBpNa.*pNaBpK - DMg.*PhiH.*pCaBpK.*pKBpNa.*pNaBpH - DMg.*PhiH.*pCaBpNa.*pKBpH.*pNaBpK + DNa.*PhiH.*pCaBpH.*pKBpMg.*pMgBpK - DNa.*PhiH.*pCaBpK.*pKBpMg.*pMgBpH - DNa.*PhiH.*pCaBpMg.*pKBpH.*pMgBpK + DK.*PhiCa.*pHBpNa.*pMgBpH.*pNaBpMg + DK.*PhiCa.*pHBpMg.*pMgBpNa.*pNaBpH + DH.*PhiCa.*pKBpNa.*pMgBpK.*pNaBpMg + DH.*PhiCa.*pKBpMg.*pMgBpNa.*pNaBpK + DMg.*PhiCa.*pHBpK.*pKBpNa.*pNaBpH + DMg.*PhiCa.*pHBpNa.*pKBpH.*pNaBpK + DNa.*PhiCa.*pHBpK.*pKBpMg.*pMgBpH + DNa.*PhiCa.*pHBpMg.*pKBpH.*pMgBpK + PhiNa.*pCaBpH.*pHBpK.*pKBpMg.*pMgBpNa - PhiNa.*pCaBpH.*pHBpNa.*pKBpMg.*pMgBpK + PhiNa.*pCaBpH.*pHBpMg.*pKBpNa.*pMgBpK + PhiNa.*pCaBpK.*pHBpNa.*pKBpMg.*pMgBpH + PhiNa.*pCaBpK.*pHBpMg.*pKBpH.*pMgBpNa - PhiNa.*pCaBpK.*pHBpMg.*pKBpNa.*pMgBpH - PhiNa.*pCaBpNa.*pHBpK.*pKBpMg.*pMgBpH - PhiNa.*pCaBpNa.*pHBpMg.*pKBpH.*pMgBpK - PhiNa.*pCaBpMg.*pHBpK.*pKBpH.*pMgBpNa + PhiNa.*pCaBpMg.*pHBpK.*pKBpNa.*pMgBpH + PhiNa.*pCaBpMg.*pHBpNa.*pKBpH.*pMgBpK + PhiMg.*pCaBpH.*pHBpK.*pKBpNa.*pNaBpMg + PhiMg.*pCaBpH.*pHBpNa.*pKBpMg.*pNaBpK - PhiMg.*pCaBpH.*pHBpMg.*pKBpNa.*pNaBpK + PhiMg.*pCaBpK.*pHBpNa.*pKBpH.*pNaBpMg - PhiMg.*pCaBpK.*pHBpNa.*pKBpMg.*pNaBpH + PhiMg.*pCaBpK.*pHBpMg.*pKBpNa.*pNaBpH - PhiMg.*pCaBpNa.*pHBpK.*pKBpH.*pNaBpMg + PhiMg.*pCaBpNa.*pHBpK.*pKBpMg.*pNaBpH + PhiMg.*pCaBpNa.*pHBpMg.*pKBpH.*pNaBpK - PhiMg.*pCaBpMg.*pHBpK.*pKBpNa.*pNaBpH - PhiMg.*pCaBpMg.*pHBpNa.*pKBpH.*pNaBpK - PhiK.*pCaBpH.*pHBpK.*pMgBpNa.*pNaBpMg + PhiK.*pCaBpH.*pHBpNa.*pMgBpK.*pNaBpMg + PhiK.*pCaBpH.*pHBpMg.*pMgBpNa.*pNaBpK - PhiK.*pCaBpK.*pHBpNa.*pMgBpH.*pNaBpMg - PhiK.*pCaBpK.*pHBpMg.*pMgBpNa.*pNaBpH + PhiK.*pCaBpNa.*pHBpK.*pMgBpH.*pNaBpMg - PhiK.*pCaBpNa.*pHBpMg.*pMgBpH.*pNaBpK + PhiK.*pCaBpNa.*pHBpMg.*pMgBpK.*pNaBpH + PhiK.*pCaBpMg.*pHBpK.*pMgBpNa.*pNaBpH + PhiK.*pCaBpMg.*pHBpNa.*pMgBpH.*pNaBpK - PhiK.*pCaBpMg.*pHBpNa.*pMgBpK.*pNaBpH - PhiH.*pCaBpH.*pKBpNa.*pMgBpK.*pNaBpMg - PhiH.*pCaBpH.*pKBpMg.*pMgBpNa.*pNaBpK - PhiH.*pCaBpK.*pKBpH.*pMgBpNa.*pNaBpMg + PhiH.*pCaBpK.*pKBpNa.*pMgBpH.*pNaBpMg + PhiH.*pCaBpK.*pKBpMg.*pMgBpNa.*pNaBpH + PhiH.*pCaBpNa.*pKBpH.*pMgBpK.*pNaBpMg + PhiH.*pCaBpNa.*pKBpMg.*pMgBpH.*pNaBpK - PhiH.*pCaBpNa.*pKBpMg.*pMgBpK.*pNaBpH + PhiH.*pCaBpMg.*pKBpH.*pMgBpNa.*pNaBpK - PhiH.*pCaBpMg.*pKBpNa.*pMgBpH.*pNaBpK + PhiH.*pCaBpMg.*pKBpNa.*pMgBpK.*pNaBpH + PhiCa.*pHBpK.*pKBpH.*pMgBpNa.*pNaBpMg - PhiCa.*pHBpK.*pKBpNa.*pMgBpH.*pNaBpMg - PhiCa.*pHBpK.*pKBpMg.*pMgBpNa.*pNaBpH - PhiCa.*pHBpNa.*pKBpH.*pMgBpK.*pNaBpMg - PhiCa.*pHBpNa.*pKBpMg.*pMgBpH.*pNaBpK + PhiCa.*pHBpNa.*pKBpMg.*pMgBpK.*pNaBpH - PhiCa.*pHBpMg.*pKBpH.*pMgBpNa.*pNaBpK + PhiCa.*pHBpMg.*pKBpNa.*pMgBpH.*pNaBpK - PhiCa.*pHBpMg.*pKBpNa.*pMgBpK.*pNaBpH + DH.*DK.*DMg.*DNa.*PhiCa - DH.*DK.*DMg.*PhiNa.*pCaBpNa - DH.*DK.*DNa.*PhiMg.*pCaBpMg - DH.*DMg.*DNa.*PhiK.*pCaBpK - DK.*DMg.*DNa.*PhiH.*pCaBpH)./(DH.*DK.*DMg.*pCaBpNa.*pNaBpCa + DH.*DK.*DNa.*pCaBpMg.*pMgBpCa + DCa.*DH.*DK.*pMgBpNa.*pNaBpMg + DK.*DMg.*DNa.*pCaBpH.*pHBpCa + DH.*DMg.*DNa.*pCaBpK.*pKBpCa + DCa.*DK.*DMg.*pHBpNa.*pNaBpH + DCa.*DH.*DMg.*pKBpNa.*pNaBpK + DCa.*DK.*DNa.*pHBpMg.*pMgBpH + DCa.*DH.*DNa.*pKBpMg.*pMgBpK + DCa.*DMg.*DNa.*pHBpK.*pKBpH - DH.*DK.*pCaBpNa.*pMgBpCa.*pNaBpMg - DH.*DK.*pCaBpMg.*pMgBpNa.*pNaBpCa - DK.*DMg.*pCaBpH.*pHBpNa.*pNaBpCa - DK.*DMg.*pCaBpNa.*pHBpCa.*pNaBpH - DH.*DMg.*pCaBpK.*pKBpNa.*pNaBpCa - DH.*DMg.*pCaBpNa.*pKBpCa.*pNaBpK - DK.*DNa.*pCaBpH.*pHBpMg.*pMgBpCa - DK.*DNa.*pCaBpMg.*pHBpCa.*pMgBpH - DH.*DNa.*pCaBpK.*pKBpMg.*pMgBpCa - DH.*DNa.*pCaBpMg.*pKBpCa.*pMgBpK - DCa.*DK.*pHBpNa.*pMgBpH.*pNaBpMg - DCa.*DK.*pHBpMg.*pMgBpNa.*pNaBpH - DCa.*DH.*pKBpNa.*pMgBpK.*pNaBpMg - DCa.*DH.*pKBpMg.*pMgBpNa.*pNaBpK - DMg.*DNa.*pCaBpH.*pHBpK.*pKBpCa - DMg.*DNa.*pCaBpK.*pHBpCa.*pKBpH - DCa.*DMg.*pHBpK.*pKBpNa.*pNaBpH - DCa.*DMg.*pHBpNa.*pKBpH.*pNaBpK - DCa.*DNa.*pHBpK.*pKBpMg.*pMgBpH - DCa.*DNa.*pHBpMg.*pKBpH.*pMgBpK - DK.*pCaBpH.*pHBpCa.*pMgBpNa.*pNaBpMg + DK.*pCaBpH.*pHBpNa.*pMgBpCa.*pNaBpMg + DK.*pCaBpH.*pHBpMg.*pMgBpNa.*pNaBpCa + DK.*pCaBpNa.*pHBpCa.*pMgBpH.*pNaBpMg - DK.*pCaBpNa.*pHBpMg.*pMgBpH.*pNaBpCa + DK.*pCaBpNa.*pHBpMg.*pMgBpCa.*pNaBpH + DK.*pCaBpMg.*pHBpCa.*pMgBpNa.*pNaBpH + DK.*pCaBpMg.*pHBpNa.*pMgBpH.*pNaBpCa - DK.*pCaBpMg.*pHBpNa.*pMgBpCa.*pNaBpH - DH.*pCaBpK.*pKBpCa.*pMgBpNa.*pNaBpMg + DH.*pCaBpK.*pKBpNa.*pMgBpCa.*pNaBpMg + DH.*pCaBpK.*pKBpMg.*pMgBpNa.*pNaBpCa + DH.*pCaBpNa.*pKBpCa.*pMgBpK.*pNaBpMg - DH.*pCaBpNa.*pKBpMg.*pMgBpK.*pNaBpCa + DH.*pCaBpNa.*pKBpMg.*pMgBpCa.*pNaBpK + DH.*pCaBpMg.*pKBpCa.*pMgBpNa.*pNaBpK + DH.*pCaBpMg.*pKBpNa.*pMgBpK.*pNaBpCa - DH.*pCaBpMg.*pKBpNa.*pMgBpCa.*pNaBpK + DMg.*pCaBpH.*pHBpK.*pKBpNa.*pNaBpCa - DMg.*pCaBpH.*pHBpCa.*pKBpNa.*pNaBpK + DMg.*pCaBpH.*pHBpNa.*pKBpCa.*pNaBpK + DMg.*pCaBpK.*pHBpCa.*pKBpNa.*pNaBpH + DMg.*pCaBpK.*pHBpNa.*pKBpH.*pNaBpCa - DMg.*pCaBpK.*pHBpNa.*pKBpCa.*pNaBpH - DMg.*pCaBpNa.*pHBpK.*pKBpH.*pNaBpCa + DMg.*pCaBpNa.*pHBpK.*pKBpCa.*pNaBpH + DMg.*pCaBpNa.*pHBpCa.*pKBpH.*pNaBpK + DNa.*pCaBpH.*pHBpK.*pKBpMg.*pMgBpCa - DNa.*pCaBpH.*pHBpCa.*pKBpMg.*pMgBpK + DNa.*pCaBpH.*pHBpMg.*pKBpCa.*pMgBpK + DNa.*pCaBpK.*pHBpCa.*pKBpMg.*pMgBpH + DNa.*pCaBpK.*pHBpMg.*pKBpH.*pMgBpCa - DNa.*pCaBpK.*pHBpMg.*pKBpCa.*pMgBpH - DNa.*pCaBpMg.*pHBpK.*pKBpH.*pMgBpCa + DNa.*pCaBpMg.*pHBpK.*pKBpCa.*pMgBpH + DNa.*pCaBpMg.*pHBpCa.*pKBpH.*pMgBpK - DCa.*pHBpK.*pKBpH.*pMgBpNa.*pNaBpMg + DCa.*pHBpK.*pKBpNa.*pMgBpH.*pNaBpMg + DCa.*pHBpK.*pKBpMg.*pMgBpNa.*pNaBpH + DCa.*pHBpNa.*pKBpH.*pMgBpK.*pNaBpMg + DCa.*pHBpNa.*pKBpMg.*pMgBpH.*pNaBpK - DCa.*pHBpNa.*pKBpMg.*pMgBpK.*pNaBpH + DCa.*pHBpMg.*pKBpH.*pMgBpNa.*pNaBpK - DCa.*pHBpMg.*pKBpNa.*pMgBpH.*pNaBpK + DCa.*pHBpMg.*pKBpNa.*pMgBpK.*pNaBpH + pCaBpH.*pHBpK.*pKBpCa.*pMgBpNa.*pNaBpMg - pCaBpH.*pHBpK.*pKBpNa.*pMgBpCa.*pNaBpMg - pCaBpH.*pHBpK.*pKBpMg.*pMgBpNa.*pNaBpCa + pCaBpH.*pHBpCa.*pKBpNa.*pMgBpK.*pNaBpMg + pCaBpH.*pHBpCa.*pKBpMg.*pMgBpNa.*pNaBpK - pCaBpH.*pHBpNa.*pKBpCa.*pMgBpK.*pNaBpMg + pCaBpH.*pHBpNa.*pKBpMg.*pMgBpK.*pNaBpCa - pCaBpH.*pHBpNa.*pKBpMg.*pMgBpCa.*pNaBpK - pCaBpH.*pHBpMg.*pKBpCa.*pMgBpNa.*pNaBpK - pCaBpH.*pHBpMg.*pKBpNa.*pMgBpK.*pNaBpCa + pCaBpH.*pHBpMg.*pKBpNa.*pMgBpCa.*pNaBpK + pCaBpK.*pHBpCa.*pKBpH.*pMgBpNa.*pNaBpMg - pCaBpK.*pHBpCa.*pKBpNa.*pMgBpH.*pNaBpMg - pCaBpK.*pHBpCa.*pKBpMg.*pMgBpNa.*pNaBpH - pCaBpK.*pHBpNa.*pKBpH.*pMgBpCa.*pNaBpMg + pCaBpK.*pHBpNa.*pKBpCa.*pMgBpH.*pNaBpMg - pCaBpK.*pHBpNa.*pKBpMg.*pMgBpH.*pNaBpCa + pCaBpK.*pHBpNa.*pKBpMg.*pMgBpCa.*pNaBpH - pCaBpK.*pHBpMg.*pKBpH.*pMgBpNa.*pNaBpCa + pCaBpK.*pHBpMg.*pKBpCa.*pMgBpNa.*pNaBpH + pCaBpK.*pHBpMg.*pKBpNa.*pMgBpH.*pNaBpCa - pCaBpK.*pHBpMg.*pKBpNa.*pMgBpCa.*pNaBpH + pCaBpNa.*pHBpK.*pKBpH.*pMgBpCa.*pNaBpMg - pCaBpNa.*pHBpK.*pKBpCa.*pMgBpH.*pNaBpMg + pCaBpNa.*pHBpK.*pKBpMg.*pMgBpH.*pNaBpCa - pCaBpNa.*pHBpK.*pKBpMg.*pMgBpCa.*pNaBpH - pCaBpNa.*pHBpCa.*pKBpH.*pMgBpK.*pNaBpMg - pCaBpNa.*pHBpCa.*pKBpMg.*pMgBpH.*pNaBpK + pCaBpNa.*pHBpCa.*pKBpMg.*pMgBpK.*pNaBpH + pCaBpNa.*pHBpMg.*pKBpH.*pMgBpK.*pNaBpCa - pCaBpNa.*pHBpMg.*pKBpH.*pMgBpCa.*pNaBpK + pCaBpNa.*pHBpMg.*pKBpCa.*pMgBpH.*pNaBpK - pCaBpNa.*pHBpMg.*pKBpCa.*pMgBpK.*pNaBpH + pCaBpMg.*pHBpK.*pKBpH.*pMgBpNa.*pNaBpCa - pCaBpMg.*pHBpK.*pKBpCa.*pMgBpNa.*pNaBpH - pCaBpMg.*pHBpK.*pKBpNa.*pMgBpH.*pNaBpCa + pCaBpMg.*pHBpK.*pKBpNa.*pMgBpCa.*pNaBpH - pCaBpMg.*pHBpCa.*pKBpH.*pMgBpNa.*pNaBpK + pCaBpMg.*pHBpCa.*pKBpNa.*pMgBpH.*pNaBpK - pCaBpMg.*pHBpCa.*pKBpNa.*pMgBpK.*pNaBpH - pCaBpMg.*pHBpNa.*pKBpH.*pMgBpK.*pNaBpCa + pCaBpMg.*pHBpNa.*pKBpH.*pMgBpCa.*pNaBpK - pCaBpMg.*pHBpNa.*pKBpCa.*pMgBpH.*pNaBpK + pCaBpMg.*pHBpNa.*pKBpCa.*pMgBpK.*pNaBpH - DCa.*DH.*DK.*DMg.*DNa);

% M/min b/c Phi
f(p.i.H_c,:) = dHdt;
f(p.i.K_c,:) = dKdt;
f(p.i.Na_c,:) = dNadt;
f(p.i.Mg_c,:) = dMgdt;
f(p.i.Ca_c,:) = dCadt;
end

%% Cation/Ligand Injection funciton used to add bolus addition
% This prevents the need to recompute the free cation concentration using
% binding polynomials
% Arbitrarily forcing injection to be finished within 1 ms to minimize
% impact of model dynamics
% the use of a smooth numerical switch is chosen over a step function to
% provide continuous derivatives

function Jt = InjectReactants(t,x,p,Jt)

d = 1e-3; % rapid injection
r = 1./(1+(t./d).^20); % define numerical switch function

Jt.Inject = p.Inject.R*(r*p.Vol.H2O_cyt./d);

end

%% Binding Polynomial Calculation
% Only 1st order binding species are considered assuming that higher
% order binding species concentractions are neglegible.
% inputs:
% l - ligand of interest
% c - compartment, 1) mtx (mito matrix), 2) ims (imm space), 
%                  3) cyt (cytosol), etc.
% x - state variable vector
% p - parameter structure
% output:
% f - free ligand conc. based on free protons and metal ion conc.

function [f, flag, new_data] = BP(l,c,x,p)

if strcmp(c,'mtx')==1
    % indices for matrix cations
    i=[p.i.H_x,p.i.K_x,p.i.Na_x,p.i.Mg_x,p.i.Ca_x];
elseif strcmp(c,'ims')==1
    % indices for IMS cations replaced w/ cytosolic cations
    i=[p.i.H_c,p.i.K_c,p.i.Na_c,p.i.Mg_c,p.i.Ca_c];
elseif strcmp(c,'cyt')==1
    % indices for cytosolic cations
    i=[p.i.H_c,p.i.K_c,p.i.Na_c,p.i.Mg_c,p.i.Ca_c]; 
else %add further compartments as necessary
    error('currently, only matrix, ims or cytosol arguments are accepted')
end

% define appropriate cations
H = x(i(1),:);
K = x(i(2),:);
Na = x(i(3),:);
Mg = x(i(4),:);
Ca = x(i(5),:);

% this is for vectorized code
f = (1 + (1./p.KD(l,p.if.H))*H + (1./p.KD(l,p.if.K))*K ...
    + (1./p.KD(l,p.if.Na)*Na) + (1./p.KD(l,p.if.Mg))*Mg ...
    + (1./p.KD(l,p.if.Ca))*Ca);

flag=0;

new_data=0;

end

%% Mitochondrial Dehydrogenases
% Qatpase and fNAD feedback (saturable)

function J = MitoDH(x,p)

% Define SVs into user friendly notation
NAD = x(p.i.NAD_x,:);   % M
NADH = x(p.i.NADH_x,:); % M
Pi = x(p.i.Pi_x,:);     % M
ATP = x(p.i.ATP_x,:);   % M
ADP = x(p.i.ADP_x,:);   % M
PYR = x(p.i.PYR_x,:);   % M
OAA = x(p.i.OAA_x,:);   % M

Q = ADP.*Pi./ATP; % feedback ratio

J = p.MitoDH.x.*Q.^p.MitoDH.n./(Q.^p.MitoDH.n + p.MitoDH.K.^p.MitoDH.n) ...
    .*(NAD./(NAD + p.MitoDH.r*NADH)).*(PYR./(PYR+p.MitoDH.Km_PYR)) ...
    .*(OAA./(OAA+p.MitoDH.Km_OAA));

end

%% MDH
% Malate Dehydrogenase (MDH)
% NAD+ + MAL2- = OAA2- + NADH + H+
% NAD + MAL = OAA + NADH (+ H^+)
% A - NAD; B - MAL; P - OAA; Q - NADH
%
% Malate dehydrogenase is the enzyme responsible for the oxidation of
% malate to form oxaloacetate and reducing equivalents for the ETS.
%
% NOTE: The following code was adopted from the TCA cycle model developed
% by Wu et al., 2007 published in JBC using the code from the model version
% Codes_TCAv2.zip obtained from http://bbc.mcw.edu/Computation/models/
% 
% Copyright:
% Jason Bazil, Ann Rundell and Greg Buzzard. Nov 2009
% Rundell Laboratory
% Weldon School of Biomedical Engineering
% Purdue University

function J = MDH(x,p)

% load substrate state variables and convert from nmol/mg to M units
H = x(p.i.H_x,:);       % M
K = x(p.i.K_x,:);       % M
NAD = x(p.i.NAD_x,:);   % M
MAL = x(p.i.MAL_x,:);   % M
OAA = x(p.i.OAA_x,:);   % M
NADH = x(p.i.NADH_x,:); % M
ATP = x(p.i.ATP_x,:);   % M
ADP = x(p.i.ADP_x,:);   % M
AMP = x(p.i.AMP_x,:);   % M
Pi = x(p.i.Pi_x,:);     % M

% redefine state variable names into generalized notation
A = NAD;
B = MAL;
P = OAA;
Q = NADH;

% dG and Keq values
dGr_mdho = p.dGf(p.if.OAA) + p.dGf(p.if.NADH) - p.dGf(p.if.NAD) ...
    - p.dGf(p.if.MAL);
Keq_mdho = exp(-dGr_mdho/p.R./p.T);
% Keq_mdh = Keq_mdho*1/H;
Keq_mdh = Keq_mdho.*(1./H).*(p.BP(p.i.OAA_x,:).*p.BP(p.i.NADH_x,:)) ...
    ./(p.BP(p.i.MAL_x,:).*p.BP(p.i.NAD_x,:));

% Km and Ki values (Molar)
KmA = p.MDH.KmNAD;
KmB = p.MDH.KmMAL;
KmP = p.MDH.KmOAA;
KmQ = p.MDH.KmNADH;
Kia = p.MDH.Kinad;
Kib = p.MDH.Kimal;
Kip = p.MDH.Kioaa;
Kiq = p.MDH.Kinadh;
% % from Kohn et al.
% KiATP = 709.3e-6;
% KiADP = 383.2e-6;
% KiAMP = 793.0e-6;
% from Oza and Shore
KiATP = p.MDH.KiATP;
KiADP = p.MDH.KiADP;
KiAMP = p.MDH.KiAMP;
KmPi = p.MDH.KPi;
bPi = p.MDH.bPi;

% inhibition/activaiton terms
% NOTE: The Wu model states all fL terms include K-bound species, but not
% in their code. This is corrected here.
fATP = ATP.*(1+H./p.KD(p.if.ATP,p.if.H)+K./p.KD(p.if.ATP,p.if.K)) ...
    ./p.BP(p.i.ATP_x,:);
fADP = ADP.*(1+H./p.KD(p.if.ADP,p.if.H)+K./p.KD(p.if.ADP,p.if.K)) ...
    ./p.BP(p.i.ADP_x,:);
fAMP = AMP.*(1+H./p.KD(p.if.AMP,p.if.H)+K./p.KD(p.if.AMP,p.if.K)) ...
    ./p.BP(p.i.AMP_x,:);

ai = 1+fATP./KiATP+fADP./KiADP+fAMP./KiAMP;

% Pi-activation (non-essential, Telegdi 1973)
PiControl = (1 + bPi.*Pi./KmPi)./(1 + Pi./KmPi);

% Vm values
Vmf = p.MDH.Vmaxf;
Vmr = Vmf./Keq_mdh.*(Kiq.*KmP./Kia./KmB);

% total reaction flux
J = (Vmf.*Vmr.*(A.*B - P.*Q./Keq_mdh)).*PiControl ...
    ./(Vmr.*Kia.*KmB.*ai+Vmr.*KmB.*A ...
    +Vmr.*KmA.*ai.*B+Vmf.*KmQ.*ai./Keq_mdh.*P+Vmf.*KmP./Keq_mdh.*Q ...
    +Vmr.*A.*B+Vmf.*KmQ./Kia./Keq_mdh.*A.*P+Vmf./Keq_mdh.*P.*Q ...
    +Vmr.*KmA./Kiq.*B.*Q+Vmr./Kip.*A.*B.*P+Vmf./Kib./Keq_mdh.*B.*P.*Q);

end

%% FH
% Fumarate hydratase (FH) or Fumarase
% FUM2- (+H2O) = MAL2-
% FUM (+ H2O) = MAL
% A - FUM; P - MAL;
%
% Fumarate hydratase is the enzyme responsible for the condensation
% reaction incorporating a water molecule into fumarate to form malate.
%
% NOTE: The following code was adopted from the TCA cycle model developed
% by Wu et al., 2007 published in JBC using the code from the model version
% Codes_TCAv2.zip obtained from http://bbc.mcw.edu/Computation/models/
% 
% Copyright:
% Jason Bazil, Ann Rundell and Greg Buzzard. Nov 2009
% Rundell Laboratory
% Weldon School of Biomedical Engineering
% Purdue University

function J = FH(x,p)

% load substrate state variables and convert from nmol/mg to M units
H = x(p.i.H_x,:);       
K = x(p.i.K_x,:);       
MAL = x(p.i.MAL_x,:);
FUM = x(p.i.FUM_x,:);
CIT = x(p.i.CIT_x,:);
ATP = x(p.i.ATP_x,:);
ADP = x(p.i.ADP_x,:);
GTP = x(p.i.GTP_x,:);
GDP = x(p.i.GDP_x,:);

% redefine state variable names into generalized notation
A = FUM;
P = MAL;

% dG and Keq values
dGr_fumo = p.dGf(p.if.MAL) - p.dGf(p.if.FUM) - p.dGf(p.if.H2O);
Keq_fumo = exp(-dGr_fumo/p.R./p.T);
% Keq_fum = Keq_fumo;
Keq_fum = Keq_fumo.*p.BP(p.i.MAL_x,:)./p.BP(p.i.FUM_x,:);

% Km and Ki values (Molar)
KmA = p.FH.KmFUM;
KmP = p.FH.KmMAL;
% KmA = 2.34e-6;
% KmP = 8e-6;
KiCIT = p.FH.KiCIT;
KiATP = p.FH.KiATP;
KiADP = p.FH.KiADP;
KiGTP = p.FH.KiGTP;
KiGDP = p.FH.KiGDP;

% inhibition coefficients
% NOTE: The Wu model states all fL terms include K-bound species, but not
% in their code. This is corrected here.  
fATP = ATP.*(1+H./p.KD(p.if.ATP,p.if.H)+K./p.KD(p.if.ATP,p.if.K)) ...
    ./p.BP(p.i.ATP_x,:);
fADP = ADP.*(1+H./p.KD(p.if.ADP,p.if.H)+K./p.KD(p.if.ADP,p.if.K)) ...
    ./p.BP(p.i.ADP_x,:);
fGTP = GTP.*(1+H./p.KD(p.if.GTP,p.if.H)+K./p.KD(p.if.GTP,p.if.K)) ...
    ./p.BP(p.i.GTP_x,:);
fGDP = GDP.*(1+H./p.KD(p.if.GDP,p.if.H)+K./p.KD(p.if.GDP,p.if.K)) ...
    ./p.BP(p.i.GDP_x,:);
ai = 1+CIT./KiCIT+fATP./KiATP+fADP./KiADP+fGTP./KiGTP+fGDP./KiGDP;

% Vm values
Vmf = p.FH.Vmaxf;
Vmr = Vmf./Keq_fum.*(KmP./KmA);

% total reaction flux
J =  (Vmf.*Vmr.*A - Vmf.*Vmr.*(P./Keq_fum)) ...
    ./(KmA.*Vmr.*ai+Vmr.*A+Vmf./Keq_fum.*P);

end

%% Malic Enzyme
% decarboxylating

function J = ME(x,p)

J = p.kf_ME.*x(p.i.MAL_x,:).*x(p.i.NAD_x,:);

end

%% CI model that includes ROS pathways (5-State Model)
% assumes Q-binding and SQ formation is rapid
% from Bazil et al. 2014

function varargout = CI(x,p)
%% define state variables
Hp = x(p.i.H_c,:);
Hn = x(p.i.H_x,:);
NAD = x(p.i.NAD_x,:);
NADH = x(p.i.NADH_x,:);
Q = x(p.i.UQ_x,:);
QH2 = x(p.i.UQH2_x,:);
dPsi = x(p.i.dPsi,:);
O2 = x(p.i.O2_x,:);
SO = x(p.i.SO_x,:);
H2O2 = x(p.i.H2O2_x,:);

% NADH = (NADH - p.CI.Kb_NADH - p.CI.B + sqrt((NADH - p.CI.Kb_NADH - p.CI.B).^2 + 4*p.CI.Kb_NADH *NADH))/2;

%% Update midpoint potentials from thermodynamic data
p.CI.Em0.Q_QH2 = (p.dGf(p.if.UQH2) - p.dGf(p.if.UQ))/(-2*p.F); % this is 18.67 mV less than value used for CI parameterization
p.CI.Em0.NADH = (p.dGf(p.if.NADH) - p.dGf(p.if.NAD))/(-2*p.F); % this is 2.3 mV higher than value used for CI parameterization
p.CI.Em0.SO = (p.dGf(p.if.SO) - p.dGf(p.if.O2))/(-1*p.F); % this is 8.3 mV higher than value used for CI parameterization
p.CI.Em0.H2O2 = (p.dGf(p.if.H2O2) - p.dGf(p.if.O2))/(-2*p.F); % this is 8.4 mV less than value used for CI parameterization

%% Update midpoint potentials for 1e- quinone couples (pH 0)
p.CI.Em0.Q_SQ = p.CI.Em0.Q_QH2 + p.R*p.T./p.F./2*log(p.CI.Kstability*1e-14); % mV
p.CI.Em0.SQ_QH2 = 2*p.CI.Em0.Q_QH2 - p.CI.Em0.Q_SQ; % mV

%% Binding Polynomials for Protonated States
PH1 = (1./(1+Hn./p.CI.KiH1));
PH2 = 1;%(Hn./p.CI.KiH2./(1+Hn./p.CI.KiH2));
PH3 = (Hn./p.CI.KiH3./(1+Hn./p.CI.KiH3));
% PH4 = (1./(1+Hn./p.f10));

%% Binding Polynomials for enzyme, substrates, products and regulators
% NADH binding constants
KdNADHo=p.CI.K_NADH_o;
KdNADo=p.CI.K_NAD_o;
KdNADHr=p.CI.K_NADH_r;
KdNADr=p.CI.K_NAD_r;
KdNADHrad=p.CI.K_NADH_rad;
KdNADrad=p.CI.K_NAD_rad;
% Quinone binding constants
KdQH2o = p.CI.K_QH2;
KdQo = p.CI.K_Q;
KdQH2r = p.CI.K_QH2;
KdQr = p.CI.K_Q;
% Binding polynomials
PNo = 1 + NADH./KdNADHo + NAD./KdNADo; % oxidized binding constants
PNr = 1 + NADH./KdNADHr + NAD./KdNADr; % reduced binding constants
PNrad = 1 + NADH./KdNADHrad + NAD./KdNADrad; % reduced binding constants
PQr = 1 + QH2./KdQH2r + Q./KdQr;% + p.rotenone/p.CI.K_rotenone; % reduced binding constants
PQo = 1 + QH2./KdQH2o + Q./KdQo;% + p.rotenone/p.CI.K_rotenone; % oxidized binding constants

muH = p.F*dPsi+p.R*p.T*log(Hp./Hn); % proton chemical potential (kJ/mol)

%% rotenone effects of SO and Q TN at Q-site

p.CI.kfQ_20 = p.CI.kfQ_20./(1+p.rotenone/p.CI.K_rotenone);
p.CI.kfQ_31 = p.CI.kfQ_31./(1+p.rotenone/p.CI.K_rotenone);
p.CI.kfQ_42 = p.CI.kfQ_42./(1+p.rotenone/p.CI.K_rotenone);

p.CI.kfSQSO_10 = p.CI.kfSQSO_10./(1+p.rotenone/p.CI.K_rotenone);
p.CI.kfSQSO_21 = p.CI.kfSQSO_21./(1+p.rotenone/p.CI.K_rotenone);
p.CI.kfSQSO_21b = p.CI.kfSQSO_21b./(1+p.rotenone/p.CI.K_rotenone);
p.CI.kfSQSO_32 = p.CI.kfSQSO_32./(1+p.rotenone/p.CI.K_rotenone);
p.CI.kfSQSO_32b = p.CI.kfSQSO_32b./(1+p.rotenone/p.CI.K_rotenone);
p.CI.kfSQSO_43 = p.CI.kfSQSO_43./(1+p.rotenone/p.CI.K_rotenone);

%% Compute pH Corrected Midpoint potentials
% NADH potential
Em.NADH = p.CI.Em0.NADH + log(10).*p.R.*p.T./p.F./2.*log10(Hn);
% FMN potentials
Em.FMNred = p.CI.Em0.FMN - p.R*p.T./2./p.F*log(Hn./10^-p.CI.FMNred.pK./(1+Hn./10^-p.CI.FMNred.pK)./Hn.^2); % FMNred./FMN, pK1 of fully reduced FMN very high (>10)0)
Em.FMNrad = p.CI.Em0.FMN2 - p.R*p.T./p.F*log((Hn./10^-(p.CI.FMNred.pK)./(1+Hn./10^-p.CI.FMNred.pK))./((Hn./10^-p.CI.FMNrad.pK)./(1+Hn./10^-p.CI.FMNrad.pK))./Hn);  % FMNred./FMNrad
Em.FMNox = p.CI.Em0.FMN1 - p.R*p.T./p.F*log((Hn./10^-p.CI.FMNrad.pK)./(1+Hn./10^-p.CI.FMNrad.pK)./Hn); %FMN./FMNrad
% N2 potential
Em.N2 = p.CI.Em0.N2 - p.R*p.T./p.F*log((Hn+10^-p.CI.N2.pKox)./(Hn+10^-p.CI.N2.pKred));
% superoxide and H2O2 potentials
Em.SO = p.CI.Em0.SO; % Em0 stays the same for pH > 6, pKa of SO is ~4.8
Em.H2O2 = p.CI.Em0.H2O2 + log(10).*p.R.*p.T./p.F.*log10(Hn); % 1st pKa is ~11
% Quinone potentials
Em.Q_QH2 = p.CI.Em0.Q_QH2 + 2*log(10)*p.R*p.T./p.F./2*log10(Hn); % assuming linked to N-side
Em.Q_SQ = p.CI.Em0.Q_SQ; % pH independent
Em.SQ_QH2 = p.CI.Em0.SQ_QH2 + 2*log(10)*p.R*p.T./p.F*log10(Hn); % assuming linked to N-side

%% State Transition Thermodynamics
% NADH-QH2 reductase related
Keq.NADHFMNred = exp((2.*p.F.*(Em.FMNred - Em.NADH))./p.R./p.T); % [Fr][NAD]./[F][NADH][H]
Keq.FMNradN2 = exp((p.F.*(Em.N2 - Em.FMNrad))./p.R./p.T); % [Frad][N2r]./[Fr]./[N2]
Keq.FMNoxN2 = exp((p.F.*(Em.N2 - Em.FMNox))./p.R./p.T); % [F][N2r]./[Frad]./[N2]
Keq.N2SQ = exp((p.F.*(Em.Q_SQ - Em.N2))./p.R./p.T); % [N2][SQ]./[N2r][Q]
Keq.N2QH2 = exp((p.F.*(Em.SQ_QH2 - Em.N2))./p.R./p.T); % [N2][QH2]./[N2r][SQ]

% superoxide and hydrogen peroxide
Keq.FMNredH2O2 = exp((2.*p.F.*(Em.H2O2 - Em.FMNred))./p.R./p.T); % [F][H2O2]./[Fr][O2]
Keq.FMNradO2 = exp((p.F.*(Em.SO - Em.FMNrad))./p.R./p.T); % [Frad][O2.-]./[Fr][O2]
Keq.FMNoxO2 = exp((p.F.*(Em.SO - Em.FMNox))./p.R./p.T); % [F][O2.-]./[Frad][O2]
Keq.N2O2 = exp((p.F.*(Em.SO - Em.N2))./p.R./p.T); % [N2][O2.-]./[N2r][O2]
Keq.SQO2 = exp((p.F.*(Em.SO - Em.Q_SQ))./p.R./p.T); % [Q][O2.-]./[SQ][O2]

%% Substates via Rapid Equilibrium

% substate equilibrium constants
K1 = Keq.FMNradN2; %[Frad][N2r]./[Fr][N2]
K2 = Keq.FMNoxN2; % [F][N2r]./[Frad][N2]
KSQ = Keq.N2SQ.*KdQr; % [N2-SQ]./[N2r-Q], KdQr = [N2r][Q]./[N2r-Q]

KQ = (Q./KdQr./PQr).*KSQ; % [N2-SQ]./[N2r]

% 0e-
PS0 = 1; % total
s0.F_N2 = 1./PS0;

% 1e-
PS1 = (1 + K2.*(1 + KQ)); % total sum(cell2mat(struct2cell(s1)))
s1.Frad_N2 = 1./PS1;
s1.F_N2r = K2.*s1.Frad_N2;
s1.F_N2_SQ = KQ.*s1.F_N2r;

% 2e-
PS2 = (1 + K1.*(1 + KQ.*(1 + K2))); % total sum(cell2mat(struct2cell(s2)))
s2.Fr_N2 = 1./PS2;
s2.Frad_N2r = K1.*s2.Fr_N2;
s2.Frad_N2_SQ = KQ.*s2.Frad_N2r;
s2.F_N2r_SQ = K2.*s2.Frad_N2_SQ;

% 3e-
PS3 = (1 + KQ.*(1 + K1)); % total sum(cell2mat(struct2cell(s3)))
s3.Fr_N2r = 1./PS3;
s3.Fr_N2_SQ = KQ.*s3.Fr_N2r;
s3.Frad_N2r_SQ = K1.*s3.Fr_N2_SQ;

% 4e-
PS4 = 1; % total
s4.Fr_N2r_SQ = 1./PS4;


%% State Transition Rates

% reverse NADH-Q reductase rates
krN0_02 = p.CI.kfNADH_02./Keq.NADHFMNred*KdNADr./KdNADHo.*PH1.*PH2;
krN0_24 = p.CI.kfNADH_24./Keq.NADHFMNred*KdNADr./KdNADHo.*PH1.*PH2;
krN0_13 = p.CI.kfNADH_13./Keq.NADHFMNred*KdNADr./KdNADHo.*PH1.*PH2;
krQ0_20 = p.CI.kfQ_20./Keq.N2QH2*KdQH2o.*(PNr.*PQo./PNo./PQr).*PH3;
krQ0_42 = p.CI.kfQ_42./Keq.N2QH2*KdQH2o.*(PNr.*PQo./PNo./PQr).*PH3;
krQ0_31 = p.CI.kfQ_31./Keq.N2QH2*KdQH2o.*(PNr.*PQo./PNo./PQr).*PH3;

% reverse ROS rates
% O2 and SQ
krSQSO_10 = p.CI.kfSQSO_10./Keq.SQO2;
krSQSO_21 = p.CI.kfSQSO_21./Keq.SQO2;
krSQSO_21b = p.CI.kfSQSO_21b./Keq.SQO2;
krSQSO_32 = p.CI.kfSQSO_32./Keq.SQO2;
krSQSO_32b = p.CI.kfSQSO_32b./Keq.SQO2;
krSQSO_43 = p.CI.kfSQSO_43./Keq.SQO2;
% O2 and N2
krN2rSO_10 = p.CI.kfN2rSO_10./Keq.N2O2;
krN2rSO_21 = p.CI.kfN2rSO_21./Keq.N2O2;
krN2rSO_21b = p.CI.kfN2rSO_21b./Keq.N2O2;
krN2rSO_32 = p.CI.kfN2rSO_32./Keq.N2O2;
krN2rSO_32b = p.CI.kfN2rSO_32b./Keq.N2O2;
krN2rSO_43 = p.CI.kfN2rSO_43./Keq.N2O2;
% O2 and FMNred
krFrSO_21 = p.CI.kfFrSO_21./Keq.FMNradO2;
krFradSO_21b = p.CI.kfFradSO_21b./Keq.FMNradO2;
krFrSO_32 = p.CI.kfFrSO_32./Keq.FMNradO2;
krFrSO_32b = p.CI.kfFrSO_32b./Keq.FMNradO2;
krFrSO_43 = p.CI.kfFrSO_43./Keq.FMNradO2;
% O2 and FMNrad
krFradSO_10 = p.CI.kfFradSO_10./Keq.FMNoxO2;
krFradSO_21 = p.CI.kfFradSO_21./Keq.FMNoxO2;
krFradSO_32 = p.CI.kfFradSO_32./Keq.FMNoxO2;
% H2O2 and FMNred
krH_20 = p.CI.kfH_20./Keq.FMNredH2O2;
krH_31 = p.CI.kfH_31./Keq.FMNredH2O2;
krH_31b = p.CI.kfH_31b./Keq.FMNredH2O2;
krH_42 = p.CI.kfH_42./Keq.FMNredH2O2;

% Nucleotide and pH effects
p.CI.kfH_20 = p.CI.kfH_20./PNr;%./(1+Hn./10^-p.CI.FMNred.pK);
p.CI.kfH_42 = p.CI.kfH_42./PNr;%./(1+Hn./10^-p.CI.FMNred.pK);
p.CI.kfH_31 = p.CI.kfH_31./PNr;%./(1+Hn./10^-p.CI.FMNred.pK);
p.CI.kfH_31b = p.CI.kfH_31b./PNr;%./(1+Hn./10^-p.CI.FMNred.pK);

p.CI.kfFrSO_21 = p.CI.kfFrSO_21./PNr./(1+Hn./10^-p.CI.FMNred.pK);
p.CI.kfFrSO_32 = p.CI.kfFrSO_32./PNr./(1+Hn./10^-p.CI.FMNred.pK);
p.CI.kfFrSO_32b = p.CI.kfFrSO_32b./PNr./(1+Hn./10^-p.CI.FMNred.pK);
p.CI.kfFrSO_43 = p.CI.kfFrSO_43./PNr./(1+Hn./10^-p.CI.FMNred.pK);

p.CI.kfFradSO_10 = p.CI.kfFradSO_10./PNrad./(1+Hn./10^-p.CI.FMNrad.pK);
p.CI.kfFradSO_21 = p.CI.kfFradSO_21./PNrad./(1+Hn./10^-p.CI.FMNrad.pK);
p.CI.kfFradSO_21b = p.CI.kfFradSO_21b./PNrad./(1+Hn./10^-p.CI.FMNrad.pK);
p.CI.kfFradSO_32 = p.CI.kfFradSO_32./PNrad./(1+Hn./10^-p.CI.FMNrad.pK);

p.CI.kfN2rSO_10 = p.CI.kfN2rSO_10./(1+Hn./10^-p.CI.N2.pKred);
p.CI.kfN2rSO_21 = p.CI.kfN2rSO_21./(1+Hn./10^-p.CI.N2.pKred);
p.CI.kfN2rSO_21b = p.CI.kfN2rSO_21b./(1+Hn./10^-p.CI.N2.pKred);
p.CI.kfN2rSO_32 = p.CI.kfN2rSO_32./(1+Hn./10^-p.CI.N2.pKred);
p.CI.kfN2rSO_32b = p.CI.kfN2rSO_32b./(1+Hn./10^-p.CI.N2.pKred);
p.CI.kfN2rSO_43 = p.CI.kfN2rSO_43./(1+Hn./10^-p.CI.N2.pKred);

% net forward rates

% membrane potential dependence
dPsiNf = 1;%exp(-4*p.CI.beta2*muH/p.R/p.T/2);
dPsiNr = 1;%exp(4*(1-p.CI.beta2)*muH/p.R/p.T/2);
dPsiQf = exp(-4*p.CI.beta1*muH/p.R/p.T);
dPsiQr = exp(4*(1-p.CI.beta1)*muH/p.R/p.T);

% NADH transitions
kfNADH_02 = p.CI.kfNADH_02.*dPsiNf.*NADH./KdNADHo./PNo.*(s0.F_N2).*PH1.*PH2;
krNADH_02 = krN0_02.*dPsiNr.*NAD./KdNADr./PNr.*(s2.Fr_N2);
kfNADH_24 = p.CI.kfNADH_24.*dPsiNf.*NADH./KdNADHo./PNo.*(s2.F_N2r_SQ).*PH1.*PH2;
krNADH_24 = krN0_24.*dPsiNr.*NAD./KdNADr./PNr.*(s4.Fr_N2r_SQ);
kfNADH_13 = p.CI.kfNADH_13.*dPsiNf.*NADH./KdNADHo./PNo.*(s1.F_N2r + s1.F_N2_SQ).*PH1.*PH2;
krNADH_13 = krN0_13.*dPsiNr.*NAD./KdNADr./PNr.*(s3.Fr_N2r + s3.Fr_N2_SQ);

% QH2 transitions
kfQ_20 = p.CI.kfQ_20.*dPsiQf.*(s2.F_N2r_SQ).*PH3;
krQ_20 = krQ0_20.*dPsiQr.*QH2./KdQH2o./PQo.*(s0.F_N2);
kfQ_42 = p.CI.kfQ_42.*dPsiQf.*(s4.Fr_N2r_SQ).*PH3;
krQ_42 = krQ0_42.*dPsiQr.*QH2./KdQH2o./PQo.*(s2.Fr_N2);
kfQ_31 = p.CI.kfQ_31.*dPsiQf.*(s3.Frad_N2r_SQ).*PH3;
krQ_31 = krQ0_31.*dPsiQr.*QH2./KdQH2o./PQo.*(s1.Frad_N2);

% H2O2 transitions
kfH2O2_20 = p.CI.kfH_20.*O2.*s2.Fr_N2;
krH2O2_20 = krH_20.*H2O2.*s0.F_N2;

kfH2O2_31 = p.CI.kfH_31.*O2.*s3.Fr_N2r + p.CI.kfH_31b.*O2.*s3.Fr_N2_SQ;
krH2O2_31 = krH_31.*H2O2.*s1.F_N2r + krH_31b.*H2O2.*s1.F_N2_SQ;

kfH2O2_42 = p.CI.kfH_42.*O2.*s4.Fr_N2r_SQ;
krH2O2_42 = krH_42.*H2O2.*(s2.F_N2r_SQ);

% SO transitions
kfSO_10 = O2.*(p.CI.kfSQSO_10.*s1.F_N2_SQ + p.CI.kfFradSO_10.*s1.Frad_N2 + p.CI.kfN2rSO_10.*s1.F_N2r);
krSO_10 = SO.*s0.F_N2.*(krSQSO_10.*(Q./KdQo./PQo) + krFradSO_10 + krN2rSO_10);

kfSO_21 = O2.*((p.CI.kfSQSO_21.*s2.Frad_N2_SQ + p.CI.kfSQSO_21b.*s2.F_N2r_SQ) + p.CI.kfFrSO_21.*s2.Fr_N2 + (p.CI.kfFradSO_21.*s2.Frad_N2r + p.CI.kfFradSO_21b.*s2.Frad_N2_SQ) + (p.CI.kfN2rSO_21.*s2.Frad_N2r + p.CI.kfN2rSO_21b.*s2.F_N2r_SQ));
krSO_21 = SO.*(krFrSO_21.*s1.Frad_N2 + krN2rSO_21.*s1.Frad_N2 + krN2rSO_21b.*s1.F_N2_SQ + krFradSO_21.*s1.F_N2r + krFradSO_21b.*s1.F_N2_SQ + (krSQSO_21.*s1.Frad_N2.*(Q./KdQo./PQo) + krSQSO_21b.*s1.F_N2r.*(Q./KdQr./PQr)));

kfSO_32 = O2.*((p.CI.kfSQSO_32.*s3.Fr_N2_SQ + p.CI.kfSQSO_32b.*s3.Frad_N2r_SQ) + (p.CI.kfFrSO_32.*s3.Fr_N2r + p.CI.kfFrSO_32b.*s3.Fr_N2_SQ) + p.CI.kfFradSO_32.*s3.Frad_N2r_SQ + (p.CI.kfN2rSO_32.*s3.Fr_N2r + p.CI.kfN2rSO_32b.*s3.Frad_N2r_SQ));
krSO_32 = SO.*(krSQSO_32.*s2.Fr_N2.*(Q./KdQo./PQo) + krSQSO_32b.*s2.Frad_N2r.*(Q./KdQr./PQr) + krFradSO_32.*s2.F_N2r_SQ + krFrSO_32.*s2.Frad_N2r + krFrSO_32b.*s2.Frad_N2_SQ + krN2rSO_32.*s2.Fr_N2 + krN2rSO_32b.*s2.Frad_N2_SQ);

kfSO_43 = O2.*((p.CI.kfSQSO_43 + p.CI.kfN2rSO_43).*(s4.Fr_N2r_SQ)  + p.CI.kfFrSO_43.*s4.Fr_N2r_SQ);
krSO_43 = SO.*(krSQSO_43.*s3.Fr_N2r.*(Q./KdQr./PQr) + krFrSO_43.*s3.Frad_N2r_SQ  + krN2rSO_43.*s3.Fr_N2_SQ);
    
% S0 <-> S2
k0_2 = kfNADH_02 + krQ_20 + krH2O2_20;
k2_0 = kfQ_20 + krNADH_02 + kfH2O2_20; 

% S2 <-> S4
k2_4 = kfNADH_24 + krQ_42 + krH2O2_42;
k4_2 = kfQ_42 + krNADH_24 + kfH2O2_42;

% S1 <-> S3
k1_3 = kfNADH_13 + krQ_31 + krH2O2_31;
k3_1 = kfQ_31 + krNADH_13 + kfH2O2_31; 

% S0 <-> S1 (superoxide)
k1_0 = kfSO_10;
k0_1 = krSO_10;

% S2 <-> S1 (superoxide)
k2_1 = kfSO_21;
k1_2 = krSO_21; % if approximateing 3 state model set equal to 1e99

% S3 <-> S2 (superoxide)
k3_2 = kfSO_32;
k2_3 = krSO_32;

% S4 <-> S3 (superoxide)
k4_3 = kfSO_43;
k3_4 = krSO_43; % if approximateing 3 state model set equal to 1e99

%% Steady-State Fractional Occupancies (solved analytically)
S0=(k1_0.*k2_0.*k3_1.*k4_2 + k1_0.*k2_0.*k3_1.*k4_3 + k1_0.*k2_0.*k3_2.*k4_2 + k1_0.*k2_1.*k3_1.*k4_2 + k1_0.*k2_0.*k3_2.*k4_3 + k1_0.*k2_1.*k3_1.*k4_3 + k1_0.*k2_1.*k3_2.*k4_2 + k1_2.*k2_0.*k3_1.*k4_2 + k1_0.*k2_0.*k3_4.*k4_2 + k1_0.*k2_1.*k3_2.*k4_3 + k1_0.*k2_3.*k3_1.*k4_2 + k1_2.*k2_0.*k3_1.*k4_3 + k1_2.*k2_0.*k3_2.*k4_2 + k1_0.*k2_1.*k3_4.*k4_2 + k1_0.*k2_3.*k3_1.*k4_3 + k1_2.*k2_0.*k3_2.*k4_3 + k1_3.*k2_0.*k3_2.*k4_2 + k1_0.*k2_4.*k3_1.*k4_3 + k1_2.*k2_0.*k3_4.*k4_2 + k1_3.*k2_0.*k3_2.*k4_3 + k1_3.*k2_0.*k3_4.*k4_2)./(k0_2.*k1_0.*k2_4.*k3_1 + k0_1.*k1_2.*k2_4.*k3_1 + k0_1.*k1_3.*k2_0.*k3_4 + k0_2.*k1_0.*k2_4.*k3_2 + k0_1.*k1_2.*k2_4.*k3_2 + k0_1.*k1_3.*k2_1.*k3_4 + k0_2.*k1_0.*k2_3.*k3_4 + k0_2.*k1_2.*k2_4.*k3_1 + k0_1.*k1_2.*k2_3.*k3_4 + k0_1.*k1_3.*k2_4.*k3_2 + k0_2.*k1_0.*k2_4.*k3_4 + k0_2.*k1_2.*k2_4.*k3_2 + k0_2.*k1_3.*k2_1.*k3_4 + k0_1.*k1_2.*k2_4.*k3_4 + k0_1.*k1_3.*k2_3.*k3_4 + k0_2.*k1_2.*k2_3.*k3_4 + k0_2.*k1_3.*k2_4.*k3_2 + k0_1.*k1_3.*k2_4.*k3_4 + k0_2.*k1_2.*k2_4.*k3_4 + k0_2.*k1_3.*k2_3.*k3_4 + k0_2.*k1_3.*k2_4.*k3_4 + k0_1.*k1_3.*k2_0.*k4_2 + k0_1.*k1_3.*k2_0.*k4_3 + k0_1.*k1_3.*k2_1.*k4_2 + k0_2.*k1_0.*k2_3.*k4_2 + k0_1.*k1_2.*k2_3.*k4_2 + k0_1.*k1_3.*k2_1.*k4_3 + k0_2.*k1_0.*k2_3.*k4_3 + k0_2.*k1_3.*k2_1.*k4_2 + k0_1.*k1_2.*k2_3.*k4_3 + k0_1.*k1_3.*k2_3.*k4_2 + k0_2.*k1_0.*k2_4.*k4_3 + k0_2.*k1_2.*k2_3.*k4_2 + k0_2.*k1_3.*k2_1.*k4_3 + k0_1.*k1_2.*k2_4.*k4_3 + k0_1.*k1_3.*k2_3.*k4_3 + k0_2.*k1_2.*k2_3.*k4_3 + k0_2.*k1_3.*k2_3.*k4_2 + k0_1.*k1_3.*k2_4.*k4_3 + k0_2.*k1_2.*k2_4.*k4_3 + k0_2.*k1_3.*k2_3.*k4_3 + k0_2.*k1_3.*k2_4.*k4_3 + k0_2.*k1_0.*k3_1.*k4_2 + k0_1.*k1_2.*k3_1.*k4_2 + k0_2.*k1_0.*k3_1.*k4_3 + k0_2.*k1_0.*k3_2.*k4_2 + k0_1.*k1_2.*k3_1.*k4_3 + k0_1.*k1_2.*k3_2.*k4_2 + k0_2.*k1_0.*k3_2.*k4_3 + k0_2.*k1_2.*k3_1.*k4_2 + k0_1.*k1_2.*k3_2.*k4_3 + k0_1.*k1_3.*k3_2.*k4_2 + k0_2.*k1_0.*k3_4.*k4_2 + k0_2.*k1_2.*k3_1.*k4_3 + k0_2.*k1_2.*k3_2.*k4_2 + k0_1.*k1_2.*k3_4.*k4_2 + k0_1.*k1_3.*k3_2.*k4_3 + k0_2.*k1_2.*k3_2.*k4_3 + k0_2.*k1_3.*k3_2.*k4_2 + k0_1.*k1_3.*k3_4.*k4_2 + k0_2.*k1_2.*k3_4.*k4_2 + k0_2.*k1_3.*k3_2.*k4_3 + k0_2.*k1_3.*k3_4.*k4_2 + k0_1.*k2_0.*k3_1.*k4_2 + k0_1.*k2_0.*k3_1.*k4_3 + k0_1.*k2_0.*k3_2.*k4_2 + k0_1.*k2_1.*k3_1.*k4_2 + k0_1.*k2_0.*k3_2.*k4_3 + k0_1.*k2_1.*k3_1.*k4_3 + k0_1.*k2_1.*k3_2.*k4_2 + k0_2.*k2_1.*k3_1.*k4_2 + k0_1.*k2_0.*k3_4.*k4_2 + k0_1.*k2_1.*k3_2.*k4_3 + k0_1.*k2_3.*k3_1.*k4_2 + k0_2.*k2_1.*k3_1.*k4_3 + k0_2.*k2_1.*k3_2.*k4_2 + k0_1.*k2_1.*k3_4.*k4_2 + k0_1.*k2_3.*k3_1.*k4_3 + k0_2.*k2_1.*k3_2.*k4_3 + k0_2.*k2_3.*k3_1.*k4_2 + k0_1.*k2_4.*k3_1.*k4_3 + k0_2.*k2_1.*k3_4.*k4_2 + k0_2.*k2_3.*k3_1.*k4_3 + k0_2.*k2_4.*k3_1.*k4_3 + k1_0.*k2_0.*k3_1.*k4_2 + k1_0.*k2_0.*k3_1.*k4_3 + k1_0.*k2_0.*k3_2.*k4_2 + k1_0.*k2_1.*k3_1.*k4_2 + k1_0.*k2_0.*k3_2.*k4_3 + k1_0.*k2_1.*k3_1.*k4_3 + k1_0.*k2_1.*k3_2.*k4_2 + k1_2.*k2_0.*k3_1.*k4_2 + k1_0.*k2_0.*k3_4.*k4_2 + k1_0.*k2_1.*k3_2.*k4_3 + k1_0.*k2_3.*k3_1.*k4_2 + k1_2.*k2_0.*k3_1.*k4_3 + k1_2.*k2_0.*k3_2.*k4_2 + k1_0.*k2_1.*k3_4.*k4_2 + k1_0.*k2_3.*k3_1.*k4_3 + k1_2.*k2_0.*k3_2.*k4_3 + k1_3.*k2_0.*k3_2.*k4_2 + k1_0.*k2_4.*k3_1.*k4_3 + k1_2.*k2_0.*k3_4.*k4_2 + k1_3.*k2_0.*k3_2.*k4_3 + k1_3.*k2_0.*k3_4.*k4_2);
S1=(k0_1.*k2_0.*k3_1.*k4_2 + k0_1.*k2_0.*k3_1.*k4_3 + k0_1.*k2_0.*k3_2.*k4_2 + k0_1.*k2_1.*k3_1.*k4_2 + k0_1.*k2_0.*k3_2.*k4_3 + k0_1.*k2_1.*k3_1.*k4_3 + k0_1.*k2_1.*k3_2.*k4_2 + k0_2.*k2_1.*k3_1.*k4_2 + k0_1.*k2_0.*k3_4.*k4_2 + k0_1.*k2_1.*k3_2.*k4_3 + k0_1.*k2_3.*k3_1.*k4_2 + k0_2.*k2_1.*k3_1.*k4_3 + k0_2.*k2_1.*k3_2.*k4_2 + k0_1.*k2_1.*k3_4.*k4_2 + k0_1.*k2_3.*k3_1.*k4_3 + k0_2.*k2_1.*k3_2.*k4_3 + k0_2.*k2_3.*k3_1.*k4_2 + k0_1.*k2_4.*k3_1.*k4_3 + k0_2.*k2_1.*k3_4.*k4_2 + k0_2.*k2_3.*k3_1.*k4_3 + k0_2.*k2_4.*k3_1.*k4_3)./(k0_2.*k1_0.*k2_4.*k3_1 + k0_1.*k1_2.*k2_4.*k3_1 + k0_1.*k1_3.*k2_0.*k3_4 + k0_2.*k1_0.*k2_4.*k3_2 + k0_1.*k1_2.*k2_4.*k3_2 + k0_1.*k1_3.*k2_1.*k3_4 + k0_2.*k1_0.*k2_3.*k3_4 + k0_2.*k1_2.*k2_4.*k3_1 + k0_1.*k1_2.*k2_3.*k3_4 + k0_1.*k1_3.*k2_4.*k3_2 + k0_2.*k1_0.*k2_4.*k3_4 + k0_2.*k1_2.*k2_4.*k3_2 + k0_2.*k1_3.*k2_1.*k3_4 + k0_1.*k1_2.*k2_4.*k3_4 + k0_1.*k1_3.*k2_3.*k3_4 + k0_2.*k1_2.*k2_3.*k3_4 + k0_2.*k1_3.*k2_4.*k3_2 + k0_1.*k1_3.*k2_4.*k3_4 + k0_2.*k1_2.*k2_4.*k3_4 + k0_2.*k1_3.*k2_3.*k3_4 + k0_2.*k1_3.*k2_4.*k3_4 + k0_1.*k1_3.*k2_0.*k4_2 + k0_1.*k1_3.*k2_0.*k4_3 + k0_1.*k1_3.*k2_1.*k4_2 + k0_2.*k1_0.*k2_3.*k4_2 + k0_1.*k1_2.*k2_3.*k4_2 + k0_1.*k1_3.*k2_1.*k4_3 + k0_2.*k1_0.*k2_3.*k4_3 + k0_2.*k1_3.*k2_1.*k4_2 + k0_1.*k1_2.*k2_3.*k4_3 + k0_1.*k1_3.*k2_3.*k4_2 + k0_2.*k1_0.*k2_4.*k4_3 + k0_2.*k1_2.*k2_3.*k4_2 + k0_2.*k1_3.*k2_1.*k4_3 + k0_1.*k1_2.*k2_4.*k4_3 + k0_1.*k1_3.*k2_3.*k4_3 + k0_2.*k1_2.*k2_3.*k4_3 + k0_2.*k1_3.*k2_3.*k4_2 + k0_1.*k1_3.*k2_4.*k4_3 + k0_2.*k1_2.*k2_4.*k4_3 + k0_2.*k1_3.*k2_3.*k4_3 + k0_2.*k1_3.*k2_4.*k4_3 + k0_2.*k1_0.*k3_1.*k4_2 + k0_1.*k1_2.*k3_1.*k4_2 + k0_2.*k1_0.*k3_1.*k4_3 + k0_2.*k1_0.*k3_2.*k4_2 + k0_1.*k1_2.*k3_1.*k4_3 + k0_1.*k1_2.*k3_2.*k4_2 + k0_2.*k1_0.*k3_2.*k4_3 + k0_2.*k1_2.*k3_1.*k4_2 + k0_1.*k1_2.*k3_2.*k4_3 + k0_1.*k1_3.*k3_2.*k4_2 + k0_2.*k1_0.*k3_4.*k4_2 + k0_2.*k1_2.*k3_1.*k4_3 + k0_2.*k1_2.*k3_2.*k4_2 + k0_1.*k1_2.*k3_4.*k4_2 + k0_1.*k1_3.*k3_2.*k4_3 + k0_2.*k1_2.*k3_2.*k4_3 + k0_2.*k1_3.*k3_2.*k4_2 + k0_1.*k1_3.*k3_4.*k4_2 + k0_2.*k1_2.*k3_4.*k4_2 + k0_2.*k1_3.*k3_2.*k4_3 + k0_2.*k1_3.*k3_4.*k4_2 + k0_1.*k2_0.*k3_1.*k4_2 + k0_1.*k2_0.*k3_1.*k4_3 + k0_1.*k2_0.*k3_2.*k4_2 + k0_1.*k2_1.*k3_1.*k4_2 + k0_1.*k2_0.*k3_2.*k4_3 + k0_1.*k2_1.*k3_1.*k4_3 + k0_1.*k2_1.*k3_2.*k4_2 + k0_2.*k2_1.*k3_1.*k4_2 + k0_1.*k2_0.*k3_4.*k4_2 + k0_1.*k2_1.*k3_2.*k4_3 + k0_1.*k2_3.*k3_1.*k4_2 + k0_2.*k2_1.*k3_1.*k4_3 + k0_2.*k2_1.*k3_2.*k4_2 + k0_1.*k2_1.*k3_4.*k4_2 + k0_1.*k2_3.*k3_1.*k4_3 + k0_2.*k2_1.*k3_2.*k4_3 + k0_2.*k2_3.*k3_1.*k4_2 + k0_1.*k2_4.*k3_1.*k4_3 + k0_2.*k2_1.*k3_4.*k4_2 + k0_2.*k2_3.*k3_1.*k4_3 + k0_2.*k2_4.*k3_1.*k4_3 + k1_0.*k2_0.*k3_1.*k4_2 + k1_0.*k2_0.*k3_1.*k4_3 + k1_0.*k2_0.*k3_2.*k4_2 + k1_0.*k2_1.*k3_1.*k4_2 + k1_0.*k2_0.*k3_2.*k4_3 + k1_0.*k2_1.*k3_1.*k4_3 + k1_0.*k2_1.*k3_2.*k4_2 + k1_2.*k2_0.*k3_1.*k4_2 + k1_0.*k2_0.*k3_4.*k4_2 + k1_0.*k2_1.*k3_2.*k4_3 + k1_0.*k2_3.*k3_1.*k4_2 + k1_2.*k2_0.*k3_1.*k4_3 + k1_2.*k2_0.*k3_2.*k4_2 + k1_0.*k2_1.*k3_4.*k4_2 + k1_0.*k2_3.*k3_1.*k4_3 + k1_2.*k2_0.*k3_2.*k4_3 + k1_3.*k2_0.*k3_2.*k4_2 + k1_0.*k2_4.*k3_1.*k4_3 + k1_2.*k2_0.*k3_4.*k4_2 + k1_3.*k2_0.*k3_2.*k4_3 + k1_3.*k2_0.*k3_4.*k4_2);
S2=(k0_2.*k1_0.*k3_1.*k4_2 + k0_1.*k1_2.*k3_1.*k4_2 + k0_2.*k1_0.*k3_1.*k4_3 + k0_2.*k1_0.*k3_2.*k4_2 + k0_1.*k1_2.*k3_1.*k4_3 + k0_1.*k1_2.*k3_2.*k4_2 + k0_2.*k1_0.*k3_2.*k4_3 + k0_2.*k1_2.*k3_1.*k4_2 + k0_1.*k1_2.*k3_2.*k4_3 + k0_1.*k1_3.*k3_2.*k4_2 + k0_2.*k1_0.*k3_4.*k4_2 + k0_2.*k1_2.*k3_1.*k4_3 + k0_2.*k1_2.*k3_2.*k4_2 + k0_1.*k1_2.*k3_4.*k4_2 + k0_1.*k1_3.*k3_2.*k4_3 + k0_2.*k1_2.*k3_2.*k4_3 + k0_2.*k1_3.*k3_2.*k4_2 + k0_1.*k1_3.*k3_4.*k4_2 + k0_2.*k1_2.*k3_4.*k4_2 + k0_2.*k1_3.*k3_2.*k4_3 + k0_2.*k1_3.*k3_4.*k4_2)./(k0_2.*k1_0.*k2_4.*k3_1 + k0_1.*k1_2.*k2_4.*k3_1 + k0_1.*k1_3.*k2_0.*k3_4 + k0_2.*k1_0.*k2_4.*k3_2 + k0_1.*k1_2.*k2_4.*k3_2 + k0_1.*k1_3.*k2_1.*k3_4 + k0_2.*k1_0.*k2_3.*k3_4 + k0_2.*k1_2.*k2_4.*k3_1 + k0_1.*k1_2.*k2_3.*k3_4 + k0_1.*k1_3.*k2_4.*k3_2 + k0_2.*k1_0.*k2_4.*k3_4 + k0_2.*k1_2.*k2_4.*k3_2 + k0_2.*k1_3.*k2_1.*k3_4 + k0_1.*k1_2.*k2_4.*k3_4 + k0_1.*k1_3.*k2_3.*k3_4 + k0_2.*k1_2.*k2_3.*k3_4 + k0_2.*k1_3.*k2_4.*k3_2 + k0_1.*k1_3.*k2_4.*k3_4 + k0_2.*k1_2.*k2_4.*k3_4 + k0_2.*k1_3.*k2_3.*k3_4 + k0_2.*k1_3.*k2_4.*k3_4 + k0_1.*k1_3.*k2_0.*k4_2 + k0_1.*k1_3.*k2_0.*k4_3 + k0_1.*k1_3.*k2_1.*k4_2 + k0_2.*k1_0.*k2_3.*k4_2 + k0_1.*k1_2.*k2_3.*k4_2 + k0_1.*k1_3.*k2_1.*k4_3 + k0_2.*k1_0.*k2_3.*k4_3 + k0_2.*k1_3.*k2_1.*k4_2 + k0_1.*k1_2.*k2_3.*k4_3 + k0_1.*k1_3.*k2_3.*k4_2 + k0_2.*k1_0.*k2_4.*k4_3 + k0_2.*k1_2.*k2_3.*k4_2 + k0_2.*k1_3.*k2_1.*k4_3 + k0_1.*k1_2.*k2_4.*k4_3 + k0_1.*k1_3.*k2_3.*k4_3 + k0_2.*k1_2.*k2_3.*k4_3 + k0_2.*k1_3.*k2_3.*k4_2 + k0_1.*k1_3.*k2_4.*k4_3 + k0_2.*k1_2.*k2_4.*k4_3 + k0_2.*k1_3.*k2_3.*k4_3 + k0_2.*k1_3.*k2_4.*k4_3 + k0_2.*k1_0.*k3_1.*k4_2 + k0_1.*k1_2.*k3_1.*k4_2 + k0_2.*k1_0.*k3_1.*k4_3 + k0_2.*k1_0.*k3_2.*k4_2 + k0_1.*k1_2.*k3_1.*k4_3 + k0_1.*k1_2.*k3_2.*k4_2 + k0_2.*k1_0.*k3_2.*k4_3 + k0_2.*k1_2.*k3_1.*k4_2 + k0_1.*k1_2.*k3_2.*k4_3 + k0_1.*k1_3.*k3_2.*k4_2 + k0_2.*k1_0.*k3_4.*k4_2 + k0_2.*k1_2.*k3_1.*k4_3 + k0_2.*k1_2.*k3_2.*k4_2 + k0_1.*k1_2.*k3_4.*k4_2 + k0_1.*k1_3.*k3_2.*k4_3 + k0_2.*k1_2.*k3_2.*k4_3 + k0_2.*k1_3.*k3_2.*k4_2 + k0_1.*k1_3.*k3_4.*k4_2 + k0_2.*k1_2.*k3_4.*k4_2 + k0_2.*k1_3.*k3_2.*k4_3 + k0_2.*k1_3.*k3_4.*k4_2 + k0_1.*k2_0.*k3_1.*k4_2 + k0_1.*k2_0.*k3_1.*k4_3 + k0_1.*k2_0.*k3_2.*k4_2 + k0_1.*k2_1.*k3_1.*k4_2 + k0_1.*k2_0.*k3_2.*k4_3 + k0_1.*k2_1.*k3_1.*k4_3 + k0_1.*k2_1.*k3_2.*k4_2 + k0_2.*k2_1.*k3_1.*k4_2 + k0_1.*k2_0.*k3_4.*k4_2 + k0_1.*k2_1.*k3_2.*k4_3 + k0_1.*k2_3.*k3_1.*k4_2 + k0_2.*k2_1.*k3_1.*k4_3 + k0_2.*k2_1.*k3_2.*k4_2 + k0_1.*k2_1.*k3_4.*k4_2 + k0_1.*k2_3.*k3_1.*k4_3 + k0_2.*k2_1.*k3_2.*k4_3 + k0_2.*k2_3.*k3_1.*k4_2 + k0_1.*k2_4.*k3_1.*k4_3 + k0_2.*k2_1.*k3_4.*k4_2 + k0_2.*k2_3.*k3_1.*k4_3 + k0_2.*k2_4.*k3_1.*k4_3 + k1_0.*k2_0.*k3_1.*k4_2 + k1_0.*k2_0.*k3_1.*k4_3 + k1_0.*k2_0.*k3_2.*k4_2 + k1_0.*k2_1.*k3_1.*k4_2 + k1_0.*k2_0.*k3_2.*k4_3 + k1_0.*k2_1.*k3_1.*k4_3 + k1_0.*k2_1.*k3_2.*k4_2 + k1_2.*k2_0.*k3_1.*k4_2 + k1_0.*k2_0.*k3_4.*k4_2 + k1_0.*k2_1.*k3_2.*k4_3 + k1_0.*k2_3.*k3_1.*k4_2 + k1_2.*k2_0.*k3_1.*k4_3 + k1_2.*k2_0.*k3_2.*k4_2 + k1_0.*k2_1.*k3_4.*k4_2 + k1_0.*k2_3.*k3_1.*k4_3 + k1_2.*k2_0.*k3_2.*k4_3 + k1_3.*k2_0.*k3_2.*k4_2 + k1_0.*k2_4.*k3_1.*k4_3 + k1_2.*k2_0.*k3_4.*k4_2 + k1_3.*k2_0.*k3_2.*k4_3 + k1_3.*k2_0.*k3_4.*k4_2);
S3=(k0_1.*k1_3.*k2_0.*k4_2 + k0_1.*k1_3.*k2_0.*k4_3 + k0_1.*k1_3.*k2_1.*k4_2 + k0_2.*k1_0.*k2_3.*k4_2 + k0_1.*k1_2.*k2_3.*k4_2 + k0_1.*k1_3.*k2_1.*k4_3 + k0_2.*k1_0.*k2_3.*k4_3 + k0_2.*k1_3.*k2_1.*k4_2 + k0_1.*k1_2.*k2_3.*k4_3 + k0_1.*k1_3.*k2_3.*k4_2 + k0_2.*k1_0.*k2_4.*k4_3 + k0_2.*k1_2.*k2_3.*k4_2 + k0_2.*k1_3.*k2_1.*k4_3 + k0_1.*k1_2.*k2_4.*k4_3 + k0_1.*k1_3.*k2_3.*k4_3 + k0_2.*k1_2.*k2_3.*k4_3 + k0_2.*k1_3.*k2_3.*k4_2 + k0_1.*k1_3.*k2_4.*k4_3 + k0_2.*k1_2.*k2_4.*k4_3 + k0_2.*k1_3.*k2_3.*k4_3 + k0_2.*k1_3.*k2_4.*k4_3)./(k0_2.*k1_0.*k2_4.*k3_1 + k0_1.*k1_2.*k2_4.*k3_1 + k0_1.*k1_3.*k2_0.*k3_4 + k0_2.*k1_0.*k2_4.*k3_2 + k0_1.*k1_2.*k2_4.*k3_2 + k0_1.*k1_3.*k2_1.*k3_4 + k0_2.*k1_0.*k2_3.*k3_4 + k0_2.*k1_2.*k2_4.*k3_1 + k0_1.*k1_2.*k2_3.*k3_4 + k0_1.*k1_3.*k2_4.*k3_2 + k0_2.*k1_0.*k2_4.*k3_4 + k0_2.*k1_2.*k2_4.*k3_2 + k0_2.*k1_3.*k2_1.*k3_4 + k0_1.*k1_2.*k2_4.*k3_4 + k0_1.*k1_3.*k2_3.*k3_4 + k0_2.*k1_2.*k2_3.*k3_4 + k0_2.*k1_3.*k2_4.*k3_2 + k0_1.*k1_3.*k2_4.*k3_4 + k0_2.*k1_2.*k2_4.*k3_4 + k0_2.*k1_3.*k2_3.*k3_4 + k0_2.*k1_3.*k2_4.*k3_4 + k0_1.*k1_3.*k2_0.*k4_2 + k0_1.*k1_3.*k2_0.*k4_3 + k0_1.*k1_3.*k2_1.*k4_2 + k0_2.*k1_0.*k2_3.*k4_2 + k0_1.*k1_2.*k2_3.*k4_2 + k0_1.*k1_3.*k2_1.*k4_3 + k0_2.*k1_0.*k2_3.*k4_3 + k0_2.*k1_3.*k2_1.*k4_2 + k0_1.*k1_2.*k2_3.*k4_3 + k0_1.*k1_3.*k2_3.*k4_2 + k0_2.*k1_0.*k2_4.*k4_3 + k0_2.*k1_2.*k2_3.*k4_2 + k0_2.*k1_3.*k2_1.*k4_3 + k0_1.*k1_2.*k2_4.*k4_3 + k0_1.*k1_3.*k2_3.*k4_3 + k0_2.*k1_2.*k2_3.*k4_3 + k0_2.*k1_3.*k2_3.*k4_2 + k0_1.*k1_3.*k2_4.*k4_3 + k0_2.*k1_2.*k2_4.*k4_3 + k0_2.*k1_3.*k2_3.*k4_3 + k0_2.*k1_3.*k2_4.*k4_3 + k0_2.*k1_0.*k3_1.*k4_2 + k0_1.*k1_2.*k3_1.*k4_2 + k0_2.*k1_0.*k3_1.*k4_3 + k0_2.*k1_0.*k3_2.*k4_2 + k0_1.*k1_2.*k3_1.*k4_3 + k0_1.*k1_2.*k3_2.*k4_2 + k0_2.*k1_0.*k3_2.*k4_3 + k0_2.*k1_2.*k3_1.*k4_2 + k0_1.*k1_2.*k3_2.*k4_3 + k0_1.*k1_3.*k3_2.*k4_2 + k0_2.*k1_0.*k3_4.*k4_2 + k0_2.*k1_2.*k3_1.*k4_3 + k0_2.*k1_2.*k3_2.*k4_2 + k0_1.*k1_2.*k3_4.*k4_2 + k0_1.*k1_3.*k3_2.*k4_3 + k0_2.*k1_2.*k3_2.*k4_3 + k0_2.*k1_3.*k3_2.*k4_2 + k0_1.*k1_3.*k3_4.*k4_2 + k0_2.*k1_2.*k3_4.*k4_2 + k0_2.*k1_3.*k3_2.*k4_3 + k0_2.*k1_3.*k3_4.*k4_2 + k0_1.*k2_0.*k3_1.*k4_2 + k0_1.*k2_0.*k3_1.*k4_3 + k0_1.*k2_0.*k3_2.*k4_2 + k0_1.*k2_1.*k3_1.*k4_2 + k0_1.*k2_0.*k3_2.*k4_3 + k0_1.*k2_1.*k3_1.*k4_3 + k0_1.*k2_1.*k3_2.*k4_2 + k0_2.*k2_1.*k3_1.*k4_2 + k0_1.*k2_0.*k3_4.*k4_2 + k0_1.*k2_1.*k3_2.*k4_3 + k0_1.*k2_3.*k3_1.*k4_2 + k0_2.*k2_1.*k3_1.*k4_3 + k0_2.*k2_1.*k3_2.*k4_2 + k0_1.*k2_1.*k3_4.*k4_2 + k0_1.*k2_3.*k3_1.*k4_3 + k0_2.*k2_1.*k3_2.*k4_3 + k0_2.*k2_3.*k3_1.*k4_2 + k0_1.*k2_4.*k3_1.*k4_3 + k0_2.*k2_1.*k3_4.*k4_2 + k0_2.*k2_3.*k3_1.*k4_3 + k0_2.*k2_4.*k3_1.*k4_3 + k1_0.*k2_0.*k3_1.*k4_2 + k1_0.*k2_0.*k3_1.*k4_3 + k1_0.*k2_0.*k3_2.*k4_2 + k1_0.*k2_1.*k3_1.*k4_2 + k1_0.*k2_0.*k3_2.*k4_3 + k1_0.*k2_1.*k3_1.*k4_3 + k1_0.*k2_1.*k3_2.*k4_2 + k1_2.*k2_0.*k3_1.*k4_2 + k1_0.*k2_0.*k3_4.*k4_2 + k1_0.*k2_1.*k3_2.*k4_3 + k1_0.*k2_3.*k3_1.*k4_2 + k1_2.*k2_0.*k3_1.*k4_3 + k1_2.*k2_0.*k3_2.*k4_2 + k1_0.*k2_1.*k3_4.*k4_2 + k1_0.*k2_3.*k3_1.*k4_3 + k1_2.*k2_0.*k3_2.*k4_3 + k1_3.*k2_0.*k3_2.*k4_2 + k1_0.*k2_4.*k3_1.*k4_3 + k1_2.*k2_0.*k3_4.*k4_2 + k1_3.*k2_0.*k3_2.*k4_3 + k1_3.*k2_0.*k3_4.*k4_2);
% S4=(k0_2.*k1_0.*k2_4.*k3_1 + k0_1.*k1_2.*k2_4.*k3_1 + k0_1.*k1_3.*k2_0.*k3_4 + k0_2.*k1_0.*k2_4.*k3_2 + k0_1.*k1_2.*k2_4.*k3_2 + k0_1.*k1_3.*k2_1.*k3_4 + k0_2.*k1_0.*k2_3.*k3_4 + k0_2.*k1_2.*k2_4.*k3_1 + k0_1.*k1_2.*k2_3.*k3_4 + k0_1.*k1_3.*k2_4.*k3_2 + k0_2.*k1_0.*k2_4.*k3_4 + k0_2.*k1_2.*k2_4.*k3_2 + k0_2.*k1_3.*k2_1.*k3_4 + k0_1.*k1_2.*k2_4.*k3_4 + k0_1.*k1_3.*k2_3.*k3_4 + k0_2.*k1_2.*k2_3.*k3_4 + k0_2.*k1_3.*k2_4.*k3_2 + k0_1.*k1_3.*k2_4.*k3_4 + k0_2.*k1_2.*k2_4.*k3_4 + k0_2.*k1_3.*k2_3.*k3_4 + k0_2.*k1_3.*k2_4.*k3_4)./(k0_2.*k1_0.*k2_4.*k3_1 + k0_1.*k1_2.*k2_4.*k3_1 + k0_1.*k1_3.*k2_0.*k3_4 + k0_2.*k1_0.*k2_4.*k3_2 + k0_1.*k1_2.*k2_4.*k3_2 + k0_1.*k1_3.*k2_1.*k3_4 + k0_2.*k1_0.*k2_3.*k3_4 + k0_2.*k1_2.*k2_4.*k3_1 + k0_1.*k1_2.*k2_3.*k3_4 + k0_1.*k1_3.*k2_4.*k3_2 + k0_2.*k1_0.*k2_4.*k3_4 + k0_2.*k1_2.*k2_4.*k3_2 + k0_2.*k1_3.*k2_1.*k3_4 + k0_1.*k1_2.*k2_4.*k3_4 + k0_1.*k1_3.*k2_3.*k3_4 + k0_2.*k1_2.*k2_3.*k3_4 + k0_2.*k1_3.*k2_4.*k3_2 + k0_1.*k1_3.*k2_4.*k3_4 + k0_2.*k1_2.*k2_4.*k3_4 + k0_2.*k1_3.*k2_3.*k3_4 + k0_2.*k1_3.*k2_4.*k3_4 + k0_1.*k1_3.*k2_0.*k4_2 + k0_1.*k1_3.*k2_0.*k4_3 + k0_1.*k1_3.*k2_1.*k4_2 + k0_2.*k1_0.*k2_3.*k4_2 + k0_1.*k1_2.*k2_3.*k4_2 + k0_1.*k1_3.*k2_1.*k4_3 + k0_2.*k1_0.*k2_3.*k4_3 + k0_2.*k1_3.*k2_1.*k4_2 + k0_1.*k1_2.*k2_3.*k4_3 + k0_1.*k1_3.*k2_3.*k4_2 + k0_2.*k1_0.*k2_4.*k4_3 + k0_2.*k1_2.*k2_3.*k4_2 + k0_2.*k1_3.*k2_1.*k4_3 + k0_1.*k1_2.*k2_4.*k4_3 + k0_1.*k1_3.*k2_3.*k4_3 + k0_2.*k1_2.*k2_3.*k4_3 + k0_2.*k1_3.*k2_3.*k4_2 + k0_1.*k1_3.*k2_4.*k4_3 + k0_2.*k1_2.*k2_4.*k4_3 + k0_2.*k1_3.*k2_3.*k4_3 + k0_2.*k1_3.*k2_4.*k4_3 + k0_2.*k1_0.*k3_1.*k4_2 + k0_1.*k1_2.*k3_1.*k4_2 + k0_2.*k1_0.*k3_1.*k4_3 + k0_2.*k1_0.*k3_2.*k4_2 + k0_1.*k1_2.*k3_1.*k4_3 + k0_1.*k1_2.*k3_2.*k4_2 + k0_2.*k1_0.*k3_2.*k4_3 + k0_2.*k1_2.*k3_1.*k4_2 + k0_1.*k1_2.*k3_2.*k4_3 + k0_1.*k1_3.*k3_2.*k4_2 + k0_2.*k1_0.*k3_4.*k4_2 + k0_2.*k1_2.*k3_1.*k4_3 + k0_2.*k1_2.*k3_2.*k4_2 + k0_1.*k1_2.*k3_4.*k4_2 + k0_1.*k1_3.*k3_2.*k4_3 + k0_2.*k1_2.*k3_2.*k4_3 + k0_2.*k1_3.*k3_2.*k4_2 + k0_1.*k1_3.*k3_4.*k4_2 + k0_2.*k1_2.*k3_4.*k4_2 + k0_2.*k1_3.*k3_2.*k4_3 + k0_2.*k1_3.*k3_4.*k4_2 + k0_1.*k2_0.*k3_1.*k4_2 + k0_1.*k2_0.*k3_1.*k4_3 + k0_1.*k2_0.*k3_2.*k4_2 + k0_1.*k2_1.*k3_1.*k4_2 + k0_1.*k2_0.*k3_2.*k4_3 + k0_1.*k2_1.*k3_1.*k4_3 + k0_1.*k2_1.*k3_2.*k4_2 + k0_2.*k2_1.*k3_1.*k4_2 + k0_1.*k2_0.*k3_4.*k4_2 + k0_1.*k2_1.*k3_2.*k4_3 + k0_1.*k2_3.*k3_1.*k4_2 + k0_2.*k2_1.*k3_1.*k4_3 + k0_2.*k2_1.*k3_2.*k4_2 + k0_1.*k2_1.*k3_4.*k4_2 + k0_1.*k2_3.*k3_1.*k4_3 + k0_2.*k2_1.*k3_2.*k4_3 + k0_2.*k2_3.*k3_1.*k4_2 + k0_1.*k2_4.*k3_1.*k4_3 + k0_2.*k2_1.*k3_4.*k4_2 + k0_2.*k2_3.*k3_1.*k4_3 + k0_2.*k2_4.*k3_1.*k4_3 + k1_0.*k2_0.*k3_1.*k4_2 + k1_0.*k2_0.*k3_1.*k4_3 + k1_0.*k2_0.*k3_2.*k4_2 + k1_0.*k2_1.*k3_1.*k4_2 + k1_0.*k2_0.*k3_2.*k4_3 + k1_0.*k2_1.*k3_1.*k4_3 + k1_0.*k2_1.*k3_2.*k4_2 + k1_2.*k2_0.*k3_1.*k4_2 + k1_0.*k2_0.*k3_4.*k4_2 + k1_0.*k2_1.*k3_2.*k4_3 + k1_0.*k2_3.*k3_1.*k4_2 + k1_2.*k2_0.*k3_1.*k4_3 + k1_2.*k2_0.*k3_2.*k4_2 + k1_0.*k2_1.*k3_4.*k4_2 + k1_0.*k2_3.*k3_1.*k4_3 + k1_2.*k2_0.*k3_2.*k4_3 + k1_3.*k2_0.*k3_2.*k4_2 + k1_0.*k2_4.*k3_1.*k4_3 + k1_2.*k2_0.*k3_4.*k4_2 + k1_3.*k2_0.*k3_2.*k4_3 + k1_3.*k2_0.*k3_4.*k4_2);
S4 = 1 - S0 - S1 - S2 - S3;

%% Turnover Rate
JNADH = p.CI.x.*(kfNADH_02.*S0 - krNADH_02.*S2 ...
    + kfNADH_24.*S2 - krNADH_24.*S4 ...
    + kfNADH_13.*S1 - krNADH_13.*S3); % NADH oxidation releases 1 proton in matrix space
JSO = p.CI.x.*(S1.*(kfSO_10 - krSO_21) + S2.*(kfSO_21 - krSO_32) + S3.*(kfSO_32 - krSO_43) + S4.*kfSO_43 - krSO_10.*S0);
JH2O2 = p.CI.x.*(S2.*(kfH2O2_20 - krH2O2_42) + S3.*kfH2O2_31 + S4.*kfH2O2_42 - krH2O2_31.*S1 - krH2O2_20.*S0);
% JQ = p.CI.x.*(kfQ_20.*S2 - krQ_20.*S0 ...
%     + kfQ_42.*S4 - krQ_42.*S2 ...
%     + kfQ_31.*S3 - krQ_31.*S1); % charge transfer and proton pumping is the same rate as JQ, 4 charges and protons moved to IMS

% for numerical stability, enforce strict stoichiometric coupling
JQ = JNADH - JSO/2 - JH2O2;

%% Output

if nargout > 1
    % pack rate constants, state fractional occupancies and dGs into g
    % NADH oxidation
    g.JNADH = JNADH;
    % Q reduction
    g.JQ = JQ;
    % ROS rates
    g.JSO = JSO;
    g.JH2O2 = JH2O2;
    g.JROS = JSO/2 + JH2O2;
    % Etot
    g.Etot = p.CI.x;
    % superoxide partial reactions
    g.JSQSO = p.CI.x.*O2.*(p.CI.kfSQSO_10.*s1.F_N2_SQ.*S1 + (p.CI.kfSQSO_21.*s2.Frad_N2_SQ + p.CI.kfSQSO_21b.*s2.F_N2r_SQ).*S2 + (p.CI.kfSQSO_32.*s3.Fr_N2_SQ + p.CI.kfSQSO_32b.*s3.Frad_N2r_SQ).*S3 + p.CI.kfSQSO_43.*s4.Fr_N2r_SQ.*S4);
    g.JFrSO = p.CI.x.*O2.*(p.CI.kfFrSO_21.*s2.Fr_N2.*S2 + (p.CI.kfFrSO_32.*s3.Fr_N2r + p.CI.kfFrSO_32b.*s3.Fr_N2_SQ).*S3 + p.CI.kfFrSO_43.*s4.Fr_N2r_SQ.*S4);
    g.JFradSO = p.CI.x.*O2.*(p.CI.kfFradSO_10.*s1.Frad_N2.*S1 + (p.CI.kfFradSO_21.*s2.Frad_N2r + p.CI.kfFradSO_21b.*s2.Frad_N2_SQ).*S2 + p.CI.kfFradSO_32.*s3.Frad_N2r_SQ.*S3);
    g.JN2rSO = p.CI.x.*O2.*(p.CI.kfN2rSO_10.*s1.F_N2r.*S1 + (p.CI.kfN2rSO_21.*s2.Frad_N2r + p.CI.kfN2rSO_21b.*s2.F_N2r_SQ).*S2 + (p.CI.kfN2rSO_32.*s3.Fr_N2r + p.CI.kfN2rSO_32b.*s3.Frad_N2r_SQ).*S3 + p.CI.kfN2rSO_43.*s4.Fr_N2r_SQ.*S4);
    g.JFrH2O2 = p.CI.x.*O2.*(p.CI.kfH_20.*s2.Fr_N2.*S2 + (p.CI.kfH_31.*s3.Fr_N2r + p.CI.kfH_31b.*s3.Fr_N2_SQ).*S3 + p.CI.kfH_42.*s4.Fr_N2r_SQ.*S4);
    % half reaction TNs
    g.Jn1 = kfNADH_02.*S0 - krNADH_02.*S2;
    g.Jn2 = kfNADH_24.*S2 - krNADH_24.*S4;
    g.Jn3 = kfNADH_13.*S1 - krNADH_13.*S3;
    g.Jq1 = kfQ_20.*S2 - krQ_20.*S0;
    g.Jq2 = kfQ_42.*S4 - krQ_42.*S2;
    g.Jq3 = kfQ_31.*S3 - krQ_31.*S1;
    g.JSO1 = S1.*kfSO_10;
    g.JSO2 =  S2.*kfSO_21;
    g.JSO3 =  S3.*kfSO_32;
    g.JSO4 = S4.*kfSO_43;
    g.JH2O22 = S2.*kfH2O2_20;
    g.JH2O23 = S3.*kfH2O2_31;
    g.JH2O24 = S4.*kfH2O2_42;
    % branch rates
    g.kfNADH_02 = kfNADH_02;
    g.krNADH_02 = krNADH_02;
    g.kfQ_20 = kfQ_20;
    g.krQ_20 = krQ_20;
    g.kfNADH_13 = kfNADH_13;
    g.krNADH_13 = krNADH_13;
    g.kfQ_31 = kfQ_31;
    g.krQ_31 = krQ_31;
    g.kfNADH_24 = kfNADH_24;
    g.krNADH_24 = krNADH_24;
    g.kfQ_42 = kfQ_42;
    g.krQ_42 = krQ_42;
    g.kSO10 = kfSO_10;
    g.kSO21 = kfSO_21;
    g.kSO32 = kfSO_32;
    g.kSO43 = kfSO_43;
    g.kH20 = kfH2O2_20;
    g.kH31 = kfH2O2_31;
    g.kH42 = kfH2O2_42;
%     % SQ only ROS
%     g.JSQ1 = (p.kfSQSO_10*s1.F_N2_SQ).*S1;
%     g.JSQ2 = (p.kfSQSO_21*(s2.Frad_N2_SQ + s2.F_N2r_SQ)).*S2;
%     g.JSQ3 = (p.kfSQSO_32*(s3.Fr_N2_SQ + s3.Frad_N2r_SQ)).*S3;
%     g.JSQ4 = ((p.kfSQSO_43).*s4.Fr_N2r_SQ).*S4;
    % parameters
    g.p = p;
    % midpoint potentials
    g.Em = Em;
    % Keqs
    g.Keq = Keq;
    % BPs
    g.PNo=PNo;
    g.PNr=PNr;
    g.PNrad=PNrad;
    g.PQo=PQo;
    g.PQr=PQr;
    % states
    g.S0=S0;
    g.S1=S1;
    g.S2=S2;
    g.S3=S3;
    g.S4=S4;
    % substates
    g.s0=s0;
    g.s1=s1;
    g.s2=s2;
    g.s3=s3;
    g.s4=s4;
    
    g.K1 = K1;
    g.K2 = K2;
    g.KSQ = KSQ;
    g.KQ = KQ;
    
    varargout = {JNADH,JQ,JSO,JH2O2,g};
else
    varargout = {JNADH};
end
end

%% This version is used to the simulate the S/Rot experiments
% Complex II
% free radical ROS generation
% 4./27./2020

function varargout = CII(x,p)

n = size(x,2); % Need this for substate array construction

%% Solution of the matrix 
O2            = x(p.i.O2_x,:);            
SO            = x(p.i.SO_x,:);            
H2O2          = x(p.i.H2O2_x,:);           
Q             = x(p.i.UQ_x,:);                
QH2           = x(p.i.UQH2_x,:); 
H             = x(p.i.H_x,:);
Succinate     = x(p.i.SUC_x,:);      
Fumarate      = x(p.i.FUM_x,:);       
Malonate      = 0;%x(p.i.Malonate,:);
Oxaloacetate  = x(p.i.OAA_x,:);
GEA           = 0;%x(p.i.GEA_x,:);
Malate        = x(p.i.MAL_x,:);

GED = 0; % no data set has any non-zero general electron donor concentrations

%% Proton dependency

xH = 1./(1+H./p.CII.KD_H);

%% Binding Polynomials for enzyme, substrates, products and regulators

BP_Q = (1 + Q./p.CII.KD_Q + QH2./p.CII.KD_QH2 + p.CII.atpenin./p.CII.KD_atpenin);

p.CII.kf_QH2 = p.CII.kf_QH2.*(1 + p.CII.atpenin./p.CII.KD_atpenin2)./(1 + p.CII.beta_atpenin*p.CII.atpenin./p.CII.KD_atpenin2);
p.CII.kf_SUC = p.CII.kf_SUC.*(1 + p.CII.atpenin./p.CII.KD_atpenin2)./(1 + p.CII.beta_atpenin*p.CII.atpenin./p.CII.KD_atpenin2).*xH;
 
BP_FAD = (1 + Succinate./p.CII.KD_SUC + Fumarate./p.CII.KD_FUM + Malate./p.CII.KD_MAL + Malonate./p.CII.KD_MALO  + Oxaloacetate./p.CII.KD_OXA); 

%% inactivation dynamics

I = Oxaloacetate./p.CII.KD_OXA./BP_FAD;

Xinact = p.CII.Ki_OAA./(I + p.CII.Ki_OAA); % assuming steady-state mechanism
p.CII.kf_SUC = p.CII.kf_SUC.*Xinact;

%% FMN Occupancy for SO and H2O2 reactions

p.CII.kf_FADH = p.CII.kf_FADH./BP_FAD./(1+H./p.CII.K_FADH);
p.CII.kf_FADH2 = p.CII.kf_FADH2./BP_FAD./(1+H./p.CII.K_FADH2);
p.CII.kf_H2O2 = p.CII.kf_H2O2./BP_FAD./(1+H./p.CII.K_FADH2);

%% Adjust redox midpoint potentials

% FAD potential as a pH dependence
Em.FAD_FADH =  p.CII.Em0.FAD_FADH + (p.R.*p.T./p.F).*log(H.*(1+p.CII.K_FADH./H)); 
Em.FADH_FADH2 = p.CII.Em0.FADH_FADH2 + (p.R.*p.T./p.F) *log(H.*(1+H./p.CII.K_FADH2)./(1+H./p.CII.K_FADH));
Em.FAD_FADH2 = p.CII.Em0.FAD_FADH2 + (p.R.*p.T./p.F./2)*log(H.^2.*(1+H./p.CII.K_FADH2).*(1+p.CII.K_FADH./H)./(1+H./p.CII.K_FADH));

Em.Q_SQ   =   p.CII.Em0.Q_SQ.*ones(1,n); % pH independent
Em.SQ_QH2 =   2*p.CII.Em0.Q_QH2 + 2*(p.R*p.T./p.F).*log(H) - Em.Q_SQ; % Em of SQ./QH2 couple at given pH

Em.Q_QH2 = p.CII.Em0.Q_QH2 + (p.R*p.T./p.F).*log(H);
Em.Fum_Suc = p.CII.Em0.Fum_Suc + (p.R*p.T./p.F).*log(H);

% general electron acceptor (phenazine or TMPD in studies used)
p.CII.Em.GEA = p.CII.Em0.GEA  + (p.R*p.T/2/p.F) * log(H);

% Midpoint potentials for electron transfer of bound states
Emb.SQ_QH2 = Em.SQ_QH2 - (p.R.*p.T./p.F).*log(p.CII.KD_QH2); % electron transfer free energy (mV)
Emb.Q_SQ = Em.Q_SQ + (p.R.*p.T./p.F).*log(p.CII.KD_Q); % electron transfer free energy (mV)
Emb.Fum_Suc = Em.Fum_Suc - (p.R.*p.T./p.F/2).*log(p.CII.KD_SUC) + (p.R.*p.T./p.F/2).*log(p.CII.KD_FUM); % electron transfer free energy (mV)

% Equilibrium constants, Keq
Keq.SO_ISC3             = exp(p.F.*(p.CII.Em.O2_SO  - p.CII.Em.ISC3)./(p.R*p.T));
Keq.SO_SQ               = exp(p.F.*(p.CII.Em.O2_SO  - Emb.Q_SQ)./(p.R*p.T));
Keq.SO_FADH             = exp(p.F.*(p.CII.Em.O2_SO - Em.FAD_FADH)./(p.R*p.T));
Keq.SO_FADH2            = exp(p.F.*(p.CII.Em.O2_SO - Em.FADH_FADH2)./(p.R*p.T));
Keq.H2O2_FADH2          = exp(2*p.F.*(p.CII.Em.O2_H2O2 - Em.FAD_FADH2)./(p.R*p.T));
Keq.QH2_ISC3            = exp(p.F.*(Emb.SQ_QH2  - p.CII.Em.ISC3)./(p.R*p.T)); % should be bound state Em for SQ_QH2
Keq.FUM_FADH2           = exp(2*p.F.*(Em.FAD_FADH2 - Emb.Fum_Suc)./(p.R*p.T)); % should be Em.FAD - Emb.Suc
Keq.GEA_ISC            = exp(2*p.F.*(p.CII.Em.GEA - (p.CII.Em.ISC3 + p.CII.Em.ISC1))./( p.R*p.T)); % since Em of ISC2 so low, using ISC1 as second e- source

%% Boltzman redox poise potentials
% compute Eh for Q./SQ reaction
Eh.FAD_FADH = Em.FAD_FADH;
Eh.FADH_FADH2 = Em.FADH_FADH2;
Eh.ISC1 = p.CII.Em.ISC1*ones(1,n);
Eh.ISC2 = p.CII.Em.ISC2*ones(1,n);
Eh.ISC3 = p.CII.Em.ISC3*ones(1,n);    
Eh.Q_SQ = Emb.Q_SQ + (p.R.*p.T./p.F).*log(Q./p.CII.KD_Q./BP_Q);

%% RT normalized DeltaG's of all substrates
% FAD_FADH, FADH_FADH2,FAD_FADH2,ISC1,ISC2,ISC3,Q_SQ;
% Eh.FAD_FADH2
DfG1 = -p.F.* [Eh.FAD_FADH;...
    Eh.FADH_FADH2;...
    Eh.ISC1;...
    Eh.ISC2;...
    Eh.ISC3;...
    Eh.Q_SQ]./p.R./p.T;

%% The fractional substrates of redox centers for each elecrtonic states
% E0
s0 = 1;
% E1
E1i = DfG1(p.CII.substates.s1,:);
s1 = exp(-E1i)./(ones(size(p.CII.substates.s1,1),1)*sum(exp(-E1i),1));
% E2
E2i = squeeze(sum(reshape(DfG1([p.CII.substates.s2],:),size(p.CII.substates.s2,1),size(p.CII.substates.s2,2),n),2));          
s2 = exp(-E2i)./(ones(size(p.CII.substates.s2,1),1)*sum(exp(-E2i),1));
% E3
E3i = squeeze(sum(reshape(DfG1([p.CII.substates.s3],:),size(p.CII.substates.s3,1),size(p.CII.substates.s3,2),n),2)); 
s3 = exp(-E3i)./(ones(size(p.CII.substates.s3,1),1)*sum(exp(-E3i),1));
% E4
E4i=squeeze(sum(reshape(DfG1([p.CII.substates.s4],:),size(p.CII.substates.s4,1),size(p.CII.substates.s4,2),n),2)); 
s4=exp(-E4i)./(ones(size(p.CII.substates.s4,1),1)*sum(exp(-E4i),1));

%% State Transition Rates
s1_FADH  = sum(s1(p.CII.s1.FADH,:),1);
s1_FAD     = sum(s1(p.CII.s1.FADox,:),1);
s1_SQ        = sum(s1(p.CII.s1.SQ,:),1);
s1_SQempty   = sum(s1(p.CII.s1.SQempty,:),1);
s1_SQempty_ISC3ox   = sum(s1(p.CII.s1.SQempty_ISC3ox,:),1);
% s1_ISC1ox_ISC3ox = sum(s1(p.CII.s1.ISC1ox_ISC3ox,:),1);
s1_ISC3     = sum(s1(p.CII.s1.ISC3,:),1);
s1_ISC3ox   = sum(s1(p.CII.s1.ISC3ox,:),1);

s2_FAD    = sum(s2(p.CII.s2.FADox,:),1);
s2_FADH     = sum(s2(p.CII.s2.FADH,:),1);
s2_FADH2    = sum(s2(p.CII.s2.FADH2,:),1);
s2_SQ       = sum(s2(p.CII.s2.SQ,:),1);
s2_SQempty  = sum(s2(p.CII.s2.SQempty,:),1);
s2_ISC3     = sum(s2(p.CII.s2.ISC3_SQempty,:),1);
s2_ISC3ox   = sum(s2(p.CII.s2.ISC3ox,:),1);
% s2_ISC1_ISC3 = sum(s2(p.CII.s2.ISC1_ISC3,:),1); 
s2_SQ_ISC3  = sum(s2(p.CII.s2.SQ_ISC3,:),1);
s2_SQempty_ISC3ox  = sum(s2(p.CII.s2.SQempty_ISC3ox,:),1);
% s2_ISC1ox_ISC3ox = sum(s2(p.CII.s2.ISC1ox_ISC3ox,:),1);

s3_FAD    = sum(s3(p.CII.s3.FADox,:),1);
s3_FADH     = sum(s3(p.CII.s3.FADH,:),1);
s3_FADH2    = sum(s3(p.CII.s3.FADH2,:),1);
s3_SQ       = sum(s3(p.CII.s3.SQ,:),1);
s3_SQempty  = sum(s3(p.CII.s3.SQempty,:),1);
s3_ISC3     = sum(s3(p.CII.s3.ISC3_SQempty,:),1);
s3_ISC3ox   = sum(s3(p.CII.s3.ISC3ox,:),1);
% s3_ISC1_ISC3 = sum(s3(p.CII.s3.ISC1_ISC3,:),1);
s3_SQ_ISC3  = sum(s3(p.CII.s3.SQ_ISC3,:),1);
% s3_ISC1ox_ISC3ox = sum(s3(p.CII.s3.ISC1ox_ISC3ox,:),1);
% s3_SQempty_ISC3ox = sum(s3(p.CII.s2.SQempty_ISC3ox,:),1);

%s4_FAD    = sum(s4(p.CII.s4.FADox,:),1);
s4_FADH     = sum(s4(p.CII.s4.FADH,:),1);
s4_FADH2    = sum(s4(p.CII.s4.FADH2,:),1);
s4_SQ       = sum(s4(p.CII.s4.SQ,:),1);
% s4_SQempty  = sum(s4(p.CII.s4.SQempty,:),1);
s4_ISC3     = sum(s4(p.CII.s4.ISC3_SQempty,:),1);
% s4_ISC3ox   = sum(s4(p.CII.s4.ISC3ox,:),1);
% s4_ISC1_ISC3 = sum(s4(p.CII.s4.ISC1_ISC3,:),1);
s4_SQ_ISC3  = sum(s4(p.CII.s4.SQ_ISC3,:),1);
% s4_ISC1ox_ISC3ox = sum(s4(p.CII.s4.ISC1ox_ISC3ox,:),1);
% s4_SQempty_ISC3ox = sum(s4(p.CII.s4.SQempty_ISC3ox,:),1);

%% Unidirectional state transition rates
% 1) E0->E1
% SO + Q -> O2 + SQ
% FAD + H + SO -> FADH + O2

k01_SO_ISC3 = (p.CII.kf_SO2./Keq.SO_ISC3).* 1./BP_Q .* SO .* s0;% ./ (1 + GEA./p.CII.KD_GEA);
k01_SO_Q = (p.CII.kf_SO./Keq.SO_SQ).* Q./p.CII.KD_Q./BP_Q .* SO .* s0;
k01_SO_FAD = (p.CII.kf_FADH./Keq.SO_FADH).* SO .* s0;
k0_1 =   k01_SO_Q + k01_SO_FAD + k01_SO_ISC3;

% 2) E1->E0
% O2 + SQ -> SO + Q
% FADH + O2 -> FAD + H + SO

k10_O2_ISC3 = p.CII.kf_SO2 .* O2 .* s1_ISC3 ./ BP_Q;% ./ (1 + GEA./p.CII.KD_GEA);
k10_O2_SQ = p.CII.kf_SO .* O2 .* s1_SQ;
k10_O2_FADH = p.CII.kf_FADH .* O2 .* s1_FADH;
k1_0 = k10_O2_SQ  + k10_O2_FADH + k10_O2_ISC3;

% 3) E0->E2                       
% H2O2 + FAD -> O2 + FADH2
% FAD + Succinate -> FADH2 + fumarate
% QH2 + [4Fe_3S]ox -> SQ + [4Fe_3S]red + 2H
% GED + [4Fe_3S]ox + [3Fe_4S]ox -> GEA  + [4Fe_3S]red + [3Fe_4S]red

k02_H2O2_FAD = (p.CII.kf_H2O2./Keq.H2O2_FADH2) .* H2O2 .* s0;
k02_SUC_FAD =  p.CII.kf_SUC.* Succinate./p.CII.KD_SUC./BP_FAD .* s0;
k02_QH2_SQempty_ISC3ox =  (p.CII.kf_QH2./ Keq.QH2_ISC3).* QH2./p.CII.KD_QH2./BP_Q .* s0;
k02_GED = (p.CII.kf_GEA./Keq.GEA_ISC).*GED;%./(GED+p.CII.KD_GEA);%.*s0;
k0_2 = k02_H2O2_FAD + k02_SUC_FAD + k02_QH2_SQempty_ISC3ox +  k02_GED; 

% 4) E2->E0
% O2 + FADH2 ->  H2O2 + FAD
% FADH2 + fumarate -> FAD + Succinate
% SQ-[4Fe_3S]red + 2H -> QH2 + [4Fe_3S]ox
% GEA + [4Fe_3S]red + [3Fe_4S]red = GED + [4Fe_3S]ox + [3Fe_4S]ox

k20_O2_FADH2 = p.CII.kf_H2O2 .* O2 .* s2_FADH2;
k20_FUM_FADH2 = (p.CII.kf_SUC./ Keq.FUM_FADH2).* Fumarate./p.CII.KD_FUM./BP_FAD .* s2_FADH2; 
k20_SQ_ISC3_QH2 =  p.CII.kf_QH2 .* s2_SQ_ISC3;
k20_GEA = p.CII.kf_GEA.*GEA;%./(GEA+p.CII.KD_GEA);%.*s2_ISC1_ISC3;
k2_0 = k20_O2_FADH2 + k20_FUM_FADH2 + k20_SQ_ISC3_QH2 + k20_GEA; 

% 5) E1->E2 
% SO + Q -> O2 + SQ
% SO + FAD + H -> FADH + O2
% SO + FADH + H -> FADH2 + O2

k12_SO_ISC3 = (p.CII.kf_SO2./Keq.SO_ISC3) .* 1./BP_Q .* SO .* s1_SQempty;% ./ (1 + GEA./p.CII.KD_GEA);
k12_SO_Q = (p.CII.kf_SO./Keq.SO_SQ) .* Q./p.CII.KD_Q./BP_Q .* SO .* s1_SQempty;
k12_SO_FAD = (p.CII.kf_FADH./Keq.SO_FADH).* SO .* s1_FAD;
k12_SO_FADH = (p.CII.kf_FADH./Keq.SO_FADH2).* SO .* s1_FADH;
k1_2 =   k12_SO_Q + k12_SO_FAD + k12_SO_FADH + k12_SO_ISC3;

% 6) E2->E1
% O2 + SQ -> SO + Q
% O2 + FADH -> FAD + H + SO
% O2 + FADH2 -> FADH + H+ SO

k21_O2_ISC3 = p.CII.kf_SO2 .* O2 .* s2_ISC3 ./  BP_Q;% ./ (1 + GEA./p.CII.KD_GEA);
k21_O2_SQ = p.CII.kf_SO .* O2 .* s2_SQ;
k21_O2_FADH = p.CII.kf_FADH .* O2 .* s2_FADH;
k21_O2_FADH2 = p.CII.kf_FADH2 .* O2 .* s2_FADH2;
k2_1 = k21_O2_SQ  + k21_O2_FADH + k21_O2_FADH2 + k21_O2_ISC3;

% 7) E1->E3
% H2O2 + FAD -> O2 + FADH2
% FAD + Succinate -> FADH2 + fumarate
% QH2 + [4Fe_3S]ox -> SQ + [4Fe_3S]red + 2H
% GED + [4Fe_3S]ox + [3Fe_4S]ox -> GEA  + [4Fe_3S]red + [3Fe_4S]red

k13_H2O2_FAD = (p.CII.kf_H2O2./Keq.H2O2_FADH2) .* H2O2 .* s1_FAD;
k13_SUC_FAD =  p.CII.kf_SUC.* Succinate./p.CII.KD_SUC./BP_FAD .* s1_FAD; 
k13_QH2_SQempty_ISC3ox =  (p.CII.kf_QH2./ Keq.QH2_ISC3).* QH2./p.CII.KD_QH2./BP_Q .* s1_SQempty_ISC3ox;
k13_GED = (p.CII.kf_GEA./Keq.GEA_ISC).*GED;%./(GED+p.CII.KD_GEA);%.*s1_ISC1ox_ISC3ox; 
k1_3 = k13_H2O2_FAD + k13_SUC_FAD + k13_QH2_SQempty_ISC3ox + k13_GED; 

% 8) E3->E1
% O2 + FADH2 ->  H2O2 + FAD
% FADH2 + fumarate -> FAD + Succinate
% SQ-[4Fe_3S]red + 2H -> QH2 + [4Fe_3S]ox
% GEA + [4Fe_3S]red + [3Fe_4S]red = GED + [4Fe_3S]ox + [3Fe_4S]ox

k31_O2_FADH2 = p.CII.kf_H2O2 .* O2 .* s3_FADH2;
k31_FUM_FADH2 = (p.CII.kf_SUC./ Keq.FUM_FADH2).* Fumarate./p.CII.KD_FUM./BP_FAD .* s3_FADH2;
k31_SQ_ISC3_QH2 =  p.CII.kf_QH2.* s3_SQ_ISC3;
k31_GEA = p.CII.kf_GEA.*GEA;%./(GEA+p.CII.KD_GEA);%.*s3_ISC1_ISC3;
k3_1 = k31_O2_FADH2 + k31_FUM_FADH2 + k31_SQ_ISC3_QH2 + k31_GEA;

% 9) E2->E3
% SO + ISC3ox -> O2 + ISC3red
% SO + Q -> O2 + SQ
% SO + FAD + H -> FADH + O2
% SO + FADH + H -> FADH2 + O2

k23_SO_ISC3 = (p.CII.kf_SO2./Keq.SO_ISC3) .* 1./BP_Q .* SO .* s2_SQempty;% ./ (1 + GEA./p.CII.KD_GEA);
k23_SO_Q = (p.CII.kf_SO./Keq.SO_SQ) .* Q./p.CII.KD_Q./BP_Q .* SO .* s2_SQempty;
k23_SO_FAD = (p.CII.kf_FADH./Keq.SO_FADH).* SO .* s2_FAD;
k23_SO_FADH = (p.CII.kf_FADH./Keq.SO_FADH2).* SO .* s2_FADH;
k2_3 =   k23_SO_Q + k23_SO_FADH + k23_SO_FAD + k23_SO_ISC3;

% 10) E3->E2
% O2 + SQ -> SO + Q
% O2 + FADH -> FAD + H+ + SO
% O2 + FADH2 -> FADH + H + SO

k32_O2_ISC3 = p.CII.kf_SO2 .* O2 .* s3_ISC3 ./ BP_Q;% ./ (1 + GEA./p.CII.KD_GEA);
k32_O2_SQ = p.CII.kf_SO .* O2 .* s3_SQ;
k32_O2_FADH = p.CII.kf_FADH .* O2 .* s3_FADH;
k32_O2_FADH2 = p.CII.kf_FADH2 .* O2 .* s3_FADH2;
k3_2= k32_O2_SQ + k32_O2_FADH + k32_O2_FADH2 + k32_O2_ISC3;

% 11) E2->E4
% H2O2 + FAD -> O2 + FADH2
% FAD + Succinate -> FADH2 + fumarate
% QH2 + [4Fe_3S]ox -> SQ + [4Fe_3S]red + 2H
% GED + [4Fe_3S]ox + [3Fe_4S]ox -> GEA  + [4Fe_3S]red + [3Fe_4S]red

k24_H2O2_FAD = (p.CII.kf_H2O2./Keq.H2O2_FADH2) .* H2O2 .* s2_FAD;
k24_SUC_FAD = p.CII.kf_SUC.* Succinate./p.CII.KD_SUC./BP_FAD .* s2_FAD; 
k24_QH2_SQempty_ISC3ox = (p.CII.kf_QH2./Keq.QH2_ISC3) .* QH2./p.CII.KD_QH2./BP_Q .* s2_SQempty_ISC3ox;
k24_GED = (p.CII.kf_GEA./Keq.GEA_ISC).*GED;%./(GED+p.CII.KD_GEA);%.*s2_ISC1ox_ISC3ox; 
k2_4 = k24_H2O2_FAD + k24_SUC_FAD + k24_QH2_SQempty_ISC3ox + k24_GED;

% 12) E4->E2
% O2 + FADH2 ->  H2O2 + FAD
% FADH2 + fumarate -> FAD + Succinate
% SQ-[4Fe_3S]red + 2H -> QH2 + [4Fe_3S]ox
% GEA + [4Fe_3S]red + [3Fe_4S]red = GED + [4Fe_3S]ox + [3Fe_4S]ox

k42_O2_FADH2 = p.CII.kf_H2O2 .* O2 .* s4_FADH2;
k42_FUM_FADH2 = (p.CII.kf_SUC./ Keq.FUM_FADH2).* Fumarate./p.CII.KD_FUM./BP_FAD .* s4_FADH2; 
k42_SQ_ISC3_QH2 =  p.CII.kf_QH2.* s4_SQ_ISC3;
k42_GEA = p.CII.kf_GEA.*GEA;%./(GEA+p.CII.KD_GEA);%.*s4_ISC1_ISC3; 
k4_2 = k42_O2_FADH2 + k42_FUM_FADH2 + k42_SQ_ISC3_QH2 + k42_GEA;

% 13) E3->E4
% SO + Q -> O2 + SQ
% SO + FAD + H -> FADH + O2
% SO + FADH + H -> FADH2 + O2

k34_SO_ISC3 = (p.CII.kf_SO2./Keq.SO_ISC3) .* 1./BP_Q .* SO .* s3_SQempty;% ./ (1 + GEA./p.CII.KD_GEA); %%
k34_SO_Q = (p.CII.kf_SO./Keq.SO_SQ) .* Q./p.CII.KD_Q./BP_Q .* SO .* s3_SQempty; %%
k34_SO_FAD = (p.CII.kf_FADH./Keq.SO_FADH).* SO .* s3_FAD;
k34_SO_FADH = p.CII.kf_FADH2 .*SO .* s3_FADH;
k3_4 =   k34_SO_Q + k34_SO_FAD + k34_SO_FADH + k34_SO_ISC3;

% 14) E4->E3
% O2 + SQ -> SO + Q
% O2 + FADH -> FAD + H + SO
% O2 + FADH2 -> FADH + H+ SO

k43_O2_ISC3 = p.CII.kf_SO2 .* O2 .* s4_ISC3 ./ BP_Q;% ./ (1 + GEA./p.CII.KD_GEA);
k43_O2_SQ = p.CII.kf_SO .* O2 .* s4_SQ;
k43_O2_FADH = p.CII.kf_FADH .* O2 .* s4_FADH;
k43_O2_FADH2 =p.CII.kf_FADH2 .* O2 .* s4_FADH2;
k4_3 = k43_O2_SQ + k43_O2_FADH + k43_O2_FADH2 + k43_O2_ISC3;

% electron occupancy steady states
D = (k0_2.*k1_0.*k2_4.*k3_1 + k0_1.*k1_2.*k2_4.*k3_1 + k0_1.*k1_3.*k2_0.*k3_4 + k0_2.*k1_0.*k2_4.*k3_2 + k0_1.*k1_2.*k2_4.*k3_2 + k0_1.*k1_3.*k2_1.*k3_4 + k0_2.*k1_0.*k2_3.*k3_4 + k0_2.*k1_2.*k2_4.*k3_1 + k0_1.*k1_2.*k2_3.*k3_4 + k0_1.*k1_3.*k2_4.*k3_2 + k0_2.*k1_0.*k2_4.*k3_4 + k0_2.*k1_2.*k2_4.*k3_2 + k0_2.*k1_3.*k2_1.*k3_4 + k0_1.*k1_2.*k2_4.*k3_4 + k0_1.*k1_3.*k2_3.*k3_4 + k0_2.*k1_2.*k2_3.*k3_4 + k0_2.*k1_3.*k2_4.*k3_2 + k0_1.*k1_3.*k2_4.*k3_4 + k0_2.*k1_2.*k2_4.*k3_4 + k0_2.*k1_3.*k2_3.*k3_4 + k0_2.*k1_3.*k2_4.*k3_4 + k0_1.*k1_3.*k2_0.*k4_2 + k0_1.*k1_3.*k2_0.*k4_3 + k0_1.*k1_3.*k2_1.*k4_2 + k0_2.*k1_0.*k2_3.*k4_2 + k0_1.*k1_2.*k2_3.*k4_2 + k0_1.*k1_3.*k2_1.*k4_3 + k0_2.*k1_0.*k2_3.*k4_3 + k0_2.*k1_3.*k2_1.*k4_2 + k0_1.*k1_2.*k2_3.*k4_3 + k0_1.*k1_3.*k2_3.*k4_2 + k0_2.*k1_0.*k2_4.*k4_3 + k0_2.*k1_2.*k2_3.*k4_2 + k0_2.*k1_3.*k2_1.*k4_3 + k0_1.*k1_2.*k2_4.*k4_3 + k0_1.*k1_3.*k2_3.*k4_3 + k0_2.*k1_2.*k2_3.*k4_3 + k0_2.*k1_3.*k2_3.*k4_2 + k0_1.*k1_3.*k2_4.*k4_3 + k0_2.*k1_2.*k2_4.*k4_3 + k0_2.*k1_3.*k2_3.*k4_3 + k0_2.*k1_3.*k2_4.*k4_3 + k0_2.*k1_0.*k3_1.*k4_2 + k0_1.*k1_2.*k3_1.*k4_2 + k0_2.*k1_0.*k3_1.*k4_3 + k0_2.*k1_0.*k3_2.*k4_2 + k0_1.*k1_2.*k3_1.*k4_3 + k0_1.*k1_2.*k3_2.*k4_2 + k0_2.*k1_0.*k3_2.*k4_3 + k0_2.*k1_2.*k3_1.*k4_2 + k0_1.*k1_2.*k3_2.*k4_3 + k0_1.*k1_3.*k3_2.*k4_2 + k0_2.*k1_0.*k3_4.*k4_2 + k0_2.*k1_2.*k3_1.*k4_3 + k0_2.*k1_2.*k3_2.*k4_2 + k0_1.*k1_2.*k3_4.*k4_2 + k0_1.*k1_3.*k3_2.*k4_3 + k0_2.*k1_2.*k3_2.*k4_3 + k0_2.*k1_3.*k3_2.*k4_2 + k0_1.*k1_3.*k3_4.*k4_2 + k0_2.*k1_2.*k3_4.*k4_2 + k0_2.*k1_3.*k3_2.*k4_3 + k0_2.*k1_3.*k3_4.*k4_2 + k0_1.*k2_0.*k3_1.*k4_2 + k0_1.*k2_0.*k3_1.*k4_3 + k0_1.*k2_0.*k3_2.*k4_2 + k0_1.*k2_1.*k3_1.*k4_2 + k0_1.*k2_0.*k3_2.*k4_3 + k0_1.*k2_1.*k3_1.*k4_3 + k0_1.*k2_1.*k3_2.*k4_2 + k0_2.*k2_1.*k3_1.*k4_2 + k0_1.*k2_0.*k3_4.*k4_2 + k0_1.*k2_1.*k3_2.*k4_3 + k0_1.*k2_3.*k3_1.*k4_2 + k0_2.*k2_1.*k3_1.*k4_3 + k0_2.*k2_1.*k3_2.*k4_2 + k0_1.*k2_1.*k3_4.*k4_2 + k0_1.*k2_3.*k3_1.*k4_3 + k0_2.*k2_1.*k3_2.*k4_3 + k0_2.*k2_3.*k3_1.*k4_2 + k0_1.*k2_4.*k3_1.*k4_3 + k0_2.*k2_1.*k3_4.*k4_2 + k0_2.*k2_3.*k3_1.*k4_3 + k0_2.*k2_4.*k3_1.*k4_3 + k1_0.*k2_0.*k3_1.*k4_2 + k1_0.*k2_0.*k3_1.*k4_3 + k1_0.*k2_0.*k3_2.*k4_2 + k1_0.*k2_1.*k3_1.*k4_2 + k1_0.*k2_0.*k3_2.*k4_3 + k1_0.*k2_1.*k3_1.*k4_3 + k1_0.*k2_1.*k3_2.*k4_2 + k1_2.*k2_0.*k3_1.*k4_2 + k1_0.*k2_0.*k3_4.*k4_2 + k1_0.*k2_1.*k3_2.*k4_3 + k1_0.*k2_3.*k3_1.*k4_2 + k1_2.*k2_0.*k3_1.*k4_3 + k1_2.*k2_0.*k3_2.*k4_2 + k1_0.*k2_1.*k3_4.*k4_2 + k1_0.*k2_3.*k3_1.*k4_3 + k1_2.*k2_0.*k3_2.*k4_3 + k1_3.*k2_0.*k3_2.*k4_2 + k1_0.*k2_4.*k3_1.*k4_3 + k1_2.*k2_0.*k3_4.*k4_2 + k1_3.*k2_0.*k3_2.*k4_3 + k1_3.*k2_0.*k3_4.*k4_2);
E0= (k1_0.*k2_0.*k3_1.*k4_2 + k1_0.*k2_0.*k3_1.*k4_3 + k1_0.*k2_0.*k3_2.*k4_2 + k1_0.*k2_1.*k3_1.*k4_2 + k1_0.*k2_0.*k3_2.*k4_3 + k1_0.*k2_1.*k3_1.*k4_3 + k1_0.*k2_1.*k3_2.*k4_2 + k1_2.*k2_0.*k3_1.*k4_2 + k1_0.*k2_0.*k3_4.*k4_2 + k1_0.*k2_1.*k3_2.*k4_3 + k1_0.*k2_3.*k3_1.*k4_2 + k1_2.*k2_0.*k3_1.*k4_3 + k1_2.*k2_0.*k3_2.*k4_2 + k1_0.*k2_1.*k3_4.*k4_2 + k1_0.*k2_3.*k3_1.*k4_3 + k1_2.*k2_0.*k3_2.*k4_3 + k1_3.*k2_0.*k3_2.*k4_2 + k1_0.*k2_4.*k3_1.*k4_3 + k1_2.*k2_0.*k3_4.*k4_2 + k1_3.*k2_0.*k3_2.*k4_3 + k1_3.*k2_0.*k3_4.*k4_2)./D;
E1= (k0_1.*k2_0.*k3_1.*k4_2 + k0_1.*k2_0.*k3_1.*k4_3 + k0_1.*k2_0.*k3_2.*k4_2 + k0_1.*k2_1.*k3_1.*k4_2 + k0_1.*k2_0.*k3_2.*k4_3 + k0_1.*k2_1.*k3_1.*k4_3 + k0_1.*k2_1.*k3_2.*k4_2 + k0_2.*k2_1.*k3_1.*k4_2 + k0_1.*k2_0.*k3_4.*k4_2 + k0_1.*k2_1.*k3_2.*k4_3 + k0_1.*k2_3.*k3_1.*k4_2 + k0_2.*k2_1.*k3_1.*k4_3 + k0_2.*k2_1.*k3_2.*k4_2 + k0_1.*k2_1.*k3_4.*k4_2 + k0_1.*k2_3.*k3_1.*k4_3 + k0_2.*k2_1.*k3_2.*k4_3 + k0_2.*k2_3.*k3_1.*k4_2 + k0_1.*k2_4.*k3_1.*k4_3 + k0_2.*k2_1.*k3_4.*k4_2 + k0_2.*k2_3.*k3_1.*k4_3 + k0_2.*k2_4.*k3_1.*k4_3)./D;
E2= (k0_2.*k1_0.*k3_1.*k4_2 + k0_1.*k1_2.*k3_1.*k4_2 + k0_2.*k1_0.*k3_1.*k4_3 + k0_2.*k1_0.*k3_2.*k4_2 + k0_1.*k1_2.*k3_1.*k4_3 + k0_1.*k1_2.*k3_2.*k4_2 + k0_2.*k1_0.*k3_2.*k4_3 + k0_2.*k1_2.*k3_1.*k4_2 + k0_1.*k1_2.*k3_2.*k4_3 + k0_1.*k1_3.*k3_2.*k4_2 + k0_2.*k1_0.*k3_4.*k4_2 + k0_2.*k1_2.*k3_1.*k4_3 + k0_2.*k1_2.*k3_2.*k4_2 + k0_1.*k1_2.*k3_4.*k4_2 + k0_1.*k1_3.*k3_2.*k4_3 + k0_2.*k1_2.*k3_2.*k4_3 + k0_2.*k1_3.*k3_2.*k4_2 + k0_1.*k1_3.*k3_4.*k4_2 + k0_2.*k1_2.*k3_4.*k4_2 + k0_2.*k1_3.*k3_2.*k4_3 + k0_2.*k1_3.*k3_4.*k4_2)./D;
E3= (k0_1.*k1_3.*k2_0.*k4_2 + k0_1.*k1_3.*k2_0.*k4_3 + k0_1.*k1_3.*k2_1.*k4_2 + k0_2.*k1_0.*k2_3.*k4_2 + k0_1.*k1_2.*k2_3.*k4_2 + k0_1.*k1_3.*k2_1.*k4_3 + k0_2.*k1_0.*k2_3.*k4_3 + k0_2.*k1_3.*k2_1.*k4_2 + k0_1.*k1_2.*k2_3.*k4_3 + k0_1.*k1_3.*k2_3.*k4_2 + k0_2.*k1_0.*k2_4.*k4_3 + k0_2.*k1_2.*k2_3.*k4_2 + k0_2.*k1_3.*k2_1.*k4_3 + k0_1.*k1_2.*k2_4.*k4_3 + k0_1.*k1_3.*k2_3.*k4_3 + k0_2.*k1_2.*k2_3.*k4_3 + k0_2.*k1_3.*k2_3.*k4_2 + k0_1.*k1_3.*k2_4.*k4_3 + k0_2.*k1_2.*k2_4.*k4_3 + k0_2.*k1_3.*k2_3.*k4_3 + k0_2.*k1_3.*k2_4.*k4_3)./D;
E4= 1 -(E0+E1+E2+E3);

%% Compute steady-state rates

% Superoxide from fully reduced flavin site (rates)
%k21_O2_FADH2 =0; k32_O2_FADH2=0; k43_O2_FADH2 =0;
%k12_SO_FADH =0 ; k23_SO_FADH = 0;  k34_SO_FADH = 0; 

% Superoxide from Q site 
%k10_O2_SQ = 0;  k21_O2_SQ = 0;  k32_O2_SQ = 0; k43_O2_SQ = 0; 
%k01_SO_Q = 0; k12_SO_Q = 0; k23_SO_Q = 0; k34_SO_Q = 0;

% succinate oxidation rate
J = p.CII.Etot.*(k02_SUC_FAD.*E0 + k13_SUC_FAD.*E1 + k24_SUC_FAD.*E2 ...
   - k20_FUM_FADH2.*E2 - k31_FUM_FADH2.*E3 - k42_FUM_FADH2.*E4);

% Superoxide formation
JSO = p.CII.Etot.*((k10_O2_ISC3 + k10_O2_SQ + k10_O2_FADH).*E1 + (k21_O2_ISC3 + k21_O2_SQ + k21_O2_FADH + k21_O2_FADH2).*E2 + (k32_O2_ISC3 + k32_O2_SQ + k32_O2_FADH + k32_O2_FADH2).*E3 + (k43_O2_ISC3 + k43_O2_SQ + k43_O2_FADH + k43_O2_FADH2).*E4...
                  - (k01_SO_ISC3 + k01_SO_Q + k01_SO_FAD).*E0 - (k12_SO_ISC3 + k12_SO_Q + k12_SO_FAD + k12_SO_FADH).*E1 - (k23_SO_ISC3 + k23_SO_Q + k23_SO_FAD + k23_SO_FADH).*E2 - (k34_SO_ISC3 + k34_SO_Q + k34_SO_FAD + k34_SO_FADH).*E3 );
            
% Hydrogen peroxide formation
JH2O2 = p.CII.Etot.*(k20_O2_FADH2.*E2 + k31_O2_FADH2.*E3 + k42_O2_FADH2.*E4 ...
                    - k02_H2O2_FAD.*E0 - k13_H2O2_FAD.*E1 - k24_H2O2_FAD.*E2);
               
%QH2 reduction              
JQH2 = p.CII.Etot.*(k20_SQ_ISC3_QH2.*E2 + k31_SQ_ISC3_QH2.*E3 + k42_SQ_ISC3_QH2.*E4...
                   - k02_QH2_SQempty_ISC3ox.*E0 - k13_QH2_SQempty_ISC3ox.*E1 - k24_QH2_SQempty_ISC3ox.*E2);
% GEA reduction
JGEA = p.CII.Etot .*(k20_GEA.*E2 + k31_GEA.*E3 + k42_GEA.*E4... 
    - k02_GED.*E0 - k13_GED.*E1 - k24_GED.*E2);

%%  the stuff might be interested  
           
JISC3_SO = p.CII.Etot.*(k10_O2_ISC3.*E1 + k21_O2_ISC3.*E2  + k32_O2_ISC3.*E3 + k43_O2_ISC3.*E4);

% SO formation from Q site
%JQSO = p.CII.Etot.*(k10_O2_SQ.*E1 + k21_O2_SQ.*E2  + k32_O2_SQ.*E3 + k43_O2_SQ.*E4...
%- k01_SO_Q.*E0 - k12_SO_Q.*E1 - k23_SO_Q.*E2 - k34_SO_Q.*E3);
JSQ_SO = p.CII.Etot.*(k10_O2_SQ.*E1 + k21_O2_SQ.*E2  + k32_O2_SQ.*E3 + k43_O2_SQ.*E4);

% Superoxide formation from FADH site 
%JFADH_SO = p.CII.Etot.*(k10_O2_FADH.*E1 + k21_O2_FADH.*E2 + k32_O2_FADH .*E3 + k43_O2_FADH.*E4...
%                - k01_SO_FAD.*E0 - k12_SO_FAD .*E1 - k23_SO_FAD.*E2 - k34_SO_FAD.*E3);
JFADH_SO = p.CII.Etot.*(k10_O2_FADH.*E1 + k21_O2_FADH.*E2 + k32_O2_FADH .*E3 + k43_O2_FADH.*E4); 

%Superoxide formation from FADH2 site 
%JFADH2_SO = p.CII.Etot.*((k21_O2_FADH2).*E2 + (k32_O2_FADH2).*E3 + (k43_O2_FADH2).*E4...
%                 - ( k12_SO_FADH.*E1 + k23_SO_FADH.*E2 + k34_SO_FADH.*E3 ));
JFADH2_SO = p.CII.Etot.*((k21_O2_FADH2).*E2 + (k32_O2_FADH2).*E3 + (k43_O2_FADH2).*E4);

% Hydrogen peroxide formation
% JFADH2_H2O2 = p.CII.Etot.*(k20_O2_FADH2.*E2 + k31_O2_FADH2.*E3 + k42_O2_FADH2.*E4 ...
%                     - k02_H2O2_FAD.*E0 - k13_H2O2_FAD.*E1 - k24_H2O2_FAD.*E2);
JFADH2_H2O2 = p.CII.Etot.*(k20_O2_FADH2.*E2 + k31_O2_FADH2.*E3 + k42_O2_FADH2.*E4);
                   

%% Collect model details and package into structure
% midpoint potentials
details.Em=Em;
details.Emb=Emb;
% equilibrium constants
details.Keq=Keq;
% substate occupancies
details.s0=s0;
details.s1=s1;
details.s2=s2;
details.s3=s3;
details.s4=s4;
% substate redox fractions
details.s1_FADH=s1_FADH;
details.s1_FAD=s1_FAD;
details.s1_SQ=s1_SQ;
details.s1_SQempty=s1_SQempty;
details.s1_SQempty_ISC3ox=s1_SQempty_ISC3ox;
details.s1_ISC3=s1_ISC3;
details.s1_ISC3ox=s1_ISC3ox;
details.s2_FAD=s2_FAD;
details.s2_FADH=s2_FADH;
details.s2_FADH2=s2_FADH2;
details.s2_SQ=s2_SQ;
details.s2_SQempty=s2_SQempty;
details.s2_ISC3=s2_ISC3;
details.s2_ISC3ox=s2_ISC3ox;
details.s2_SQ_ISC3=s2_SQ_ISC3;
details.s2_SQempty_ISC3ox=s2_SQempty_ISC3ox;
details.s3_FAD=s3_FAD;
details.s3_FADH=s3_FADH;
details.s3_FADH2=s3_FADH2;
details.s3_SQ=s3_SQ;
details.s3_SQempty=s3_SQempty;
details.s3_ISC3=s3_ISC3;
details.s3_ISC3ox=s3_ISC3ox;
details.s3_SQ_ISC3=s3_SQ_ISC3;
details.s4_FADH=s4_FADH;
details.s4_FADH2=s4_FADH2;
details.s4_SQ=s4_SQ;
details.s4_ISC3=s4_ISC3;
details.s4_SQ_ISC3=s4_SQ_ISC3;
% state occupancies
details.E0=E0;
details.E1=E1;
details.E2=E2;
details.E3=E3;
details.E4=E4;
%total redox state fractions
details.E1_FADH=E1.*s1_FADH;
details.E1_SQ=E1.*s1_SQ;
details.E2_FADH=E2.*s2_FADH;
details.E2_FADH2=E2.*s2_FADH2;
details.E2_SQ=E2.*s2_SQ;
details.E3_FADH=E3.*s3_FADH;
details.E3_FADH2=E3.*s3_FADH2;
details.E3_SQ=E3.*s3_SQ;
details.E4_FADH=E4.*s4_FADH;
details.E4_FADH2=E4.*s4_FADH2;
details.E4_SQ=E4.*s4_SQ;
%total redox fractions
details.FADH=E1.*s1_FADH + E2.*s2_FADH + E3.*s3_FADH + E4.*s4_FADH;
details.SQ=E1.*s1_SQ + E2.*s2_SQ + E3.*s3_SQ + E4.*s4_SQ;
details.FADH2=E2.*s2_FADH2 + E3.*s3_FADH2 + E4.*s4_FADH2;
% turnover at each site
details.J=J;
details.JSO=JSO;
details.JH2O2=JH2O2;
details.JQH2=JQH2;
details.JGEA=JGEA;
% Partial reactions 
% 1)E0->E1
details.k01_SO_Q = k01_SO_Q;
details.k01_SO_FAD = k01_SO_FAD;
details.k0_1 =k0_1;
% 2)E1->E0
details.k10_O2_SQ = k10_O2_SQ;
details.k10_O2_FADH = k10_O2_FADH;
details.k1_0 = k1_0;
% 3)E0->E2
details.k02_H2O2_FAD = k02_H2O2_FAD;
details.k02_SUC_FAD = k02_SUC_FAD ;
details.k02_QH2_SQempty_ISC3ox = k02_QH2_SQempty_ISC3ox;
details.k02_GED = k02_GED; 
details.k0_2 = k0_2; 
% 4)E2->E0
details.k20_O2_FADH2 = k20_O2_FADH2;
details.k20_FUM_FADH2 = k20_FUM_FADH2; 
details.k20_SQ_ISC3_QH2 =  k20_SQ_ISC3_QH2; 
details.k20_GED = k20_GEA; 
details.k2_0 = k2_0; 
% 5) E1->E2 
details.k12_SO_Q = k12_SO_Q;
details.k12_SO_FAD = k12_SO_FAD;
details.k12_SO_FADH = k12_SO_FADH;
details.k1_2 =   k1_2;
% 6)E2->E1
details.k21_O2_SQ = k21_O2_SQ;
details.k21_O2_FADH = k21_O2_FADH;
details.k21_O2_FADH2 = k21_O2_FADH2;
details.k2_1 = k2_1;
% 7)E1->E3
details.k13_H2O2_FAD = k13_H2O2_FAD;
details.k13_SUC_FAD =  k13_SUC_FAD; 
details.k13_QH2_SQempty_ISC3ox =  k13_QH2_SQempty_ISC3ox;
details.k13_GED = k13_GED; 
details.k1_3 = k1_3; 
% 8)E3->E1
details.k31_O2_FADH2 = k31_O2_FADH2;
details.k31_FUM_FADH2 = k31_FUM_FADH2;
details.k31_SQ_ISC3_QH2 =  k31_SQ_ISC3_QH2;
details.k31_GED = k31_GEA;
details.k3_1 = k3_1;
% 9)E2->E3
details.k23_SO_Q = k23_SO_Q;
details.k23_SO_FAD = k23_SO_FAD;
details.k23_SO_FADH = k23_SO_FADH;
details.k2_3 =   k2_3;
% 10)E3->E2
details.k32_O2_SQ = k32_O2_SQ;
details.k32_O2_FADH = k32_O2_FADH;
details.k32_O2_FADH2 = k32_O2_FADH2;
details.k3_2= k3_2;
% 11)E2->E4
details.k24_H2O2_FAD = k24_H2O2_FAD;
details.k24_SUC_FAD = k24_SUC_FAD; 
details.k24_QH2_SQempty_ISC3ox = k24_QH2_SQempty_ISC3ox;
details.k24_GED = k24_GED;
details.k2_4 = k2_4;
% 12)E4->E2
details.k42_O2_FADH2 = k42_O2_FADH2;
details.k42_FUM_FADH2 = k42_FUM_FADH2; 
details.k42_SQ_ISC3_QH2 =  k42_SQ_ISC3_QH2;
details.k42_GED = k42_GEA;
details.k4_2 = k4_2;
% 13)E3->E4
details.k34_SO_Q = k34_SO_Q;
details.k34_SO_FAD = k34_SO_FAD ;
details.k34_SO_FADH =k34_SO_FADH;
details.k3_4 =   k3_4;
% 14)E4->E3
details.k43_O2_SQ = k43_O2_SQ;
details.k43_O2_FADH = k43_O2_FADH;
details.k43_O2_FADH2 = k43_O2_FADH2;
details.k4_3 = k4_3;

details.JISC3_SO=JISC3_SO;
details.JSQ_SO=JSQ_SO;
details.JFADH_SO=JFADH_SO;
details.JFADH2_SO=JFADH2_SO;
details.JFADH2_H2O2=JFADH2_H2O2;
details.x = x;


%% output
if nargout > 1
    varargout = {J, JSO, JH2O2, JQH2, JGEA, Xinact, details};
else
    varargout = {J};
end

end

%% bc1 Dimer Random Model
% from Bazil et al. 2017

function varargout = CIII(x,p)
% define state variables

dPsi = x(p.i.dPsi,:); % mV
Hp = x(p.i.H_c,:); % M
Hn = x(p.i.H_x,:); % M
Q = x(p.i.UQ_x,:); % M
QH2 = x(p.i.UQH2_x,:); % M
c3 = x(p.i.c3_x,:); % M
c2 = x(p.i.c2_x,:); % M
O2 = x(p.i.O2_x,:); % M
SO = (x(p.i.SO_x,:)+x(p.i.SO_c,:))/2;

n=size(x,2); % need this for substate array construction

p.CIII.Etot=p.CIII.Etot/2; % convert to dimer conc.

% Binding Polynomials for enzyme, substrates, products and regulators

% Qp-site
P_Qp = (1 + Q./p.CIII.KQp + QH2./p.CIII.KQH2p);
% Cytc c-site
P_c = (1 + c2./p.CIII.Kc2 + c3./p.CIII.Kc3);
% Qn-site
P_Qn = (1 + Q./p.CIII.KQn + QH2./p.CIII.KQH2n);
if p.CIII.AA>0 && p.CIII.AAstim==0
    P_Qn = inf; % prevent Q or QH2 binding to Qn-site
end

fAA01 = (1+p.CIII.AA/p.CIII.KAA)/(1+p.CIII.beta_AA01*p.CIII.AA/p.CIII.KAA);
fAA12 = (1+p.CIII.AA/p.CIII.KAA)/(1+p.CIII.beta_AA12*p.CIII.AA/p.CIII.KAA);
fAA23 = (1+p.CIII.AA/p.CIII.KAA)/(1+p.CIII.beta_AA23*p.CIII.AA/p.CIII.KAA);
fAA34 = (1+p.CIII.AA/p.CIII.KAA)/(1+p.CIII.beta_AA34*p.CIII.AA/p.CIII.KAA);
fAA45 = (1+p.CIII.AA/p.CIII.KAA)/(1+p.CIII.beta_AA45*p.CIII.AA/p.CIII.KAA);

if p.CIII.AA>0 && p.CIII.AAstim==0
    P_Qn = inf; % prevent Q or QH2 binding to Qn-site
    p.CIII.kfQH2p_01 = p.CIII.kfQH2p_01*fAA01;
    p.CIII.kfQH2p_12 = p.CIII.kfQH2p_12*fAA12;
    p.CIII.kfQH2p_23 = p.CIII.kfQH2p_23*fAA23;
    p.CIII.kfQH2p_34 = p.CIII.kfQH2p_34*fAA34;
    p.CIII.kfQH2p_45 = p.CIII.kfQH2p_45*fAA45;
end

% Update midpoint potentials from thermodynamic data
p.CIII.Em0.Q_QH2 = (p.dGf(p.if.UQH2) - p.dGf(p.if.UQ))/(-2*p.F);
p.CIII.Em0.c = (p.dGf(p.if.c2) - p.dGf(p.if.c3))/(-1*p.F);
p.CIII.Em.SO = (p.dGf(p.if.SO) - p.dGf(p.if.O2))/(-1*p.F); % pH independent in physiological range

% Midpoint potentials

% SQ potentials
p.CIII.Em0.Q_SQp = p.CIII.Em0.Q_QH2 + p.R*p.T/p.F/2*log(10)*log10(p.CIII.Kstabo) - p.R*p.T/p.F*log(10)*7; % mV
p.CIII.Em0.SQ_QH2p = p.CIII.Em0.Q_QH2 - p.R*p.T/p.F/2*log(10)*log10(p.CIII.Kstabo) + p.R*p.T/p.F*log(10)*7; % mV
p.CIII.Em0.Q_SQn = p.CIII.Em0.Q_QH2 + p.R*p.T/p.F/2*log(10)*log10(p.CIII.Kstabi) - p.R*p.T/p.F*log(10)*7; % mV
p.CIII.Em0.SQ_QH2n = p.CIII.Em0.Q_QH2 - p.R*p.T/p.F/2*log(10)*log10(p.CIII.Kstabi) + p.R*p.T/p.F*log(10)*7; % mV

% Qp-site
Em.c = p.CIII.Em0.c; % pH independent
Em.Q_QH2p = p.CIII.Em0.Q_QH2 + p.R*p.T/p.F*log(Hp);
Em.Q_SQp = p.CIII.Em0.Q_SQp.*ones(1,n); % pH independent
Em.SQ_QH2p = p.CIII.Em0.SQ_QH2p + 2*p.R*p.T/p.F*log(Hp);
Em.bL = p.CIII.Em0.bL - log(10).*p.R.*p.T./p.F.*log10((Hp+10.^-p.CIII.pK.bLox)./(Hp+10.^-p.CIII.pK.bLred)); % assuming linked to P-side, equiv to log10((Hp./10.^-p.CIII.pK.bLred./P_bLred)./(Hp./10.^-p.CIII.pK.bLox./P_bLox))

% Qn-site
Em.bH = p.CIII.Em0.bH - log(10).*p.R.*p.T./p.F.*log10((Hn+10.^-p.CIII.pK.bHox)./(Hn+10.^-p.CIII.pK.bHred)); % assuming linked to N-side, equiv to log10((Hn./10.^-p.CIII.pK.bHred./P_bHred)./(Hn./10.^-p.CIII.pK.bHox./P_bHox))
Em.Q_QH2n = p.CIII.Em0.Q_QH2 + p.R*p.T/p.F*log(Hn);
Em.Q_SQn = p.CIII.Em0.Q_SQn.*ones(1,n); % pH independent
Em.SQ_QH2n = p.CIII.Em0.SQ_QH2n + 2*p.R*p.T/p.F*log(Hn);

% Midpoint potentials for electron transfer of bound states
% dG = dG_ET + dG_binding - dG_unbinding
% dG_binding = RTln(Kd) or -RTln(Ka) where Kd is dissociation constant and Ka is association constant (1./Kd)
Emb.c = Em.c + p.R.*p.T./p.F.*log(p.CIII.Kc3) - p.R.*p.T./p.F.*log(p.CIII.Kc2); % electron transfer free energy (mV)
Emb.SQ_QH2p = Em.SQ_QH2p - p.R.*p.T./p.F.*log(p.CIII.KQH2p); % electron transfer free energy (mV)
Emb.Q_SQp = Em.Q_SQp + p.R.*p.T./p.F.*log(p.CIII.KQp); % electron transfer free energy (mV)
Emb.SQ_QH2n = Em.SQ_QH2n - p.R.*p.T./p.F.*log(p.CIII.KQH2n); % electron transfer free energy (mV)
Emb.Q_SQn = Em.Q_SQn + p.R.*p.T./p.F.*log(p.CIII.KQn); % electron transfer free energy (mV)

%% Equilibrium constants for electron transfer

% QH2 oxidation at Qp-site
Keqb.QH2c2 = exp(p.F.*(Emb.c - Emb.SQ_QH2p)./p.R./p.T); % [c2+][SQp]./[c3+][QH2] Keq of electron transfer of bound-state

% Q reduction at Qn-site
Keqb.bHQH2 = exp(p.F.*(Emb.SQ_QH2n - Em.bH)./p.R./p.T); % [bHox][QH2]./[bHred][SQn] Keq of electron transfer of bound-state

% superoxide at Qp-site
Keq.SQO2 = exp((p.F.*(p.CIII.Em.SO - Emb.Q_SQp))./p.R./p.T); % [Q][O2.-]./[SQp][O2] Keq for overall rxn

% Substates via Rapid Equilibrium

% monomer 1
Eh.Q_SQna = Emb.Q_SQn + p.R.*p.T./p.F.*log(Q./p.CIII.KQn./P_Qn);
Eh.bHa = Em.bH - 2.*(1-p.CIII.beta).*dPsi;
Eh.bLa = Em.bL;
Eh.Q_SQpa = Emb.Q_SQp + p.R.*p.T./p.F.*log(Q./p.CIII.KQp./P_Qp);
% monomer 2
Eh.Q_SQnb = Emb.Q_SQn + p.R.*p.T./p.F.*log(Q./p.CIII.KQn./P_Qn);
Eh.bHb = Em.bH - 2.*(1-p.CIII.beta).*dPsi;
Eh.bLb = Em.bL;
Eh.Q_SQpb = Emb.Q_SQp + p.R.*p.T./p.F.*log(Q./p.CIII.KQp./P_Qp);

% Qp-site inactive unless AA bound to on monomer
Eh.Q_SQpb = -inf*ones(size(Eh.Q_SQpb));

% assumes Qn-site for one monomer is off and other Qp-site is on when AA present
if p.CIII.AAstim == 1
Eh.Q_SQpb = Emb.Q_SQp + p.R.*p.T./p.F.*log(Q./p.CIII.KQp./P_Qp);
Eh.Q_SQnb = -inf*ones(size(Eh.Q_SQnb));
end

% formation energies for each redox center
% index order is given in p.CIII.substates field
dfG1 = -p.F.*[Eh.Q_SQna; Eh.bHa; Eh.bLa; Eh.Q_SQpa; Eh.bLb; Eh.Q_SQpb; Eh.bHb; Eh.Q_SQnb]./p.R./p.T;

% 0 e-
s0 = 1;

% 1 e-
E1i=dfG1(p.CIII.substates.s1,:);
s1=exp(-E1i)./(ones(size(p.CIII.substates.s1,1),1)*sum(exp(-E1i),1));

% 2 e-
E2i=squeeze(sum(reshape(dfG1([p.CIII.substates.s2],:),size(p.CIII.substates.s2,1),size(p.CIII.substates.s2,2),n),2)); % this assumes no interaction between redox centers
% add Coulombic effects for states containing bH(red) and bL(red) on same monomer
E2i(p.CIII.coulomb.s2,:) = E2i(p.CIII.coulomb.s2,:) + p.CIII.dfG_bHredbLred/p.R/p.T;
E2i(p.CIII.coulomb3.s2,:) = E2i(p.CIII.coulomb3.s2,:) + p.CIII.dfG_bLredbLred/p.R/p.T;
s2=exp(-E2i)./(ones(size(p.CIII.substates.s2,1),1)*sum(exp(-E2i),1));

% 3 e-
E3i=squeeze(sum(reshape(dfG1([p.CIII.substates.s3],:),size(p.CIII.substates.s3,1),size(p.CIII.substates.s3,2),n),2)); % this assumes no interaction between redox centers
% add Coulombic effects for states containing bH(red) and bL(red) on same monomer
E3i(p.CIII.coulomb.s3,:) = E3i(p.CIII.coulomb.s3,:) + p.CIII.dfG_bHredbLred/p.R/p.T;
E3i(p.CIII.coulomb3.s3,:) = E3i(p.CIII.coulomb3.s3,:) + p.CIII.dfG_bLredbLred/p.R/p.T;
s3=exp(-E3i)./(ones(size(p.CIII.substates.s3,1),1)*sum(exp(-E3i),1));

% 4 e-
E4i=squeeze(sum(reshape(dfG1([p.CIII.substates.s4],:),size(p.CIII.substates.s4,1),size(p.CIII.substates.s4,2),n),2)); % this assumes no interaction between redox centers
% add Coulombic effects for states containing bH(red) and bL(red) on same monomer
E4i(p.CIII.coulomb.s4,:) = E4i(p.CIII.coulomb.s4,:) + p.CIII.dfG_bHredbLred/p.R/p.T;
E4i(p.CIII.coulomb2.s4,:) = E4i(p.CIII.coulomb2.s4,:) + p.CIII.dfG_bHredbLred/p.R/p.T;
E4i(p.CIII.coulomb3.s4,:) = E4i(p.CIII.coulomb3.s4,:) + p.CIII.dfG_bLredbLred/p.R/p.T;
s4=exp(-E4i)./(ones(size(p.CIII.substates.s4,1),1)*sum(exp(-E4i),1));

% 5 e-
E5i=squeeze(sum(reshape(dfG1([p.CIII.substates.s5],:),size(p.CIII.substates.s5,1),size(p.CIII.substates.s5,2),n),2)); % this assumes no interaction between redox centers
% add Coulombic effects for states containing bH(red) and bL(red) on same monomer
E5i(p.CIII.coulomb.s5,:) = E5i(p.CIII.coulomb.s5,:) + p.CIII.dfG_bHredbLred/p.R/p.T;
E5i(p.CIII.coulomb2.s5,:) = E5i(p.CIII.coulomb2.s5,:) + p.CIII.dfG_bHredbLred/p.R/p.T;
E5i(p.CIII.coulomb3.s5,:) = E5i(p.CIII.coulomb3.s5,:) + p.CIII.dfG_bLredbLred/p.R/p.T;
s5=exp(-E5i)./(ones(size(p.CIII.substates.s5,1),1)*sum(exp(-E5i),1));

% ISP protonation state
PISP = Hp./10.^-p.CIII.pK.ISPox2./(1+Hp./10.^-p.CIII.pK.ISPox1)./(1+Hp./10.^-p.CIII.pK.ISPox2);

% State Transition Rates

% unidirectional state transition rates

% --- S0->S1 ---

% QH2 oxidation at Qp
% QH2 + c3+ + bLox -> Q + bLred + c2+ 2Hp+
kfQH2p_01a = p.CIII.kfQH2p_01.*QH2./p.CIII.KQH2p./P_Qp.*PISP.*c3./p.CIII.Kc3./P_c.*s0; % QH2 oxidation rate on monomer a for S0
kfQH2p_01b = 0;
if p.CIII.AAstim
    kfQH2p_01b = p.CIII.kfQH2p_01.*QH2./p.CIII.KQH2p./P_Qp.*PISP.*c3./p.CIII.Kc3./P_c.*s0.*fAA01; % QH2 oxidation rate on monomer b for S0
end
kfQH2p_01 = kfQH2p_01a + kfQH2p_01b; % total QH2 oxidation rate for S0

% reverse of SO formation
% SO + Q -> SQp + O2
krSO_10a = p.CIII.kfSO./Keq.SQO2.*SO.*Q./p.CIII.KQp./P_Qp.*s0; % reverse rxn on monomer a for S0
krSO_10b = 0;
if p.CIII.AAstim
    krSO_10b = p.CIII.kfSO./Keq.SQO2.*SO.*Q./p.CIII.KQp./P_Qp.*s0.*fAA01; % reverse rxn on monomer b for S0
end
krSO_10 = krSO_10a + krSO_10b; % total reverse SO formation rate for S0

k0_1 = kfQH2p_01 + krSO_10; % total of all rates from S0 to S1

% --- S0<-S1 ---

% reverse of QH2 oxidation at Qp
% Q + bLred + c2+ + 2Hp+ -> QH2 + bLox + c3+
fSQpa1 = sum(s1(p.CIII.s1.Qpa,:),1); % fraction of S1 SQp is reduced on monomer a
fSQpb1 = sum(s1(p.CIII.s1.Qpb,:),1); % fraction of S1 SQp is reduced on monomer b
krQH2p_01a = p.CIII.kfQH2p_01./Keqb.QH2c2.*PISP.*c2./p.CIII.Kc2./P_c.*fSQpa1; % reverse rxn on monomer a for S1
krQH2p_01b = 0;
if p.CIII.AAstim
krQH2p_01b = p.CIII.kfQH2p_01./Keqb.QH2c2.*PISP.*c2./p.CIII.Kc2./P_c.*fSQpb1.*fAA01; % reverse rxn on monomer b for S1
end
krQH2p_01 = krQH2p_01a + krQH2p_01b; % sum rates from both monomers

% SO formation
% SQp + O2 -> SO + Q
kfSO_10a = p.CIII.kfSO.*O2.*fSQpa1; % SO formation rate on monomer a for S1
kfSO_10b = 0;
if p.CIII.AAstim
kfSO_10b = p.CIII.kfSO.*O2.*fSQpb1.*fAA01; % SO formation rate on monomer b for S1
end
kfSO_10 = kfSO_10a + kfSO_10b; % total SO formation rate for S1

k1_0 = krQH2p_01 + kfSO_10; % total of all rates from S1 to S0

% --- S0->S2 ---

% QH2 oxidation at Qn
% QH2 + bHox -> SQ-bHred + 2Hn+
krQH2n_20a = p.CIII.kfQH2n_20./Keqb.bHQH2.*exp(p.CIII.beta.*2.*p.F.*dPsi./p.R./p.T./2).*QH2./p.CIII.KQH2n./P_Qn.*s0; % QH2 oxidation at Qn on monomer a for S0
krQH2n_20b = p.CIII.kfQH2n_20./Keqb.bHQH2.*exp(p.CIII.beta.*2.*p.F.*dPsi./p.R./p.T./2).*QH2./p.CIII.KQH2n./P_Qn.*s0; % QH2 oxidation at Qn on monomer b for S0    ##################### manually turned off for monomer test ####################
if p.CIII.AAstim
    krQH2n_20b = 0;
end
krQH2n_20 = krQH2n_20a + krQH2n_20b; % total QH2 oxidation at Qn

k0_2 = krQH2n_20; % total of all rates from S0 to S2

% --- S0<-S2 ---

% SQ reduction at Qn
% SQn + bHred + 2Hn+ -> QH2 + bHox
fSQnbHreda2 = sum(s2(p.CIII.s2.bHSQna,:),1); % fraction of S2 SQn-bHred on monomer a
fSQnbHredb2 = sum(s2(p.CIII.s2.bHSQnb,:),1); % fraction of S2 SQn-bHred on monomer b
kfQH2n_20a = p.CIII.kfQH2n_20.*exp(-p.CIII.beta.*2.*p.F.*dPsi./p.R./p.T./2).*fSQnbHreda2; % rate of QH2 formation at n on monomer a for S2
kfQH2n_20b = p.CIII.kfQH2n_20.*exp(-p.CIII.beta.*2.*p.F.*dPsi./p.R./p.T./2).*fSQnbHredb2; % rate of QH2 formation at n on monomer b for S2
kfQH2n_20 = kfQH2n_20a + kfQH2n_20b; % total rate of QH2 formation for S2

k2_0 = kfQH2n_20; % total of all rates from S2 to S0

% --- S1->S2 ---

% QH2 oxidation at Qp
% QH2 + c3+ + bLox -> Q + bLred + c2+ 2Hp+
fSQpemptya1 = sum(s1(p.CIII.s1.Qpa_e,:),1); % fraction S1 SQp site empty in monomer a
fSQpemptyb1 = sum(s1(p.CIII.s1.Qpb_e,:),1); % fraction S1 SQp site empty in monomer b
kfQH2p_12a = p.CIII.kfQH2p_12.*QH2./p.CIII.KQH2p./P_Qp.*PISP.*c3./p.CIII.Kc3./P_c.*fSQpemptya1; % QH2 oxidation rate on monomer a for S1
kfQH2p_12b=0;
if p.CIII.AAstim
kfQH2p_12b = p.CIII.kfQH2p_12.*QH2./p.CIII.KQH2p./P_Qp.*PISP.*c3./p.CIII.Kc3./P_c.*fSQpemptyb1.*fAA12; % QH2 oxidation rate on monomer b for S1
end
kfQH2p_12 = kfQH2p_12a + kfQH2p_12b; % total QH2 oxidation rate for S1

% reverse of SO formation
% SO + Q -> SQp + O2
krSO_21a = p.CIII.kfSO./Keq.SQO2.*SO.*Q./p.CIII.KQp./P_Qp.*fSQpemptya1; % reverse rxn on monomer a for S1
krSO_21b=0;
if p.CIII.AAstim
krSO_21b = p.CIII.kfSO./Keq.SQO2.*SO.*Q./p.CIII.KQp./P_Qp.*fSQpemptyb1.*fAA12; % reverse rxn on monomer b for S1    
end
krSO_21 = krSO_21a + krSO_21b; % total reverse SO formation rate for S1

k1_2 = kfQH2p_12 + krSO_21; % total of all rates from S1 to S2

% --- S1<-S2 ---

% reverse of QH2 oxidation at Qp
% Q + bLred + c2+ + 2Hp+ -> QH2 + bLox + c3+   
fSQpa2 = sum(s2(p.CIII.s2.Qpa,:),1); % fraction of S2 SQp on monomer a
fSQpb2 = sum(s2(p.CIII.s2.Qpb,:),1); % fraction of S2 SQp on monomer b
krQH2p_12a = p.CIII.kfQH2p_12./Keqb.QH2c2.*PISP.*c2./p.CIII.Kc2./P_c.*fSQpa2; % reverse rxn on monomer a for S2
krQH2p_12b = 0;
if p.CIII.AAstim
krQH2p_12b = p.CIII.kfQH2p_12./Keqb.QH2c2.*PISP.*c2./p.CIII.Kc2./P_c.*fSQpb2.*fAA12; % reverse rxn on monomer b for S2
end
krQH2p_12 = krQH2p_12a + krQH2p_12b; % sum rates from both monomers

% SO formation
% SQp + O2 -> SO + Q
kfSO_21a = p.CIII.kfSO.*O2.*fSQpa2; % SO formation rate on monomer a for S1
kfSO_21b = 0;
if p.CIII.AAstim
kfSO_21b = p.CIII.kfSO.*O2.*fSQpb2.*fAA12; % SO formation rate on monomer b for S1
end
kfSO_21 = kfSO_21a + kfSO_21b; % total SO formation rate for S1

k2_1 = krQH2p_12 + kfSO_21; % total of all rates from S1 to S0

% --- S2->S3 ---

% QH2 oxidation at Qp
% QH2 + c3+ + bLox -> Q + bLred + c2+ 2Hp+
fSQpemptya2 = sum(s2(p.CIII.s2.Qpa_e,:),1); % fraction S2 SQp site empty on monomer a
fSQpemptyb2 = sum(s2(p.CIII.s2.Qpb_e,:),1); % fraction S2 SQp site empty on monomer b
kfQH2p_23a = p.CIII.kfQH2p_23.*QH2./p.CIII.KQH2p./P_Qp.*PISP.*c3./p.CIII.Kc3./P_c.*fSQpemptya2; % QH2 oxidation rate on monomer a for S1
kfQH2p_23b=0;
if p.CIII.AAstim
kfQH2p_23b = p.CIII.kfQH2p_23.*QH2./p.CIII.KQH2p./P_Qp.*PISP.*c3./p.CIII.Kc3./P_c.*fSQpemptyb2.*fAA23; % QH2 oxidation rate on monomer b for S1
end
kfQH2p_23 = kfQH2p_23a + kfQH2p_23b; % total QH2 oxidation rate for S1

% reverse of SO formation
% SO + Q -> SQp + O2
krSO_32a = p.CIII.kfSO./Keq.SQO2.*SO.*Q./p.CIII.KQp./P_Qp.*fSQpemptya2; % reverse rxn on monomer a for S3
krSO_32b=0;
if p.CIII.AAstim
krSO_32b = p.CIII.kfSO./Keq.SQO2.*SO.*Q./p.CIII.KQp./P_Qp.*fSQpemptyb2.*fAA23; % reverse rxn on monomer b for S3
end
krSO_32 = krSO_32a + krSO_32b; % total reverse SO formation rate for S1

k2_3 = kfQH2p_23 + krSO_32; % total of all rates from S2 to S3

% --- S2<-S3 ---

% reverse of QH2 oxidation at Qp
% Q + bLred + c2+ + 2Hp+ -> QH2 + bLox + c3+
fSQpa3 = sum(s3(p.CIII.s3.Qpa,:),1); % fraction of S3 SQp on monomer a
fSQpb3 = sum(s3(p.CIII.s3.Qpb,:),1); % fraction of S3 SQp on monomer b
krQH2p_23a = p.CIII.kfQH2p_23./Keqb.QH2c2.*PISP.*c2./p.CIII.Kc2./P_c.*fSQpa3; % reverse rxn on monomer a for S3
krQH2p_23b = 0;
if p.CIII.AAstim
krQH2p_23b = p.CIII.kfQH2p_23./Keqb.QH2c2.*PISP.*c2./p.CIII.Kc2./P_c.*fSQpb3.*fAA23; % reverse rxn on monomer b for S3
end
krQH2p_23 = krQH2p_23a + krQH2p_23b; % sum rates from both monomers

% SO formation
% SQp + O2 -> SO + Q
kfSO_32a = p.CIII.kfSO.*O2.*fSQpa3; % SO formation rate on monomer a for S3
kfSO_32b = 0;
if p.CIII.AAstim
kfSO_32b = p.CIII.kfSO.*O2.*fSQpb3.*fAA23; % SO formation rate on monomer b for S3
end
kfSO_32 = kfSO_32a + kfSO_32b; % total SO formation rate for S3

k3_2 = krQH2p_23 + kfSO_32; % total of all rates from S1 to S0

% --- S1<-S3 ---

% SQ reduction at Qn
% SQn + bHred + 2Hn+ -> QH2 + bHox
fSQnbHreda3 = sum(s3(p.CIII.s3.bHSQna,:),1); % fraction of S3 SQn-bHred-any on monomer a
fSQnbHredb3 = sum(s3(p.CIII.s3.bHSQnb,:),1); % fraction of S3 SQn-bHred-any on monomer b
kfQH2n_31a = p.CIII.kfQH2n_31.*exp(-p.CIII.beta.*2.*p.F.*dPsi./p.R./p.T./2).*fSQnbHreda3; % rate of QH2 formation at n on monomer a for S3
kfQH2n_31b = p.CIII.kfQH2n_31.*exp(-p.CIII.beta.*2.*p.F.*dPsi./p.R./p.T./2).*fSQnbHredb3; % rate of QH2 formation at n on monomer b for S3
kfQH2n_31 = kfQH2n_31a + kfQH2n_31b; % total rate of QH2 formation for S3

k3_1 = kfQH2n_31; % total of all rates from S3 to S1

% --- S1->S3 ---

% QH2 oxidation at Qn
% QH2 + bHox -> SQ-bHred + 2Hn+
fQnemptybHoxa1 = sum(s1(p.CIII.s1.bHSQna_e,:),1); % fraction of S1 there is no SQn or bLred on monomer a
fQnemptybHoxb1 = sum(s1(p.CIII.s1.bHSQnb_e,:),1); % fraction of S1 there is no SQn or bLred on monomer b
krQH2n_31a = p.CIII.kfQH2n_31./Keqb.bHQH2.*exp(p.CIII.beta.*2.*p.F.*dPsi./p.R./p.T./2).*QH2./p.CIII.KQH2n./P_Qn.*fQnemptybHoxa1; % QH2 oxidation at Qn on monomer a for S1
krQH2n_31b = p.CIII.kfQH2n_31./Keqb.bHQH2.*exp(p.CIII.beta.*2.*p.F.*dPsi./p.R./p.T./2).*QH2./p.CIII.KQH2n./P_Qn.*fQnemptybHoxb1; % QH2 oxidation at Qn on monomer b for S1    ##################### manually turned off for monomer test ####################
if p.CIII.AAstim
    krQH2n_31b = 0;
end

krQH2n_31 = krQH2n_31a + krQH2n_31b; % total QH2 oxidation at Qn

k1_3 = krQH2n_31; % total of all rates from S0 to S2

% --- S3->S4 ---

% QH2 oxidation at Qp
% QH2 + c3+ + bLox -> Q + bLred + c2+ 2Hp+
fSQpemptya3 = sum(s3(p.CIII.s3.Qpa_e,:),1); % fraction S3 SQp site empty on monomer a
fSQpemptyb3 = sum(s3(p.CIII.s3.Qpb_e,:),1); % fraction S3 SQp site empty on monomer b
kfQH2p_34a = p.CIII.kfQH2p_34.*QH2./p.CIII.KQH2p./P_Qp.*PISP.*c3./p.CIII.Kc3./P_c.*fSQpemptya3; % QH2 oxidation rate on monomer a for S3
kfQH2p_34b=0;
if p.CIII.AAstim
kfQH2p_34b = p.CIII.kfQH2p_34.*QH2./p.CIII.KQH2p./P_Qp.*PISP.*c3./p.CIII.Kc3./P_c.*fSQpemptyb3.*fAA34; % QH2 oxidation rate on monomer b for S3
end
kfQH2p_34 = kfQH2p_34a + kfQH2p_34b; % total QH2 oxidation rate for S1

% reverse of SO formation
% SO + Q -> SQp + O2
krSO_43a = p.CIII.kfSO./Keq.SQO2.*SO.*Q./p.CIII.KQp./P_Qp.*fSQpemptya3; % reverse rxn on monomer a for S3
krSO_43b=0;
if p.CIII.AAstim
krSO_43b = p.CIII.kfSO./Keq.SQO2.*SO.*Q./p.CIII.KQp./P_Qp.*fSQpemptyb3.*fAA34; % reverse rxn on monomer b for S3
end
krSO_43 = krSO_43a + krSO_43b; % total reverse SO formation rate for S1

k3_4 = kfQH2p_34 + krSO_43; % total of all rates from S2 to S3

% --- S3<-S4 ---

% reverse of QH2 oxidation at Qp
% Q + bLred + c2+ + 2Hp+ -> QH2 + bLox + c3+
fSQpa4 = sum(s4(p.CIII.s4.Qpa,:),1); % fraction of S4 SQp on monomer a
fSQpb4 = sum(s4(p.CIII.s4.Qpb,:),1); % fraction of S4 SQp on monomer b
krQH2p_34a = p.CIII.kfQH2p_34./Keqb.QH2c2.*PISP.*c2./p.CIII.Kc2./P_c.*fSQpa4; % reverse rxn on monomer a for S3
krQH2p_34b = 0;
if p.CIII.AAstim
krQH2p_34b = p.CIII.kfQH2p_34./Keqb.QH2c2.*PISP.*c2./p.CIII.Kc2./P_c.*fSQpb4.*fAA34; % reverse rxn on monomer b for S3
end
krQH2p_34 = krQH2p_34a + krQH2p_34b; % sum rates from both monomers

% SO formation
% SQp + O2 -> SO + Q
kfSO_43a = p.CIII.kfSO.*O2.*fSQpa4; % SO formation rate on monomer a for S3
kfSO_43b = 0;
if p.CIII.AAstim
kfSO_43b = p.CIII.kfSO.*O2.*fSQpb4.*fAA34; % SO formation rate on monomer b for S3
end
kfSO_43 = kfSO_43a + kfSO_43b; % total SO formation rate for S3

k4_3 = krQH2p_34 + kfSO_43; % total of all rates from S1 to S0

% --- S2<-S4 ---

% SQ reduction at Qn
% SQn + bHred + 2Hn+ -> QH2 + bHox
fSQnbHreda4 = sum(s4(p.CIII.s4.bHSQna,:),1); % fraction of S4 SQn-bHred-any on monomer a
fSQnbHredb4 = sum(s4(p.CIII.s4.bHSQnb,:),1); % fraction of S4 SQn-bHred-any on monomer b
kfQH2n_42a = p.CIII.kfQH2n_42.*exp(-p.CIII.beta.*2.*p.F.*dPsi./p.R./p.T./2).*fSQnbHreda4; % rate of QH2 formation at n on monomer a for S4
kfQH2n_42b = p.CIII.kfQH2n_42.*exp(-p.CIII.beta.*2.*p.F.*dPsi./p.R./p.T./2).*fSQnbHredb4; % rate of QH2 formation at n on monomer b for S4
kfQH2n_42 = kfQH2n_42a + kfQH2n_42b; % total rate of QH2 formation for S4

k4_2 = kfQH2n_42; % total of all rates from S3 to S1

% --- S2->S4 ---
% QH2 oxidation at Qn
% QH2 + bHox -> SQ-bHred + 2Hn+
fQnemptybHoxa2 = sum(s2(p.CIII.s2.bHSQna_e,:),1); % fraction of S2 there is no SQn or bLred on monomer a
fQnemptybHoxb2 = sum(s2(p.CIII.s2.bHSQnb_e,:),1); % fraction of S2 there is no SQn or bLred on monomer b
krQH2n_42a = p.CIII.kfQH2n_42./Keqb.bHQH2.*exp(p.CIII.beta.*2.*p.F.*dPsi./p.R./p.T./2).*QH2./p.CIII.KQH2n./P_Qn.*fQnemptybHoxa2; % QH2 oxidation at Qn on monomer a for S2
krQH2n_42b = p.CIII.kfQH2n_42./Keqb.bHQH2.*exp(p.CIII.beta.*2.*p.F.*dPsi./p.R./p.T./2).*QH2./p.CIII.KQH2n./P_Qn.*fQnemptybHoxb2; % QH2 oxidation at Qn on monomer b for S2     ##################### manually turned off for monomer test ####################
if p.CIII.AAstim
    krQH2n_42b = 0;
end

krQH2n_42 = krQH2n_42a + krQH2n_42b; % total QH2 oxidation at Qn

k2_4 = krQH2n_42; % total of all rates from S0 to S2

% --- S4->S5 ---

% QH2 oxidation at Qp
% QH2 + c3+ + bLox -> Q + bLred + c2+ 2Hp+
fSQpemptya4 = sum(s4(p.CIII.s4.Qpa_e,:),1); % fraction SQp site empty on monomer a
fSQpemptyb4 = sum(s4(p.CIII.s4.Qpb_e,:),1); % fraction SQp site empty on monomer b
kfQH2p_45a = p.CIII.kfQH2p_45.*QH2./p.CIII.KQH2p./P_Qp.*PISP.*c3./p.CIII.Kc3./P_c.*fSQpemptya4; % QH2 oxidation rate on monomer a
kfQH2p_45b = 0;
if p.CIII.AAstim
kfQH2p_45b = p.CIII.kfQH2p_45.*QH2./p.CIII.KQH2p./P_Qp.*PISP.*c3./p.CIII.Kc3./P_c.*fSQpemptyb4.*fAA45; % QH2 oxidation rate on monomer b
end
kfQH2p_45 = kfQH2p_45a + kfQH2p_45b; % total QH2 oxidation rate for S4

% reverse of SO formation
% SO + Q -> SQp + O2
krSO_54a = p.CIII.kfSO./Keq.SQO2.*SO.*Q./p.CIII.KQp./P_Qp.*fSQpemptya4; % reverse rxn on monomer a
krSO_54b = 0;
if p.CIII.AAstim
krSO_54b = p.CIII.kfSO./Keq.SQO2.*SO.*Q./p.CIII.KQp./P_Qp.*fSQpemptyb4.*fAA45; % reverse rxn on monomer b
end
krSO_54 = krSO_54a + krSO_54b; % total reverse SO formation rate

k4_5 = kfQH2p_45 + krSO_54; % total of all rates from S4 to S5

% --- S4<-S5 ---

% reverse of QH2 oxidation at Qp
% Q + bLred + c2+ + 2Hp+ -> QH2 + bLox + c3+
fSQpa5 = sum(s5(p.CIII.s5.Qpa,:),1); % fraction of SQp on monomer a
fSQpb5 = sum(s5(p.CIII.s5.Qpb,:),1); % fraction of SQp on monomer b
krQH2p_45a = p.CIII.kfQH2p_45./Keqb.QH2c2.*PISP.*c2./p.CIII.Kc2./P_c.*fSQpa5; % reverse rxn on monomer a
krQH2p_45b = 0;
if p.CIII.AAstim
krQH2p_45b = p.CIII.kfQH2p_45./Keqb.QH2c2.*PISP.*c2./p.CIII.Kc2./P_c.*fSQpb5.*fAA45; % reverse rxn on monomer b
end
krQH2p_45 = krQH2p_45a + krQH2p_45b; % sum rates from both monomers

% SO formation
% SQp + O2 -> SO + Q
kfSO_54a = p.CIII.kfSO.*O2.*fSQpa5; % SO formation rate on monomer a
kfSO_54b = 0;
if p.CIII.AAstim
kfSO_54b = p.CIII.kfSO.*O2.*fSQpb5.*fAA45; % SO formation rate on monomer b
end
kfSO_54 = kfSO_54a + kfSO_54b; % total SO formation rate

k5_4 = krQH2p_45 + kfSO_54; % total of all rates from S5 to S4

% --- S3<-S5 ---

% SQ reduction at Qn
% SQn + bHred + 2Hn+ -> QH2 + bHox
fSQnbHreda5 = sum(s5(p.CIII.s5.bHSQna,:),1); % fraction of SQn-bHred-any on monomer a
fSQnbHredb5 = sum(s5(p.CIII.s5.bHSQnb,:),1); % fraction of SQn-bHred-any on monomer b
kfQH2n_53a = p.CIII.kfQH2n_53.*exp(-p.CIII.beta.*2.*p.F.*dPsi./p.R./p.T./2).*fSQnbHreda5; % rate of QH2 formation at n on monomer a
kfQH2n_53b = p.CIII.kfQH2n_53.*exp(-p.CIII.beta.*2.*p.F.*dPsi./p.R./p.T./2).*fSQnbHredb5; % rate of QH2 formation at n on monomer b
kfQH2n_53 = kfQH2n_53a + kfQH2n_53b; % total rate of QH2 formation

k5_3 = kfQH2n_53; % total of all rates from S5 to S3

% --- S3->S5 ---
% QH2 oxidation at Qn
% QH2 + bHox -> SQ-bHred + 2Hn+
fQnemptybHoxa3 = sum(s3(p.CIII.s3.bHSQna_e,:),1); % fraction there is no SQn or bLred on monomer a
fQnemptybHoxb3 = sum(s3(p.CIII.s3.bHSQnb_e,:),1); % fraction there is no SQn or bLred on monomer b
krQH2n_53a = p.CIII.kfQH2n_53./Keqb.bHQH2.*exp(p.CIII.beta.*2.*p.F.*dPsi./p.R./p.T./2).*QH2./p.CIII.KQH2n./P_Qn.*fQnemptybHoxa3; % QH2 oxidation at Qn on monomer a
krQH2n_53b = p.CIII.kfQH2n_53./Keqb.bHQH2.*exp(p.CIII.beta.*2.*p.F.*dPsi./p.R./p.T./2).*QH2./p.CIII.KQH2n./P_Qn.*fQnemptybHoxb3; % QH2 oxidation at Qn on monomer b     ##################### manually turned off for monomer test ####################
if p.CIII.AAstim
    krQH2n_53b=0;
end
krQH2n_53 = krQH2n_53a + krQH2n_53b; % total QH2 oxidation at Qn

k3_5 = krQH2n_53; % total of all rates from S3 to S5

% Steady-State Fractional Occupancies (solved analytically)

D=(k0_1.*k1_3.*k2_0.*k3_5.*k4_2 + k0_1.*k1_3.*k2_0.*k3_5.*k4_3 + k0_1.*k1_3.*k2_1.*k3_5.*k4_2 + k0_2.*k1_0.*k2_3.*k3_5.*k4_2 + k0_2.*k1_0.*k2_4.*k3_1.*k4_5 + k0_1.*k1_2.*k2_3.*k3_5.*k4_2 + k0_1.*k1_2.*k2_4.*k3_1.*k4_5 + k0_1.*k1_3.*k2_0.*k3_4.*k4_5 + k0_1.*k1_3.*k2_1.*k3_5.*k4_3 + k0_2.*k1_0.*k2_3.*k3_5.*k4_3 + k0_2.*k1_0.*k2_4.*k3_2.*k4_5 + k0_2.*k1_3.*k2_1.*k3_5.*k4_2 + k0_1.*k1_2.*k2_3.*k3_5.*k4_3 + k0_1.*k1_2.*k2_4.*k3_2.*k4_5 + k0_1.*k1_3.*k2_0.*k3_5.*k4_5 + k0_1.*k1_3.*k2_1.*k3_4.*k4_5 + k0_1.*k1_3.*k2_3.*k3_5.*k4_2 + k0_2.*k1_0.*k2_3.*k3_4.*k4_5 + k0_2.*k1_0.*k2_4.*k3_5.*k4_3 + k0_2.*k1_2.*k2_3.*k3_5.*k4_2 + k0_2.*k1_2.*k2_4.*k3_1.*k4_5 + k0_2.*k1_3.*k2_1.*k3_5.*k4_3 + k0_1.*k1_2.*k2_3.*k3_4.*k4_5 + k0_1.*k1_2.*k2_4.*k3_5.*k4_3 + k0_1.*k1_3.*k2_1.*k3_5.*k4_5 + k0_1.*k1_3.*k2_3.*k3_5.*k4_3 + k0_1.*k1_3.*k2_4.*k3_2.*k4_5 + k0_2.*k1_0.*k2_3.*k3_5.*k4_5 + k0_2.*k1_0.*k2_4.*k3_4.*k4_5 + k0_2.*k1_2.*k2_3.*k3_5.*k4_3 + k0_2.*k1_2.*k2_4.*k3_2.*k4_5 + k0_2.*k1_3.*k2_1.*k3_4.*k4_5 + k0_2.*k1_3.*k2_3.*k3_5.*k4_2 + k0_1.*k1_2.*k2_3.*k3_5.*k4_5 + k0_1.*k1_2.*k2_4.*k3_4.*k4_5 + k0_1.*k1_3.*k2_3.*k3_4.*k4_5 + k0_1.*k1_3.*k2_4.*k3_5.*k4_3 + k0_2.*k1_0.*k2_4.*k3_5.*k4_5 + k0_2.*k1_2.*k2_3.*k3_4.*k4_5 + k0_2.*k1_2.*k2_4.*k3_5.*k4_3 + k0_2.*k1_3.*k2_1.*k3_5.*k4_5 + k0_2.*k1_3.*k2_3.*k3_5.*k4_3 + k0_2.*k1_3.*k2_4.*k3_2.*k4_5 + k0_1.*k1_2.*k2_4.*k3_5.*k4_5 + k0_1.*k1_3.*k2_3.*k3_5.*k4_5 + k0_1.*k1_3.*k2_4.*k3_4.*k4_5 + k0_2.*k1_2.*k2_3.*k3_5.*k4_5 + k0_2.*k1_2.*k2_4.*k3_4.*k4_5 + k0_2.*k1_3.*k2_3.*k3_4.*k4_5 + k0_2.*k1_3.*k2_4.*k3_5.*k4_3 + k0_1.*k1_3.*k2_4.*k3_5.*k4_5 + k0_2.*k1_2.*k2_4.*k3_5.*k4_5 + k0_2.*k1_3.*k2_3.*k3_5.*k4_5 + k0_2.*k1_3.*k2_4.*k3_4.*k4_5 + k0_2.*k1_3.*k2_4.*k3_5.*k4_5 + k0_2.*k1_0.*k2_4.*k3_1.*k5_3 + k0_1.*k1_2.*k2_4.*k3_1.*k5_3 + k0_1.*k1_3.*k2_0.*k3_4.*k5_3 + k0_2.*k1_0.*k2_4.*k3_1.*k5_4 + k0_2.*k1_0.*k2_4.*k3_2.*k5_3 + k0_1.*k1_2.*k2_4.*k3_1.*k5_4 + k0_1.*k1_2.*k2_4.*k3_2.*k5_3 + k0_1.*k1_3.*k2_0.*k3_4.*k5_4 + k0_1.*k1_3.*k2_1.*k3_4.*k5_3 + k0_2.*k1_0.*k2_3.*k3_4.*k5_3 + k0_2.*k1_0.*k2_4.*k3_2.*k5_4 + k0_2.*k1_2.*k2_4.*k3_1.*k5_3 + k0_1.*k1_2.*k2_3.*k3_4.*k5_3 + k0_1.*k1_2.*k2_4.*k3_2.*k5_4 + k0_1.*k1_3.*k2_0.*k3_5.*k5_4 + k0_1.*k1_3.*k2_1.*k3_4.*k5_4 + k0_1.*k1_3.*k2_4.*k3_2.*k5_3 + k0_2.*k1_0.*k2_3.*k3_4.*k5_4 + k0_2.*k1_0.*k2_4.*k3_4.*k5_3 + k0_2.*k1_2.*k2_4.*k3_1.*k5_4 + k0_2.*k1_2.*k2_4.*k3_2.*k5_3 + k0_2.*k1_3.*k2_1.*k3_4.*k5_3 + k0_1.*k1_2.*k2_3.*k3_4.*k5_4 + k0_1.*k1_2.*k2_4.*k3_4.*k5_3 + k0_1.*k1_3.*k2_1.*k3_5.*k5_4 + k0_1.*k1_3.*k2_3.*k3_4.*k5_3 + k0_1.*k1_3.*k2_4.*k3_2.*k5_4 + k0_2.*k1_0.*k2_3.*k3_5.*k5_4 + k0_2.*k1_0.*k2_4.*k3_4.*k5_4 + k0_2.*k1_2.*k2_3.*k3_4.*k5_3 + k0_2.*k1_2.*k2_4.*k3_2.*k5_4 + k0_2.*k1_3.*k2_1.*k3_4.*k5_4 + k0_2.*k1_3.*k2_4.*k3_2.*k5_3 + k0_1.*k1_2.*k2_3.*k3_5.*k5_4 + k0_1.*k1_2.*k2_4.*k3_4.*k5_4 + k0_1.*k1_3.*k2_3.*k3_4.*k5_4 + k0_1.*k1_3.*k2_4.*k3_4.*k5_3 + k0_2.*k1_0.*k2_4.*k3_5.*k5_4 + k0_2.*k1_2.*k2_3.*k3_4.*k5_4 + k0_2.*k1_2.*k2_4.*k3_4.*k5_3 + k0_2.*k1_3.*k2_1.*k3_5.*k5_4 + k0_2.*k1_3.*k2_3.*k3_4.*k5_3 + k0_2.*k1_3.*k2_4.*k3_2.*k5_4 + k0_1.*k1_2.*k2_4.*k3_5.*k5_4 + k0_1.*k1_3.*k2_3.*k3_5.*k5_4 + k0_1.*k1_3.*k2_4.*k3_4.*k5_4 + k0_2.*k1_2.*k2_3.*k3_5.*k5_4 + k0_2.*k1_2.*k2_4.*k3_4.*k5_4 + k0_2.*k1_3.*k2_3.*k3_4.*k5_4 + k0_2.*k1_3.*k2_4.*k3_4.*k5_3 + k0_1.*k1_3.*k2_4.*k3_5.*k5_4 + k0_2.*k1_2.*k2_4.*k3_5.*k5_4 + k0_2.*k1_3.*k2_3.*k3_5.*k5_4 + k0_2.*k1_3.*k2_4.*k3_4.*k5_4 + k0_2.*k1_3.*k2_4.*k3_5.*k5_4 + k0_1.*k1_3.*k2_0.*k4_2.*k5_3 + k0_1.*k1_3.*k2_0.*k4_2.*k5_4 + k0_1.*k1_3.*k2_0.*k4_3.*k5_3 + k0_1.*k1_3.*k2_1.*k4_2.*k5_3 + k0_2.*k1_0.*k2_3.*k4_2.*k5_3 + k0_1.*k1_2.*k2_3.*k4_2.*k5_3 + k0_1.*k1_3.*k2_0.*k4_3.*k5_4 + k0_1.*k1_3.*k2_1.*k4_2.*k5_4 + k0_1.*k1_3.*k2_1.*k4_3.*k5_3 + k0_2.*k1_0.*k2_3.*k4_2.*k5_4 + k0_2.*k1_0.*k2_3.*k4_3.*k5_3 + k0_2.*k1_3.*k2_1.*k4_2.*k5_3 + k0_1.*k1_2.*k2_3.*k4_2.*k5_4 + k0_1.*k1_2.*k2_3.*k4_3.*k5_3 + k0_1.*k1_3.*k2_0.*k4_5.*k5_3 + k0_1.*k1_3.*k2_1.*k4_3.*k5_4 + k0_1.*k1_3.*k2_3.*k4_2.*k5_3 + k0_2.*k1_0.*k2_3.*k4_3.*k5_4 + k0_2.*k1_0.*k2_4.*k4_3.*k5_3 + k0_2.*k1_2.*k2_3.*k4_2.*k5_3 + k0_2.*k1_3.*k2_1.*k4_2.*k5_4 + k0_2.*k1_3.*k2_1.*k4_3.*k5_3 + k0_1.*k1_2.*k2_3.*k4_3.*k5_4 + k0_1.*k1_2.*k2_4.*k4_3.*k5_3 + k0_1.*k1_3.*k2_1.*k4_5.*k5_3 + k0_1.*k1_3.*k2_3.*k4_2.*k5_4 + k0_1.*k1_3.*k2_3.*k4_3.*k5_3 + k0_2.*k1_0.*k2_3.*k4_5.*k5_3 + k0_2.*k1_0.*k2_4.*k4_3.*k5_4 + k0_2.*k1_2.*k2_3.*k4_2.*k5_4 + k0_2.*k1_2.*k2_3.*k4_3.*k5_3 + k0_2.*k1_3.*k2_1.*k4_3.*k5_4 + k0_2.*k1_3.*k2_3.*k4_2.*k5_3 + k0_1.*k1_2.*k2_3.*k4_5.*k5_3 + k0_1.*k1_2.*k2_4.*k4_3.*k5_4 + k0_1.*k1_3.*k2_3.*k4_3.*k5_4 + k0_1.*k1_3.*k2_4.*k4_3.*k5_3 + k0_2.*k1_0.*k2_4.*k4_5.*k5_3 + k0_2.*k1_2.*k2_3.*k4_3.*k5_4 + k0_2.*k1_2.*k2_4.*k4_3.*k5_3 + k0_2.*k1_3.*k2_1.*k4_5.*k5_3 + k0_2.*k1_3.*k2_3.*k4_2.*k5_4 + k0_2.*k1_3.*k2_3.*k4_3.*k5_3 + k0_1.*k1_2.*k2_4.*k4_5.*k5_3 + k0_1.*k1_3.*k2_3.*k4_5.*k5_3 + k0_1.*k1_3.*k2_4.*k4_3.*k5_4 + k0_2.*k1_2.*k2_3.*k4_5.*k5_3 + k0_2.*k1_2.*k2_4.*k4_3.*k5_4 + k0_2.*k1_3.*k2_3.*k4_3.*k5_4 + k0_2.*k1_3.*k2_4.*k4_3.*k5_3 + k0_1.*k1_3.*k2_4.*k4_5.*k5_3 + k0_2.*k1_2.*k2_4.*k4_5.*k5_3 + k0_2.*k1_3.*k2_3.*k4_5.*k5_3 + k0_2.*k1_3.*k2_4.*k4_3.*k5_4 + k0_2.*k1_3.*k2_4.*k4_5.*k5_3 + k0_2.*k1_0.*k3_1.*k4_2.*k5_3 + k0_1.*k1_2.*k3_1.*k4_2.*k5_3 + k0_2.*k1_0.*k3_1.*k4_2.*k5_4 + k0_2.*k1_0.*k3_1.*k4_3.*k5_3 + k0_2.*k1_0.*k3_2.*k4_2.*k5_3 + k0_1.*k1_2.*k3_1.*k4_2.*k5_4 + k0_1.*k1_2.*k3_1.*k4_3.*k5_3 + k0_1.*k1_2.*k3_2.*k4_2.*k5_3 + k0_2.*k1_0.*k3_1.*k4_3.*k5_4 + k0_2.*k1_0.*k3_2.*k4_2.*k5_4 + k0_2.*k1_0.*k3_2.*k4_3.*k5_3 + k0_2.*k1_2.*k3_1.*k4_2.*k5_3 + k0_1.*k1_2.*k3_1.*k4_3.*k5_4 + k0_1.*k1_2.*k3_2.*k4_2.*k5_4 + k0_1.*k1_2.*k3_2.*k4_3.*k5_3 + k0_1.*k1_3.*k3_2.*k4_2.*k5_3 + k0_2.*k1_0.*k3_1.*k4_5.*k5_3 + k0_2.*k1_0.*k3_2.*k4_3.*k5_4 + k0_2.*k1_0.*k3_4.*k4_2.*k5_3 + k0_2.*k1_2.*k3_1.*k4_2.*k5_4 + k0_2.*k1_2.*k3_1.*k4_3.*k5_3 + k0_2.*k1_2.*k3_2.*k4_2.*k5_3 + k0_1.*k1_2.*k3_1.*k4_5.*k5_3 + k0_1.*k1_2.*k3_2.*k4_3.*k5_4 + k0_1.*k1_2.*k3_4.*k4_2.*k5_3 + k0_1.*k1_3.*k3_2.*k4_2.*k5_4 + k0_1.*k1_3.*k3_2.*k4_3.*k5_3 + k0_2.*k1_0.*k3_2.*k4_5.*k5_3 + k0_2.*k1_0.*k3_4.*k4_2.*k5_4 + k0_2.*k1_2.*k3_1.*k4_3.*k5_4 + k0_2.*k1_2.*k3_2.*k4_2.*k5_4 + k0_2.*k1_2.*k3_2.*k4_3.*k5_3 + k0_2.*k1_3.*k3_2.*k4_2.*k5_3 + k0_1.*k1_2.*k3_2.*k4_5.*k5_3 + k0_1.*k1_2.*k3_4.*k4_2.*k5_4 + k0_1.*k1_3.*k3_2.*k4_3.*k5_4 + k0_1.*k1_3.*k3_4.*k4_2.*k5_3 + k0_2.*k1_0.*k3_5.*k4_2.*k5_4 + k0_2.*k1_2.*k3_1.*k4_5.*k5_3 + k0_2.*k1_2.*k3_2.*k4_3.*k5_4 + k0_2.*k1_2.*k3_4.*k4_2.*k5_3 + k0_2.*k1_3.*k3_2.*k4_2.*k5_4 + k0_2.*k1_3.*k3_2.*k4_3.*k5_3 + k0_1.*k1_2.*k3_5.*k4_2.*k5_4 + k0_1.*k1_3.*k3_2.*k4_5.*k5_3 + k0_1.*k1_3.*k3_4.*k4_2.*k5_4 + k0_2.*k1_2.*k3_2.*k4_5.*k5_3 + k0_2.*k1_2.*k3_4.*k4_2.*k5_4 + k0_2.*k1_3.*k3_2.*k4_3.*k5_4 + k0_2.*k1_3.*k3_4.*k4_2.*k5_3 + k0_1.*k1_3.*k3_5.*k4_2.*k5_4 + k0_2.*k1_2.*k3_5.*k4_2.*k5_4 + k0_2.*k1_3.*k3_2.*k4_5.*k5_3 + k0_2.*k1_3.*k3_4.*k4_2.*k5_4 + k0_2.*k1_3.*k3_5.*k4_2.*k5_4 + k0_1.*k2_0.*k3_1.*k4_2.*k5_3 + k0_1.*k2_0.*k3_1.*k4_2.*k5_4 + k0_1.*k2_0.*k3_1.*k4_3.*k5_3 + k0_1.*k2_0.*k3_2.*k4_2.*k5_3 + k0_1.*k2_1.*k3_1.*k4_2.*k5_3 + k0_1.*k2_0.*k3_1.*k4_3.*k5_4 + k0_1.*k2_0.*k3_2.*k4_2.*k5_4 + k0_1.*k2_0.*k3_2.*k4_3.*k5_3 + k0_1.*k2_1.*k3_1.*k4_2.*k5_4 + k0_1.*k2_1.*k3_1.*k4_3.*k5_3 + k0_1.*k2_1.*k3_2.*k4_2.*k5_3 + k0_2.*k2_1.*k3_1.*k4_2.*k5_3 + k0_1.*k2_0.*k3_1.*k4_5.*k5_3 + k0_1.*k2_0.*k3_2.*k4_3.*k5_4 + k0_1.*k2_0.*k3_4.*k4_2.*k5_3 + k0_1.*k2_1.*k3_1.*k4_3.*k5_4 + k0_1.*k2_1.*k3_2.*k4_2.*k5_4 + k0_1.*k2_1.*k3_2.*k4_3.*k5_3 + k0_1.*k2_3.*k3_1.*k4_2.*k5_3 + k0_2.*k2_1.*k3_1.*k4_2.*k5_4 + k0_2.*k2_1.*k3_1.*k4_3.*k5_3 + k0_2.*k2_1.*k3_2.*k4_2.*k5_3 + k0_1.*k2_0.*k3_2.*k4_5.*k5_3 + k0_1.*k2_0.*k3_4.*k4_2.*k5_4 + k0_1.*k2_1.*k3_1.*k4_5.*k5_3 + k0_1.*k2_1.*k3_2.*k4_3.*k5_4 + k0_1.*k2_1.*k3_4.*k4_2.*k5_3 + k0_1.*k2_3.*k3_1.*k4_2.*k5_4 + k0_1.*k2_3.*k3_1.*k4_3.*k5_3 + k0_2.*k2_1.*k3_1.*k4_3.*k5_4 + k0_2.*k2_1.*k3_2.*k4_2.*k5_4 + k0_2.*k2_1.*k3_2.*k4_3.*k5_3 + k0_2.*k2_3.*k3_1.*k4_2.*k5_3 + k0_1.*k2_0.*k3_5.*k4_2.*k5_4 + k0_1.*k2_1.*k3_2.*k4_5.*k5_3 + k0_1.*k2_1.*k3_4.*k4_2.*k5_4 + k0_1.*k2_3.*k3_1.*k4_3.*k5_4 + k0_1.*k2_4.*k3_1.*k4_3.*k5_3 + k0_2.*k2_1.*k3_1.*k4_5.*k5_3 + k0_2.*k2_1.*k3_2.*k4_3.*k5_4 + k0_2.*k2_1.*k3_4.*k4_2.*k5_3 + k0_2.*k2_3.*k3_1.*k4_2.*k5_4 + k0_2.*k2_3.*k3_1.*k4_3.*k5_3 + k0_1.*k2_1.*k3_5.*k4_2.*k5_4 + k0_1.*k2_3.*k3_1.*k4_5.*k5_3 + k0_1.*k2_4.*k3_1.*k4_3.*k5_4 + k0_2.*k2_1.*k3_2.*k4_5.*k5_3 + k0_2.*k2_1.*k3_4.*k4_2.*k5_4 + k0_2.*k2_3.*k3_1.*k4_3.*k5_4 + k0_2.*k2_4.*k3_1.*k4_3.*k5_3 + k0_1.*k2_4.*k3_1.*k4_5.*k5_3 + k0_2.*k2_1.*k3_5.*k4_2.*k5_4 + k0_2.*k2_3.*k3_1.*k4_5.*k5_3 + k0_2.*k2_4.*k3_1.*k4_3.*k5_4 + k0_2.*k2_4.*k3_1.*k4_5.*k5_3 + k1_0.*k2_0.*k3_1.*k4_2.*k5_3 + k1_0.*k2_0.*k3_1.*k4_2.*k5_4 + k1_0.*k2_0.*k3_1.*k4_3.*k5_3 + k1_0.*k2_0.*k3_2.*k4_2.*k5_3 + k1_0.*k2_1.*k3_1.*k4_2.*k5_3 + k1_0.*k2_0.*k3_1.*k4_3.*k5_4 + k1_0.*k2_0.*k3_2.*k4_2.*k5_4 + k1_0.*k2_0.*k3_2.*k4_3.*k5_3 + k1_0.*k2_1.*k3_1.*k4_2.*k5_4 + k1_0.*k2_1.*k3_1.*k4_3.*k5_3 + k1_0.*k2_1.*k3_2.*k4_2.*k5_3 + k1_2.*k2_0.*k3_1.*k4_2.*k5_3 + k1_0.*k2_0.*k3_1.*k4_5.*k5_3 + k1_0.*k2_0.*k3_2.*k4_3.*k5_4 + k1_0.*k2_0.*k3_4.*k4_2.*k5_3 + k1_0.*k2_1.*k3_1.*k4_3.*k5_4 + k1_0.*k2_1.*k3_2.*k4_2.*k5_4 + k1_0.*k2_1.*k3_2.*k4_3.*k5_3 + k1_0.*k2_3.*k3_1.*k4_2.*k5_3 + k1_2.*k2_0.*k3_1.*k4_2.*k5_4 + k1_2.*k2_0.*k3_1.*k4_3.*k5_3 + k1_2.*k2_0.*k3_2.*k4_2.*k5_3 + k1_0.*k2_0.*k3_2.*k4_5.*k5_3 + k1_0.*k2_0.*k3_4.*k4_2.*k5_4 + k1_0.*k2_1.*k3_1.*k4_5.*k5_3 + k1_0.*k2_1.*k3_2.*k4_3.*k5_4 + k1_0.*k2_1.*k3_4.*k4_2.*k5_3 + k1_0.*k2_3.*k3_1.*k4_2.*k5_4 + k1_0.*k2_3.*k3_1.*k4_3.*k5_3 + k1_2.*k2_0.*k3_1.*k4_3.*k5_4 + k1_2.*k2_0.*k3_2.*k4_2.*k5_4 + k1_2.*k2_0.*k3_2.*k4_3.*k5_3 + k1_3.*k2_0.*k3_2.*k4_2.*k5_3 + k1_0.*k2_0.*k3_5.*k4_2.*k5_4 + k1_0.*k2_1.*k3_2.*k4_5.*k5_3 + k1_0.*k2_1.*k3_4.*k4_2.*k5_4 + k1_0.*k2_3.*k3_1.*k4_3.*k5_4 + k1_0.*k2_4.*k3_1.*k4_3.*k5_3 + k1_2.*k2_0.*k3_1.*k4_5.*k5_3 + k1_2.*k2_0.*k3_2.*k4_3.*k5_4 + k1_2.*k2_0.*k3_4.*k4_2.*k5_3 + k1_3.*k2_0.*k3_2.*k4_2.*k5_4 + k1_3.*k2_0.*k3_2.*k4_3.*k5_3 + k1_0.*k2_1.*k3_5.*k4_2.*k5_4 + k1_0.*k2_3.*k3_1.*k4_5.*k5_3 + k1_0.*k2_4.*k3_1.*k4_3.*k5_4 + k1_2.*k2_0.*k3_2.*k4_5.*k5_3 + k1_2.*k2_0.*k3_4.*k4_2.*k5_4 + k1_3.*k2_0.*k3_2.*k4_3.*k5_4 + k1_3.*k2_0.*k3_4.*k4_2.*k5_3 + k1_0.*k2_4.*k3_1.*k4_5.*k5_3 + k1_2.*k2_0.*k3_5.*k4_2.*k5_4 + k1_3.*k2_0.*k3_2.*k4_5.*k5_3 + k1_3.*k2_0.*k3_4.*k4_2.*k5_4 + k1_3.*k2_0.*k3_5.*k4_2.*k5_4);
S0=(k1_0.*k2_0.*k3_1.*k4_2.*k5_3 + k1_0.*k2_0.*k3_1.*k4_2.*k5_4 + k1_0.*k2_0.*k3_1.*k4_3.*k5_3 + k1_0.*k2_0.*k3_2.*k4_2.*k5_3 + k1_0.*k2_1.*k3_1.*k4_2.*k5_3 + k1_0.*k2_0.*k3_1.*k4_3.*k5_4 + k1_0.*k2_0.*k3_2.*k4_2.*k5_4 + k1_0.*k2_0.*k3_2.*k4_3.*k5_3 + k1_0.*k2_1.*k3_1.*k4_2.*k5_4 + k1_0.*k2_1.*k3_1.*k4_3.*k5_3 + k1_0.*k2_1.*k3_2.*k4_2.*k5_3 + k1_2.*k2_0.*k3_1.*k4_2.*k5_3 + k1_0.*k2_0.*k3_1.*k4_5.*k5_3 + k1_0.*k2_0.*k3_2.*k4_3.*k5_4 + k1_0.*k2_0.*k3_4.*k4_2.*k5_3 + k1_0.*k2_1.*k3_1.*k4_3.*k5_4 + k1_0.*k2_1.*k3_2.*k4_2.*k5_4 + k1_0.*k2_1.*k3_2.*k4_3.*k5_3 + k1_0.*k2_3.*k3_1.*k4_2.*k5_3 + k1_2.*k2_0.*k3_1.*k4_2.*k5_4 + k1_2.*k2_0.*k3_1.*k4_3.*k5_3 + k1_2.*k2_0.*k3_2.*k4_2.*k5_3 + k1_0.*k2_0.*k3_2.*k4_5.*k5_3 + k1_0.*k2_0.*k3_4.*k4_2.*k5_4 + k1_0.*k2_1.*k3_1.*k4_5.*k5_3 + k1_0.*k2_1.*k3_2.*k4_3.*k5_4 + k1_0.*k2_1.*k3_4.*k4_2.*k5_3 + k1_0.*k2_3.*k3_1.*k4_2.*k5_4 + k1_0.*k2_3.*k3_1.*k4_3.*k5_3 + k1_2.*k2_0.*k3_1.*k4_3.*k5_4 + k1_2.*k2_0.*k3_2.*k4_2.*k5_4 + k1_2.*k2_0.*k3_2.*k4_3.*k5_3 + k1_3.*k2_0.*k3_2.*k4_2.*k5_3 + k1_0.*k2_0.*k3_5.*k4_2.*k5_4 + k1_0.*k2_1.*k3_2.*k4_5.*k5_3 + k1_0.*k2_1.*k3_4.*k4_2.*k5_4 + k1_0.*k2_3.*k3_1.*k4_3.*k5_4 + k1_0.*k2_4.*k3_1.*k4_3.*k5_3 + k1_2.*k2_0.*k3_1.*k4_5.*k5_3 + k1_2.*k2_0.*k3_2.*k4_3.*k5_4 + k1_2.*k2_0.*k3_4.*k4_2.*k5_3 + k1_3.*k2_0.*k3_2.*k4_2.*k5_4 + k1_3.*k2_0.*k3_2.*k4_3.*k5_3 + k1_0.*k2_1.*k3_5.*k4_2.*k5_4 + k1_0.*k2_3.*k3_1.*k4_5.*k5_3 + k1_0.*k2_4.*k3_1.*k4_3.*k5_4 + k1_2.*k2_0.*k3_2.*k4_5.*k5_3 + k1_2.*k2_0.*k3_4.*k4_2.*k5_4 + k1_3.*k2_0.*k3_2.*k4_3.*k5_4 + k1_3.*k2_0.*k3_4.*k4_2.*k5_3 + k1_0.*k2_4.*k3_1.*k4_5.*k5_3 + k1_2.*k2_0.*k3_5.*k4_2.*k5_4 + k1_3.*k2_0.*k3_2.*k4_5.*k5_3 + k1_3.*k2_0.*k3_4.*k4_2.*k5_4 + k1_3.*k2_0.*k3_5.*k4_2.*k5_4)./D;
S1=(k0_1.*k2_0.*k3_1.*k4_2.*k5_3 + k0_1.*k2_0.*k3_1.*k4_2.*k5_4 + k0_1.*k2_0.*k3_1.*k4_3.*k5_3 + k0_1.*k2_0.*k3_2.*k4_2.*k5_3 + k0_1.*k2_1.*k3_1.*k4_2.*k5_3 + k0_1.*k2_0.*k3_1.*k4_3.*k5_4 + k0_1.*k2_0.*k3_2.*k4_2.*k5_4 + k0_1.*k2_0.*k3_2.*k4_3.*k5_3 + k0_1.*k2_1.*k3_1.*k4_2.*k5_4 + k0_1.*k2_1.*k3_1.*k4_3.*k5_3 + k0_1.*k2_1.*k3_2.*k4_2.*k5_3 + k0_2.*k2_1.*k3_1.*k4_2.*k5_3 + k0_1.*k2_0.*k3_1.*k4_5.*k5_3 + k0_1.*k2_0.*k3_2.*k4_3.*k5_4 + k0_1.*k2_0.*k3_4.*k4_2.*k5_3 + k0_1.*k2_1.*k3_1.*k4_3.*k5_4 + k0_1.*k2_1.*k3_2.*k4_2.*k5_4 + k0_1.*k2_1.*k3_2.*k4_3.*k5_3 + k0_1.*k2_3.*k3_1.*k4_2.*k5_3 + k0_2.*k2_1.*k3_1.*k4_2.*k5_4 + k0_2.*k2_1.*k3_1.*k4_3.*k5_3 + k0_2.*k2_1.*k3_2.*k4_2.*k5_3 + k0_1.*k2_0.*k3_2.*k4_5.*k5_3 + k0_1.*k2_0.*k3_4.*k4_2.*k5_4 + k0_1.*k2_1.*k3_1.*k4_5.*k5_3 + k0_1.*k2_1.*k3_2.*k4_3.*k5_4 + k0_1.*k2_1.*k3_4.*k4_2.*k5_3 + k0_1.*k2_3.*k3_1.*k4_2.*k5_4 + k0_1.*k2_3.*k3_1.*k4_3.*k5_3 + k0_2.*k2_1.*k3_1.*k4_3.*k5_4 + k0_2.*k2_1.*k3_2.*k4_2.*k5_4 + k0_2.*k2_1.*k3_2.*k4_3.*k5_3 + k0_2.*k2_3.*k3_1.*k4_2.*k5_3 + k0_1.*k2_0.*k3_5.*k4_2.*k5_4 + k0_1.*k2_1.*k3_2.*k4_5.*k5_3 + k0_1.*k2_1.*k3_4.*k4_2.*k5_4 + k0_1.*k2_3.*k3_1.*k4_3.*k5_4 + k0_1.*k2_4.*k3_1.*k4_3.*k5_3 + k0_2.*k2_1.*k3_1.*k4_5.*k5_3 + k0_2.*k2_1.*k3_2.*k4_3.*k5_4 + k0_2.*k2_1.*k3_4.*k4_2.*k5_3 + k0_2.*k2_3.*k3_1.*k4_2.*k5_4 + k0_2.*k2_3.*k3_1.*k4_3.*k5_3 + k0_1.*k2_1.*k3_5.*k4_2.*k5_4 + k0_1.*k2_3.*k3_1.*k4_5.*k5_3 + k0_1.*k2_4.*k3_1.*k4_3.*k5_4 + k0_2.*k2_1.*k3_2.*k4_5.*k5_3 + k0_2.*k2_1.*k3_4.*k4_2.*k5_4 + k0_2.*k2_3.*k3_1.*k4_3.*k5_4 + k0_2.*k2_4.*k3_1.*k4_3.*k5_3 + k0_1.*k2_4.*k3_1.*k4_5.*k5_3 + k0_2.*k2_1.*k3_5.*k4_2.*k5_4 + k0_2.*k2_3.*k3_1.*k4_5.*k5_3 + k0_2.*k2_4.*k3_1.*k4_3.*k5_4 + k0_2.*k2_4.*k3_1.*k4_5.*k5_3)./D;
S2=(k0_2.*k1_0.*k3_1.*k4_2.*k5_3 + k0_1.*k1_2.*k3_1.*k4_2.*k5_3 + k0_2.*k1_0.*k3_1.*k4_2.*k5_4 + k0_2.*k1_0.*k3_1.*k4_3.*k5_3 + k0_2.*k1_0.*k3_2.*k4_2.*k5_3 + k0_1.*k1_2.*k3_1.*k4_2.*k5_4 + k0_1.*k1_2.*k3_1.*k4_3.*k5_3 + k0_1.*k1_2.*k3_2.*k4_2.*k5_3 + k0_2.*k1_0.*k3_1.*k4_3.*k5_4 + k0_2.*k1_0.*k3_2.*k4_2.*k5_4 + k0_2.*k1_0.*k3_2.*k4_3.*k5_3 + k0_2.*k1_2.*k3_1.*k4_2.*k5_3 + k0_1.*k1_2.*k3_1.*k4_3.*k5_4 + k0_1.*k1_2.*k3_2.*k4_2.*k5_4 + k0_1.*k1_2.*k3_2.*k4_3.*k5_3 + k0_1.*k1_3.*k3_2.*k4_2.*k5_3 + k0_2.*k1_0.*k3_1.*k4_5.*k5_3 + k0_2.*k1_0.*k3_2.*k4_3.*k5_4 + k0_2.*k1_0.*k3_4.*k4_2.*k5_3 + k0_2.*k1_2.*k3_1.*k4_2.*k5_4 + k0_2.*k1_2.*k3_1.*k4_3.*k5_3 + k0_2.*k1_2.*k3_2.*k4_2.*k5_3 + k0_1.*k1_2.*k3_1.*k4_5.*k5_3 + k0_1.*k1_2.*k3_2.*k4_3.*k5_4 + k0_1.*k1_2.*k3_4.*k4_2.*k5_3 + k0_1.*k1_3.*k3_2.*k4_2.*k5_4 + k0_1.*k1_3.*k3_2.*k4_3.*k5_3 + k0_2.*k1_0.*k3_2.*k4_5.*k5_3 + k0_2.*k1_0.*k3_4.*k4_2.*k5_4 + k0_2.*k1_2.*k3_1.*k4_3.*k5_4 + k0_2.*k1_2.*k3_2.*k4_2.*k5_4 + k0_2.*k1_2.*k3_2.*k4_3.*k5_3 + k0_2.*k1_3.*k3_2.*k4_2.*k5_3 + k0_1.*k1_2.*k3_2.*k4_5.*k5_3 + k0_1.*k1_2.*k3_4.*k4_2.*k5_4 + k0_1.*k1_3.*k3_2.*k4_3.*k5_4 + k0_1.*k1_3.*k3_4.*k4_2.*k5_3 + k0_2.*k1_0.*k3_5.*k4_2.*k5_4 + k0_2.*k1_2.*k3_1.*k4_5.*k5_3 + k0_2.*k1_2.*k3_2.*k4_3.*k5_4 + k0_2.*k1_2.*k3_4.*k4_2.*k5_3 + k0_2.*k1_3.*k3_2.*k4_2.*k5_4 + k0_2.*k1_3.*k3_2.*k4_3.*k5_3 + k0_1.*k1_2.*k3_5.*k4_2.*k5_4 + k0_1.*k1_3.*k3_2.*k4_5.*k5_3 + k0_1.*k1_3.*k3_4.*k4_2.*k5_4 + k0_2.*k1_2.*k3_2.*k4_5.*k5_3 + k0_2.*k1_2.*k3_4.*k4_2.*k5_4 + k0_2.*k1_3.*k3_2.*k4_3.*k5_4 + k0_2.*k1_3.*k3_4.*k4_2.*k5_3 + k0_1.*k1_3.*k3_5.*k4_2.*k5_4 + k0_2.*k1_2.*k3_5.*k4_2.*k5_4 + k0_2.*k1_3.*k3_2.*k4_5.*k5_3 + k0_2.*k1_3.*k3_4.*k4_2.*k5_4 + k0_2.*k1_3.*k3_5.*k4_2.*k5_4)./D;
S3=(k0_1.*k1_3.*k2_0.*k4_2.*k5_3 + k0_1.*k1_3.*k2_0.*k4_2.*k5_4 + k0_1.*k1_3.*k2_0.*k4_3.*k5_3 + k0_1.*k1_3.*k2_1.*k4_2.*k5_3 + k0_2.*k1_0.*k2_3.*k4_2.*k5_3 + k0_1.*k1_2.*k2_3.*k4_2.*k5_3 + k0_1.*k1_3.*k2_0.*k4_3.*k5_4 + k0_1.*k1_3.*k2_1.*k4_2.*k5_4 + k0_1.*k1_3.*k2_1.*k4_3.*k5_3 + k0_2.*k1_0.*k2_3.*k4_2.*k5_4 + k0_2.*k1_0.*k2_3.*k4_3.*k5_3 + k0_2.*k1_3.*k2_1.*k4_2.*k5_3 + k0_1.*k1_2.*k2_3.*k4_2.*k5_4 + k0_1.*k1_2.*k2_3.*k4_3.*k5_3 + k0_1.*k1_3.*k2_0.*k4_5.*k5_3 + k0_1.*k1_3.*k2_1.*k4_3.*k5_4 + k0_1.*k1_3.*k2_3.*k4_2.*k5_3 + k0_2.*k1_0.*k2_3.*k4_3.*k5_4 + k0_2.*k1_0.*k2_4.*k4_3.*k5_3 + k0_2.*k1_2.*k2_3.*k4_2.*k5_3 + k0_2.*k1_3.*k2_1.*k4_2.*k5_4 + k0_2.*k1_3.*k2_1.*k4_3.*k5_3 + k0_1.*k1_2.*k2_3.*k4_3.*k5_4 + k0_1.*k1_2.*k2_4.*k4_3.*k5_3 + k0_1.*k1_3.*k2_1.*k4_5.*k5_3 + k0_1.*k1_3.*k2_3.*k4_2.*k5_4 + k0_1.*k1_3.*k2_3.*k4_3.*k5_3 + k0_2.*k1_0.*k2_3.*k4_5.*k5_3 + k0_2.*k1_0.*k2_4.*k4_3.*k5_4 + k0_2.*k1_2.*k2_3.*k4_2.*k5_4 + k0_2.*k1_2.*k2_3.*k4_3.*k5_3 + k0_2.*k1_3.*k2_1.*k4_3.*k5_4 + k0_2.*k1_3.*k2_3.*k4_2.*k5_3 + k0_1.*k1_2.*k2_3.*k4_5.*k5_3 + k0_1.*k1_2.*k2_4.*k4_3.*k5_4 + k0_1.*k1_3.*k2_3.*k4_3.*k5_4 + k0_1.*k1_3.*k2_4.*k4_3.*k5_3 + k0_2.*k1_0.*k2_4.*k4_5.*k5_3 + k0_2.*k1_2.*k2_3.*k4_3.*k5_4 + k0_2.*k1_2.*k2_4.*k4_3.*k5_3 + k0_2.*k1_3.*k2_1.*k4_5.*k5_3 + k0_2.*k1_3.*k2_3.*k4_2.*k5_4 + k0_2.*k1_3.*k2_3.*k4_3.*k5_3 + k0_1.*k1_2.*k2_4.*k4_5.*k5_3 + k0_1.*k1_3.*k2_3.*k4_5.*k5_3 + k0_1.*k1_3.*k2_4.*k4_3.*k5_4 + k0_2.*k1_2.*k2_3.*k4_5.*k5_3 + k0_2.*k1_2.*k2_4.*k4_3.*k5_4 + k0_2.*k1_3.*k2_3.*k4_3.*k5_4 + k0_2.*k1_3.*k2_4.*k4_3.*k5_3 + k0_1.*k1_3.*k2_4.*k4_5.*k5_3 + k0_2.*k1_2.*k2_4.*k4_5.*k5_3 + k0_2.*k1_3.*k2_3.*k4_5.*k5_3 + k0_2.*k1_3.*k2_4.*k4_3.*k5_4 + k0_2.*k1_3.*k2_4.*k4_5.*k5_3)./D;
S4=(k0_2.*k1_0.*k2_4.*k3_1.*k5_3 + k0_1.*k1_2.*k2_4.*k3_1.*k5_3 + k0_1.*k1_3.*k2_0.*k3_4.*k5_3 + k0_2.*k1_0.*k2_4.*k3_1.*k5_4 + k0_2.*k1_0.*k2_4.*k3_2.*k5_3 + k0_1.*k1_2.*k2_4.*k3_1.*k5_4 + k0_1.*k1_2.*k2_4.*k3_2.*k5_3 + k0_1.*k1_3.*k2_0.*k3_4.*k5_4 + k0_1.*k1_3.*k2_1.*k3_4.*k5_3 + k0_2.*k1_0.*k2_3.*k3_4.*k5_3 + k0_2.*k1_0.*k2_4.*k3_2.*k5_4 + k0_2.*k1_2.*k2_4.*k3_1.*k5_3 + k0_1.*k1_2.*k2_3.*k3_4.*k5_3 + k0_1.*k1_2.*k2_4.*k3_2.*k5_4 + k0_1.*k1_3.*k2_0.*k3_5.*k5_4 + k0_1.*k1_3.*k2_1.*k3_4.*k5_4 + k0_1.*k1_3.*k2_4.*k3_2.*k5_3 + k0_2.*k1_0.*k2_3.*k3_4.*k5_4 + k0_2.*k1_0.*k2_4.*k3_4.*k5_3 + k0_2.*k1_2.*k2_4.*k3_1.*k5_4 + k0_2.*k1_2.*k2_4.*k3_2.*k5_3 + k0_2.*k1_3.*k2_1.*k3_4.*k5_3 + k0_1.*k1_2.*k2_3.*k3_4.*k5_4 + k0_1.*k1_2.*k2_4.*k3_4.*k5_3 + k0_1.*k1_3.*k2_1.*k3_5.*k5_4 + k0_1.*k1_3.*k2_3.*k3_4.*k5_3 + k0_1.*k1_3.*k2_4.*k3_2.*k5_4 + k0_2.*k1_0.*k2_3.*k3_5.*k5_4 + k0_2.*k1_0.*k2_4.*k3_4.*k5_4 + k0_2.*k1_2.*k2_3.*k3_4.*k5_3 + k0_2.*k1_2.*k2_4.*k3_2.*k5_4 + k0_2.*k1_3.*k2_1.*k3_4.*k5_4 + k0_2.*k1_3.*k2_4.*k3_2.*k5_3 + k0_1.*k1_2.*k2_3.*k3_5.*k5_4 + k0_1.*k1_2.*k2_4.*k3_4.*k5_4 + k0_1.*k1_3.*k2_3.*k3_4.*k5_4 + k0_1.*k1_3.*k2_4.*k3_4.*k5_3 + k0_2.*k1_0.*k2_4.*k3_5.*k5_4 + k0_2.*k1_2.*k2_3.*k3_4.*k5_4 + k0_2.*k1_2.*k2_4.*k3_4.*k5_3 + k0_2.*k1_3.*k2_1.*k3_5.*k5_4 + k0_2.*k1_3.*k2_3.*k3_4.*k5_3 + k0_2.*k1_3.*k2_4.*k3_2.*k5_4 + k0_1.*k1_2.*k2_4.*k3_5.*k5_4 + k0_1.*k1_3.*k2_3.*k3_5.*k5_4 + k0_1.*k1_3.*k2_4.*k3_4.*k5_4 + k0_2.*k1_2.*k2_3.*k3_5.*k5_4 + k0_2.*k1_2.*k2_4.*k3_4.*k5_4 + k0_2.*k1_3.*k2_3.*k3_4.*k5_4 + k0_2.*k1_3.*k2_4.*k3_4.*k5_3 + k0_1.*k1_3.*k2_4.*k3_5.*k5_4 + k0_2.*k1_2.*k2_4.*k3_5.*k5_4 + k0_2.*k1_3.*k2_3.*k3_5.*k5_4 + k0_2.*k1_3.*k2_4.*k3_4.*k5_4 + k0_2.*k1_3.*k2_4.*k3_5.*k5_4)./D;
S5=1-S0-S1-S2-S3-S4;

% cyt c reduction
Jc = p.CIII.Etot.*(kfQH2p_01.*S0 + kfQH2p_12.*S1 + kfQH2p_23.*S2 + kfQH2p_34.*S3 - krQH2p_01.*S1 - krQH2p_12.*S2 - krQH2p_23.*S3 - krQH2p_34.*S4 + kfQH2p_45.*S4 - krQH2p_45.*S5);

% QH2 oxidaiton at Qp
JQp = Jc;

% superoxide formation
JSO = p.CIII.Etot.*(kfSO_10.*S1 + kfSO_21.*S2 + kfSO_32.*S3 + kfSO_43.*S4 + kfSO_54.*S5 - krSO_10.*S0 - krSO_21.*S1 - krSO_32.*S2 - krSO_43.*S3 - krSO_54.*S4);

% Q reduction at Qn
JQn = (Jc - JSO)/2;
% JQn = p.CIII.Etot.*(kfQH2n_20.*S2 + kfQH2n_31.*S3 + kfQH2n_42.*S4 - krQH2n_20.*S0 - krQH2n_31.*S1 - krQH2n_42.*S2 + kfQH2n_53.*S5 - krQH2n_53.*S3); % equal to above

% Collect model details and package into structure

% midpoint potentials
details.Em=Em;
details.Emb=Emb;
% equilibrium constants
details.Keq=Keq;
details.Keqb=Keqb;
% binding polynomials
details.P_Qp=P_Qp;
details.P_Qn=P_Qn;
details.P_c=P_c;
% substate occupancies
details.s0=s0;
details.s1=s1;
details.s2=s2;
details.s3=s3;
details.s4=s4;
details.s5=s5;
% state occupancies
details.S0=S0;
details.S1=S1;
details.S2=S2;
details.S3=S3;
details.S4=S4;
details.S5=S5;
% turnover at each site
details.Jc=Jc;
details.JQp=JQp;
details.JQn=JQn;
details.JSO=JSO;
% partial reactions
details.kfQH2n_20 = kfQH2n_20;
details.krQH2n_20 = krQH2n_20;
details.kfQH2n_31 = kfQH2n_31;
details.krQH2n_31 = krQH2n_31;
details.kfQH2n_42 = kfQH2n_42;
details.krQH2n_42 = krQH2n_42;
details.kfQH2n_53 = kfQH2n_53;
details.krQH2n_53 = krQH2n_53;
details.kfQH2p_01 = kfQH2p_01;
details.krQH2p_01 = krQH2p_01;
details.kfQH2p_12 = kfQH2p_12;
details.krQH2p_12 = krQH2p_12;
details.kfQH2p_23 = kfQH2p_23;
details.krQH2p_23 = krQH2p_23;
details.kfQH2p_34 = kfQH2p_34;
details.krQH2p_34 = krQH2p_34;
details.kfQH2p_45 = kfQH2p_45;
details.krQH2p_45 = krQH2p_45;
details.kfSO_10 = kfSO_10;
details.kfSO_21 = kfSO_21;
details.kfSO_32 = kfSO_32;
details.kfSO_43 = kfSO_43;
details.kfSO_54 = kfSO_54;
details.krSO_10 = krSO_10;
details.krSO_21 = krSO_21;
details.krSO_32 = krSO_32;
details.krSO_43 = krSO_43;
details.krSO_54 = krSO_54;
% free energy spans
dG = -2*p.F*(Em.c - Em.Q_QH2p) - 2*p.F*(Em.Q_QH2n - Em.Q_QH2p) + 2*p.F*dPsi + p.R*p.T*log(c2.^2.*Q./c3.^2./QH2); % dGr in absence of O2 and SO
dGso = -p.F*(Em.c + p.CIII.Em.SO - 2*Em.Q_QH2p) + p.R*p.T*log(c2.*Q.*SO./c3./QH2./O2); % dGr of only SO related reactions
dGtot = -2*p.F*(Em.c - Em.Q_QH2p) - p.F*(Em.c + p.CIII.Em.SO - 2*Em.Q_QH2p) - 2*p.F*(Em.Q_QH2n - Em.Q_QH2p) + 2*p.F*dPsi + p.R*p.T*log(c2.^3.*Q.*SO./c3.^3./QH2./O2); % full dGr
details.dG = dG;
details.dGso = dGso;
details.dGtot = dGtot;

% Output

if nargout > 1
    varargout = {Jc, JQp, JQn, JSO, details};
else
    varargout = {Jc};
end

end

%% Complex IV
% based on Brand's JO vs c2+ vs dPsi surface

function J = CIV(x,p)

% Define SVs into generalized notation
c2 = x(p.i.c2_x,:);         % M
c3 = x(p.i.c3_x,:);         % M
O2 = x(p.i.O2_x,:);         % M
Hx = x(p.i.H_x,:);          % M
Hi = x(p.i.H_c,:);          % M
dPsi = x(p.i.dPsi,:);       % mV

%Calculate pH independnt dG0
dG0 = 2*p.dGf(p.if.c3) + p.dGf(p.if.H2O) ...
    - 2*p.dGf(p.if.c2) - 0.5*p.dGf(p.if.O2);

AC4=dG0+4*p.F*dPsi;  % partial dG spanning complex, 4H+/2e-
Keq=exp(-AC4./p.R./p.T).*(Hx.^4./Hi.^2);

KM = p.CIV.KM; % M
beta = p.CIV.beta; % unitless

J = p.CIV.x.*(O2./(O2+p.CIV.k_O2)).*(c2.^2./(c2.^2 + KM^2*(1+beta*exp(dPsi*2*p.F/p.R/p.T)))).*(1 - (c3.^2./c2.^2./O2.^.5)./Keq);

end

%% H leak model based on data from Serviddio and Sastre in Advanced Protocols in Oxidative Stress II
% energy barrier makes inward fluxes more sensitive to dPsi, b=3/4
% data from 7.4 on page 117
% collected at 37 C without nigericin
% but also fits data from Porter et al. 1999 Fig 2a
% rat liver mito at 37 C with nigericin

function J = mHleak(x,p)

% Define SVs into generalized notation
Hx = x(p.i.H_x,:);    % M
Hi = x(p.i.H_c,:);    % M
DPsi = x(p.i.dPsi,:);   % mV

x_HLE = p.mHleak.x;

b=3/4; % based on fit to data given above

J = x_HLE.*(Hi.*exp(p.F.*DPsi./p.R./p.T*b) ...
    -Hx.*exp(-p.F.*DPsi./p.R./p.T*(1-b)));

end

%% Mass action ATP Synthase function

function J = F1FO(x,p)

% Define SVs into generalized notation
ADP = x(p.i.ADP_x,:);   % M
Pi = x(p.i.Pi_x,:);     % M
ATP = x(p.i.ATP_x,:);   % M
Hx = x(p.i.H_x,:);      % M
Hi = x(p.i.H_c,:);      % M
dPsi = x(p.i.dPsi,:);   % mV

% define reference reacation dG
dGr = p.dGf(p.if.ATP) + p.dGf(p.if.H2O) - p.dGf(p.if.ADP) ...
    - p.dGf(p.if.Pi); % kJ/mol

% define total free energy adding contribution to proton motive force
Keq0 = exp(-(dGr - (p.F1FO.nH)*p.F*dPsi)./p.R./p.T);

% define overall equlibrium constant
Keq = Keq0.*(Hi.^(p.F1FO.nH)./Hx.^(p.F1FO.nH-1)).*BP(p.if.ATP,'mtx',x,p) ...
    ./BP(p.if.ADP,'mtx',x,p)./BP(p.if.Pi,'mtx',x,p);

k1_F1FO = p.F1FO.x;

J = k1_F1FO.*(Keq.*ADP.*Pi-ATP);

end

%% modified Metelkin ANT model from Bazil et al. 2010

function J = ANT(x,p)

% load substrate state variables
ATPe = x(p.i.ATP_c,:); % M
ADPe = x(p.i.ADP_c,:); % M
ATPm = x(p.i.ATP_x,:); % M
ADPm = x(p.i.ADP_x,:); % M
dPsi = -x(p.i.dPsi,:);  % mV

% calculate ATP^4- and ADP^3- concentrations
fATPe = ATPe./BP(p.if.ATP,'cyt',x,p);
fADPe = ADPe./BP(p.if.ADP,'cyt',x,p);
fATPm = ATPm./BP(p.if.ATP,'mtx',x,p);
fADPm = ADPm./BP(p.if.ADP,'mtx',x,p);

% ATP/ATP exchange
% k1 = p.ANT.k10.*exp((4.*p.ANT.a1-4.*p.ANT.a2+p.ANT.a3) ...
%     .*dPsi.*p.F/p.R./p.T);
% ADPe/ATPm exchange
k2 = p.ANT.k20.*exp((3*p.ANT.a1-4*p.ANT.a2+p.ANT.a3) ...
    .*dPsi.*p.F./p.R./p.T);
% ATPe/ADPm exchange (solved for using Haldane constraint)
k3 = p.ANT.k30.*exp((4.*p.ANT.a1-3.*p.ANT.a2+p.ANT.a3) ...
    .*dPsi.*p.F./p.R./p.T);
% ADP/ADP exchange
% k4 = p.ANT.k40.*exp((3.*p.ANT.a1-3.*p.ANT.a2+p.ANT.a3) ...
%     .*dPsi.*p.F/p.R./p.T);


% define AdN binding constants
KTe = p.ANT.KT0e.*exp(-4.*p.ANT.deltaT_o.*dPsi.*p.F./p.R./p.T);
KDe = p.ANT.KD0e.*exp(-3.*p.ANT.deltaD_o.*dPsi.*p.F./p.R./p.T);

% define the equilibrium constant (Keq = (ToDi/TiDo)eq = k2KDiKTo/k3KTiKDo)
Keq = exp(-dPsi.*p.F./p.R./p.T); 
% solve for k3 using Haldane constraint
q = k3.*KDe./k2./KTe.*Keq; %(q=KDm./KTm)

% define enzyme-substrate states
den=(1+fATPe./KTe + fADPe./KDe).*(q.*fATPm + fADPm);

% define exchange rates
%Ex1 = k1.*q.*fATPm.*fATPe./KTe;
Ex2 = k2.*q.*fATPm.*fADPe./KDe;
Ex3 = k3.*fADPm.*fATPe./KTe;
%Ex4 = k4.*fADPm.*fADPe./KDe;

% net hetero-exchange defined as
J = p.ANT.Etot.*(Ex2-Ex3)./den; % nmol/mg/min

end

%% Inorganic Phosphate Carrier

function J = PiC(x,p)

% Define SVs into generalized notation
Pii = x(p.i.Pi_c,:);
Pix = x(p.i.Pi_x,:);
Hi = x(p.i.H_c,:);
Hx = x(p.i.H_x,:);

A = Pii.*Hi./p.KD(p.if.Pi,p.if.H)./BP(p.if.Pi,'cyt',x,p);
P = Pix.*Hx./p.KD(p.if.Pi,p.if.H)./BP(p.if.Pi,'mtx',x,p);

x_PIH = p.PIC.x;
k_PIH = p.PIC.kPi;

J = x_PIH./k_PIH.*(Hi.*A - Hx.*P)./(1+A/k_PIH)./(1+P/k_PIH);

end

%% Dicarboxylate Carrier
% Dicarboxylate Carrier (DCC)
% The DCC exchanges TCA cycle intermediates malate (MAL), succinate (SUC)
% and hydrogen phosphate (HPO4). The enzyme rxn was modeled based on a
% rapid equilibrium bi-bi mechanism. The kinetic parameter set was fit to
% the expercental data from Indiveri et al. 
% In the heart, DCC can exchange either MAL or SUC for Pi with ~ equal
% affinity. The enzyme turn-over is comparable to other transporters (ANT,
% GAE, AKG/MAL exchanger), but the enzyme is expressed at lower level in
% the heart. 

function J = DCC(x,p)

% Load substrate state variables and convert from nmol/mg to M units
Pic = x(p.i.Pi_c,:);      
Pix = x(p.i.Pi_x,:);        
MALc = x(p.i.MAL_c,:);    
MALx = x(p.i.MAL_x,:);      
SUCc = x(p.i.SUC_c,:);   
SUCx = x(p.i.SUC_x,:);      

% Calculate ionic species involved in reactions
HPO4c = Pic./p.BP(p.i.Pi_c,:);
HPO4x = Pix./p.BP(p.i.Pi_x,:);
fMALc = MALc./p.BP(p.i.MAL_c,:);
fMALx = MALx./p.BP(p.i.MAL_x,:);
fSUCc = SUCc./p.BP(p.i.SUC_c,:);
fSUCx = SUCx./p.BP(p.i.SUC_x,:);

% Redefine state variable names into generalized notation
S1o = fMALc;
S2o = fSUCc;
S3o = HPO4c;
S1i = fMALx;
S2i = fSUCx;
S3i = HPO4x;

% redefine parameter names into generalized notation
KS1o = p.DCC.K_MAL;
KS1i = KS1o; % assume binding symmetry
KS2o = p.DCC.K_SUC;
KS2i = KS2o; % assume binding symmetry
KS3o = p.DCC.K_Pi;
KS3i = KS3o; % assume binding symmetry

% Keq = 1

Do = 1 + S1o/KS1o + S2o/KS2o + S3o/KS3o;
Di = 1 + S1i/KS1i + S2i/KS2i + S3i/KS3i;

% Assumed binding of substrates are independent of each other and forward
% and reverse rates are equal, can make all variable with proper
% constraints if required. also, we are not requiring Pi to be the
% antiporter conterpart and allow homo and hetero exchange

k=1; % assume symmetric flipping rates
k12 = k.*(S1o./KS1o./Do + S2o./KS2o./Do + S3o./KS3o./Do); % flux o -> i
k21 = k.*(S1i./KS1i./Di + S2i./KS2i./Di + S3i./KS3i./Di); % flux i -> o

% steady-state enzyme fractions
ETo = k21./(k12 + k21);
ETi = 1 - ETo;

% Calculate net fluxes of individual substrates through DCC
J.MAL = p.DCC.Vmax.*(S1o/KS1o.*ETo./Do - S1i/KS1i.*ETi./Di);
J.SUC = p.DCC.Vmax.*(S2o/KS2o.*ETo./Do - S2i/KS2i.*ETi./Di);
J.Pi = p.DCC.Vmax.*(S3o/KS3o.*ETo./Do - S3i/KS3i.*ETi./Di);

end

%% Monocarboxylate Transporter

% PYR^{-}-H^{+} co-transporter (PYRH)

function J = MCT(x,p)

H_c = x(p.i.H_c,:);
H_x = x(p.i.H_x,:);
PYR_c = x(p.i.PYR_c,:);
PYR_x = x(p.i.PYR_x,:);

fPYR_c = PYR_c./p.BP(p.i.PYR_c,:); % free pyruvate concentration outside mitos
fPYR_x = PYR_x./p.BP(p.i.PYR_x,:); % free pyruvate concentration inside matrix

J = p.MCT.x.*(fPYR_c.*H_c - fPYR_x.*H_x);

end

%% Mass action KHE model

function J = KHE(x,p)

J = p.mKHE.x*(x(p.i.K_x,:).*x(p.i.H_c,:) - x(p.i.K_c,:).*x(p.i.H_x,:));

end

%% Simple MCU model

function J = MCU(x,p)

% Define SVs into generalized notation
Ca_x = x(p.i.Ca_x,:);    % M
Ca_c = x(p.i.Ca_c,:);    % M
dPsi = x(p.i.dPsi,:);    % mV

KCa = p.MCU.KCa;

u = 2*dPsi.*p.F./p.R./p.T; % normalized potential

n = p.MCU.x.*(exp(u/2).*(Ca_c./KCa).^p.MCU.nCa - exp(-u/2).*(Ca_x./KCa).^p.MCU.nCa);

d= (1 + (Ca_c./KCa).^p.MCU.nCa + (Ca_x./KCa).^p.MCU.nCa);

J = n./d;

end

%% NCLX from Bazil et al. 2010 without calcium activation

function J = NCLX(x,p)

% Define SVs into generalized notation
Caim=x(p.i.Ca_c,:);     % M
Naim = x(p.i.Na_c,:);   % M
Cam = x(p.i.Ca_x,:);    % M
Nam = x(p.i.Na_x,:);    % M
dPsi = x(p.i.dPsi,:);   % mV

u=dPsi.*p.F./p.R./p.T;

a = p.NCLX.x.*(exp(u.*2/4).*Naim.^3.*Cam./(p.NCLX.KNa.^3.*p.NCLX.KCa) ...
    - exp(-u.*2/4).*Nam.^3.*Caim./(p.NCLX.KNa.^3.*p.NCLX.KCa));

b = 1 + Naim.^3./p.NCLX.KNa.^3 + Cam./p.NCLX.KCa ...
    + Naim.^3.*Cam./(p.NCLX.KNa.^3.*p.NCLX.KCa) ...
    + Nam.^3./p.NCLX.KNa.^3 + Caim./p.NCLX.KCa ...
    + Nam.^3.*Caim./(p.NCLX.KNa.^3.*p.NCLX.KCa);

J=a./b;

end

%% Mitochondrial NaH Exchanger
% from Bazil et al. 2010

function J = NHE(x,p)

% load substrate state variables and convert from nmol/mg to M units
Hc = x(p.i.H_c,:);     % M
Nac = x(p.i.Na_c,:);   % M
Hx = x(p.i.H_x,:);     % M
Nax = x(p.i.Na_x,:);   % M

KmNam = p.NHE.KmNa.*Hx./(Hx+p.NHE.KmH);
KmNaim = p.NHE.KmNa.*Hc./(Hc+p.NHE.KmH);

a = p.NHE.x.*(Hx.^p.NHE.n)./(Hx.^p.NHE.n+p.NHE.KmHreg.^p.NHE.n);
b = Hc.*Nax./(p.NHE.KmH.*KmNam) - Hx.*Nac./(p.NHE.KmH.*KmNaim);
c = 1 + Hc./p.NHE.KmH + Nax./KmNam + Hc.*Nax./(p.NHE.KmH.*KmNam) ...
    + Hx./p.NHE.KmH + Nac./KmNaim + Hx.*Nac./(p.NHE.KmH.*KmNaim);

J = a.*b./c;

end

%% Mass action model for calcium phosphate formation

function f = CaPi(x,p,kf,K,n,m)

H = x(p.i.H_x,:);     % M
Ca = x(p.i.Ca_x,:);   % M
Pi = x(p.i.Pi_x,:);   % M
CaPi = x(p.i.CaPi,:); % M

f = kf*(Ca.^n.*(Pi./p.BP(p.if.Pi,:)).^m - CaPi.*H.^(2*n-2*m)./K);

end

%% Hybrid mass action/kinetic ATPase function

function J = ATPase(x,p)

ATP = x(p.i.ATP_c,:);   % M
ADP = x(p.i.ADP_c,:);   % M
Pi = x(p.i.Pi_c,:);     % M
H = x(p.i.H_c,:);       % M

ATP=max(1e-12,ATP);

dGr_atpase = p.dGf(p.if.ADP) + p.dGf(p.if.Pi) - p.dGf(p.if.ATP) ...
    - p.dGf(p.if.H2O);
Keq_atpase = exp(-dGr_atpase/p.R./p.T);
Keq = Keq_atpase.*(1./H) ...
    .*(BP(p.if.ADP,'cyt',x,p).*BP(p.if.Pi,'cyt',x,p)) ...
    ./BP(p.if.ATP,'cyt',x,p);

J = p.ATPase.x*p.ATPase.Etot./(1+x(p.i.ADP_c,:)/p.ATPase.Ki_ADP).*(1 - ADP.*Pi./ATP./Keq);

end
```
