## Supplementary material for "Identifying Site-specific Superoxide and Hydrogen Peroxide Production Rates from the Mitochondrial Electron Transport System Using a Computational Strategy": Model Codes: Generate_Figures.m

```
%% Plot Model Outputs
% Inputs 
% data - data structure
% p - parameter structure
% Load data and parameters for data and p structures, respectively
%
% Outputs
% figures used in main text and supplemental material, note that some
% legends and axes labels were done outside of Matlab
%
% Copyright
% Jason Bazil 2021
% Bazil Laboratory
% Department of Physiology
% Michigan State University

function model = Generate_Figures(data,p)
%% Setup simulations

% Set model
fun = @ETS_ROS_model; 
 
modeloutputs = {@PM,@SR,@S,@PMS,@PM_O2,@SRot_O2,@MAL,@PM_Fig10};

model = getoutput(fun,p,modeloutputs);

% define model indices
iPM = find(strcmp(model.label,'PM'));
iSR = find(strcmp(model.label,'SR'));
iS = find(strcmp(model.label,'S'));
iPMS = find(strcmp(model.label,'PMS'));
iMAL = find(strcmp(model.label,'MAL'));
iPM_O2 = find(strcmp(model.label,'PM_O2'));
iSR_O2 = find(strcmp(model.label,'SR_O2'));
iPM_Fig10 = find(strcmp(model.label,'PM_Fig10'));

% set color scheme
c = [0    0.4470    0.7410
    0.8500    0.3250    0.0980
    0.9290    0.6940    0.1250
    0.4940    0.1840    0.5560
    0.4660    0.6740    0.1880
    0.3010    0.7450    0.9330
    0.6350    0.0780    0.1840];

%% Figure 1: Model schematic

%% FIGURE 2: In-silico reproduction of JO2 (nmol/mg/min)

% Panel A: P/M - JO2
figure
hold on
box on
% Model respiratory profile 
plot(model.output{iPM}{1},-model.output{iPM}{4}.dO2*1e7,'color',c(1,:),'linewidth',8) % Model parameters are based on nmol/mg/min, so no further conversion is necessary. 
% Data 
yMAL_leak = [data.PM.JO2_leak1]; % These data were already multiplied by 0.6 to get the correct unit of nmol/mg/min
yPM_leak2 = [data.PM.JO2_leak2];
yMAL_oxphos = [data.PM.JO2_oxphos];
% Individual data points
plot(4.5+(0.1-0.2*rand(1,numel(yMAL_leak))),yMAL_leak,'ok','markersize',20,'linewidth',2)
plot(9.5+(0.1-0.2*rand(1,numel(yPM_leak2))),yPM_leak2,'ok','markersize',20,'linewidth',2)
plot(10.5+(0.1-0.2*rand(1,numel(yMAL_oxphos))),yMAL_oxphos,'ok','markersize',20,'linewidth',2)
% Error bars
errorbar(4.5,mean(yMAL_leak),std(yMAL_leak),'.','color',[0 0 0],'markersize',1,'linewidth',2)
errorbar(9.5,mean(yPM_leak2),std(yPM_leak2),'.','color',[0 0 0],'markersize',1,'linewidth',2)
errorbar(10.5,mean(yMAL_oxphos),std(yMAL_oxphos),'.','color',[0 0 0],'markersize',1,'linewidth',2)
% Axes 
set(gcf, 'Position', [1 1 1920 1080]);
ylabel({'J_{O_2}(nmol O_2/mg/min)'},'color',[0 0 0],'fontsize',60);
ylim([0 450])
xlim([0 17])
yticks([0 100 200 300 400])
xticks([0 5 10 15])
xticklabels([0 5 10 15])
xlabel({'Time (min)'});
set(gca,'FontName','Georgia','FontSize',40);
set(gcf,'color','none'),set(gca,'color','none')

% Panel D: P/M - JH2O2
figure
hold on
box on
% Model JH2O2 profile 
plot(model.output{iPM}{1},model.output{iPM}{4}.HrP*1e10,'color',c(1,:),'linewidth',8)
% Data 
yMAL_leak = [data.PM.JH2O2_leak1];
yPM_leak2 = [data.PM.JH2O2_leak2];
yMAL_oxphos = [data.PM.JH2O2_oxphos];
% Individual data points
plot(4.5+(0.1-0.2*rand(1,numel(yMAL_leak))),yMAL_leak,'ok','markersize',20,'linewidth',2)
plot(9.5+(0.1-0.2*rand(1,numel(yPM_leak2))),yPM_leak2,'ok','markersize',20,'linewidth',2)
plot(10.5+(0.1-0.2*rand(1,numel(yMAL_oxphos))),yMAL_oxphos,'ok','markersize',20,'linewidth',2)
% Error bars
errorbar(4.5,mean(yMAL_leak),std(yMAL_leak),'.','color',[0 0 0],'markersize',1,'linewidth',2)
errorbar(9.5,mean(yPM_leak2),std(yPM_leak2),'.','color',[0 0 0],'markersize',1,'linewidth',2)
errorbar(10.5,mean(yMAL_oxphos),std(yMAL_oxphos),'.','color',[0 0 0],'markersize',1,'linewidth',2)
% Axes 
set(gcf, 'Position', [1 1 1920 1080]); 
ylabel({'J_{H_2O_2}(pmol/mg/min)'},'color',[0 0 0],'fontsize',60);
xlabel({'Time (min)'},'color',[0 0 0],'fontsize',60);
ylim([0 150])
xlim([0 17])
yticks([0 20 40 60 80 100 120 140])
xticks([0 5 10 15])
xticklabels([0 5 10 15])
set(gca,'FontName','Georgia','FontSize',40);
set(gcf,'color','none'),set(gca,'color','none')

% Panel B: S/R - JO2
figure
hold on
box on
% Model Respiratory Profile 
plot(model.output{iSR}{1},-model.output{iSR}{4}.dO2*1e7,'color',c(2,:),'linewidth',8)
% Data 
ySR_leak1 = [data.SR.JO2_leak1];
ySR_leak2 = [data.SR.JO2_leak2];
ySR_oxphos = [data.SR.JO2_oxphos];
% Individual data points
plot(2.5+(0.1-0.2*rand(1,numel(ySR_leak1))),ySR_leak1,'ok','markersize',20,'linewidth',2)
plot(4.5+(0.1-0.2*rand(1,numel(ySR_leak2))),ySR_leak2,'ok','markersize',20,'linewidth',2)
plot(5.5+(0.1-0.2*rand(1,numel(ySR_oxphos))),ySR_oxphos,'ok','markersize',20,'linewidth',2)
% Error bars
errorbar(2.5,mean(ySR_leak1),std(ySR_leak1),'.','color',[0 0 0],'markersize',1,'linewidth',2)
errorbar(4.5,mean(ySR_leak2),std(ySR_leak2),'.','color',[0 0 0],'markersize',1,'linewidth',2)
errorbar(5.5,mean(ySR_oxphos),std(ySR_oxphos),'.','color',[0 0 0],'markersize',1,'linewidth',2)
% Axes 
set(gcf, 'Position', [1 1 1920 1080]); 
ylabel({'J_{O_2}(nmol O_2/mg/min)'},'color',[0 0 0],'fontsize',60);
xlabel({'Time (min)'},'color',[0 0 0],'fontsize',60);
ylim([0 450])
xlim([0 8.5])
yticks([0 100 200 300 400])
xticks([0 2 4 6 8 10])
xticklabels([0 2 4 6 8 10]);
set(gca,'FontName','Georgia','FontSize',40);
set(gcf,'color','none'),set(gca,'color','none')

% Pabel E: S/R - JH2O2
figure
hold on
box on
% Model JH2O2 profile 
plot(model.output{iSR}{1},model.output{iSR}{4}.HrP*1e10,'color',c(2,:),'linewidth',8)
% Data 
ySR_leak1 = [data.SR.JH2O2_leak1];
ySR_leak2 = [data.SR.JH2O2_leak2];
ySR_oxphos = [data.SR.JH2O2_oxphos];
% Individual data points
plot(2.5+(0.1-0.2*rand(1,numel(ySR_leak1))),ySR_leak1,'ok','markersize',20,'linewidth',2)
plot(4.5+(0.1-0.2*rand(1,numel(ySR_leak2))),ySR_leak2,'ok','markersize',20,'linewidth',2)
plot(5.5+(0.1-0.2*rand(1,numel(ySR_oxphos))),ySR_oxphos,'ok','markersize',20,'linewidth',2)
% Error bars
errorbar(2.5,mean(ySR_leak1),std(ySR_leak1),'.','color',[0 0 0],'markersize',1,'linewidth',2)
errorbar(4.5,mean(ySR_leak2),std(ySR_leak2),'.','color',[0 0 0],'markersize',1,'linewidth',2)
errorbar(5.5,mean(ySR_oxphos),std(ySR_oxphos),'.','color',[0 0 0],'markersize',1,'linewidth',2)
% Axes 
set(gcf, 'Position', [1 1 1920 1080]); 
ylabel({'J_{H_2O_2}(pmol/mg/min)'},'color',[0 0 0],'fontsize',60);
xlabel({'Time (min)'},'color',[0 0 0],'fontsize',60);
ylim([0 300])
xlim([0 8.5])
yticks([0 50 100 150 200 250 300])
xticks([0 2 4 6 8])
xticklabels([0 2 4 6 8])
set(gca,'FontName','Georgia','FontSize',40);
set(gcf,'color','none'),set(gca,'color','none')

% Panel C: Succinate - JO2
figure
hold on
box on
% Model respiratory profile 
plot(model.output{iS}{1},-model.output{iS}{4}.dO2*1e7,'color',c(3,:),'linewidth',8) % Model parameters are based on nmol/mg/min, so no further conversion is necessary. 
% Data 
ySuc_leak1 = [data.S.JO2_leak1]; % These data were already multiplied by 0.6 to get the correct unit of nmol/mg/min
ySuc_leak2 = [data.S.JO2_leak2];
ySuc_oxphos = [data.S.JO2_oxphos];
% Individual data points
plot(2+(0.1-0.2*rand(1,numel(ySuc_leak1))),ySuc_leak1,'ok','markersize',20,'linewidth',2)
plot(4+(0.1-0.2*rand(1,numel(ySuc_leak2))),ySuc_leak2,'ok','markersize',20,'linewidth',2)
plot(5.5+(0.1-0.2*rand(1,numel(ySuc_oxphos))),ySuc_oxphos,'ok','markersize',20,'linewidth',2)
% Error bars
errorbar(2,mean(ySuc_leak1),std(ySuc_leak1),'.','color',[0 0 0],'markersize',1,'linewidth',2)
errorbar(4,mean(ySuc_leak2),std(ySuc_leak2),'.','color',[0 0 0],'markersize',1,'linewidth',2)
errorbar(5.5,mean(ySuc_oxphos),std(ySuc_oxphos),'.','color',[0 0 0],'markersize',1,'linewidth',2)
% Axes 
set(gcf, 'Position', [1 1 1920 1080]); 
ylabel({'J_{O_2}(nmol O_2/mg/min)'},'color',[0 0 0],'fontsize',60);
ylim([0 450])
xlim([0 8.5])
yticks([0 100 200 300 400])
xticks([0 2 4 6 8])
xticklabels([0 2 4 6 8])
xlabel({'Time (min)'});
set(gca,'FontName','Georgia','FontSize',40);
set(gcf,'color','none'),set(gca,'color','none')

% Panel F: Succinate - JH2O2
figure
hold on
box on
% Model JH2O2 profile 
plot(model.output{iS}{1},model.output{iS}{4}.HrP*1e-3/0.1*1e12,'color',c(3,:),'linewidth',8)
% Data 
yS_leak1 = [data.S.JH2O2_leak1];
yS_leak2 = [data.S.JH2O2_leak2];
yS_oxphos = [data.S.JH2O2_oxphos];
% Individual data points
plot(2+(0.1-0.2*rand(1,numel(yS_leak1))),yS_leak1,'ok','markersize',20,'linewidth',2)
plot(4+(0.1-0.2*rand(1,numel(yS_leak2))),yS_leak2,'ok','markersize',20,'linewidth',2)
plot(5.5+(0.1-0.2*rand(1,numel(yS_oxphos))),yS_oxphos,'ok','markersize',20,'linewidth',2)
% Error bars
errorbar(2,mean(yS_leak1),std(yS_leak1),'.','color',[0 0 0],'markersize',1,'linewidth',2)
errorbar(4,mean(yS_leak2),std(yS_leak2),'.','color',[0 0 0],'markersize',1,'linewidth',2)
errorbar(5.5,mean(yS_oxphos),std(yS_oxphos),'.','color',[0 0 0],'markersize',1,'linewidth',2)
% Axes 
set(gcf, 'Position', [1 1 1920 1080]); 
ylabel({'J_{H_2O_2}(pmol/mg/min)'},'color',[0 0 0],'fontsize',60);
xlabel({'Time (min)'},'color',[0 0 0],'fontsize',60);
ylim([0 7000])
xlim([0 8.5])
yticks([0 1000 2000 3000 4000 5000 6000])
xticks([0 2 4 6 8])
xticklabels([0 2 4 6 8])
set(gca,'FontName','Georgia','FontSize',40,'Color',[0 0 0]);
set(gcf,'color','none'),set(gca,'color','none')

%% FIG 3: NADH Reduction 

% Panel A: P/M   
figure
hold on
box on
% Model NADH dynamics
plot(model.output{iPM}{1},model.output{iPM}{2}(:,p.i.NADH_x)./p.Ntot*100,'color',c(1,:),'linewidth',8)
% Data
N_PMleak = [data.PM.NADH_leak];
N_PMoxphos = [data.PM.NADH_oxphos];
% Individual data points
plot(4.5+(0.1-0.2*rand(1,numel(N_PMleak))),N_PMleak,'ok','markersize',20,'linewidth',2)
plot(10.5+(0.1-0.2*rand(1,numel(N_PMoxphos))),N_PMoxphos,'ok','markersize',20,'linewidth',2)
% Error bars
errorbar(4.5,mean(N_PMleak),std(N_PMleak),'.','color',[0 0 0],'markersize',1,'linewidth',2)
errorbar(10.5,mean(N_PMoxphos),std(N_PMoxphos),'.','color',[0 0 0],'markersize',1,'linewidth',2)
% Axes 
set(gcf, 'Position', [1 1 1920 1080]); % Maximize figure.
ylabel({'% NADH Reduction'},'color',[0 0 0],'fontsize',60);
xlabel({'Time (min)'},'color',[0 0 0],'fontsize',60);
ylim([0 125])
xlim([1e-9 17])
yticks([0 25 50 75 100 125])
xticks([0 5 10 15 20])
xticklabels([0 5 10 15 20])
set(gca,'FontName','Georgia','FontSize',40);
set(gcf,'color','none'),set(gca,'color','none')

% Panel B: S/R
figure
hold on
box on
% Model NADH dynamics
plot(model.output{iSR}{1},model.output{iSR}{2}(:,p.i.NADH_x)./p.Ntot*100,'color',c(2,:),'linewidth',8)
plot(data.SNdyn(:,1),data.SNdyn(:,2)*100,'o','color',[0 0 0],'markersize',20,'linewidth',2)
set(gcf, 'Position', [1 1 1920 1080]); % Maximize figure.
ylabel({'% NADH Reduction'},'color',[0 0 0],'fontsize',60);
xlabel({'Time (min)'},'color',[0 0 0],'fontsize',60);
ylim([0 125])
xlim([1e-9 8.5])
yticks([0 25 50 75 100 125])
xticks([0 2 4 6 8])
xticklabels([0 2 4 6 8]);
set(gca,'FontName','Georgia','FontSize',40);
set(gcf,'color','none'),set(gca,'color','none')

%% MODEL VALIDATION
%% FIG 4: P/M/S JO2, JH2O2 and [O2]

% Panel A: JO2
figure
hold on
box on
% Model respiratory profile 
plot(model.output{iPMS}{1},-model.output{iPMS}{4}.dO2*1e7,'color',c(4,:),'linewidth',8)
% Data 
yPMS_leak1 = [data.PMS.JO2_leak1];
yPMS_leak2 = [data.PMS.JO2_leak2];
yPMS_oxphos = [data.PMS.JO2_oxphos];
% Individual data points
plot(1.75+(0.1-0.2*rand(1,numel(yPMS_leak1))),yPMS_leak1,'ok','markersize',20,'linewidth',2)
plot(3+(0.1-0.2*rand(1,numel(yPMS_leak2))),yPMS_leak2,'ok','markersize',20,'linewidth',2)
plot(4.5+(0.1-0.2*rand(1,numel(yPMS_oxphos))),yPMS_oxphos,'ok','markersize',20,'linewidth',2)
% Error bars
errorbar(1.75,mean(yPMS_leak1),std(yPMS_leak1),'.','color',[0 0 0],'markersize',1,'linewidth',2)
errorbar(3,mean(yPMS_leak2),std(yPMS_leak2),'.','color',[0 0 0],'markersize',1,'linewidth',2)
errorbar(4.5,mean(yPMS_oxphos),std(yPMS_oxphos),'.','color',[0 0 0],'markersize',1,'linewidth',2)
% Axes 
set(gcf, 'Position', [1 1 1920 1080]); % Maximize figure.
% ylabel({'J_{O_2}(nmol O_2/mg/min)'},'color',[0 0 0],'fontsize',60);
xlabel({'Time (min)'},'color',[0 0 0],'fontsize',60);
ylim([0 900])
xlim([0 6.5])
yticks([0 200 400 600 800 1000])
xticks([0 2 4 6])
xticklabels([0 2 4 6]);
set(gca,'FontName','Georgia','FontSize',40);
set(gcf,'color','none'),set(gca,'color','none')

% Panel B: JH2O2
figure
hold on
box on
% Model H2O2 dynamics 
plot(model.output{iPMS}{1},model.output{iPMS}{4}.HrP*1e-3/0.1*1e12,'color',c(4,:),'linewidth',8)
% Data 
yPMS_leak1 = [data.PMS.JH2O2_leak1];
yPMS_leak2 = [data.PMS.JH2O2_leak2];
yPMS_oxphos = [data.PMS.JH2O2_oxphos];
% Individual data points
plot(1.75+(0.1-0.2*rand(1,numel(yPMS_leak1))),yPMS_leak1,'ok','markersize',20,'linewidth',2)
plot(3+(0.1-0.2*rand(1,numel(yPMS_leak2))),yPMS_leak2,'ok','markersize',20,'linewidth',2)
plot(4.5+(0.1-0.2*rand(1,numel(yPMS_oxphos))),yPMS_oxphos,'ok','markersize',20,'linewidth',2)
% Error bars
errorbar(1.75,mean(yPMS_leak1),std(yPMS_leak1),'.','color',[0 0 0],'markersize',1,'linewidth',2)
errorbar(3,mean(yPMS_leak2),std(yPMS_leak2),'.','color',[0 0 0],'markersize',1,'linewidth',2)
errorbar(4.5,mean(yPMS_oxphos),std(yPMS_oxphos),'.','color',[0 0 0],'markersize',1,'linewidth',2)
% Axes 
set(gcf, 'Position', [1 1 1920 1080]); % Maximize figure.
% ylabel({'J_{H_2O_2}(pmol/mg/min)'},'color',[0 0 0],'fontsize',60);
xlabel({'Time (min)'},'color',[0 0 0],'fontsize',60);
ylim([0 6000])
xlim([0 6.5])
yticks([0 1000 2000 3000 4000 5000])
xticks([0 2 4 6])
xticklabels([0 2 4 6]);
set(gca,'FontName','Georgia','FontSize',40);
set(gcf,'color','none'),set(gca,'color','none')

% Panel C: [O2]
figure
hold on
box on
% Model [O2] dynamics 
plot(model.output{iPMS}{1},model.output{iPMS}{2}(:,p.i.O2_x).*1e6,'color',c(4,:),'linewidth',8)
% Data 
yPMS_leak1 = [data.PMS.O2_leak1];
yPMS_leak2 = [data.PMS.O2_leak2];
yPMS_oxphos = [data.PMS.O2_oxphos];
% Individual data points
plot(2+(0.1-0.2*rand(1,numel(yPMS_leak1))),yPMS_leak1,'ok','markersize',20,'linewidth',2)
plot(4+(0.1-0.2*rand(1,numel(yPMS_leak2))),yPMS_leak2,'ok','markersize',20,'linewidth',2)
plot(5+(0.1-0.2*rand(1,numel(yPMS_oxphos))),yPMS_oxphos,'ok','markersize',20,'linewidth',2)
% Error bars
errorbar(2,mean(yPMS_leak1),std(yPMS_leak1),'.','color',[0 0 0],'markersize',1,'linewidth',2)
errorbar(4,mean(yPMS_leak2),std(yPMS_leak2),'.','color',[0 0 0],'markersize',1,'linewidth',2)
errorbar(5,mean(yPMS_oxphos),std(yPMS_oxphos),'.','color',[0 0 0],'markersize',1,'linewidth',2)
% Axes 
set(gcf, 'Position', [1 1 1920 1080]); % Maximize figure.
% ylabel({'[O_2](\muM)'},'color',[0 0 0],'fontsize',60);
xlabel({'Time (min)'},'color',[0 0 0],'fontsize',60);
ylim([0 200])
xlim([0 6.5])
yticks([0 50 100 150 200])
xticks([0 2 4 6])
xticklabels([0 2 4 6]);
set(gca,'FontName','Georgia','FontSize',40);
set(gcf,'color','none'),set(gca,'color','none')

%% FIG 5: JH2O2 v.s. [O2]

in = model.output{iPM_O2}{1}>1; % ignore 1st 1 min of simulation time

% Panel A: P/M
figure
hold on
box on
plot(model.output{iPM_O2}{2}(in,p.i.O2_x)*1e6,model.output{iPM_O2}{4}.HrP(in)*1e-3/0.1*1e12,'color',c(1,:),'linewidth',4) % converted to the correct unit in the fitting f(x)
% errorbar(data.Y{1,id(i)}(:,1),data.Y{1,id(i)}(:,2),data.Y{1,id(i)}(:,3),'o','MarkerSize',18,'color',[0 0 0],'linewidth',2)
plot(data.PM_O2(:,1),data.PM_O2(:,2),'o','color',[0 0 0],'markersize',20,'linewidth',2)
set(gcf, 'Position', get(0,'Screensize')); % Maximize figure.
ylabel({'J_{H_2O_2}(pmol/mg/min)'},'color',[0 0 0],'fontsize',60);
xlabel({'[O_2] (\muM)'},'color',[0 0 0],'fontsize',60);
xlim([0 200])
ylim([0 150])
set(gca,'FontSize',54,'FontName','Times New Roman')
set(gcf,'color','none'),set(gca,'color','none')

in = model.output{iSR_O2}{1}>1; % ignore 1st 1 min of simulation time

% Panel B: S/R
figure
hold on
box on
plot(model.output{iSR_O2}{2}(in,p.i.O2_x)*1e6,model.output{iSR_O2}{4}.HrP(in)*1e-3/0.1*1e12,'color',c(2,:),'linewidth',4) % converted to the correct unit in the fitting f(x)
% errorbar(data.Y{1,id(i)}(:,1),data.Y{1,id(i)}(:,2),data.Y{1,id(i)}(:,3),'o','MarkerSize',18,'color',[0 0 0],'linewidth',2)
plot(data.SR_O2(:,1),data.SR_O2(:,2),'o','color',[0 0 0],'markersize',20,'linewidth',2)
set(gcf, 'Position', get(0,'Screensize')); % Maximize figure.
ylabel({'J_{H_2O_2}(pmol/mg/min)'},'color',[0 0 0],'fontsize',60);
xlabel({'[O_2] (\muM)'},'color',[0 0 0],'fontsize',60);
xlim([0 200])
ylim([0 300])
set(gca,'FontSize',54,'FontName','Times New Roman')
set(gcf,'color','none'),set(gca,'color','none')

% Panel B inset
figure
hold on
box on
plot(model.output{iSR_O2}{2}(in,p.i.O2_x)*1e6,1./model.output{iSR_O2}{4}.CI.d.PNr(in)*100,'color',c(2,:),'linewidth',4)
xlim([1e-9 200])
ylabel({'% free FMN_{red}'},'color',[0 0 0],'fontsize',60);
xlabel({'[O_2] (\muM)'},'color',[0 0 0],'fontsize',60);
set(gca,'FontSize',54,'FontName','Times New Roman')
set(gcf,'color','none'),set(gca,'color','none')
set(gcf, 'Position', get(0,'Screensize')); % Maximize figure.

%% MODEL PREDICTION
%% FIG 7. Model analysis to explain the monotonicity of JH2O2 on [O2]

in = model.output{iPM_O2}{1}>1; % ignore 1st 1 min of simulation time

% P/M
% JROS & Scavenging activity
fig = figure;
left_color = [0 0 0];
right_color = [0 0 0];
set(fig,'defaultAxesColorOrder',[left_color; right_color]);
hold on
box on
yyaxis left
% Scavenging Rate
plot(model.output{iPM_O2}{2}(in,p.i.O2_x).*1e6,p.Vol.H2O_mtx*model.output{iPM_O2}{4}.H2O2scavenge(in),'color',[0 0 0],'linewidth',6)
% total SO
plot(model.output{iPM_O2}{2}(in,p.i.O2_x).*1e6,(model.output{iPM_O2}{4}.CI.JSO(in)/2 + model.output{iPM_O2}{4}.CII.JSO(in)/2 + model.output{iPM_O2}{4}.CIII.JSO(in)/2),'-','color',c(1,:),'linewidth',6)
% total H2O2 
plot(model.output{iPM_O2}{2}(in,p.i.O2_x).*1e6,(model.output{iPM_O2}{4}.CI.JH2O2(in) + model.output{iPM_O2}{4}.CII.JH2O2(in)),'--','color',c(1,:),'linewidth',6)
ylabel({'J_{ROS}(nmol/min/mg)'},'color',[0 0 0],'fontsize',60);
xlabel({'[O_2] (\muM)'},'color',[0 0 0],'fontsize',60);
set(gca,'FontSize',45,'FontName','Times New Roman')
ylim([0 1])
xlim([0 200])
yyaxis right
ylim([0 1/(p.Vmax_scavening*p.Vol.H2O_mtx)])
ylabel({'Scavenging Capacity'},'color',[0 0 0],'fontsize',60);
set(gcf,'color','none'),set(gca,'color','none')
set(gcf, 'Position', [1 1 1920 1080]); % Maximize figure.

% Hydrogen peroxide by IF and IIF
figure 
hold on
box on
plot(model.output{iPM_O2}{2}(in,p.i.O2_x).*1e6,model.output{iPM_O2}{4}.CI.d.JFrH2O2(in),'-','color',c(1,:),'linewidth',6)
plot(model.output{iPM_O2}{2}(in,p.i.O2_x).*1e6,model.output{iPM_O2}{4}.CII.d.JFADH2_H2O2(in),'--','color',c(1,:),'linewidth',6)
% plot(model.output{1,18}(:,p.i.O2_x).*1e6,model.output{1,20}.CI.JH2O2 + model.output{1,20}.CII.JH2O2,'color',[0 0 0],'linewidth',3)
ylabel({'J_{H2O2}(nmol/min.g)'},'color',[0 0 0],'fontsize',45);
xlabel({'[O_2] (\muM)'},'color',[0 0 0],'fontsize',45);
ylim([0 0.2])
xlim([0 200])
set(gca,'FontSize',45,'FontName','Times New Roman')
set(gcf,'color','none'),set(gca,'color','none')
set(gcf, 'Position', [1 1 1920 1080]); % Maximize figure.

% NADH & Q pool redox state
figure
hold on
box on
plot(model.output{iPM_O2}{2}(in,p.i.O2_x).*1e6,model.output{iPM_O2}{2}(in,p.i.NADH_x)/p.Ntot,'color',c(5,:),'linewidth',6)
set(gcf, 'Position', [1 1 1920 1080]); % Maximize figure.
ylabel({'Fractional NADH'},'color',[0 0 0],'fontsize',45);
xlabel({'[O_2] (\muM)'},'color',[0 0 0],'fontsize',45);
ylim([0 1])
xlim([0 200])
set(gca,'FontSize',45,'FontName','Times New Roman')
plot(model.output{iPM_O2}{2}(in,p.i.O2_x).*1e6,model.output{iPM_O2}{2}(in,p.i.UQH2_x)/p.Qtot,'color',c(6,:),'linewidth',6)
ylabel({'Fraction Reduced'},'color',[0 0 0],'fontsize',45);
set(gcf,'color','none'),set(gca,'color','none')

in = model.output{iSR_O2}{1}>1; % ignore 1st 1 min of simulation time

% S/R
% JROS & Scavenging activity
fig = figure;
left_color = [0 0 0];
right_color = [0 0 0];
set(fig,'defaultAxesColorOrder',[left_color; right_color]);
hold on
box on
yyaxis left
% Scavenging Rate
plot(model.output{iSR_O2}{2}(in,p.i.O2_x).*1e6,p.Vol.H2O_mtx*model.output{iSR_O2}{4}.H2O2scavenge(in),'color',[0 0 0],'linewidth',6)
% total SO
plot(model.output{iSR_O2}{2}(in,p.i.O2_x).*1e6,(model.output{iSR_O2}{4}.CI.JSO(in)/2 + model.output{iSR_O2}{4}.CII.JSO(in)/2 + model.output{iSR_O2}{4}.CIII.JSO(in)/2),'-','color',c(2,:),'linewidth',6)
% total H2O2 
plot(model.output{iSR_O2}{2}(in,p.i.O2_x).*1e6,(model.output{iSR_O2}{4}.CI.JH2O2(in) + model.output{iSR_O2}{4}.CII.JH2O2(in)),'--','color',c(2,:),'linewidth',6)
ylabel({'J_{ROS}(nmol/min/mg)'},'color',[0 0 0],'fontsize',60);
xlabel({'[O_2] (\muM)'},'color',[0 0 0],'fontsize',60);
set(gca,'FontSize',45,'FontName','Times New Roman')
ylim([0 5])
xlim([0 200])
yyaxis right
ylim([0 5/(p.Vmax_scavening*p.Vol.H2O_mtx)])
ylabel({'Scavenging Capacity'},'color',[0 0 0],'fontsize',60);
set(gcf,'color','none'),set(gca,'color','none')
set(gcf, 'Position', [1 1 1920 1080]); % Maximize figure.

% Hydrogen peroxide by IF and IIF
figure 
hold on
box on
plot(model.output{iSR_O2}{2}(in,p.i.O2_x).*1e6,model.output{iSR_O2}{4}.CI.d.JFrH2O2(in),'-','color',c(2,:),'linewidth',6)
plot(model.output{iSR_O2}{2}(in,p.i.O2_x).*1e6,model.output{iSR_O2}{4}.CII.d.JFADH2_H2O2(in),'--','color',c(2,:),'linewidth',6)
% plot(model.output{1,18}(:,p.i.O2_x).*1e6,model.output{1,20}.CI.JH2O2 + model.output{1,20}.CII.JH2O2,'color',[0 0 0],'linewidth',3)
ylabel({'J_{H2O2}(nmol/min.g)'},'color',[0 0 0],'fontsize',45);
xlabel({'[O_2] (\muM)'},'color',[0 0 0],'fontsize',45);
% ylim([0 0.2])
xlim([0 200])
set(gca,'FontSize',45,'FontName','Times New Roman')
set(gcf,'color','none'),set(gca,'color','none')
set(gcf, 'Position', [1 1 1920 1080]); % Maximize figure.

% NADH & Q pool redox state
figure
hold on
box on
plot(model.output{iSR_O2}{2}(in,p.i.O2_x).*1e6,model.output{iSR_O2}{2}(in,p.i.NADH_x)/p.Ntot,'color',c(5,:),'linewidth',6)
set(gcf, 'Position', [1 1 1920 1080]); % Maximize figure.
ylabel({'Fractional NADH'},'color',[0 0 0],'fontsize',45);
xlabel({'[O_2] (\muM)'},'color',[0 0 0],'fontsize',45);
ylim([0 1])
xlim([0 200])
set(gca,'FontSize',45,'FontName','Times New Roman')
plot(model.output{iSR_O2}{2}(in,p.i.O2_x).*1e6,model.output{iSR_O2}{2}(in,p.i.UQH2_x)/p.Qtot,'color',c(6,:),'linewidth',6)
ylabel({'Fraction Reduced'},'color',[0 0 0],'fontsize',45);
set(gcf,'color','none'),set(gca,'color','none')


%% FIG 8: Site-specific SO production 
% Panel A: LEAK

% Time intervals
ileak_PM = (model.output{iPM}{1} >= 4 & model.output{iPM}{1} < 5)';  
ileak_SR = (model.output{iSR}{1} >= 2.5 & model.output{iSR}{1} < 3)';
ileak_S = (model.output{iS}{1} >= 2.5 & model.output{iS}{1} < 3)';
ileak_PMS = (model.output{iPMS}{1} >= 2 & model.output{iPMS}{1} < 2.5)';

% P/M
CIa_SQSO = mean(model.output{iPM}{4}.CI.d.JSQSO(:,ileak_PM)); % SO from the semiquinone
CIa_FrSO = mean(model.output{iPM}{4}.CI.d.JFrSO(:,ileak_PM)); % SO from the fully reduced FMN
CIa_FradSO = mean(model.output{iPM}{4}.CI.d.JFradSO(:,ileak_PM)); % SO from the FMN radical
CIIa_ISC3SO = mean(model.output{iPM}{4}.CII.d.JISC3_SO(:,ileak_PM)); % SO from the [3Fe-4S] cluster
CIIa_FradSO = mean(model.output{iPM}{4}.CII.d.JFADH_SO(:,ileak_PM)); % SO from the FMN radical
CIIIa_QpSO = mean(model.output{iPM}{4}.CIII.d.JSO(:,ileak_PM));

% S/R
CIb_SQSO = mean(model.output{iSR}{4}.CI.d.JSQSO(:,ileak_SR)); % SO from the semiquinone
CIb_FrSO = mean(model.output{iSR}{4}.CI.d.JFrSO(:,ileak_SR)); % SO from the fully reduced FMN
CIb_FradSO = mean(model.output{iSR}{4}.CI.d.JFradSO(:,ileak_SR)); % SO from the FMN radical
CIIb_ISC3SO = mean(model.output{iSR}{4}.CII.d.JISC3_SO(:,ileak_SR)); % SO from the [3Fe-4S] cluster
CIIb_FradSO = mean(model.output{iSR}{4}.CII.d.JFADH_SO(:,ileak_SR)); % SO from the FMN radical
CIIIb_QpSO = mean(model.output{iSR}{4}.CIII.d.JSO(:,ileak_SR));

% S
CIc_SQSO = mean(model.output{iS}{4}.CI.d.JSQSO(:,ileak_S)); % SO from the semiquinone
CIc_FrSO = mean(model.output{iS}{4}.CI.d.JFrSO(:,ileak_S)); % SO from the fully reduced FMN
CIc_FradSO = mean(model.output{iS}{4}.CI.d.JFradSO(:,ileak_S)); % SO from the FMN radical
CIIc_ISC3SO = mean(model.output{iS}{4}.CII.d.JISC3_SO(:,ileak_S)); % SO from the [3Fe-4S] cluster
CIIc_FradSO = mean(model.output{iS}{4}.CII.d.JFADH_SO(:,ileak_S)); % SO from the FMN radical
CIIIc_QpSO = mean(model.output{iS}{4}.CIII.d.JSO(:,ileak_S));

% P/M/S
CId_SQSO = mean(model.output{iPMS}{4}.CI.d.JSQSO(:,ileak_PMS)); % SO from the semiquinone
CId_FrSO = mean(model.output{iPMS}{4}.CI.d.JFrSO(:,ileak_PMS)); % SO from the fully reduced FMN
CId_FradSO = mean(model.output{iPMS}{4}.CI.d.JFradSO(:,ileak_PMS)); % SO from the FMN radical
CIId_ISC3SO = mean(model.output{iPMS}{4}.CII.d.JISC3_SO(:,ileak_PMS)); % SO from the [3Fe-4S] cluster
CIId_FradSO = mean(model.output{iPMS}{4}.CII.d.JFADH_SO(:,ileak_PMS)); % SO from the FMN radical
CIIId_QpSO = mean(model.output{iPMS}{4}.CIII.d.JSO(:,ileak_PMS));

% Plot
figure
hold on
box on
%P/M
i=1;
bar(1,CIa_SQSO,'FaceColor',c(i,:),'EdgeColor',c(i,:))
bar(5.5,CIa_FrSO,'FaceColor',c(i,:),'EdgeColor',c(i,:))
bar(10,CIa_FradSO,'FaceColor',c(i,:),'EdgeColor',c(i,:))
bar(14,CIIa_ISC3SO,'FaceColor',c(i,:),'EdgeColor',c(i,:))
bar(19,CIIa_FradSO,'FaceColor',c(i,:),'EdgeColor',c(i,:))
bar(23.5,CIIIa_QpSO,'FaceColor',c(i,:),'EdgeColor',c(i,:))
%S/R
i=2;
bar(2,CIb_SQSO,'FaceColor',c(i,:),'EdgeColor',c(i,:))
bar(6.5,CIb_FrSO,'FaceColor',c(i,:),'EdgeColor',c(i,:))
bar(11,CIb_FradSO,'FaceColor',c(i,:),'EdgeColor',c(i,:))
bar(15,CIIb_ISC3SO,'FaceColor',c(i,:),'EdgeColor',c(i,:))
bar(20,CIIb_FradSO,'FaceColor',c(i,:),'EdgeColor',c(i,:))
bar(24.5,CIIIb_QpSO,'FaceColor',c(i,:),'EdgeColor',c(i,:))
%S
i=3;
bar(3,CIc_SQSO,'FaceColor',c(i,:),'EdgeColor',c(i,:))
bar(7.5,CIc_FrSO,'FaceColor',c(i,:),'EdgeColor',c(i,:))
bar(12,CIc_FradSO,'FaceColor',c(i,:),'EdgeColor',c(i,:))
bar(16,CIIc_ISC3SO,'FaceColor',c(i,:),'EdgeColor',c(i,:))
bar(21,CIIc_FradSO,'FaceColor',c(i,:),'EdgeColor',c(i,:))
bar(25.5,CIIIc_QpSO,'FaceColor',c(i,:),'EdgeColor',c(i,:))% P/M/S
i=4;
bar(4,CId_SQSO,'FaceColor',c(i,:),'EdgeColor',c(i,:))
bar(8.5,CId_FrSO,'FaceColor',c(i,:),'EdgeColor',c(i,:))
bar(13,CId_FradSO,'FaceColor',c(i,:),'EdgeColor',c(i,:))
bar(17,CIId_ISC3SO,'FaceColor',c(i,:),'EdgeColor',c(i,:))
bar(22,CIId_FradSO,'FaceColor',c(i,:),'EdgeColor',c(i,:))
bar(26.5,CIIId_QpSO,'FaceColor',c(i,:),'EdgeColor',c(i,:))

% Axes
set(gcf, 'Position', get(0,'Screensize'));
% ylabel({'J_{SO}(nmol/mg/min)'},'color',[0 0 0],'fontsize',60);
xticks([2.5 7 11.5 16 20.5 25 30 34.5])
yticks([0 5 10 15])
xticklabels({'I_Q','FMNH_2','FMNH^{.-}','[3Fe-4S]','FADH^{.-}','Q_p'})
xlim([0 27.5])
ylim([0 17])
set(gca,'FontSize',38,'FontName','Times New Roman')
set(gcf,'color','none'),set(gca,'color','none')

% FIGURE 8A - inset 
% same plot as above except ylim([0 0.8]);yticks([0 0.2 0.4 0.6 0.8]);


% Axes
set(gcf, 'Position', get(0,'Screensize')); % Maximize figure.
% ylabel({'J_{SO}(nmol/mg/min)'},'color',[0 0 0],'fontsize',60);
xticks([2.5 7 11.5 16 20.5 25])
xticklabels({'FMNH^{.-}','[3Fe-4S]','FADH^{.-}','Q_p'})
ytickformat('%,.1f')
xlim([0 18])
ylim([0 0.8])
set(gca,'FontSize',38,'FontName','Times New Roman')
set(gcf,'color','none'),set(gca,'color','none')


% Panel B: OXPHOS
% Time intervals
ioxphos_PM = (model.output{iPM}{1} >= 10 & model.output{iPM}{1} < 10.5)';  
ioxphos_SR = (model.output{iSR}{1} >= 5 & model.output{iSR}{1} < 5.5)';
ioxphos_S = (model.output{iS}{1} >= 5 & model.output{iS}{1} < 5.5)';
ioxphos_PMS = (model.output{iPMS}{1} >= 4 & model.output{iPMS}{1} < 4.5)';

% P/M
CIa_SQSO = mean(model.output{iPM}{4}.CI.d.JSQSO(:,ioxphos_PM)); % SO from the semiquinone
CIa_FrSO = mean(model.output{iPM}{4}.CI.d.JFrSO(:,ioxphos_PM)); % SO from the fully reduced FMN
CIa_FradSO = mean(model.output{iPM}{4}.CI.d.JFradSO(:,ioxphos_PM)); % SO from the FMN radical
CIIa_ISC3SO = mean(model.output{iPM}{4}.CII.d.JISC3_SO(:,ioxphos_PM)); % SO from the [3Fe-4S] cluster
CIIa_FradSO = mean(model.output{iPM}{4}.CII.d.JFADH_SO(:,ioxphos_PM)); % SO from the FMN radical
CIIIa_QpSO = mean(model.output{iPM}{4}.CIII.d.JSO(:,ioxphos_PM));

% S/R
CIb_SQSO = mean(model.output{iSR}{4}.CI.d.JSQSO(:,ioxphos_SR)); % SO from the semiquinone
CIb_FrSO = mean(model.output{iSR}{4}.CI.d.JFrSO(:,ioxphos_SR)); % SO from the fully reduced FMN
CIb_FradSO = mean(model.output{iSR}{4}.CI.d.JFradSO(:,ioxphos_SR)); % SO from the FMN radical
CIIb_ISC3SO = mean(model.output{iSR}{4}.CII.d.JISC3_SO(:,ioxphos_SR)); % SO from the [3Fe-4S] cluster
CIIb_FradSO = mean(model.output{iSR}{4}.CII.d.JFADH_SO(:,ioxphos_SR)); % SO from the FMN radical
CIIIb_QpSO = mean(model.output{iSR}{4}.CIII.d.JSO(:,ioxphos_SR));

% S
CIc_SQSO = mean(model.output{iS}{4}.CI.d.JSQSO(:,ioxphos_S)); % SO from the semiquinone
CIc_FrSO = mean(model.output{iS}{4}.CI.d.JFrSO(:,ioxphos_S)); % SO from the fully reduced FMN
CIc_FradSO = mean(model.output{iS}{4}.CI.d.JFradSO(:,ioxphos_S)); % SO from the FMN radical
CIIc_ISC3SO = mean(model.output{iS}{4}.CII.d.JISC3_SO(:,ioxphos_S)); % SO from the [3Fe-4S] cluster
CIIc_FradSO = mean(model.output{iS}{4}.CII.d.JFADH_SO(:,ioxphos_S)); % SO from the FMN radical
CIIIc_QpSO = mean(model.output{iS}{4}.CIII.d.JSO(:,ioxphos_S));

% P/M/S
CId_SQSO = mean(model.output{iPMS}{4}.CI.d.JSQSO(:,ioxphos_PMS)); % SO from the semiquinone
CId_FrSO = mean(model.output{iPMS}{4}.CI.d.JFrSO(:,ioxphos_PMS)); % SO from the fully reduced FMN
CId_FradSO = mean(model.output{iPMS}{4}.CI.d.JFradSO(:,ioxphos_PMS)); % SO from the FMN radical
CIId_ISC3SO = mean(model.output{iPMS}{4}.CII.d.JISC3_SO(:,ioxphos_PMS)); % SO from the [3Fe-4S] cluster
CIId_FradSO = mean(model.output{iPMS}{4}.CII.d.JFADH_SO(:,ioxphos_PMS)); % SO from the FMN radical
CIIId_QpSO = mean(model.output{iPMS}{4}.CIII.d.JSO(:,ioxphos_PMS));

% Plot
figure
hold on
box on
%P/M
i=1;
bar(1,CIa_SQSO,'FaceColor',c(i,:),'EdgeColor',c(i,:))
bar(5.5,CIa_FrSO,'FaceColor',c(i,:),'EdgeColor',c(i,:))
bar(10,CIa_FradSO,'FaceColor',c(i,:),'EdgeColor',c(i,:))
bar(14,CIIa_ISC3SO,'FaceColor',c(i,:),'EdgeColor',c(i,:))
bar(19,CIIa_FradSO,'FaceColor',c(i,:),'EdgeColor',c(i,:))
bar(23.5,CIIIa_QpSO,'FaceColor',c(i,:),'EdgeColor',c(i,:))
%S/R
i=2;
bar(2,CIb_SQSO,'FaceColor',c(i,:),'EdgeColor',c(i,:))
bar(6.5,CIb_FrSO,'FaceColor',c(i,:),'EdgeColor',c(i,:))
bar(11,CIb_FradSO,'FaceColor',c(i,:),'EdgeColor',c(i,:))
bar(15,CIIb_ISC3SO,'FaceColor',c(i,:),'EdgeColor',c(i,:))
bar(20,CIIb_FradSO,'FaceColor',c(i,:),'EdgeColor',c(i,:))
bar(24.5,CIIIb_QpSO,'FaceColor',c(i,:),'EdgeColor',c(i,:))
%S
i=3;
bar(3,CIc_SQSO,'FaceColor',c(i,:),'EdgeColor',c(i,:))
bar(7.5,CIc_FrSO,'FaceColor',c(i,:),'EdgeColor',c(i,:))
bar(12,CIc_FradSO,'FaceColor',c(i,:),'EdgeColor',c(i,:))
bar(16,CIIc_ISC3SO,'FaceColor',c(i,:),'EdgeColor',c(i,:))
bar(21,CIIc_FradSO,'FaceColor',c(i,:),'EdgeColor',c(i,:))
bar(25.5,CIIIc_QpSO,'FaceColor',c(i,:),'EdgeColor',c(i,:))% P/M/S
i=4;
bar(4,CId_SQSO,'FaceColor',c(i,:),'EdgeColor',c(i,:))
bar(8.5,CId_FrSO,'FaceColor',c(i,:),'EdgeColor',c(i,:))
bar(13,CId_FradSO,'FaceColor',c(i,:),'EdgeColor',c(i,:))
bar(17,CIId_ISC3SO,'FaceColor',c(i,:),'EdgeColor',c(i,:))
bar(22,CIId_FradSO,'FaceColor',c(i,:),'EdgeColor',c(i,:))
bar(26.5,CIIId_QpSO,'FaceColor',c(i,:),'EdgeColor',c(i,:))

% Axes
set(gcf, 'Position', get(0,'Screensize')); 
% ylabel({'J_{SO}(nmol/mg/min)'},'color',[0 0 0],'fontsize',60);
xticks([2.5 7 11.5 16 20.5 25 30 34.5])
xticklabels({'I_Q','FMNH_2','FMNH^{.-}','[3Fe-4S]','FADH^{.-}','Q_p'})
xlim([0 27.5])
ylim([0 3.5])
set(gca,'FontSize',38,'FontName','Times New Roman')
set(gcf,'color','none'),set(gca,'color','none')

%% FIG 9: Site-specific H2O2

% Panel A: FMNH2
% LEAK
CIa_FrH2O2 = mean(model.output{iPM}{4}.CI.d.JFrH2O2(:,ileak_PM)); % SO from the FMN radical
CIb_FrH2O2 = mean(model.output{iSR}{4}.CI.d.JFrH2O2(:,ileak_SR)); % SO from the FMN radical
CIc_FrH2O2 = mean(model.output{iS}{4}.CI.d.JFrH2O2(:,ileak_S)); % SO from the FMN radical
CId_FrH2O2 = mean(model.output{iPMS}{4}.CI.d.JFrH2O2(:,ileak_PMS)); % SO from the FMN radical
% OXPHOS
CIa_FrH2O2o = mean(model.output{iPM}{4}.CI.d.JFrH2O2(1,ioxphos_PM)); 
CIb_FrH2O2o = mean(model.output{iSR}{4}.CI.d.JFrH2O2(1,ioxphos_SR));
CIc_FrH2O2o = mean(model.output{iS}{4}.CI.d.JFrH2O2(1,ioxphos_S)); 
CId_FrH2O2o = mean(model.output{iPMS}{4}.CI.d.JFrH2O2(1,ioxphos_PMS)); 

% Plot
figure
hold on
box on
% FMNH2
i=1;bar(1,CIa_FrH2O2,'FaceColor',c(i,:),'EdgeColor',c(i,:)) % P/M
i=2;bar(2,CIb_FrH2O2,'FaceColor',c(i,:),'EdgeColor',c(i,:)) % S/R
i=3;bar(3,CIc_FrH2O2,'FaceColor',c(i,:),'EdgeColor',c(i,:)) % S
i=4;bar(4,CId_FrH2O2,'FaceColor',c(i,:),'EdgeColor',c(i,:)) % P/M/S
i=1;bar(6,CIa_FrH2O2o,'FaceColor',c(i,:),'EdgeColor',c(i,:)) % P/M
i=2;bar(7,CIb_FrH2O2o,'FaceColor',c(i,:),'EdgeColor',c(i,:)) % S/R
i=3;bar(8,CIc_FrH2O2o,'FaceColor',c(i,:),'EdgeColor',c(i,:)) % S
i=4;bar(9,CId_FrH2O2o,'FaceColor',c(i,:),'EdgeColor',c(i,:)) % P/M/S
% Axes
set(gcf, 'Position', get(0,'Screensize')); % Maximize figure.
% ylabel({'J_{H2O2}(nmol/mg/min)'},'color',[0 0 0],'fontsize',60);
xticks([2.5 7.5])
ylim([0 18])
xticklabels({'Leak','Oxphos'})
set(gca,'FontSize',50,'FontName','Times New Roman')
set(gcf,'color','none'),set(gca,'color','none')
set(gcf, 'Position', get(0,'Screensize')); 


% Panel B: FMNH2
% LEAK
CIIa_FrH2O2 = mean(model.output{iPM}{4}.CII.d.JFADH2_H2O2(:,ileak_PM)); % SO from the fully reduced FMN, INSIGNIFICANT
CIIb_FrH2O2 = mean(model.output{iSR}{4}.CII.d.JFADH2_H2O2(:,ileak_SR)); % SO from the fully reduced FMN 
CIIc_FrH2O2 = mean(model.output{iS}{4}.CII.d.JFADH2_H2O2(:,ileak_S)); % SO from the fully reduced FMN 
CIId_FrH2O2 = mean(model.output{iPMS}{4}.CII.d.JFADH2_H2O2(:,ileak_PMS)); % SO from the fully reduced FMN 
% OXPHOS
CIIa_FrH2O2o = mean(model.output{iPM}{4}.CII.d.JFADH2_H2O2(1,ioxphos_PM)); 
CIIb_FrH2O2o = mean(model.output{iSR}{4}.CII.d.JFADH2_H2O2(1,ioxphos_SR)); 
CIIc_FrH2O2o = mean(model.output{iS}{4}.CII.d.JFADH2_H2O2(1,ioxphos_S)); 
CIId_FrH2O2o = mean(model.output{iPMS}{4}.CII.d.JFADH2_H2O2(1,ioxphos_PMS)); 

% Plot
figure
hold on
box on
% FMNH2
i=1;bar(1,CIIa_FrH2O2,'FaceColor',c(i,:),'EdgeColor',c(i,:)) % P/M
i=2;bar(2,CIIb_FrH2O2,'FaceColor',c(i,:),'EdgeColor',c(i,:)) % S/R
i=3;bar(3,CIIc_FrH2O2,'FaceColor',c(i,:),'EdgeColor',c(i,:)) % S
i=4;bar(4,CIId_FrH2O2,'FaceColor',c(i,:),'EdgeColor',c(i,:)) % P/M/S
i=1;bar(6,CIIa_FrH2O2o,'FaceColor',c(i,:),'EdgeColor',c(i,:)) % P/M
i=2;bar(7,CIIb_FrH2O2o,'FaceColor',c(i,:),'EdgeColor',c(i,:)) % S/R
i=3;bar(8,CIIc_FrH2O2o,'FaceColor',c(i,:),'EdgeColor',c(i,:)) % S
i=4;bar(9,CIId_FrH2O2o,'FaceColor',c(i,:),'EdgeColor',c(i,:)) % P/M/S
xticks([2.5 7])
% ylim([0 2.5])
xticklabels({'Leak','Oxphos'})
% FADH2
xlim([0 10])
ylim([0 1e-3])
set(gca,'FontSize',50,'FontName','Times New Roman')
set(gcf,'color','none'),set(gca,'color','none')
set(gcf, 'Position', get(0,'Screensize')); 

%% FIG 9: Scavenging System Activity
% Panel A: P/M
figure 
hold on
box on
plot(model.output{iPM}{1},model.output{iPM}{4}.H2O2scavenge/p.Vmax_scavening,'color',c(1,:),'linewidth',6)
% Axes 
set(gcf, 'Position', [1 1 1920 1080]); 
ylabel({'Scavenging Capacity'},'color',[0 0 0],'fontsize',60);
xlabel({'Time (min)'},'color',[0 0 0],'fontsize',60);
xlim([0 17])
set(gca,'FontSize',45,'FontName','Times New Roman')
set(gcf,'color','none'),set(gca,'color','none')

% Panel B: S/R
figure 
hold on
box on
plot(model.output{iSR}{1},model.output{iSR}{4}.H2O2scavenge/p.Vmax_scavening,'color',c(2,:),'linewidth',6)
% Axes 
set(gcf, 'Position', [1 1 1920 1080]);
ylabel({'Scavenging Capacity'},'color',[0 0 0],'fontsize',60);
xlabel({'Time (min)'},'color',[0 0 0],'fontsize',60);
xlim([0 8.5])
set(gca,'FontSize',45,'FontName','Times New Roman')
set(gcf,'color','none'),set(gca,'color','none')

% Panel C: S
figure 
hold on
box on
plot(model.output{iS}{1},model.output{iS}{4}.H2O2scavenge/p.Vmax_scavening,'color',c(3,:),'linewidth',6)
% Axes 
set(gcf, 'Position', [1 1 1920 1080]);
ylabel({'Scavenging Capacity'},'color',[0 0 0],'fontsize',60);
xlabel({'Time (min)'},'color',[0 0 0],'fontsize',60);
xlim([0 8.5])
set(gca,'FontSize',45,'FontName','Times New Roman')
set(gcf,'color','none'),set(gca,'color','none')

% Panel D: P/M/S
figure 
hold on
box on
plot(model.output{iPMS}{1},model.output{iPMS}{4}.H2O2scavenge/p.Vmax_scavening,'color',c(4,:),'linewidth',6)
% Axes 
set(gcf, 'Position', [1 1 1920 1080]);
ylabel({'Scavenging Capacity'},'color',[0 0 0],'fontsize',60);
xlabel({'Time (min)'},'color',[0 0 0],'fontsize',60);
xlim([0 6.5])
set(gca,'FontSize',45,'FontName','Times New Roman')
set(gcf,'color','none'),set(gca,'color','none')

%% Supplemental Figure (MAL)

figure
hold on
box on
% Model respiratory profile 
plot(model.output{iMAL}{1},-model.output{iMAL}{4}.dO2*1e7,'color',c(1,:),'linewidth',8) % Model parameters are based on nmol/mg/min, so no further conversion is necessary. 
% Data 
yMAL_leak = [data.MAL.JO2_leak]; % These data were already multiplied by 0.6 to get the correct unit of nmol/mg/min
yMAL_oxphos = [data.MAL.JO2_oxphos];
% Individual data points
plot(5+(0.1-0.2*rand(1,numel(yMAL_leak))),yMAL_leak,'ok','markersize',20,'linewidth',2)
plot(11+(0.1-0.2*rand(1,numel(yMAL_oxphos))),yMAL_oxphos,'ok','markersize',20,'linewidth',2)
% Error bars
errorbar(5,mean(yMAL_leak),std(yMAL_leak),'.','color',[0 0 0],'markersize',1,'linewidth',2)
errorbar(11,mean(yMAL_oxphos),std(yMAL_oxphos),'.','color',[0 0 0],'markersize',1,'linewidth',2)
% Axes 
set(gcf, 'Position', [1 1 1920 1080]);
ylabel({'J_{O_2}(nmol O_2/mg/min)'},'color',[0 0 0],'fontsize',60);
ylim([0 75])
xlim([0 20])
xlabel({'Time (min)'});
set(gca,'FontName','Georgia','FontSize',40);
set(gcf,'color','none'),set(gca,'color','none')

%% FIG 10: JSO and JH2O2 as a f(x) of dPsi, %NADH, and %QH2 

dlgTitle    = 'User Question';
dlgQuestion = 'The 4D plots take quite a while. Do you wish to continue?';
choice = questdlg(dlgQuestion,dlgTitle,'Yes','No', 'No');

if strcmp(choice,'Yes')
    
% Extracting steady-state variables at 5 minutes and  varying dPSi, NADH,
% and QH2 to assess the effects of these variables on SO and H2O2 

% Picking the first SV set at t = 5 min
i = find(model.output{iPM_Fig10}{1}>=5,1); 
x0 = model.output{iPM_Fig10}{2}(i,:)'; % SV structure: rows = time, columns = SVs
x00 = x0([p.i.NADH_x,p.i.UQH2_x,p.i.dPsi]);

% Setting dPsi, NADH, and QH2
dPsi = linspace(0,220,221);
NADH = linspace(0,p.Ntot,101); % M (3 mM)
QH2 = linspace(0,p.Qtot,101); % M (20 mM)
 
x0 = x0*ones(1,numel(dPsi));
% set dPsi in x0
x0(p.i.dPsi,:) = dPsi;

p.rotenone = 0;
p.exp = 3; % flag needed for model

J={};H=[]; % initialize J and H
% Set NADH/NAD and QH2/Q ratios (this takes a while)
f = waitbar(0,'Please wait...');
for i=1:numel(NADH)
for j=1:numel(QH2)
    x0(p.i.NADH_x,:) = NADH(i);
    x0(p.i.NAD_x,:) =  p.Ntot - NADH(i);
    x0(p.i.UQH2_x,:) = QH2(j);
    x0(p.i.UQ_x,:) =  p.Qtot - QH2(j);
    [~,Z] = ETS_ROS_model(0,x0,p);
%     Z=rmfield(Z,{'AK','CK','Inject','FCCP2','FCCP','MCU','NCLX','mNHE','mCHE2','mCHE3','RaM','mKleak','MitoDH_fat'});
    H.CI=Z.CI;
    H.CII=Z.CII;
    H.CIII=Z.CIII;
    J{i,j}=H;
    
    waitbar((j+101*(i-1))/(101*101),f,'Please wait...');
end
end
delete(f)

% build vectors for 3D scatter plot
N = zeros(numel(dPsi),numel(NADH),numel(QH2));
Q = zeros(numel(dPsi),numel(NADH),numel(QH2));
JROS = zeros(numel(dPsi),numel(NADH),numel(QH2));
JCI_SO = zeros(numel(dPsi),numel(NADH),numel(QH2));
JCI_H2O2 = zeros(numel(dPsi),numel(NADH),numel(QH2));
JCII_SO = zeros(numel(dPsi),numel(NADH),numel(QH2));
JCII_H2O2 = zeros(numel(dPsi),numel(NADH),numel(QH2));
JCIII_SO = zeros(numel(dPsi),numel(NADH),numel(QH2));
for i=1:numel(NADH)
for j=1:numel(QH2)
    N(:,i,j) = ones(numel(dPsi),1)*NADH(i);
    Q(:,i,j) = ones(numel(dPsi),1)*QH2(j);
    JROS(:,i,j) = J{i,j}.CI.JSO/2+J{i,j}.CI.JH2O2 + J{i,j}.CII.JSO/2+J{i,j}.CII.JH2O2 + J{i,j}.CIII.JSO/2;
    JCI_SO(:,i,j) = J{i,j}.CI.JSO;
    JCI_H2O2(:,i,j) = J{i,j}.CI.JH2O2;
    JCII_SO(:,i,j) = J{i,j}.CII.JSO;
    JCII_H2O2(:,i,j) = J{i,j}.CII.JH2O2;
    JCIII_SO(:,i,j) = J{i,j}.CIII.JSO;
end
end

% scatter3 plot prep
D=repmat(dPsi',101*201,1);
N=N(:);
Q=Q(:);
JROS=JROS(:);
JCI_SO = JCI_SO(:);
JCI_H2O2 = JCI_H2O2(:);
JCII_SO = JCII_SO(:);
JCII_H2O2 = JCII_H2O2(:);
JCIII_SO = JCIII_SO(:);
JCI_SO(JCI_SO<0)=1e-15; % to preven imag when taking logs
JCI_H2O2(JCI_H2O2<0)=1e-15; % to preven imag when taking logs
JCIII_SO(JCIII_SO<0)=1e-15; % to preven imag when taking logs
JROS(JROS<0)=1e-15;

% size determined by JROS value
Si=100*JROS; % to bound and scale things properly
Si(Si<=0)=1e-3; % set lower limit

% [~,ii]=datasample(N,100000,'Replace',false);
load rng_ii ii % load ii (same indices used in paper for 4D plots)

B=[max(log10(JROS))*1.2-4 max(log10(JROS))*1.2];

% JROS

end

%% Wrapper function to simulate experimental protocols

function model = getoutput(fun,p,fittingfunctions)

%% Initialize model cell array
model.label = {}; % string used to identify model outputs
model.output = {}; % model output


%% Loop through fittingfunctions
for i=1:numel(fittingfunctions)
    model = fittingfunctions{i}(fun,p,model);
end
end

%% Generalized initial conditions
% condition specific initial conditions set in output functions

function x0 = Model_ICs(p)

% Algebraic related SVs adjusted for IC consistency
x0=1e-12*ones(p.i.length,1);

% Non-solute SVs
x0(p.i.dPsi)=0;
% Matrix SVs
x0(p.i.H_x)=1e-7;
x0(p.i.Mg_x) = 0.7e-3; %  Jung et al, 1990; Biochem.
x0(p.i.K_x)=0.13;
x0(p.i.NAD_x)=p.Ntot-1e-12;
x0(p.i.NADH_x)=1e-12;
x0(p.i.ATP_x)=1e-12;
x0(p.i.ADP_x)=p.Atot-1e-12;
x0(p.i.Pi_x)=1e-4;
x0(p.i.UQ_x)=p.Qtot-1e-12;
x0(p.i.UQH2_x)=10^-12;
x0(p.i.O2_x)=20e-6;
% IMS SVs
x0(p.i.c3_x)=p.Ctot-1e-12;
x0(p.i.c2_x)=1e-12;
% Cytosolic SVs
x0(p.i.H_c)=1e-7;
x0(p.i.Mg_c)=1e-9;
x0(p.i.K_c)=0.13;
x0(p.i.ATP_c)=1e-9;
x0(p.i.ADP_c)=1e-9;
x0(p.i.Pi_c)=1e-6;
x0(p.i.O2_c)=200e-9;

end

%% Simulate P/M condition

function model = PM(fun,p,model)
%% Initialize Model Parameters and ICs

xi0 = Model_ICs(p); % general model ICs

%% set model flags

p.exp = 2; % only clamp extra-mitochondrial proton concentration
p.rotenone = 0; % M

%% Set water/mito volume fraction (based on mito load)

p.Vol.H2O_cyt = 1e6/0.1; % nl buffer / mg based on 0.1 mg/ml mitochondria

%% Set extra-mitochonrial buffering parameters

p.cBuff.Htot = 20e-3; % MOPS
p.cBuff.KD_H = 10^-7.2; % MOPS pKa

%% Initialize Integrator Options

% Mass matrix
M=eye(p.i.length);M(p.i.ATP_x,:)=0;M(p.i.NADH_x,:)=0;M(p.i.UQH2_x,:)=0;
M(p.i.c2_x,:)=0;

% define integration absolute tolerances for matrix, ims and cytosol SVs
AbsTol_vec=1e-12.*ones(1,p.i.length);
% set global integrator options
options = odeset('RelTol',1e-3,'AbsTol',AbsTol_vec,'Vectorized','on', ...
    'MassSingular','yes','Mass',M,'MStateDependence','none','BDF','on');

%% Set Model ICs

xi0(p.i.O2_x)=188e-6; % closer to buffer [O2]
xi0(p.i.H_c) = 10^-7.1;
xi0(p.i.Mg_c) = 1e-3;
xi0(p.i.K_c) = 0.13;
xi0(p.i.Na_c) = 20e-3;
xi0(p.i.K_x) = 0.075;
xi0(p.i.Pi_c) = 5e-3;
xi0(p.i.PYR_c) = 5e-3;
xi0(p.i.MAL_c) = 1e-3;

%% Simulate Experiments

ADP = 500e-6; % M
p.ATPase.Etot = 0; % initialize to zero
p.cBuff.EGTA= 1e-3; % M
% initializaiton phase
[t1,x1] = ode15s(fun,[0 5],xi0,options,p);
xi2=x1(end,:);
% calcium uptake and cycling phase(if loading mitos with calcium)
[t2,x2] = ode15s(fun,[0 5],xi2,options,p);
%set ICs for ADP phosphorylation phase
xi3=x2(end,:);
p.ATPase.Etot = 0.05*1e-6/1e-3; % approx (variable dependent on mito prep)
% ADP phosphorylation phase
p.Inject.R(p.if.ADP)=ADP; % define ADP injection parameters
[t3,x3] = ode15s(fun,[0 10],xi3(end,:),options,p);
p.Inject.R(p.if.ADP)=0; % reset to zero
p.ATPase.Etot = 0; % reset back to zero
% compile outputs
t=[t1;t2+t1(end);t3+t1(end)+t2(end)];
x=[x1;x2;x3];
% evaluate LHS and Js
[f, J] = fun(t',x.',p);

%% Set Model Output

% define model output according to data
model.label{numel(model.label)+1} = 'PM';
% set t, x, f and J
model.output{numel(model.output)+1}={t x f J};

end

%% Simulate S/R condition

function model = SR(fun,p,model)
%% Initialize Model Parameters and ICs

xi0 = Model_ICs(p); % general model ICs

%% set model flags

p.exp = 2; % only clamp extra-mitochondrial proton concentration
p.rotenone = 1e-6; % M

%% Set water/mito volume fraction (based on mito load)

p.Vol.H2O_cyt = 1e6/0.1; % nl buffer / mg based on 0.1 mg/ml mitochondria

%% Set extra-mitochonrial buffering parameters

p.cBuff.Htot = 20e-3; % MOPS
p.cBuff.KD_H = 10^-7.2; % MOPS pKa

%% Initialize Integrator Options

% Mass matrix
M=eye(p.i.length);M(p.i.ATP_x,:)=0;M(p.i.NADH_x,:)=0;M(p.i.UQH2_x,:)=0;
M(p.i.c2_x,:)=0;

% define integration absolute tolerances for matrix, ims and cytosol SVs
AbsTol_vec=1e-12.*ones(1,p.i.length);
% set global integrator options
options = odeset('RelTol',1e-3,'AbsTol',AbsTol_vec,'Vectorized','on', ...
    'MassSingular','yes','Mass',M,'MStateDependence','none','BDF','on');

%% Set Model ICs

xi0(p.i.O2_x)=188e-6; % closer to buffer [O2]
xi0(p.i.H_c) = 10^-7.1;
xi0(p.i.Mg_c) = 1e-3;
xi0(p.i.K_c) = 0.13;
xi0(p.i.Na_c) = 20e-3;
xi0(p.i.K_x) = 0.075;
xi0(p.i.Pi_c) = 5e-3;
xi0(p.i.SUC_c) = 10e-3;
xi0(p.i.MAL_c) = 1e-12;

%% Simulate Experiments

ADP = 500e-6; % M
p.ATPase.Etot = 0;
p.cBuff.EGTA= 1e-3; % M
% initializaiton phase
[t1,x1] = ode15s(fun,[0 2.5],xi0,options,p);
xi2=x1(end,:);
% calcium uptake phase(if loading with calcium)
[t2,x2] = ode15s(fun,[0 2.5],xi2,options,p);
%set ICs for ADP phosphorylation phase
xi3=x2(end,:);
p.ATPase.Etot = 0.05*1e-6/1e-3; % approx (variable dependent on mito prep)
p.Inject.R(p.if.Ca) = 0;
% ADP phosphorylation phase
p.Inject.R(p.if.ADP)=ADP;
[t3,x3] = ode15s(fun,[0 5],xi3(end,:),options,p);
p.Inject.R(p.if.ADP)=0;
p.ATPase.Etot = 0;
% compile outputs
t=[t1;t2+t1(end);t3+t1(end)+t2(end)];
x=[x1;x2;x3];
[f, J] = fun(t',x.',p);

%% Set Model Output

% define model output according to data
model.label{numel(model.label)+1} = 'SR';
% set t, x, f and J
model.output{numel(model.output)+1}={t x f J};

end

%% Simulate S condition

function model = S(fun,p,model)
%% Initialize Model Parameters and ICs

xi0 = Model_ICs(p); % general model ICs

%% set model flags

p.exp = 2; % only clamp extra-mitochondrial proton concentration
p.rotenone = 0; % M

%% Set water/mito volume fraction (based on mito load)

p.Vol.H2O_cyt = 1e6/0.1; % nl buffer / mg based on 0.1 mg/ml mitochondria

%% Set extra-mitochonrial buffering parameters

p.cBuff.Htot = 20e-3; % MOPS
p.cBuff.KD_H = 10^-7.2; % MOPS pKa

%% Initialize Integrator Options

% Mass matrix
M=eye(p.i.length);M(p.i.ATP_x,:)=0;M(p.i.NADH_x,:)=0;M(p.i.UQH2_x,:)=0;
M(p.i.c2_x,:)=0;

% define integration absolute tolerances for matrix, ims and cytosol SVs
AbsTol_vec=1e-12.*ones(1,p.i.length);
% set global integrator options
options = odeset('RelTol',1e-3,'AbsTol',AbsTol_vec,'Vectorized','on', ...
    'MassSingular','yes','Mass',M,'MStateDependence','none','BDF','on');

%% Set Model ICs

xi0(p.i.O2_x)=188e-6; % closer to buffer [O2]
xi0(p.i.H_c) = 10^-7.1;
xi0(p.i.Mg_c) = 1e-3;
xi0(p.i.K_c) = 0.13;
xi0(p.i.Na_c) = 20e-3;
xi0(p.i.K_x) = 0.075;
xi0(p.i.Pi_c) = 5e-3;
xi0(p.i.SUC_c) = 10e-3;
xi0(p.i.MAL_c) = 1e-12;

%% Simulate Experiments

ADP = 500e-6; % M
p.ATPase.Etot = 0;
p.cBuff.EGTA= 1e-3; % M
% initializaiton phase
[t1,x1] = ode15s(fun,[0 2.5],xi0,options,p);
xi2=x1(end,:);
% calcium uptake phase(if loading with calcium)
[t2,x2] = ode15s(fun,[0 2.5],xi2,options,p);
%set ICs for ADP phosphorylation phase
xi3=x2(end,:);
p.ATPase.Etot = 0.05*1e-6/1e-3; % approx (variable dependent on mito prep)
p.Inject.R(p.if.Ca) = 0;
% ADP phosphorylation phase
p.Inject.R(p.if.ADP)=ADP;
[t3,x3] = ode15s(fun,[0 5],xi3(end,:),options,p);
p.Inject.R(p.if.ADP)=0;
p.ATPase.Etot = 0;
% compile outputs
t=[t1;t2+t1(end);t3+t1(end)+t2(end)];
x=[x1;x2;x3];
[f, J] = fun(t',x.',p);

%% Set Model Output

% define model output according to data
model.label{numel(model.label)+1} = 'S';
% set t, x, f and J
model.output{numel(model.output)+1}={t x f J};

end

%% Simulate PMS condition

function model = PMS(fun,p,model)
%% Initialize Model Parameters and ICs

xi0 = Model_ICs(p); % general model ICs

%% set model flags

p.exp = 2; % only clamp extra-mitochondrial proton concentration
p.rotenone = 0; % M

%% Set water/mito volume fraction (based on mito load)

p.Vol.H2O_cyt = 1e6/0.1; % nl buffer / mg based on 0.1 mg/ml mitochondria

%% Set extra-mitochonrial buffering parameters

p.cBuff.Htot = 20e-3; % MOPS
p.cBuff.KD_H = 10^-7.2; % MOPS pKa

%% Initialize Integrator Options

% Mass matrix
M=eye(p.i.length);M(p.i.ATP_x,:)=0;M(p.i.NADH_x,:)=0;M(p.i.UQH2_x,:)=0;
M(p.i.c2_x,:)=0;

% define integration absolute tolerances for matrix, ims and cytosol SVs
AbsTol_vec=1e-12.*ones(1,p.i.length);
% set global integrator options
options = odeset('RelTol',1e-3,'AbsTol',AbsTol_vec,'Vectorized','on', ...
    'MassSingular','yes','Mass',M,'MStateDependence','none','BDF','on');

%% Set Model ICs

xi0(p.i.O2_x)=188e-6; % closer to buffer [O2]
xi0(p.i.H_c) = 10^-7.1;
xi0(p.i.Mg_c) = 1e-3;
xi0(p.i.K_c) = 0.13;
xi0(p.i.Na_c) = 20e-3;
xi0(p.i.K_x) = 0.075;
xi0(p.i.Pi_c) = 5e-3;
xi0(p.i.PYR_c) = 5e-3;
xi0(p.i.MAL_c) = 1e-3;
xi0(p.i.SUC_c) = 10e-3;

%% Simulate Experiments

ADP = 500e-6; % M
p.ATPase.Etot = 0;
p.cBuff.EGTA= 1e-3; % M
% initializaiton phase
[t1,x1] = ode15s(fun,[0 2],xi0,options,p);
xi2=x1(end,:);
% calcium uptake phase(if loading with calcium)
[t2,x2] = ode15s(fun,[0 2],xi2,options,p);
%set ICs for ADP phosphorylation phase
xi3=x2(end,:);
p.ATPase.Etot = 0.05*1e-6/1e-3; % approx (variable dependent on mito prep)
p.Inject.R(p.if.Ca) = 0;
% ADP phosphorylation phase
p.Inject.R(p.if.ADP)=ADP;
[t3,x3] = ode15s(fun,[0 4],xi3(end,:),options,p);
p.Inject.R(p.if.ADP)=0;
p.ATPase.Etot = 0;
% compile outputs
t=[t1;t2+t1(end);t3+t1(end)+t2(end)];
x=[x1;x2;x3];
[f, J] = fun(t',x.',p);

%% Set Model Output

% define model output according to data
model.label{numel(model.label)+1} = 'PMS';
% set t, x, f and J
model.output{numel(model.output)+1}={t x f J};

end

%% Simulate P/M condition for various O2 concentrations

function model = PM_O2(fun,p,model)
%% Initialize Model Parameters and ICs

xi0 = Model_ICs(p); % general model ICs

%% set model flags

p.exp = 2; % only clamp extra-mitochondrial proton concentration
p.rotenone = 0; % M

%% Set water/mito volume fraction (based on mito load)

p.Vol.H2O_cyt = 1e6/0.1; % nl buffer / mg based on 0.1 mg/ml mitochondria

%% Set extra-mitochonrial buffering parameters

p.cBuff.Htot = 20e-3; % MOPS
p.cBuff.KD_H = 10^-7.2; % MOPS pKa

%% Initialize Integrator Options

% Mass matrix
M=eye(p.i.length);M(p.i.ATP_x,:)=0;M(p.i.NADH_x,:)=0;M(p.i.UQH2_x,:)=0;
M(p.i.c2_x,:)=0;

% define integration absolute tolerances for matrix, ims and cytosol SVs
AbsTol_vec=1e-12.*ones(1,p.i.length);
% set global integrator options
options = odeset('RelTol',1e-3,'AbsTol',AbsTol_vec,'Vectorized','on', ...
    'MassSingular','yes','Mass',M,'MStateDependence','none','BDF','on');

%% Set Model ICs

p.cBuff.EGTA= 1e-3; % M
xi0(p.i.O2_x)=188e-6; % closer to buffer [O2]
xi0(p.i.H_c) = 10^-7.1;
xi0(p.i.Mg_c) = 1e-3;
xi0(p.i.K_c) = 0.13;
xi0(p.i.Na_c) = 20e-3;
xi0(p.i.K_x) = 0.075;
xi0(p.i.Pi_c) = 5e-3;
xi0(p.i.PYR_c) = 5e-3;
xi0(p.i.MAL_c) = 1e-3;

%% Simulate model to anoxia

% initializaiton phase
[t,x] = ode15s(fun,[0 90],xi0,options,p);
[f, J] = fun(t',x.',p);

%% Set Model Output

% define model output according to data
model.label{numel(model.label)+1} = 'PM_O2';
% set t, x, f and J
model.output{numel(model.output)+1}={t x f J};

end

%% Simulate S/R condition for various O2 concentrations

function model = SRot_O2(fun,p,model)
%% Initialize Model Parameters and ICs

xi0 = Model_ICs(p); % general model ICs

%% set model flags

p.exp = 2; % only clamp extra-mitochondrial proton concentration
p.rotenone = 1e-6; % M

%% Set water/mito volume fraction (based on mito load)

p.Vol.H2O_cyt = 1e6/0.1; % nl buffer / mg based on 0.1 mg/ml mitochondria

%% Set extra-mitochonrial buffering parameters

p.cBuff.Htot = 20e-3; % MOPS
p.cBuff.KD_H = 10^-7.2; % MOPS pKa

%% Initialize Integrator Options

% Mass matrix
M=eye(p.i.length);M(p.i.ATP_x,:)=0;M(p.i.NADH_x,:)=0;M(p.i.UQH2_x,:)=0;
M(p.i.c2_x,:)=0;

% define integration absolute tolerances for matrix, ims and cytosol SVs
AbsTol_vec=1e-12.*ones(1,p.i.length);
% set global integrator options
options = odeset('RelTol',1e-3,'AbsTol',AbsTol_vec,'Vectorized','on', ...
    'MassSingular','yes','Mass',M,'MStateDependence','none','BDF','on');

%% Set Model ICs

p.cBuff.EGTA= 1e-3; % M
xi0(p.i.O2_x)=188e-6; % closer to buffer [O2]
xi0(p.i.H_c) = 10^-7.1;
xi0(p.i.Mg_c) = 1e-3;
xi0(p.i.K_c) = 0.13;
xi0(p.i.Na_c) = 20e-3;
xi0(p.i.K_x) = 0.075;
xi0(p.i.Pi_c) = 5e-3;
xi0(p.i.SUC_c) = 10e-3;
xi0(p.i.MAL_c) = 1e-12;

%% Simulate model to anoxia

% initialization phase
[t,x] = ode15s(fun,[0 40],xi0,options,p);
[f, J] = fun(t',x.',p);

%% Set Model Output

% define model output according to data
model.label{numel(model.label)+1} = 'SR_O2';
% set t, x, f and J
model.output{numel(model.output)+1}={t x f J};

end

%% Simulate M conditon

function model = MAL(fun,p,model)
%% Initialize Model Parameters and ICs
xi0 = Model_ICs(p); % general model ICs (MAKE GEN AND ADD BOSE ICS)

p.exp = 2; % only clamp extra-mitochondrial proton concentration
p.rotenone = 0; % M

%% Set water/mito volume fraction (based on mito load)
p.Vol.H2O_cyt = 1e6/0.1; % nl buffer / mg based on 0.1 mg/ml mitochondria

p.cBuff.Htot = 20e-3; % MOPS
p.cBuff.KD_H = 10^-7.2; % MOPS pKa

%% Initialize Integrator Options

% Mass matrix
M=eye(p.i.length);
M(p.i.ATP_x,:)=0;
M(p.i.NADH_x,:)=0;
M(p.i.UQH2_x,:)=0;
M(p.i.c2_x,:)=0;

% define integration absolute tolerances for matrix, ims and cytosol SVs
AbsTol_vec=1e-9.*ones(1,p.i.length);
% set global integrator options
options = odeset('RelTol',1e-3,'AbsTol',AbsTol_vec,'Vectorized','on', ...
    'MassSingular','yes','Mass',M,'MStateDependence','none','BDF','on');

%% Set Model ICs
xi0(p.i.O2_x)=188e-6; % closer to buffer [O2]
xi0(p.i.H_c) = 10^-7.1;
xi0(p.i.dPsi) = 0;
xi0(p.i.Mg_c) = 1e-3;
xi0(p.i.ATP_c) = 1e-16;
xi0(p.i.K_c) = 0.13;
xi0(p.i.Na_c) = 20e-3;
xi0(p.i.K_x) = 0.075;
xi0(p.i.Pi_c) = 5e-3;
xi0(p.i.PYR_c) = 0e-3;
xi0(p.i.MAL_c) = 1e-3;

%% Simulate Experiments
ADP = 500e-6; % M
p.ATPase.Etot = 0;
p.cBuff.EGTA= 1e-3; % M
% initializaiton phase
[t1,x1] = ode15s(fun,[0 5],xi0,options,p);
xi2=x1(end,:);
% calcium uptake phase(if loading with calcium)
[t2,x2] = ode15s(fun,[0 5],xi2,options,p);
%set ICs for ADP phosphorylation phase
xi3=x2(end,:);
p.ATPase.Etot = 0.05*1e-6/1e-3; % approx (variable dependent on mito prep)
p.Inject.R(p.if.Ca) = 0;
% ADP phosphorylation phase
p.Inject.R(p.if.ADP)=ADP;
[t3,x3] = ode15s(fun,[0 10],xi3(end,:),options,p);
p.Inject.R(p.if.ADP)=0;
p.ATPase.Etot = 0;
% compile outputs
t=[t1;t2+t1(end);t3+t1(end)+t2(end)];
x=[x1;x2;x3];
[f, J] = fun(t',x.',p);

%% Set Model Output

% define model output according to data
model.label{numel(model.label)+1} = 'MAL';
% set t, x, f and J
model.output{numel(model.output)+1}={t x f J};

end

%% Simulat P/M condition (with fixed PYR, MAL, and O2)

function model = PM_Fig10(fun,p,model)

%% Initialize Model Parameters and ICs
xi0 = Model_ICs(p); % general model ICs (MAKE GEN AND ADD BOSE ICS)

p.exp = 3; % keep substrates saturating, fix pH, and fix O2
p.rotenone = 0; % M

%% Set water/mito volume fraction (based on mito load)
p.Vol.H2O_cyt = 1e6/0.1; % nl buffer / mg based on 0.1 mg/ml mitochondria

p.cBuff.Htot = 20e-3; % MOPS
p.cBuff.KD_H = 10^-7.2; % MOPS pKa

%% Initialize Integrator Options

% Mass matrix
M=eye(p.i.length);
M(p.i.ATP_x,:)=0;
M(p.i.NADH_x,:)=0;
M(p.i.UQH2_x,:)=0;
M(p.i.c2_x,:)=0;

% define integration absolute tolerances for matrix, ims and cytosol SVs
AbsTol_vec=1e-12.*ones(1,p.i.length);
% set global integrator options
options = odeset('RelTol',1e-3,'AbsTol',AbsTol_vec,'Vectorized','on', ...
    'MassSingular','yes','Mass',M,'MStateDependence','none','BDF','on');

%% Set Model ICs
xi0(p.i.O2_x) = 20e-6; % more physiological O2 concentration in heart
xi0(p.i.H_c) = 10^-7.1;
xi0(p.i.dPsi) = 0;
xi0(p.i.Mg_c) = 1e-3;
xi0(p.i.ATP_c) = 1e-16;
xi0(p.i.K_c) = 0.13;
xi0(p.i.Na_c) = 20e-3;
xi0(p.i.K_x) = 0.075;
xi0(p.i.Pi_c) = 5e-3;
xi0(p.i.PYR_c) = 5e-3;
xi0(p.i.MAL_c) = 1e-3;

%% Simulate Experiments
ADP = 500e-6; % M
p.ATPase.Etot = 0;
p.cBuff.EGTA= 1e-3; % M
% initializaiton phase
[t1,x1] = ode15s(fun,[0 5],xi0,options,p);
xi2=x1(end,:);
% calcium uptake phase(if loading with calcium)
[t2,x2] = ode15s(fun,[0 5],xi2,options,p);
%set ICs for ADP phosphorylation phase
xi3=x2(end,:);
p.ATPase.Etot = 0.05*1e-6/1e-3; % approx (variable dependent on mito prep)
p.Inject.R(p.if.Ca) = 0;
% ADP phosphorylation phase
p.Inject.R(p.if.ADP)=ADP;
[t3,x3] = ode15s(fun,[0 10],xi3(end,:),options,p);
p.Inject.R(p.if.ADP)=0;
p.ATPase.Etot = 0;
% compile outputs
t=[t1;t2+t1(end);t3+t1(end)+t2(end)];
x=[x1;x2;x3];
[f, J] = fun(t',x.',p);

%% Set Model Output

% define model output according to data
model.label{numel(model.label)+1} = 'PM_Fig10';
% set t, x, f and J
model.output{numel(model.output)+1}={t x f J};

end

function create4Dfigure(X1, Y1, Z1, S1, C1, B, x00)
%CREATEFIGURE(X1, Y1, Z1, S1, C1)
%  X1:  scatter3 x
%  Y1:  scatter3 y
%  Z1:  scatter3 z
%  S1:  scatter3 s
%  C1:  scatter3 c

%  Auto-generated by MATLAB on 14-May-2021 21:07:54

% Create figure
figure1 = figure('Color','none');

% Create axes
axes1 = axes('Parent',figure1);
hold(axes1,'on');

% Create scatter3
scatter3(X1,Y1,Z1,S1,C1,'MarkerFaceAlpha',0.25,'MarkerFaceColor','flat',...
    'MarkerEdgeAlpha',0,...
    'MarkerEdgeColor','none');

% Create zlabel
zlabel('\Delta\Psi (mV)','FontName','Times New Roman','Color',[1 1 1]);

% Create ylabel
ylabel('%QH_2','FontName','Times New Roman','Color',[1 1 1]);

% Create xlabel
xlabel('%NADH','FontName','Times New Roman','Color',[1 1 1]);
zlim([100 220])
view(axes1,[150 33]);
grid(axes1,'on');
hold(axes1,'off');
% Set the remaining axes properties
set(axes1,'CLim',[B(1) B(2)],'Color','none','Colormap',...
    [0.244160786271095 0.152360782027245 0.662260830402374;0.245987892150879 0.155552506446838 0.671863496303558;0.247776344418526 0.15877303481102 0.681466221809387;0.249526545405388 0.162022367119789 0.691068947315216;0.251238942146301 0.165300488471985 0.700671672821045;0.252913951873779 0.168607398867607 0.710274338722229;0.254551976919174 0.171943113207817 0.719877064228058;0.256153464317322 0.175307631492615 0.729479789733887;0.257718801498413 0.178700938820839 0.739082515239716;0.259248435497284 0.182123050093651 0.7486851811409;0.260742783546448 0.185573950409889 0.758287906646729;0.262202262878418 0.189053654670715 0.767890632152557;0.26362732052803 0.192562147974968 0.777493357658386;0.257618427276611 0.191558972001076 0.778151512145996;0.251578241586685 0.190553814172745 0.778809666633606;0.245506763458252 0.189546689391136 0.779467880725861;0.239403903484344 0.188537582755089 0.78012603521347;0.233269616961479 0.187526494264603 0.780784249305725;0.227103844285011 0.186513438820839 0.781442403793335;0.220906555652618 0.185498401522636 0.78210061788559;0.214677661657333 0.184481397271156 0.782758772373199;0.208417147397995 0.183462411165237 0.783416986465454;0.202124938368797 0.182441458106041 0.784075140953064;0.195800974965096 0.181418523192406 0.784733355045319;0.189445227384567 0.180393606424332 0.785391509532928;0.183057636022568 0.17936672270298 0.786049723625183;0.17833785712719 0.180037572979927 0.786707878112793;0.177307024598122 0.184427350759506 0.787366092205048;0.176274210214615 0.188845172524452 0.788024246692657;0.175239413976669 0.193291112780571 0.788682460784912;0.174202650785446 0.197765201330185 0.789340615272522;0.173163920640945 0.2022675126791 0.789998829364777;0.172123193740845 0.206798076629639 0.790656983852386;0.171080499887466 0.211356967687607 0.791315197944641;0.170035839080811 0.215944215655327 0.791973352432251;0.168989196419716 0.220559895038605 0.792631566524506;0.167940571904182 0.225204050540924 0.793289721012115;0.166889980435371 0.229876726865768 0.79394793510437;0.165837407112122 0.234577983617783 0.79460608959198;0.164782866835594 0.239307880401611 0.795264303684235;0.163726344704628 0.244066461920738 0.795922458171844;0.162667840719223 0.248853772878647 0.796580672264099;0.16160736978054 0.253669887781143 0.797238826751709;0.16054493188858 0.258514821529388 0.797897040843964;0.15948049724102 0.263388693332672 0.798555195331573;0.158414095640182 0.268291473388672 0.799213409423828;0.157345727086067 0.273223280906677 0.799871563911438;0.156275376677513 0.278184115886688 0.800529778003693;0.15520304441452 0.283174067735672 0.801187932491302;0.15412874519825 0.288193196058273 0.801846146583557;0.153052464127541 0.293241530656815 0.802504301071167;0.151974216103554 0.298319101333618 0.803162515163422;0.150893986225128 0.303426027297974 0.803820669651031;0.149811774492264 0.308562308549881 0.804478883743286;0.148727595806122 0.313728004693985 0.805137038230896;0.147641435265541 0.31892317533493 0.805795252323151;0.146553307771683 0.324147880077362 0.80645340681076;0.145463198423386 0.329402148723602 0.807111620903015;0.144371122121811 0.334686070680618 0.807769775390625;0.143277063965797 0.339999675750732 0.80842798948288;0.142181023955345 0.345343023538589 0.80908614397049;0.141083016991615 0.350716143846512 0.809744358062744;0.139983028173447 0.356119096279144 0.810402512550354;0.138881072402 0.361551970243454 0.811060726642609;0.137777134776115 0.367014795541763 0.811718881130219;0.136671215295792 0.372507601976395 0.812377035617828;0.13556332886219 0.378030478954315 0.813035249710083;0.13445346057415 0.383583456277847 0.813693404197693;0.133341625332832 0.389166593551636 0.814351618289948;0.132227808237076 0.394779920578003 0.815009772777557;0.13111200928688 0.400423526763916 0.815667986869812;0.129994243383408 0.40609747171402 0.816326141357422;0.128874510526657 0.411801755428314 0.816984355449677;0.127752795815468 0.417536467313766 0.817642509937286;0.12662909924984 0.423301666975021 0.818300724029541;0.125503420829773 0.429097384214401 0.818958878517151;0.124375782907009 0.434923678636551 0.819617092609406;0.123246155679226 0.440780609846115 0.820275247097015;0.122114561498165 0.446668237447739 0.82093346118927;0.120980992913246 0.452586591243744 0.82159161567688;0.119845442473888 0.458535730838776 0.822249829769135;0.118707917630672 0.464515715837479 0.822907984256744;0.117568418383598 0.470526576042175 0.823566198348999;0.116426944732666 0.476568430662155 0.824224352836609;0.115283496677876 0.482641249895096 0.824882566928864;0.114138066768646 0.488745123147964 0.825540721416473;0.112990662455559 0.494880110025406 0.826198935508728;0.111841291189194 0.501046240329742 0.826857089996338;0.11068993806839 0.507243633270264 0.827515304088593;0.109536603093147 0.513472259044647 0.828173458576202;0.108381301164627 0.519732177257538 0.828831672668457;0.107224024832249 0.526023507118225 0.829489827156067;0.106064766645432 0.532346248626709 0.830148041248322;0.104903534054756 0.53870040178299 0.830806195735931;0.103740327060223 0.545086145401001 0.831464409828186;0.102575145661831 0.551503479480743 0.832122564315796;0.101407989859581 0.557952404022217 0.832780778408051;0.100238852202892 0.564433038234711 0.83343893289566;0.0990677401423454 0.570945382118225 0.834097146987915;0.0978946536779404 0.57748955488205 0.834755301475525;0.0967195928096771 0.584065556526184 0.83541351556778;0.0955425575375557 0.590673446655273 0.836071670055389;0.0943635478615761 0.597313284873962 0.836729884147644;0.0931825563311577 0.603985130786896 0.837388038635254;0.0919995978474617 0.610688984394073 0.838046252727509;0.0908146575093269 0.61742502450943 0.838704407215118;0.089627742767334 0.624193131923676 0.839362621307373;0.0884388461709023 0.630993545055389 0.840020775794983;0.0872479826211929 0.637826144695282 0.840678989887238;0.0860551446676254 0.644691109657288 0.841337144374847;0.0848603248596191 0.651588380336761 0.841995358467102;0.0836635306477547 0.658518135547638 0.842653512954712;0.082464762032032 0.665480375289917 0.843311727046967;0.0812640190124512 0.672475099563599 0.843969881534576;0.0800612941384315 0.679502427577972 0.844628095626831;0.0788566023111343 0.686562359333038 0.845286250114441;0.0776499286293983 0.693655014038086 0.845944464206696;0.0674284473061562 0.696953058242798 0.839918613433838;0.0573461949825287 0.700385451316833 0.83389276266098;0.0474031828343868 0.703946530818939 0.827866971492767;0.03759940341115 0.707630932331085 0.82184112071991;0.0279348604381084 0.711433112621307 0.815815329551697;0.0192775689065456 0.714674592018127 0.809512555599213;0.0123623115941882 0.717817604541779 0.80309921503067;0.00738342106342316 0.720885097980499 0.79657518863678;0.0044031604193151 0.723868131637573 0.789965927600861;0.00339387194253504 0.726792454719543 0.783326148986816;0.00443505588918924 0.729632675647736 0.776654601097107;0.00761206354945898 0.73239654302597 0.769897103309631;0.012982465326786 0.735102415084839 0.763065934181213;0.0206828378140926 0.737715721130371 0.756176829338074;0.0309630744159222 0.740260481834412 0.749233722686768;0.0328734405338764 0.749926388263702 0.735706925392151;0.03478729352355 0.750619113445282 0.713285744190216;0.036704633384943 0.751311838626862 0.690943837165833;0.0386254638433456 0.752004563808441 0.668681561946869;0.0405497848987579 0.752697288990021 0.646499156951904;0.0424775928258896 0.7533900141716 0.624396979808807;0.0444088876247406 0.75408273935318 0.602375388145447;0.0463436767458916 0.75477546453476 0.580434739589691;0.0482819490134716 0.755468189716339 0.558575391769409;0.0502237118780613 0.756160855293274 0.536797642707825;0.0521689653396606 0.756853580474854 0.515101790428162;0.0541177056729794 0.757546305656433 0.493488252162933;0.0560699328780174 0.758239030838013 0.471957325935364;0.0580256506800652 0.758931756019592 0.450509369373322;0.0599848590791225 0.759624481201172 0.429144710302353;0.0619475543498993 0.760317206382751 0.407863676548004;0.0639137402176857 0.761009931564331 0.386666595935822;0.0658834129571915 0.761702656745911 0.365553855895996;0.0678565725684166 0.762395322322845 0.344525754451752;0.0698332190513611 0.763088047504425 0.323582649230957;0.0718133598566055 0.763780772686005 0.302724868059158;0.0737969875335693 0.764473497867584 0.281952738761902;0.0757841020822525 0.765166223049164 0.261266618967056;0.0777747109532356 0.765858948230743 0.240666851401329;0.0797688066959381 0.766551673412323 0.220153763890266;0.08176638931036 0.767244398593903 0.199727684259415;0.0837674587965012 0.767937123775482 0.179388955235481;0.0857720151543617 0.768629789352417 0.159137934446335;0.0877800658345222 0.769322514533997 0.13897493481636;0.0897916033864021 0.770015239715576 0.118900321424007;0.0918066278100014 0.770707964897156 0.0989144071936607;0.108632750809193 0.771400690078735 0.0938251465559006;0.132484242320061 0.772093415260315 0.0958471447229385;0.156252980232239 0.772786140441895 0.0978726372122765;0.1799386292696 0.773478865623474 0.0999016240239143;0.203540861606598 0.774171590805054 0.101934090256691;0.227059334516525 0.774864256381989 0.103970050811768;0.250493705272675 0.775556981563568 0.106009498238564;0.273843616247177 0.776249706745148 0.108052432537079;0.297108769416809 0.776942431926727 0.110098853707314;0.320288836956024 0.777635157108307 0.112148769199848;0.343383401632309 0.778327882289886 0.114202171564102;0.366392195224762 0.779020607471466 0.116259060800076;0.389314860105515 0.779713332653046 0.118319444358349;0.41215106844902 0.780406057834625 0.120383314788342;0.434900462627411 0.78109872341156 0.122450664639473;0.457562685012817 0.78179144859314 0.124521516263485;0.480137467384338 0.782484173774719 0.126595854759216;0.50262439250946 0.783176898956299 0.128673672676086;0.52502316236496 0.783869624137878 0.130754977464676;0.547333478927612 0.784562349319458 0.132839784026146;0.569554924964905 0.785255074501038 0.134928077459335;0.591687202453613 0.785947799682617 0.137019857764244;0.613729953765869 0.786640524864197 0.139115124940872;0.635682821273804 0.787333190441132 0.14121387898922;0.657545566558838 0.788025915622711 0.143316119909286;0.679317772388458 0.788718640804291 0.145421847701073;0.700999081134796 0.78941136598587 0.147531077265739;0.722589194774628 0.79010409116745 0.149643778800964;0.744087755680084 0.79079681634903 0.15175998210907;0.765494465827942 0.791489541530609 0.153879672288895;0.786808967590332 0.792182266712189 0.156002849340439;0.792874991893768 0.777719080448151 0.158129513263702;0.793567717075348 0.757975518703461 0.160259664058685;0.805684030056 0.755361974239349 0.157389789819717;0.81753271818161 0.752796590328217 0.155962035059929;0.827666401863098 0.752123892307281 0.160788252949715;0.837800085544586 0.751296699047089 0.165685966610909;0.847933769226074 0.750318288803101 0.17065517604351;0.858067452907562 0.74919193983078 0.175695866346359;0.86820113658905 0.747920989990234 0.180808067321777;0.878334820270538 0.746508717536926 0.185991749167442;0.888468503952026 0.744958400726318 0.191246941685677;0.898602187633514 0.743273437023163 0.196573615074158;0.908735871315002 0.741457104682922 0.201971784234047;0.91886955499649 0.73951268196106 0.207441449165344;0.929003238677979 0.737443506717682 0.21298260986805;0.939136922359467 0.735252916812897 0.218595266342163;0.949270606040955 0.732944130897522 0.224279403686523;0.959404289722443 0.73052054643631 0.230035051703453;0.968122959136963 0.731046497821808 0.236939057707787;0.976598262786865 0.732291340827942 0.242168441414833;0.984302461147308 0.734619498252869 0.245080649852753;0.990729451179504 0.738054275512695 0.244619235396385;0.995505750179291 0.7423996925354 0.241526961326599;0.998070657253265 0.747583270072937 0.23729906976223;0.997749328613281 0.752790808677673 0.233176067471504;0.997427999973297 0.758066654205322 0.229055657982826;0.997106671333313 0.763410627841949 0.224937871098518;0.996785402297974 0.768822729587555 0.220822706818581;0.99646407365799 0.774302899837494 0.216710135340691;0.996142745018005 0.779851019382477 0.212600186467171;0.995821475982666 0.785466969013214 0.20849284529686;0.995500147342682 0.791150808334351 0.204388126730919;0.995178818702698 0.796902418136597 0.200286015868187;0.994857490062714 0.802721679210663 0.196186512708664;0.994536221027374 0.808608591556549 0.19208961725235;0.99421489238739 0.814563035964966 0.187995329499245;0.993893563747406 0.820584952831268 0.18390366435051;0.993572294712067 0.826674282550812 0.179814606904984;0.993250966072083 0.832830965518951 0.175728172063828;0.992929637432098 0.839054942131042 0.171644330024719;0.992608308792114 0.845346093177795 0.167563110589981;0.992287039756775 0.851704359054565 0.163484498858452;0.991965711116791 0.858129680156708 0.159408509731293;0.991644382476807 0.864622056484222 0.155335113406181;0.991323113441467 0.871181309223175 0.15126433968544;0.991001784801483 0.877807438373566 0.147196188569069;0.990680456161499 0.884500324726105 0.143130630254745;0.990359127521515 0.891259968280792 0.139067694544792;0.990037858486176 0.898086190223694 0.135007366538048;0.989716529846191 0.904979050159454 0.130949646234512;0.989395201206207 0.911938369274139 0.126894548535347;0.989073932170868 0.918964207172394 0.122842058539391;0.988752603530884 0.926056325435638 0.118792176246643;0.9884312748909 0.933214783668518 0.114744909107685;0.988109946250916 0.940439462661743 0.110700249671936;0.987788677215576 0.947730302810669 0.106658205389977;0.987467348575592 0.955087244510651 0.102618768811226;0.987146019935608 0.962510168552399 0.0985819473862648;0.986824750900269 0.969999074935913 0.0945477411150932;0.986503422260284 0.977553904056549 0.0905161425471306;0.9861820936203 0.985174477100372 0.0864871591329575;0.978860795497894 0.985860824584961 0.0824607834219933],...
    'FontName','Times New Roman','FontSize',45,'XColor',[1 1 1],'YColor',...
    [1 1 1],'ZColor',[1 1 1]);
% Create colorbar
colorbar(axes1,'Position',...
    [0.938909722222221 0.137508983529538 0.0111111111111111 0.787491016470461],...
    'LineWidth',1,...
    'Color',[1 1 1]);

hold on
scatter3(100*x00(1)/3e-3,100*x00(2)/20e-3,x00(3),50,[1 1 1],'MarkerFaceAlpha',1,'MarkerFaceColor','flat',...
    'MarkerEdgeAlpha',0,...
    'MarkerEdgeColor','none');
hold off

set(figure1, 'Position', get(0,'Screensize')); % Maximize figure.
set(figure1,'color','none'),set(axes1,'color','none')

end

end
```
